# Expanded FLP toolbox for spatiotemporal protein degradation and transcriptomic profiling in *C. elegans*

**DOI:** 10.1101/2021.12.21.473632

**Authors:** Adrián Fragoso-Luna, Raquel Romero-Bueno, Michael Eibl, Cristina Ayuso, Celia Muñoz-Jiménez, Vladimir Benes, Ildefonso Cases, Peter Askjaer

**Author notes:** These authors contributed equally (Adrián Fragoso-Luna, Raquel Romero-Bueno, Michael Eibl).

## Abstract

Control of gene expression in specific tissues and/or at certain stages of development allows the study and manipulation of gene function with high precision. Site-specific genome recombination by the Flippase (FLP) and Cre enzymes has proven particularly relevant. Joint efforts of many research groups have led to the creation of efficient FLP and Cre drivers to regulate gene expression in a variety of tissues in *Caenorhabditis elegans*. Here, we extend this toolkit by the addition of FLP lines that drive recombination specifically in distal tip cells, the somatic gonad, coelomocytes and the epithelial P lineage. In some cases, recombination-mediated gene knockouts do not completely deplete protein levels due to persistence of long-lived proteins. To overcome this, we developed a spatiotemporally regulated degradation system for GFP fusion proteins (GFPdeg) based on FLP-mediated recombination. Using two stable nuclear pore proteins, MEL-28/ELYS and NPP-2/NUP85 as examples, we report the benefit of combining tissue-specific gene knockout and protein degradation to achieve complete protein depletion. We also demonstrate that FLP-mediated recombination can be utilized to identify transcriptomes in a *C. elegans* tissue of interest. We have adapted RNA polymerase DamID (RAPID) for the FLP toolbox and by focusing on a well-characterized tissue, the hypodermis, we show that the vast majority of genes identified by RAPID are known to be expressed in this tissue. These tools allow combining FLP activity for simultaneous gene inactivation and transcriptomic profiling, thus enabling the inquiry of gene function in various complex biological processes.

## Introduction

The nematode *C. elegans* is a popular model organism for genetic and biomedical research. A variety of methods have been developed to perturb gene expression and protein abundance in specific tissues and/or at particular developmental stages of *C. elegans*. These include tissue-specific RNA interference (RNAi) (Watts *et al*. 2020), conditional protein degradation (Zhang *et al*. 2015; Wang *et al*. 2017; Hills-Muckey *et al*. 2021) and site-specific recombinases (Nance and Frokjaer-Jensen 2019; Driesschaert *et al*. 2021). Especially Cyclization recombination (Cre) and Flippase (FLP) enzymes are popular tools for the control of gene expression in a spatiotemporal manner (Hubbard 2014; Driesschaert *et al*. 2021). While Cre was derived from the P1 bacteriophage (Sauer 1987), FLP originates from *Saccharomyces cerevisiae* (Senecoff *et al*. 1985). The Cre and FLP recombinases recognize 34 bp sequences, that are called loxP (locus of crossing-over in P1) and FRT (FLP recognition target), respectively. DNA located between two target sequences is either excised or inverted depending on their orientation. Both systems have been used efficiently in *C. elegans*, for example to induce gene knockout (KO) (Ruijtenberg and van den Heuvel 2015; Konietzka *et al*. 2020), control ectopic expression (Schmitt *et al*. 2012), ablation of neurotransmission (Davis *et al*. 2008), determination of gene expression (Gomez-Saldivar *et al*. 2020), mapping of chromatin organization (Cabianca *et al*. 2019; Harr *et al*. 2020) and transgenesis (Nonet 2020). Moreover, by utilizing both Cre and FLP, further versatility and cell specificity can be achieved (Ge *et al*. 2020; Nonet 2020). Thanks to community efforts, a growing number of characterized and stable Cre and FLP drivers are available, which facilitates the development and implementation of recombination-based tools (Voutev and Hubbard 2008; Kage-Nakadai *et al*. 2014; Ruijtenberg and van den Heuvel 2015; Munoz-Jimenez *et al*. 2017; Macias-Leon and Askjaer 2018; Ayuso and Askjaer 2019). Most recently, a light-inducible Cre recombinase has been added to the toolbox (Davis *et al*. 2021).

In some cases, the study of gene function at specific developmental time points can be compromised by remaining protein levels after gene perturbation. At a time point of interest, remaining protein might occlude the true phenotype associated with the gene’s loss of function by RNAi or conditional KO. This may be particularly relevant for long-lived proteins, such as histones and nuclear pore complex (NPC) proteins, that sometimes show lifelong persistence (Savas *et al*. 2012; Toyama *et al*. 2013). Besides, re-initiation of translation in the case of frameshift-mediated KO appears to be more prevalent than previously anticipated (Cohen *et al*. 2019). The combination of gene KO methods with conditional protein degradation may help to overcome this hurdle and completely deplete protein levels.

Tools for targeted protein degradation are constantly being developed and adapted for *C. elegans* and other model organisms (Nance and Frokjaer-Jensen 2019). Auxin-inducible degradation (AID) of proteins tagged with a 44 amino acid AID degron offers both temporal and spatial control through exposure to auxin and expression of *Arabidopsis thaliana* TIR1 protein, respectively (Zhang *et al*. 2015). Wang et al. (Wang *et al*. 2017) developed a system for the degradation of green fluorescent protein (GFP)-tagged proteins in *C. elegans* building on previous protein depletion systems (Caussinus *et al*. 2011; Armenti *et al*. 2014). In this GFP degradation (GFPdeg) system, a camelid single-domain antibody fragment that specifically binds GFP (“GFP nanobody”) is fused to the SOCS-box adaptor protein ZIF-1, that recruits CUL-2/Cullin-2, an E3 ligase. This leads to ubiquitination and proteasomal degradation of the GFP-tagged protein. An advantage of GFPdeg is that it is immediately applicable to any existing line expressing a GFP-tagged protein, but it offers only spatial control in its current version. Degradation kinetics in the AID or GFPdeg systems depend on the synthesis and turnover rates of the target protein, which sometimes renders protein depletion incomplete. Therefore, using AID or GFPdeg on its own may also not be sufficient for the inquiry of some biological processes and could benefit from the combination with genetic tools, such as site-specific recombination.

Here, we report an extension of the FLP-based toolbox for *C. elegans* by four means. Firstly, we describe novel integrated single-copy FLP drivers that efficiently induce recombination in the somatic gonad, distal tip cells, the P lineage or coelomocytes. We also report an improved FLP driver for intestinal cells and demonstrate that equally high recombination efficiency can be achieved for FLP drivers on different autosomes. These drivers increase the number of cell types amenable for easy FLP-mediated spatiotemporal control of gene expression. Secondly, we adapt the GFPdeg system for FLP-based regulation, thereby enabling spatiotemporal control of GFP fusion protein degradation. We show that inducible GFPdeg is equally efficient as the original constitutive version. Thirdly, using two endogenously GFP-tagged nuclear pore proteins, MEL-28/ELYS and NPP-2/NUP85, as test cases, we provide evidence that combined FLP-mediated gene KO and protein degradation is rapid and highly efficient to deplete stable proteins from a specific tissue of interest. The possibility of using a single FLP driver to control both conditional gene KO and protein degradation will facilitate the inquiry of tissue-specific gene function in complex biological processes. Lastly, we present the application of FLP-controlled RNA polymerase DamID (RAPID) (Gomez-Saldivar *et al*. 2020) for identification of transcribed genes in the hypodermis. We propose that combined with the growing list of tissue-specific FLP and Cre drivers, RAPID is an efficient and attractive method to explore transcription profiles across tissues without the need of physical isolation of cells.

## Material and Methods

### Plasmids

Promoter fragments for FLP expression constructs were PCR-amplified with *Pfu* polymerase using wild type *C. elegans* genomic DNA as template and inserted as *Not*I/*Kpn*I fragments into *Not*I/*Kpn*I of plasmid pBN338 (Munoz-Jimenez *et al*. 2017). Details are in Supplementary Tables S2 and S3. Exceptions were (1) pBN527 *gpa-14p::FLP::SL2::mNG* that was generated by first combining a 2.4 kb *Not*I/*Cla*I restriction fragment from pNP259 (Schmitt *et al*. 2012) with a 0.6 kb *Cla*I/*Kpn*I PCR fragment followed by insertion into *Not*I/*Kpn*I of pBN338 and (2) pBN487 *unc-122p::FLP::SL2::mNG* that was generated by inserting a *Not*I/*BsrG*I PCR fragment into *Not*I/*Acc65*I of pBN338. For generation of pBN532 *hlh-12p::FLP::SL2::mTagBFP2*, an intergenic *gpd-2/-3* sequence was amplified with primers B1012 and B1364 (see Supplementary Table S1) using pBN338 as template. A sequence encoding mTag2BFP2 was amplified with B1365 and B1366 using pMINIT mTagBFP2 (Porta-de-la-Riva *et al*. 2021) as template. The two PCR fragments were PCR-stitched with primers B1012 and B1366 and inserted by NEBuilder HiFi DNA Assembly (New England Biolabs) into *Mlu*I of pBN523. To generate plasmids pBN548 *dat-1p::FLP::SL2::mNG* and pBN549 *hlh-12p::FLP::SL2::mTagBFP2* for integration into universal MosSCI landing sites (Frokjaer-Jensen *et al*. 2014), *Xma*I/*Not*I fragments from pBN338 and pBN532, respectively, were inserted into *Ngo*MIV/*Not*I of pBN8 (Rodenas *et al*. 2012).

Targeting plasmids for SapTrap CRISPR/Cas9 gene modification were obtained as described (Schwartz and Jorgensen 2016). Details for construction of plasmids pBN309 *mCh::baf-1*, pBN433 *g>f>p::npp-2* (“>” denotes an FRT site) and pBN493 *gf>p::baf-1* are provided in Supplementary Tables S1 and S3. The plasmids direct the insertion of fluorescent tags immediately downstream of the endogenous start codon. Plasmid pBN351 for insertion of *g>f>p* immediately upstream of the endogenous *mel-28* start codon was generated from plasmids pMLS256, pMLS288 and pBN312 (Schwartz and Jorgensen 2016; Munoz-Jimenez *et al*. 2017), sgRNA oligonucleotides B1050 and B1051, 5’ homology arm (499 bp; amplified with B1042 and B1043 using plasmid #1397 (Gomez-Saldivar *et al*. 2016) as template) and 3’ homology arm (258 bp; amplified with B1044 and B1045 using plasmid #1397 as template) (see Supplementary Tables S1 and S3). Donor plasmid pBN477 containing FRT in the second *gfp* intron was constructed by inserting a 574 bp *Xho*I/*Hind*III fragment from pBN312 into *Xho*I/*Hind*III of pMLS252 (Schwartz and Jorgensen 2016).

Plasmid pBN488 *hsp16.41p>mCh::his-58>vhhGFP4::zif-1* for targeted degradation of GFP fusion proteins was constructed by amplifying *vhhGFP4::zif-1* with primers B1333 and B1334 using plasmid pNH102 (a kind gift from Ngang Heok Tang and Andrew Chisholm) as template. The PCR product was digested with *Xho*I and *Nhe*I and inserted into plasmid pBN209 *hsp16.41p>mCh::his-58>dam::emr-1* (Cabianca *et al*. 2019) digested with the same enzymes to replace *dam::emr-1*.

Plasmid pBN537 *hsp16.41p>mCh::his-58>dam::rpb-6* for RAPID was constructed by amplifying *rpb-6* with primers B1456 and B1457 using genomic DNA as template. The PCR product was digested with *Ngo*MIV and *Nhe*I and inserted into plasmid pBN209 digested with the same enzymes to replace *emr-1*.

### Nematode culture, strains and genome engineering

Worm strains were maintained on Nematode Growth Medium (NGM) petri dishes seeded with *Escherichia coli* OP50 (Stiernagle 2006). Unless otherwise noted, strains were maintained at 16°C. The N2 strain was used as wild type reference. Other strains are listed in Supplementary Table S4.

Single-copy insertion of FLP expression constructs into the *cxTi10882* locus on chrIV or the *oxTi365* locus on chrV was done by microinjection into the gonads of EG6700 or EG8082 young adult hermaphrodites (Frokjaer-Jensen *et al*. 2008; Frokjaer-Jensen *et al*. 2014; Dobrzynska *et al*. 2016). Plasmids carrying FLP and *unc-119(+)* as selection marker were injected at 50 ng/μl together with pCFJ601 *eft-3p::Mos1* ((Frokjaer-Jensen *et al*. 2012), 50 ng/μl), pBN1 *lmn-1p::mCh::his-58* ((Rodenas *et al*. 2012), 10 ng/μl), pCFJ90 *myo-2p::mCh* ((Frokjaer-Jensen *et al*. 2008), 2.5 ng/μl) and pCFJ104 *myo-3p::mCh* ((Frokjaer-Jensen *et al*. 2008), 5 ng/μl). Integration of the FLP expression constructs was confirmed by detection of fluorescent mNeonGreen (mNG) or mTagBFP2 in the target tissue of wild-type moving animals without ectopic mCherry expression from the co-injection markers. For FLP constructs without fluorescent co-expression markers, correct expression pattern was confirmed after crossing with the dual-color FRT reporter (see Confocal Microscopy below).

Similarly, single-copy insertion of vhhGFP4::ZIF-1 and Dam::RPB-6 expression constructs into the *ttTi5605* locus on chrII was done by microinjection into the gonads of EG4322 young adult hermaphrodites. Injection markers and transposase plasmid were injected at same concentrations as above. Integrity of the integrated transgenes was confirmed by PCR, and, after crossing to FLP drivers, functional assays (degradation of GFP fusion proteins for the GFPdeg system; see Confocal Microscopy below).

CRISPR/Cas9-mediated genome modification to generate strains BN581 *baf-1(bq13[mCh::baf-1])* III, BN1164 *baf-1(bq47[gf>p::baf-1])* Ill, BN902 *mel-28(bq17[g>f>p::mel-28])* III and BN1044 *npp-2(bq38[g>f>p::npp-2])* I was performed by microinjection of targeting plasmids pBN309, pBN493, pBN351 and pBN433, respectively, into the gonads of HT1593 *unc-119(ed3)* III young adult hermaphrodites (Schwartz and Jorgensen 2016). Targeting plasmids were injected at 65 ng/μl together with plasmids #1286 *eft-3p::Cas9* ((Friedland *et al*. 2013), 25 ng/μl), pBN1 (10 ng/μl), pCFJ90 (2.5 ng/μl) and pCFJ104 (5 ng/μl). Successful modification of the target loci was confirmed by detection of fluorescent fusion proteins in the nuclear envelope throughout the body of wild-type moving animals without ectopic mCherry expression from the co-injection markers. The *unc-119(+)* selection marker was next excised by microinjection with Cre expression plasmid pMLS328 (Schwartz and Jorgensen 2016). Finally, uncoordinated hermaphrodites were outcrossed to N2 males to remove the *unc-119(ed3)* allele.

Insertion of an FRT in the *baf-1* 3’-UTR was performed by microinjection of BN1164 *baf-1(bq47[gf>p::baf-1])* young adult hermaphrodites with a mixture containing 10 μM *baf-1* crRNA:trRNA duplex (B1312+B1067), 2.5 μM *dpy-10* crRNA:trRNA duplex (B1068+B1067), 3 μM recombinant Cas9, 4 μM *baf-1* ssDNA repair template (B1313) and 1 μM *dpy-10* ssDNA repair template (B725). Recombinant Cas9 was produced by the CABD Proteomics and Biochemistry Facility. All DNA and RNA oligonucleotides were purchased from IDT.

### Confocal microscopy

To induce expression of either the dual-color FRT reporter or the GFPdeg system, NGM plates with nematodes were incubated for 15 min floating in a 34°C water bath. For evaluation of recombination efficiency and specificity, nematodes were left to recover at 20°C for 2-3 hours prior to imaging. For GFP degradation experiments, recovery was at 25°C for 1-24 hours as indicated for each experiment. For imaging, nematodes were anesthetized in 5 μl 10 mM levamisole HCl on 2% agarose pads and covered by a cover slip. Images were acquired with a Nikon Eclipse Ti microscope equipped with Plan Fluor 40×/1.3 and Plan Apo VC 60×/1.4 objectives and an A1R scanner using a pinhole of 1.2 airy units (Munoz-Jimenez *et al*. 2017). Cells with GFP signal as an indication of a recombination event in the dual-color reporter as well as mNG co-expression with FLP were evaluated from z-stacks acquired with the Nikon NIS software. To analyze the reduction of protein levels of GFP fusion proteins after FLP recombination and/or the induction of the GFP degradation system, specific regions and focal planes were localized to acquire reproducible sets of images. These are indicated in the text.

Fluorescence intensity was quantified in Fiji/ImageJ 2.1.0/1.53c (Schindelin *et al*. 2012) by tracing and measuring the signal at the nuclear envelope with a freehand line of width 3. After subtracting the background measured in animals without expression of GFP, the data were analyzed in R Studio (1.3.1093 (Team 2020a); running R 4.0.2 (Team 2020b)). Data were visualized either with base plotting or using ggplot2 (Wickham 2016), mainly within the ggpubr framework (version 0.4.0 (Kassambara 2020)). For statistical analysis, Shapiro-Wilk test of normality and Levene’s test for homogeneity of variance across groups were performed on the data sets to determine if either ANOVA and pair-wise t tests or Kruskal-Wallis rank sum test combined with Dunn’s test for pairwise multiple comparisons of the ranked data were most suitable. In all cases, p values were adjusted for multiple comparisons (Benjamini & Hochberg method).

### Brood size experiments

BN580 *g>f>p::baf-1*, BN1182 *mCh::baf-1/g>f>p::baf-1; nhx-2p::FLP* and BN1223 *mCh::baf-1/gf>p::baf-1>; nhx-2p::FLP* worms were cultured at 16°C until L4 larval stage. For BN1182 and BN1223, homozygous *mCh::baf-1, g>f>p::baf-1* and *gf>p::baf-1>* animals were selected under a fluorescence stereoscope. In each replica, 8-10 L4 larvae per strain were transferred to individual NGM plates and incubated at 20°C. During the reproductive period, adults were transferred to new plates each day and the numbers of progenies were recorded. Adults were censored if they desiccated on the walls of the plates or were damaged during transfer. Statistical analysis was done in R using Wilcoxon rank sum tests and adjusted for multiple comparisons (Benjamini & Hochberg method).

### Lifespan experiments

Worms were cultured at 16°C until L4 larval stage. In each replica, 20 L4 larvae per strain were transferred to newly seeded NGM plates and incubated for 15 min floating in a 34°C water bath to induce the GFPdeg system. They were then continuously incubated at 25°C. During the reproductive period, the live adults were transferred to new plates each day to not confuse founders with their progeny. Throughout the experiment, the number of live and dead animals were recorded. Animals were censored if they disappeared, were damaged during transfer or died due to internal hatching. Statistical analysis was done in R using the survival package (version 3.2-7 (Therneau 2020)), which calculates Chi-square statistics based on a log-rank test.

### RAPID

Worms were grown at 20°C for at least two generations on NGM plates seeded with *E. coli* GM119 *dam*-bacteria. Eggs were obtained by sodium hypochlorite treatment (Stiernagle 2006) and incubated in M9 containing 0.01% Tween-20 (M9-T) overnight at 20°C without food. For each strain and replica, four 100 mm NGM plates with a lawn of GM119 were seeded each with approximately 1000 synchronized L1 larvae and incubated at 20°C for 48 h until reaching L4 stage. Larvae were washed off the plates with M9-T, collected in 15 ml tubes and washed 8 times in M9-T. Next, worms were transferred to 1.5 ml tubes washed twice in M9-T. The supernatant was aspired to leave ~30 μl and tubes were snap frozen in liquid nitrogen and stored at −80°C until used.

To purify genomic DNA, samples were lysed in 5 freeze/thaw cycles (1 min in liquid nitrogen; 3 min in 37°C shaker) and processed with DNeasy Blood and Tissue Kit (QIAGEN #69504). Two hundred ng of genomic DNA was used as input in the DamID protocol that included an improved pool of AdR primers as described (de la Cruz Ruiz *et al*. 2022). Dam-methylated fragments were amplified with 14 PCR cycles and quality checked on an Agilent Bioanalyzer 2100. Next, libraries were prepared using 50 ng of PCR-amplified fragments and NEBNext® Ultra™ II DNA Library Prep Kit for Illumina (E7645S). Library amplicons size range distribution was checked as before, pooled and re-purified with 1x volume SpeedBeads (de la Cruz Ruiz *et al*. 2022) followed by sequencing at EMBL GeneCore using a NextSeq 500 (Illumina).

Sequencing fastq files were filtered to discard reads that do not start with the DamID adapter followed by GATC (“CGCGGCCGAG*GATC*”) using R Studio and the damid.seq.R pipeline version 0.1.3 (available at github.com/damidseq/RDamIDSeq) (Sharma *et al*. 2016). Retained fragments were mapped to *C. elegans* UCSC genome version ce11 with the same pipeline, which produced 9.5-41.9 million uniquely mapped reads per sample. Normalization of Dam::RPB-6 reads to GFP::Dam reads was performed using the damidseq_pipeline package (https://owenjm.github.io/damidseq_pipeline/) (Marshall and Brand 2015) on the bam files generated by the damid.seq.R script. Gene occupancy was computed as log2(Dam::RPB-6/GFP::Dam) with polii.gene.call (https://github.com/owenjm/polii.gene.call) using bedgraph output files from the damidseq_pipeline and WBCel235 gene annotation. A false discovery rate (FDR) of 0.05 was used as threshold to call transcribed genes. Average profile plots and heatmap were generated with SeqPlots (Version 3.0.12) using bedgraph output files from the damidseq_pipeline and WBCel235 gene annotation; H3K36me3 signal along either all protein coding genes or top20 transcribed genes on chromosome I (included in the software) was used as comparison (Stempor and Ahringer 2016). Genome browser views and Venn diagrams were generated with Integrative Genomics Viewer (IGV Web App) (Thorvaldsdottir *et al*. 2013) and jvenn (Web App) (Bardou *et al*. 2014), respectively. Statistical significance of overlap between protein coding gene lists was calculated with Fisher exact test for independence in 2×2 contingency tables using R Studio and a total count of protein coding genes of 19,975.

### Data availability

See Table S1, Table S2, Table S3 and Table S4 for complete lists of primers, promoters, plasmids and strains, respectively. Representative strains will be made immediately available through the *Caenorhabditis* Genetics Center; plasmids and other strains can be requested from the authors. Tables S5-S7 contain lists of hypodermal genes identified by RAPID (Table S5), their overlap with other hypodermal gene lists (Table S6) and their expression level in single-cell RNAseq (Table S7). Raw and processed RAPID files are deposited at Gene Expression Omnibus accession GSE214300 (https://www.ncbi.nlm.nih.gov/geo/).

## Results

### Characterization of new FLP lines

Previously, a series of efficient tissue-specific FLP-lines were generated by Mos1-mediated Single Copy Insertion (MosSCI) (Munoz-Jimenez *et al*. 2017; Macias-Leon and Askjaer 2018; Ayuso and Askjaer 2019; Van de Walle *et al*. 2020). To expand the existing toolkit, we created additional MosSCI lines that express FLP in specific cell types and evaluated recombination efficiency with the heat-inducible dual color reporter *bqSi294* that produces green nuclei (GFP::HIS-58) upon successful recombination and red nuclei (mCherry::HIS-58) elsewhere (Figure 1A) (Munoz-Jimenez *et al*. 2017). In most constructs (see below), an SL2 trans-splicing sequence followed by the gene encoding the fluorescent protein mNeonGreen (mNG) was inserted downstream of FLP (FLP::SL2::mNG), enabling visualization of FLP expression by co-expression of mNG (Munoz-Jimenez *et al*. 2017).

**Figure 1.**
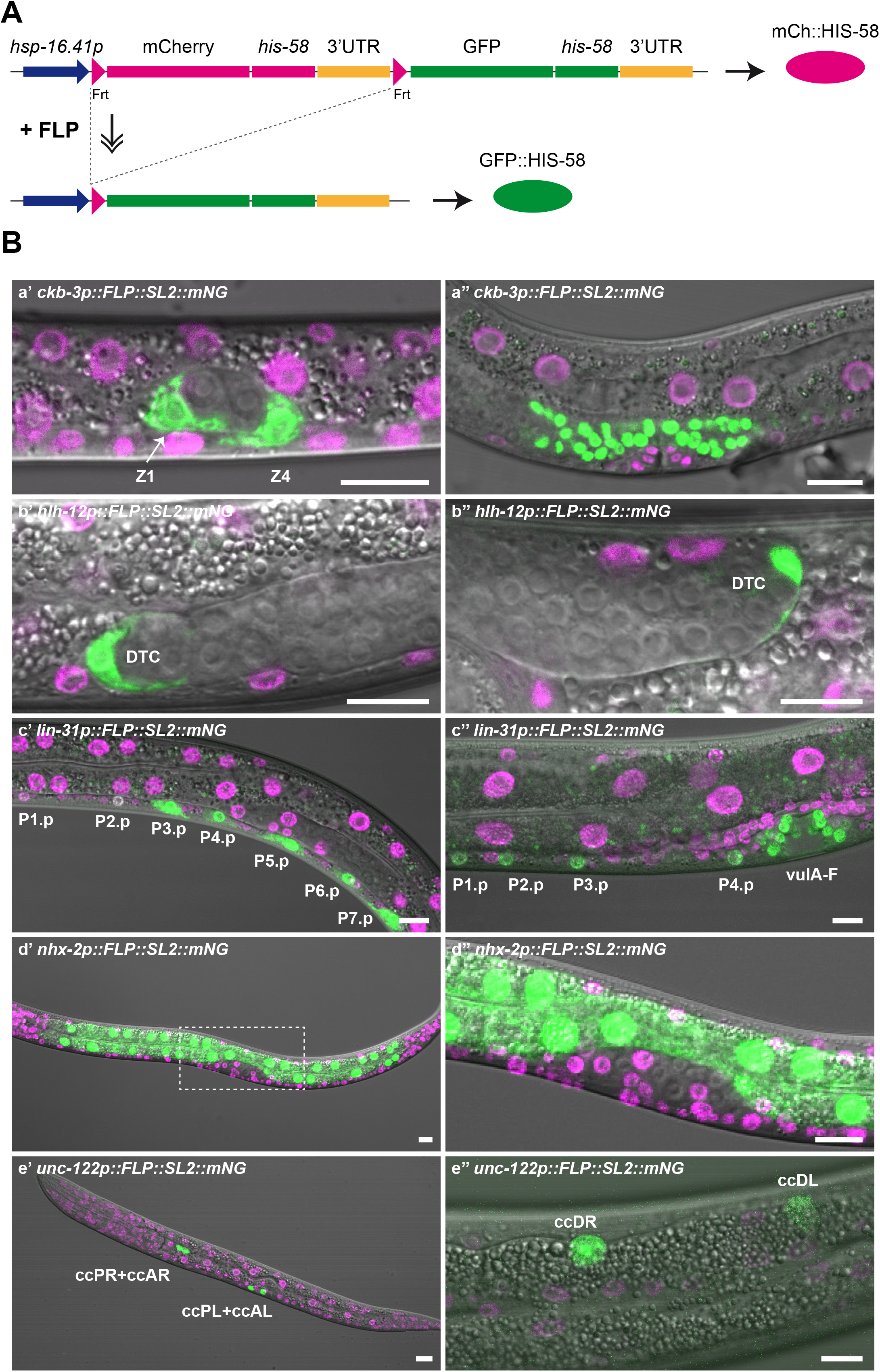
Characterization of novel FLP lines. (A) Schematic representation of the dual color reporter used to evaluate recombination activity. Upon temperature induction of the *hsp-16.41* promoter, the reporter expresses red (mCh) or green (GFP) histone HIS-58 in cell lineages without or with FLP-mediated recombination, respectively. (B) Evaluation of FLP lines crossed with the dual color reporter. Although the FLP lines co-express diffuse mNG in the cytoplasm, GFP::HIS-58 is clearly visible in the expected nuclei. (Ba) *ckb-3p::FLP::SL2::mNG*: maximum projections of 2-3 confocal sections of BN854 L1 larva (Ba’) and larva at L3/L4 molt (Ba’’). The Z1 and Z4 somatic gonad precursor cells are indicated. (Bb) *hlh-12p::FLP::SL2::mNG*: single confocal sections of BN1204 L2 larva (Bb’) and young adult (Bb’’). The distal tip cells (DTCs) are indicated. (Bc)*lin-31p::FLP::SL2::mNG*: maximum projections of three confocal sections of BN1023 L2 larva (Bc’) and L4 larva (Bc’’). The posterior daughters of P1-P7 and vulA-vulF forming the vulva are indicated. (Bd) *nhx-2p::FLP::SL2::mNG*: maximum projection of five confocal sections of BN999 L1 larva (Bd’; indicated area shown enlarged in Bd’’). (Be) *unc-122p::FLP::SL2::mNG*: maximum projections of four confocal sections of BN1029 L1 larva (Be’) and L4 larva (Be’’). The three pairs of coelomocytes are indicated. Scale bars 10 μm.

We first generated a line that expresses FLP under control of the *ckb-3* promoter, which is active in the Z1 and Z4 cells that compose the somatic gonadal primordium of the hermaphrodite in early L1 larvae (Kroetz and Zarkower 2015). When crossed with the dual color reporter, we detected recombination in all 36 L1 larvae analyzed; in 35 larvae both Z1 and Z4 expressed GFP::HIS-58, and one larva expressed GFP::HIS-58 in a single cell (Figure 1B; Table 1). The majority of Z1/Z4 cells expressed GFP::HIS-58 only, indicating that recombination had taken place at both alleles (67/72 cells = 93%), whereas both green and red histones were observed in 4 cells. Of note, there were never more than the expected two green nuclei at this developmental stage (early L1), which highlights the tissue specificity of the *ckb-3p::FLP* line. The high efficiency of the line in the somatic gonadal primordium implies that it is very suitable for manipulation of all five tissues of the reproductive system: distal tip cells, gonadal sheath, spermatheca, spermatheca-uterine valve and the uterus (Figure 1B) (Hubbard and Greenstein 2005).

**Table 1.** Efficiency of FLP-mediated recombination.

| Promoter<sup>a</sup> | Cell type | Stage<sup>b</sup> | Observed<sup>c</sup> | Expected | Efficiency |
| --- | --- | --- | --- | --- | --- |
| *ckb-3* | Somatic gonad | Early L1 | 2.0±0.2 (36) | 2 | 99% |
| *dat-1* | Dopaminergic neurons | L3-L4 | 8.0±0.0 (11) | 8 | 100% |
| *hlh-12* | Distal tip cell | L2-adult | 2.0±0.0 (24; chrIV<sup>a</sup>)<br>2.0±0.0 (10; chrV<sup>a</sup>) | 2 | 100% |
| *lin-31* | P1.p-P11.p (P5.p-P7p) | L2-L4 | 3.0±0.0 (14) | 3<sup>d</sup> | 100% |
|  |  | L3 | 12.0±0.0 (13) | 12<sup>d</sup> | 100% |
|  |  | L4 | 22.0±0.0 (3) | 22<sup>d</sup> | 100% |
| *nhx-2* | Intestine | L1 | 20.0±0.0 (10) | 20 (19-22) | 100% |
|  |  | L2 | 31.9±0.6 (17) | 32 (19-22) | 100% |
| *unc-122* | Coelomocytes | L1 | 4.0±0.0 (8) | 4 | 100% |
|  |  | L3-L4 | 6.0±0.0 (23) | 6 | 100% |

<sup>a</sup> Single-copy FLP constructs inserted on chrIV except *dat-1p::FLP* and one of the *hlh-12p::FLP* constructs that were inserted on chrV.
<sup>b</sup> Developmental stage that was used for determination of recombination efficiency.
<sup>c</sup> Average number of GFP positive nuclei ± SD; number in brackets: *n*.
<sup>d</sup> Efficiencies were calculated based on recombination in central Pn.p cells and their descendants.

The distal tip cells (DTCs) play important roles during development as the leading cells that guide migration of the gonad arms in hermaphrodites and also in control of germ cell proliferation (Hubbard and Greenstein 2005). To express FLP specifically in DTCs, we used the *hlh-12* promoter which is expressed from L2 stage (Tamai and Nishiwaki 2007). We observed recombination in both DTCs in all hermaphrodites analyzed (*n*=24, L2-adult stage; Figure 1B; Table 1). Except for one DTC in a young L2 larva, recombination had taken place at both alleles (47/48 cells = 98%) and was not observed in other cell types, demonstrating the high efficiency and specificity of the *hlh-12p::FLP* line.

We then generated a *lin-31p::FLP* line to enable recombination in the vulval precursor cells (VPCs). LIN-31 is expressed in P1.p-P11.p cells from the middle of L1 stage, when these cells are born (Tan *et al*. 1998). We focused our attention on the central VPCs in L2 larvae and on VPC descendants during vulval morphogenesis in L3 and L4 larvae. Importantly, recombination of both alleles was observed in all VPCs and VPC descendants in all animals (*n*=30, L2-adult stage; Figure 1B; Table 1). We also observed GFP::HIS-58 expression in a few head neurons, which is in agreement with a previous report on *lin-31* expression (Supplementary Figure S1A)(Reece-Hoyes *et al*. 2007). Finally, we noted weak GFP::HIS-58 signal in hyp7 nuclei in late L3-L4 larvae, which we assume originates from fusion of descendants of P3.p, P4.p and P8.p with the hyp7 syncytium (Supplementary Figure S1A).

In the first version of our FLP toolkit, we used the *elt-2* promoter to drive recombination in the intestine with ~100% efficiency (Munoz-Jimenez *et al*. 2017). However, because recombination was also detected in the somatic gonad, we generated an alternative line based on the *nhx-2* promoter (Nehrke and Melvin 2002). In young L1 larvae, the number of intestinal nuclei ranges from 19 to 22 and increases to 30-34 in L2 larvae (McGhee 2007). Recombination efficiency reached 98% based on green-only nuclei and 100% when also including nuclei with expression of both green and red histones (*n*=27, Figure 1B; Table 1). This demonstrates high recombination efficiency, comparable to the *elt-2p::FLP* line. Importantly, we did not detect recombination outside the intestine with the *nhx-2p::FLP* line, so we recommend to use this line.

We next generated an *unc-122p::FLP* line to enable recombination in coelomocytes (CCs), which are specialized cells thought to be involved in the innate immune response and scavenging (Loria *et al*. 2004; Altun and Hall 2009). Two anterior CC pairs are born during embryogenesis whereas a posterior pair is born right before the L1/L2 molt. Consistently, in 8 of 8 L1s we observed 4 GFP::HIS-58 expressing CCs, and in 23 of 23 L3-L4 larvae 6 GFP::HIS-58 expressing CCs (Figure 1B; Table 1). Expression of mCh::HIS-58 was not detected in the CCs, indicating that both alleles had recombined, whereas GFP::HIS-58 expression was restricted to the CCs.

We also attempted to generate additional lines to target specific neuronal subtypes, including cholinergic neurons (using the *unc-17* promoter) and *gpa-14*-expressing neurons. However, although both lines expressed the marker mNG in the expected tissues, they also showed either recombination in unexpected cell types (for *unc-17p::FLP*) or inefficient recombination (for *gpa-14p::FLP*) and were not characterized in detail (Supplementary Figure S1B-C; see Discussion).

In the strains described above, co-expression of mNG serves to visualize when and where FLP is expressed. However, in some experiments, the signal from mNG might interfere with other fluorescent proteins. To circumvent this, we replaced mNG with mTagBFP2 that emits blue light. When placed under control of the *hlh-12* promoter, we observed mTagBFP2 expression and FLP-mediated recombination in both DTCs of all animals (*n*=12; Supplementary Figure S1D). The sequences encoding mNG and mTagBFP2 in the MosSCI plasmids are flanked by *Mlu*I restriction sites, which facilitates the exchange of the former for the latter in the creation of new strains.

All lines described so far involve FLP expression constructs on chromosome IV. To increase the versatility of the FLP toolkit, we subcloned *dat-1p::FLP* and *hlh-12p::FLP* constructs to *ttTi5605* targeting vectors which enable insertion into universal MosSCI sites on any autosome (Frokjaer-Jensen *et al*. 2014). After insertion of the FLP expression constructs on chromosome V we tested their efficiency on the dual-color reporter and found that they both induce specific recombination in all expected cells (Table 1). This suggests that other existing FLP constructs can easily be adapted for insertion into different loci without compromising their efficiency.

### Conditional gene knockout by GFP exon removal

We previously designed a conditional knockout (KO) cassette based on GFP containing FRT sites in introns 1 and 2 (Figure 2A) (Munoz-Jimenez *et al*. 2017). Excision of GFP exon 2 by FLP causes a frame shift and a premature termination codon, which most likely targets the mRNA for nonsense-mediated decay (Arribere *et al*. 2020). The premature termination codon precedes three short open reading frames that are out of frame with respect to the gene downstream of GFP. Together, this should completely prevent expression of the tagged protein. To test this experimentally, we compared the consequence of either complete removal of the *baf-1* gene (*gf>p::baf-1>;* “>” denotes the position of FRT; see Figure 2A) or removal of GFP exon 2 in an endogenously tagged *gfp::baf-1* line (*g>f>p::baf-1*). BAF-1 is an essential and ubiquitously expressed nuclear envelope protein (Gorjanacz *et al*. 2007). In both cases, we used the *nhx-2p::FLP* line described above to induce *baf-1* KO specifically in intestinal cells, which negatively affects health (Munoz-Jimenez *et al*. 2017). As controls we included (i) the *g>f>p::baf-1* line without FLP expression and (ii) an endogenously tagged *mCh::baf-1* line without FRT sequences but with FLP expression. While hermaphrodites of the two control lines produced an identical number of progeny (*g>f>p::baf-1* 299±25 [mean±SD], n=10; *mCh::baf-1* 301±55, n=15), fertility was reduced equally by 31% in the two conditional knockout lines (*g>f>p::baf-1; nhx-2p::FLP* 207±71, n=15; *gf>p::baf-1>; nhx-2p::FLP* 206±68, n=12)(Figure 2B). Based on this, we conclude that excision of the second exon of the GFP KO cassette causes an equally strong loss of function phenotype as complete gene removal. Moreover, it provides the advantage that insertion of the GFP KO cassette in the 5’-end of a target gene in a single step is easier than inserting both a GFP tag (with a single FRT) and a second FRT inside or downstream of the gene.

**Figure 2.**
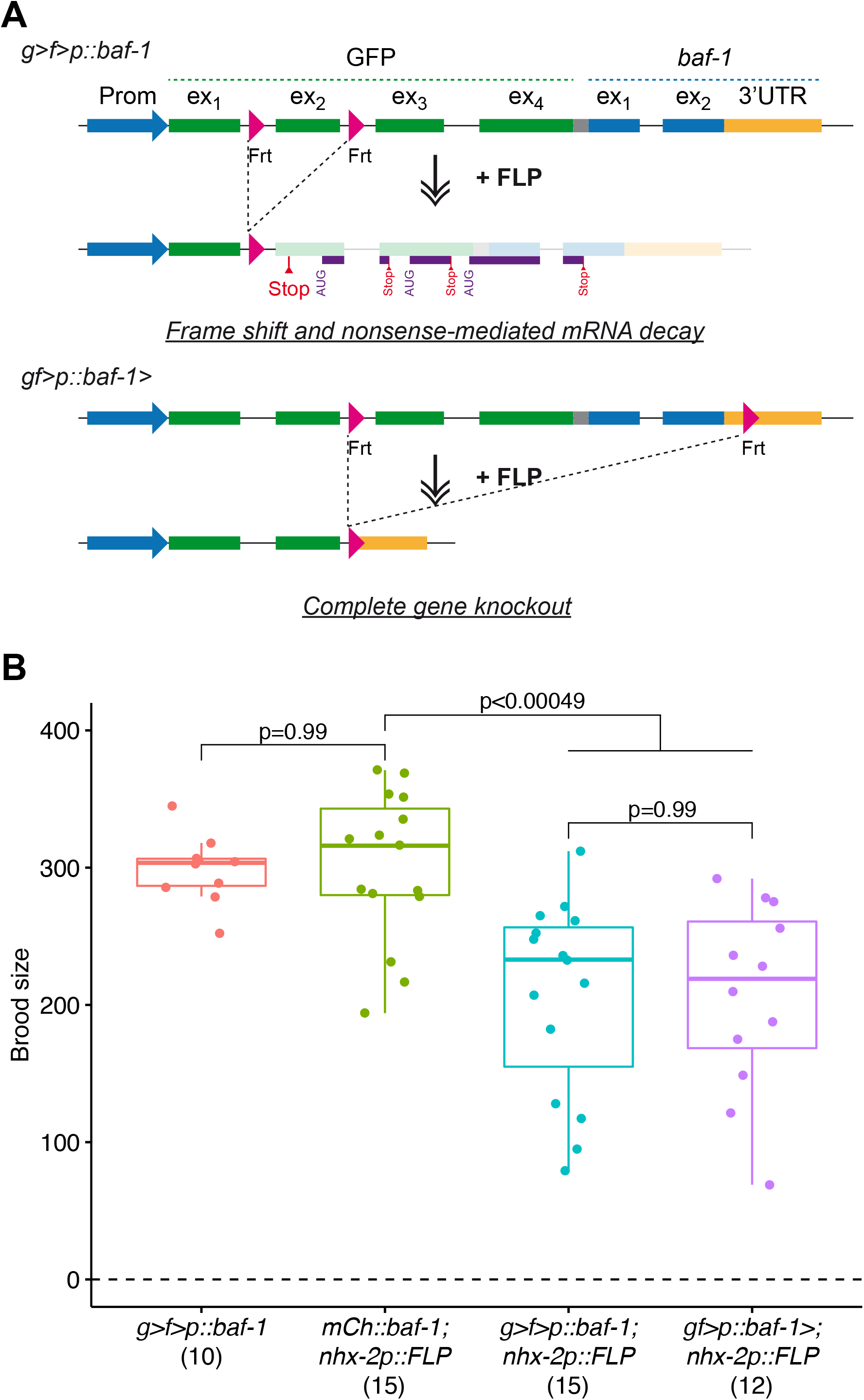
Comparison of GFP KO cassette and complete gene KO. (A) Schematic representation of the two KO strategies. In the first strategy (top), a GFP KO cassette with FRTs in *gfp* introns 1 and 2 is inserted immediately downstream of, and in frame with, the start codon of a gene of interest (here *baf-1*). FLP-mediated recombination leads to a frame shift and a premature termination codon (“Stop”). In case the ribosome remains associated with the mRNA, the three downstream open reading frames are all out-of-frame with *baf-1* (purple lines). In the second strategy (bottom), the GFP cassette contains a single FRT and a second FRT is inserted in the 3’-UTR of the gene of interest. FLP-mediated recombination leads to complete gene removal. (B) Brood sizes were determined for control animals (*g>f>p::baf-1* and *mCh::baf-1; nhx-2p::FLP*) and for animals with conditional KO of *baf-1* in the intestine (*g>f>p::baf-1; nhx-2p::FLP* [first strategy] and *gf>p::baf-1>; nhx-2p::FLP* [second strategy]; strains BN580, BN1182 and BN1223). Each point represents the number of progenies produced by an individual worm; *n* for each genotype is indicated in parentheses. P values from pair-wise t tests and adjusted for multiple comparisons (Benjamini & Hochberg method) are indicated. In these and subsequent boxplots, the midline represents the median and the upper and lower limits of the box correspond to the third and first quartile, respectively. The whiskers extend to the lowest and higher observations but maximum up to 1.5 times the interquartile range.

While conditional gene KO prevents synthesis of new transcripts, mRNA and protein molecules already existing in the cell might persist for days, weeks or even months (Toyama *et al*. 2013). This may obscure both investigation of gene function and potential therapeutic interventions. To address this, we evaluated the kinetics of GFP::MEL-28 depletion from hypodermal cells of *g>f>p::mel-28; dpy-7p::FLP* hermaphrodites. MEL-28 is a component of the long-lived NUP107 nuclear pore subcomplex and is required for DNA segregation and post-mitotic nuclear assembly (Fernandez and Piano 2006; Galy *et al*. 2006; Toyama *et al*. 2013). Although the *dpy-7* promoter is active already during embryogenesis to drive FLP expression, ~95% of GFP::MEL-28 was still present in L1 larvae as compared to the control *g>f>p::mel-28* line without FLP expression (Figure 3; compare Control and KO in 3B). In contrast, in L2 larvae and young adults a dramatic reduction of the median GFP::MEL-28 signal to ~12% and ~3%, respectively, was observed specifically in the hypodermis (Figure 3). This demonstrates the potential of conditional KO methods to evaluate the function of long-lived proteins in late larvae and adults (see also below), whereas they might be less suitable in early larvae.

**Figure 3.**
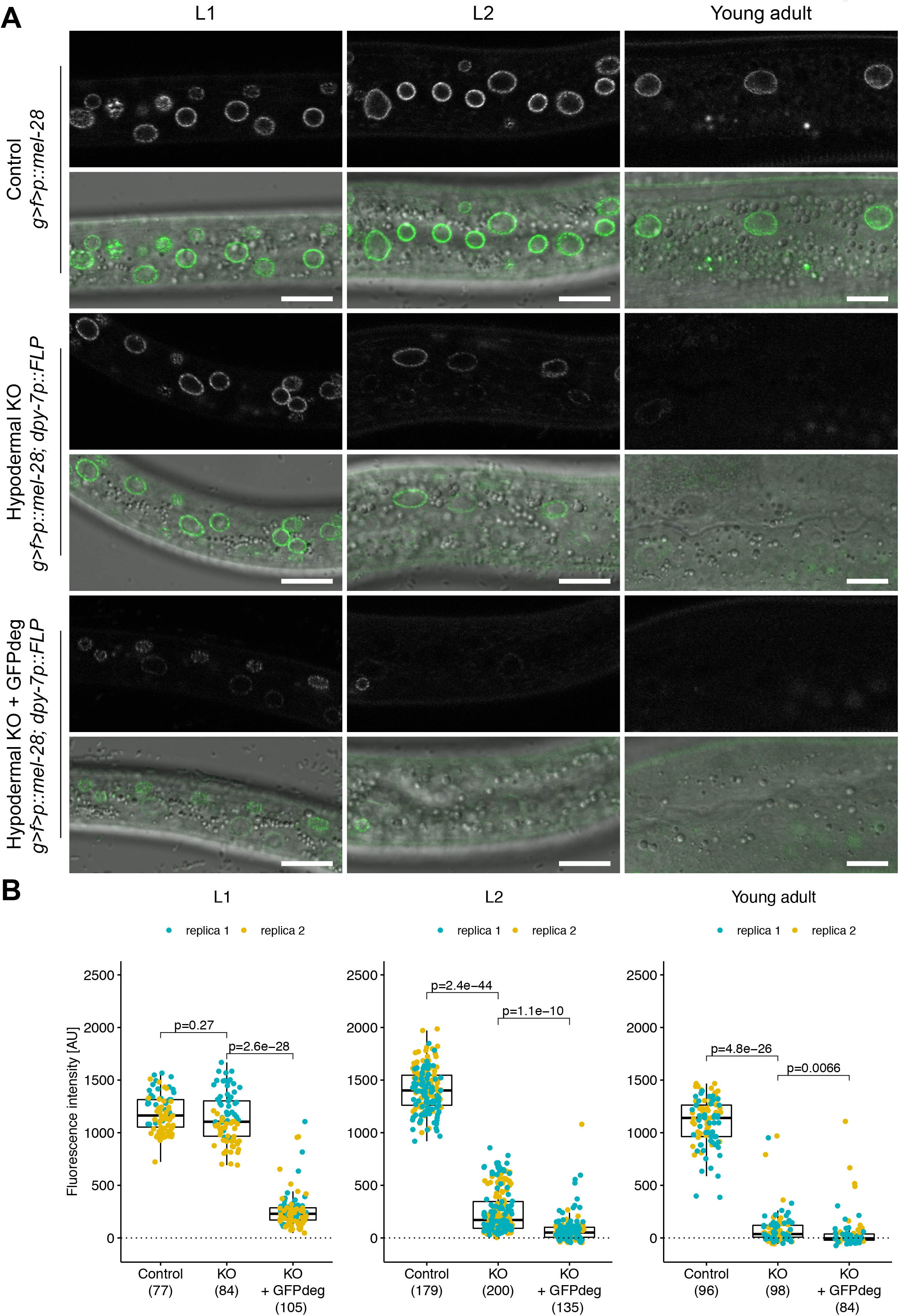
KO of GFP::MEL-28 in hypodermal nuclei. (A) Confocal micrographs of *g>f>p::mel-28* L1 larvae, L2 larvae and young adults either not expressing FLP (top panels; strain BN793) or expressing FLP in the hypodermis either without (middle panels; *dpy-7p::FLP* driver; strain BN1121) or with GFPdeg in the hypodermis (bottom panels; *dpy-7p::FLP* driver; strain BN1122). A single focal plane of the hypodermis is shown. Scale bar 10 μm. (B) Quantification of GFP::MEL-28 at the nuclear envelope of hypodermal nuclei. Each point represents a single nucleus. Data were acquired from at least 10 animals in two independent experiments; *n* for each genotype is indicated in parentheses. P values from Kruskal-Wallis rank sum tests and adjusted with Dunn’s test for multiple comparisons (Benjamini & Hochberg method) are indicated.

### FLP-mediated protein degradation

To enable analysis of long-lived proteins during development, we developed a system for FLP-mediated, inducible and rapid protein degradation from specific cells. A potent system for constitutive tissue-specific degradation of GFP-tagged proteins based on expression of vhhGFP4::ZIF-1, a GFP-binding nanobody fused to ZIF-1, was recently developed in *C. elegans* (Wang *et al*. 2017). To achieve both temporal and spatial control of degradation of GFP-tagged proteins (hereafter termed GFPdeg), we replaced the *gfp::his-58* sequence of the dual color reporter with a sequence encoding vhhGFP4::ZIF-1. In this design, tissue-specificity is controlled by the choice of a particular FLP-expressing line, whereas temporal control is provided by the heat-inducible *hsp-16.41* promoter (Figure 4A).

**Figure 4.**
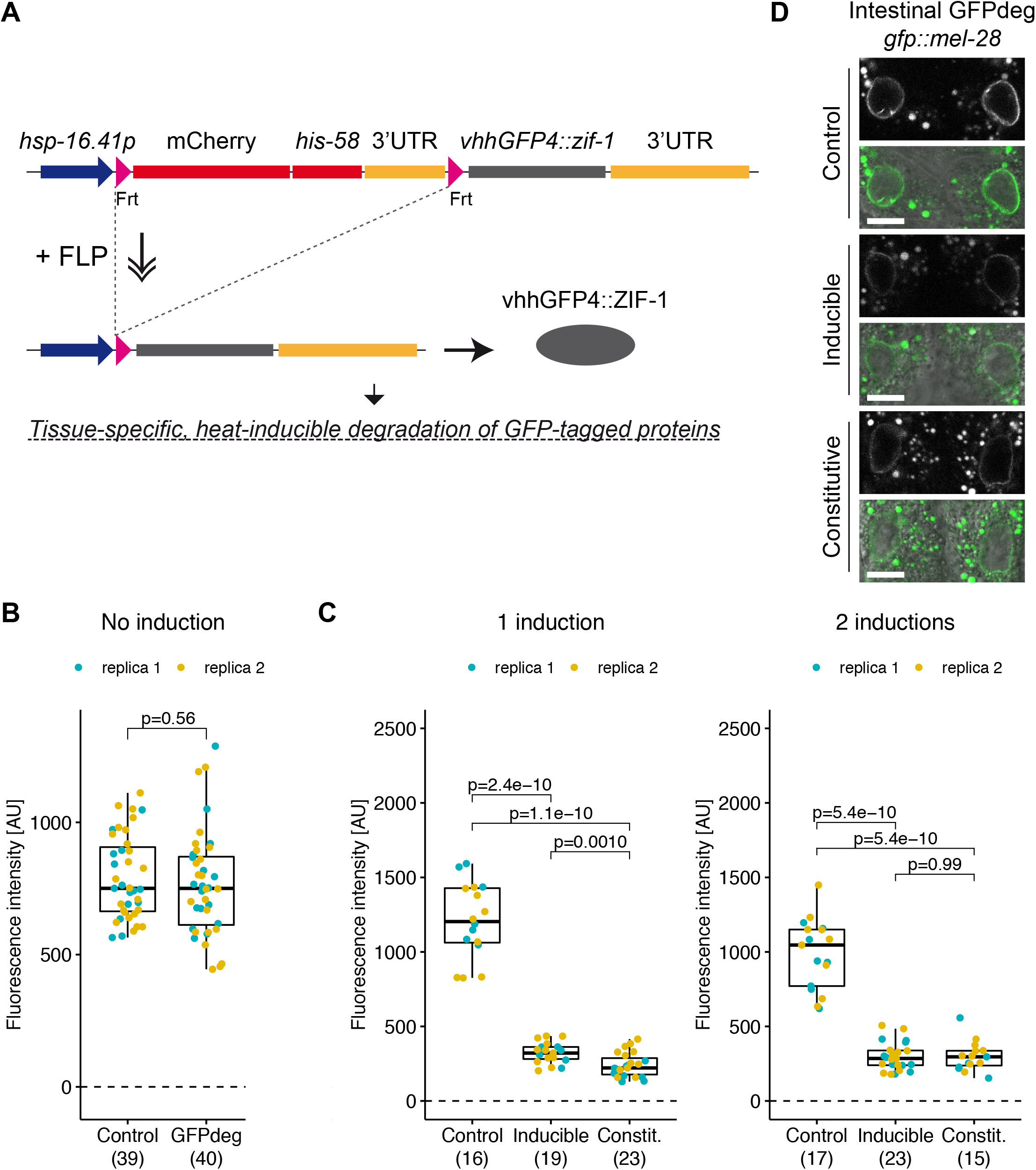
Degradation of GFP::MEL-28 in intestinal cells. (A) Schematic representation of the GFPdeg strategy. The vhhGFP4::ZIF-1 fusion protein targets GFP-tagged proteins for rapid degradation. Temporal and spatial control is provided by the *hsp-16.41* promoter and tissue-specific FLP expression, respectively. (B) Quantification of GFP::MEL-28 at the nuclear envelope of hypodermal nuclei of young adults either without the GFPdeg system (Control; strain BN452) or with the GFPdeg system but without induction (GFPdeg; strain BN1084). P value from t test is indicated. (C) Quantification of GFP::MEL-28 at the nuclear envelope of anterior intestinal nuclei of young adults after 1 or 2 rounds of GFPdeg induction in the intestine (*nhx-2p::FLP* driver; strain BN1118). Control (strain BN452) and constitutive degradation (*elt-2p::vhhGFP4::zif-1;* strain BN746) are included for comparison. Graph details are explained in Figure 3. P values from pair-wise t tests and adjusted for multiple comparisons (Benjamini & Hochberg method) are indicated. (D) Confocal micrographs of *gfp::mel-28* young adults without GFPdeg (top panels; strain BN452), inducible intestinal GFPdeg (middle panels; strain BN1118) or constitutive intestinal GFPdeg (bottom panels; strain BN746). A single focal plane of the anterior pair of intestinal nuclei is shown. Scale bar 10 μm.

We first determined that basal expression from the *hsp-16.41* promoter in the absence of heat induction does not cause depletion of GFP::MEL-28 in the hypodermis (Figure 4B). This implies that the system can be used to evaluate the function of essential proteins late in development by raising the animals at 16°C until induction. We next investigated whether the heat-induced GFPdeg system is equally efficient as the original constitutive variant (Wang *et al*. 2017). We focused on degradation of GFP::MEL-28 in intestinal cells. GFP intensity was quantified in one or two intestinal cells closest to the pharynx and to the microscope lens to ensure consistent measurements. Other intestinal cells often lie below the gonad, impeding reproducible quantification. Three hours after a short heat pulse (15 min; 34°C) the median GFP::MEL-28 intensity was reduced to ~27% in the inducible GFPdeg strain, whereas slightly less protein remained in the constitutive GFPdeg strain (~18%; Figure 4C, left graph). By exposing the animals to two 15 min heat pulses at 34°C, one 24h and another 3h prior to imaging, the efficiencies of the inducible and constitutive GFPdeg systems were equal (p=0.91; Figure 4C, right graph; Figure 4D). This demonstrates, that additional temporal control of the tissue-specific GFPdeg module can be achieved without compromising degradation efficiency. Depending on the protein of interest, induction conditions may have to be optimized. For instance, the efficiency of endogenous GFP::MEL-28 degradation in the intestine by either the constitutive or the inducible system (Figure 4C-D) is lower than the depletion reported for endogenous GFP::MDF-1 based on the identical constitutive GFPdeg strain (Wang *et al*. 2017). To evaluate the degradation dynamics in more details, we measured GFP::MEL-28 levels in hypodermal cells at different time frames after a single induction of GFPdeg (15 min; 34°C). Hypodermis-specific expression of vhhGFP4::zif-1 was ensured by excision of *mCh::his-58* with FLP under the control of the *dpy-7* promoter. All images were taken from a region of the hypodermis closest to the lens and just posterior to the pharynx. One hour after induction, the median level of GFP::MEL-28 was reduced to ~55% and further down to ~19-20% after 2-4 h (Figure 5). A possible explanation for the remaining GFP::MEL-28 even at the optimal time points (2-4 h) could be novel protein synthesis since the degradation system will only be active for a finite time after induction. Indeed, substantial protein re-synthesis was observed 24 h after heat shock (Figure 5). The time frame of optimal degradation efficiency allows some flexibility in protocol design. However, these protocols, especially number and timing of heat inductions, presumably need to be tailored to the protein of interest and its turnover.

**Figure 5.**
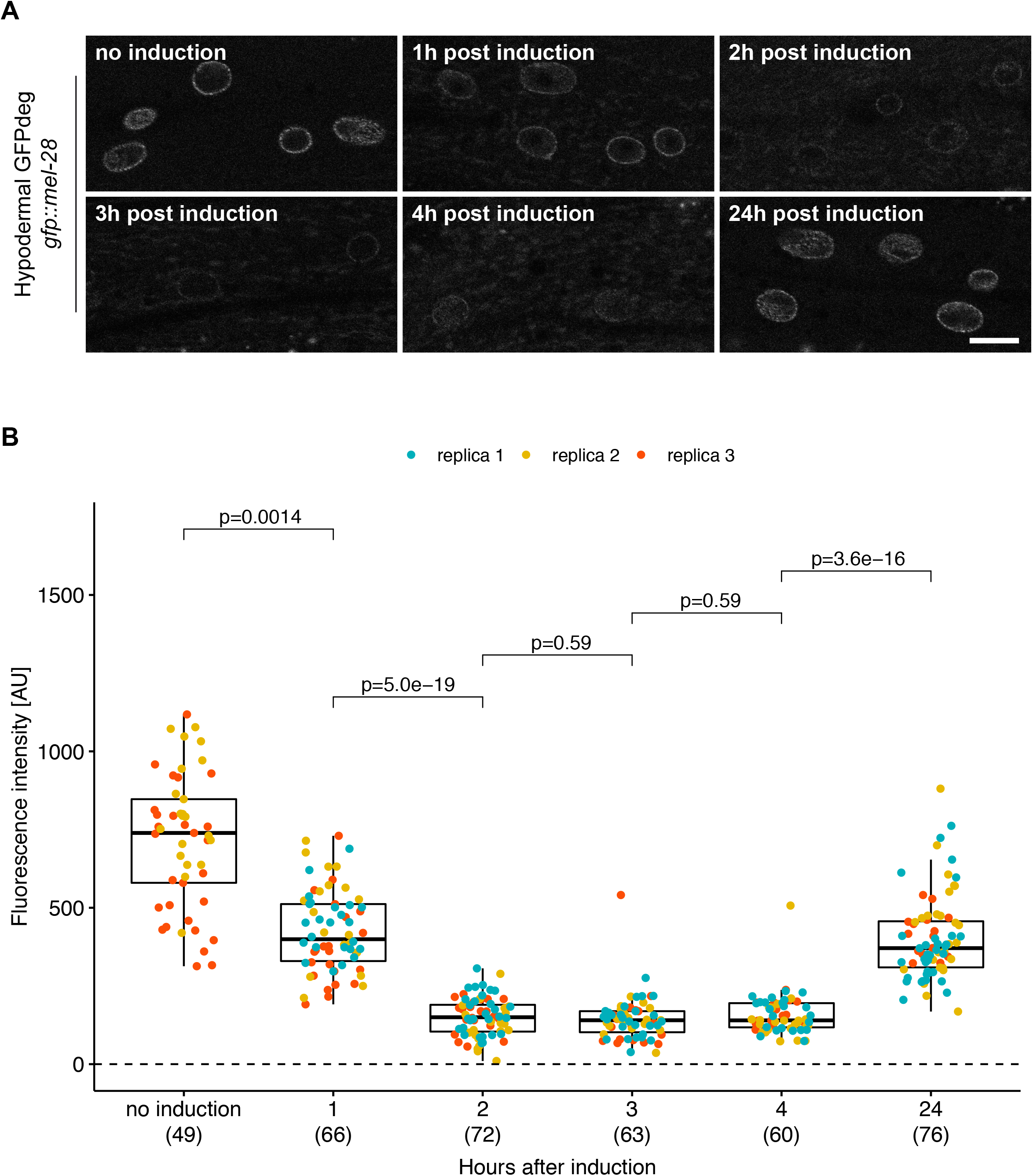
Kinetics of GFP::MEL-28 degradation in hypodermal cells. (A) Confocal micrographs of *gfp::mel-28* young adults either without induction or 1-24 h after induction of GFPdeg in the hypodermis (*dpy-7p::FLP* driver; strain BN1084). A single focal plane of the hypodermis is shown. Scale bar 10 μm. (B) Quantification of GFP::MEL-28 at the nuclear envelope of hypodermal nuclei. Graph details are explained in Figure 3.

### Combination of FLP-mediated gene knockout and protein degradation

Having available two FLP-controlled systems for inhibition of protein expression, we decided to test the efficiency of combining them. In this setup, the same FLP driver controls both conditional gene KO and protein degradation (Figure 6A). As a proof of principle, we evaluated the combined action of the GFP KO cassette and GFPdeg on the expression of endogenously GFP-tagged MEL-28 in the hypodermis. GFPdeg was induced by exposing young L1s to a short heat pulse (15 min; 34°C) followed by incubation at 16°C. The effect was first evaluated in L1 larvae after 3 h and again in L2 larvae as well as young adults. In contrast to the GFP KO system which alone had insignificant effects in L1 larvae, the combination of the two systems led to ~80% reduction of median GFP::MEL-28 signal (Figure 3). In L2 larvae, GFP::MEL-28 levels in the hypodermis were further decreased to ~3% and undetectable in most young adults. Thus, in all three life stages tested, the combination of conditional gene knockout and protein degradation provided the most efficient inhibition of protein expression. We also compared the efficiency of either conditional gene KO alone or in combination with targeted protein degradation by inserting the GFP KO cassette in frame after the start codon of another stable nuclear pore protein, NPP-2/NUP85. In young adults, the conditional KO caused a reduction of ~97% of GFP::NPP-2 in the hypodermis, whereas only ~1.5% remained when conditional KO was combined with the GFPdeg system (Figure 6B-C).

**Figure 6.**
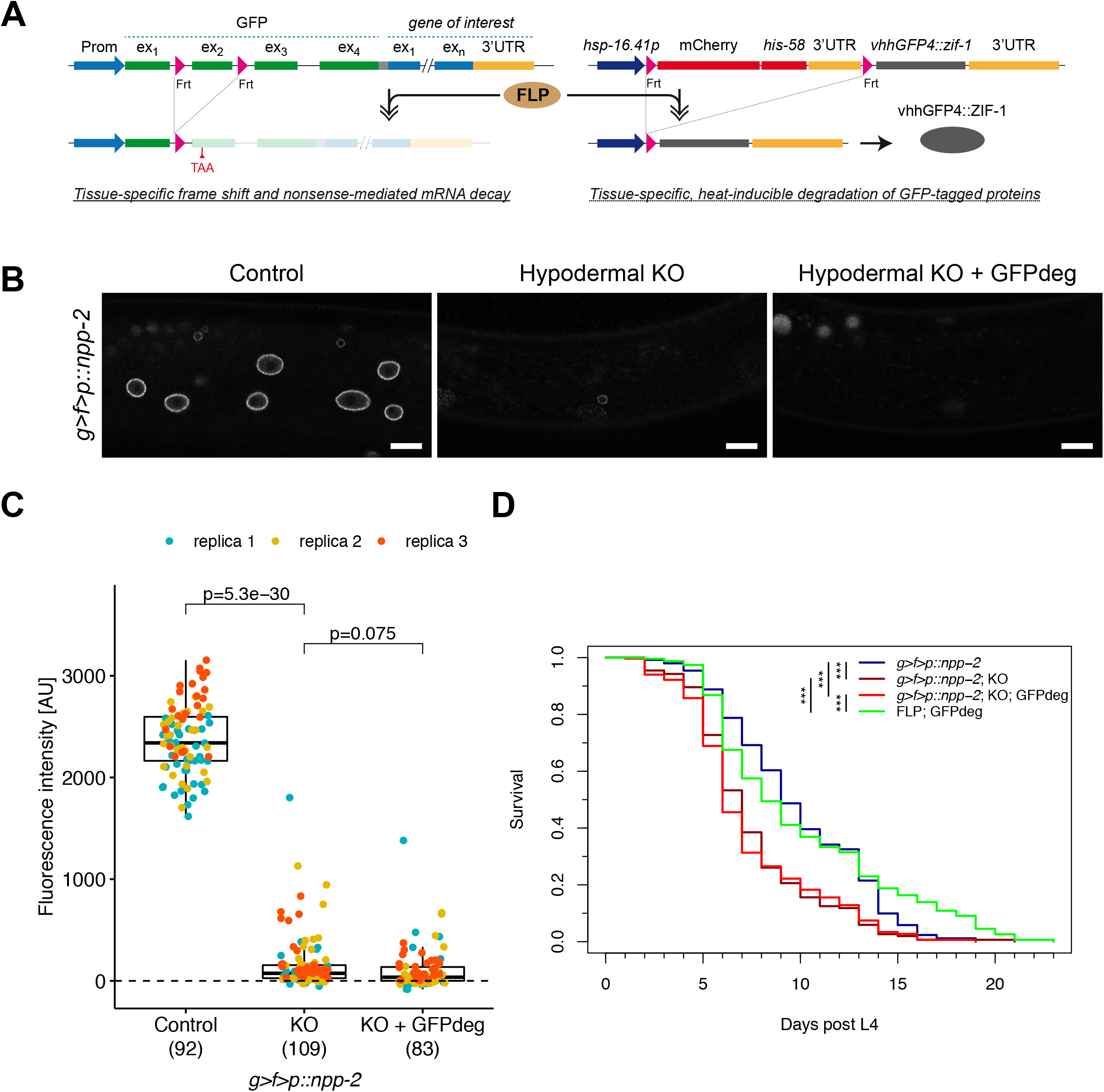
Efficient protein depletion by combined gene KO and GFPdeg. (A) Schematic representation of the combined KO and GFPdeg strategy. A single FLP driver controls in a tissue-specific manner both KO (left) and protein degradation (right) of a gene tagged with the GFP KO cassette. (B) Confocal micrographs of *g>f>p::npp-2* L4 larvae either not expressing FLP (left panel; strain BN1082), expressing FLP in the hypodermis (middle panel; *dpy-7p::FLP* driver; strain BN1111) or expressing FLP and vhhGFP4::ZIF-1 in the hypodermis (right panel; *dpy-7p::FLP* driver; strain BN1086). A single focal plane of the hypodermis is shown. Scale bar 10 μm. (C) Quantification of GFP::NPP-2 at the nuclear envelope of hypodermal nuclei. Graph details are explained in Figure 3. (D) Survival curves for animals expressing GFP::NPP-2 alone (blue; strain BN1082) or combined with either FLP in the hypodermis (dark red; strain BN1111) or FLP and GFPdeg in the hypodermis (red; strain BN1086). A strain with hypodermal FLP and GFPdeg but without target was included as control (green; BN1097). The lifespan of the two strains with depletion of GFP::NPP-2 in the hypodermis was significantly shorter compared to the two control strains (*** indicates p<1e-08). More than 175 animals were analysed for each genotype.

To test if these dramatic reductions of endogenous GFP::NPP-2 in the hypodermis had consequences on the health of the animals, we determined their lifespan at 25°C. We followed >175 animals for each genotype and observed a 22-33% reduction in median lifespan from 9 days post L4 in GFP::NPP-2 control animals (95% confidence interval [CI] = 9-10 days) to 7 days in conditional KO animals (95% CI = 6-7 days) and to 6 days in animals with combined conditional KO and GFPdeg systems (95% CI = 6-7 days) (Figure 6D. Expression of FLP and GFPdeg in the absence of the GFP KO cassette had no effect on the lifespan (p=0.3). The statistically insignificant shift to the left of the lifespan curve for the animals with the combined conditional KO and GFPdeg (red line) as compared to the animals with the conditional KO only (dark red line; p=0.5) is in line with the unsubstantial difference in GFP::NPP-2 signal intensity between the two strains. We conclude that conditional NPP-2 KO in the hypodermis causes a dramatic shortening of lifespan, illustrating the suitability of the combination of the GFP KO cassette and the GFPdeg system for evaluation of physiological consequences of protein depletion in specific tissues.

### Tissue-specific transcriptomic profiling by RAPID

Various complementary methods are available for identification of gene expression profiles in specific cell types in *C. elegans*, including physical isolation of cells or nuclei of interest (Haenni *et al*. 2012; Steiner *et al*. 2012; Kaletsky *et al*. 2018), affinity purification of RNAs from specific cells (Blazie *et al*. 2017), trans-splicing-based RNA tagging (Ma *et al*. 2016), and mapping of RNA polymerase occupancy (Gomez-Saldivar *et al*. 2020; Katsanos *et al*. 2021). RNA polymerase DamID (RAPID) (Gomez-Saldivar *et al*. 2020) and Targeted DamID (TaDa) (Katsanos *et al*.2021) are based on tissue-specific basal expression of *E. coli* Dam methyltransferase fused to the RNA polymerase subunit RPB-6. The Dam::RPB-6 fusion protein provides an *in vivo* footprint of RNA polymerases on transcribed genes by specific methylation of GATC sequences that are identified by enzymatic and deep sequencing techniques (Marshall *et al*. 2016; Askjaer and Harr 2021).

RAPID relies on recombinases to achieve tissue specificity and has been used in body wall muscles, intestine and specific neurons (Gomez-Saldivar *et al*. 2020). To increase the number of tissues immediately amenable to RAPID, we adapted it for FLP-mediated spatiotemporal control. We cloned and inserted a *dam::rpb-6* fusion gene downstream of the FRT-flanked *mCh::his-58* cassette used above and under control of the *hsp-16.41* promoter (Figure 7A). We introduced a single copy of this construct into the *C. elegans* genome and crossed the resulting line with a *dpy-7p::FLP* driver. Nematodes were cultured at 20°C to ensure low levels of Dam::RPB-6 expression in the hypodermis and total genomic DNA was purified from L4 larvae. DNA from animals expressing GFP::Dam was used to control for unspecific methylation. The genome-wide association profile of Dam::RPB-6 was determined by deep sequencing (see Materials and Methods), which revealed a list of 2331 protein coding genes with FDR < 0.05 (Table S5). When plotting the RAPID signal along all protein coding genes, an increase in signal is observed from the transcriptional start site (TSS) to the transcriptional end site (TES), similar to previous RAPID and TaDa observations (Gomez-Saldivar *et al*. 2020; Katsanos *et al*. 2021)(Figure 7B, left). However, clustering the signal from individual genes using k-means algorithm (with default number of clusters = 5 (Stempor and Ahringer 2016)) suggests that the average signal masks multiple different patterns of RAPID signal along genes (Figure 7B, middle). A more uniform RAPID signal from TSS to TES is also observed when averaging the 2331 genes determined as expressed in the hypodermis with FDR < 0.05 (Figure 7B, right). Among the transcribed genes are several known and expected hypodermal genes, such as *col-39, col-79, col-80* and *sqt-1* that express collagens as well as ubiquitously expressed genes *eef-1A.1* and *fib-1* (Figure 7C).

**Figure 7.**
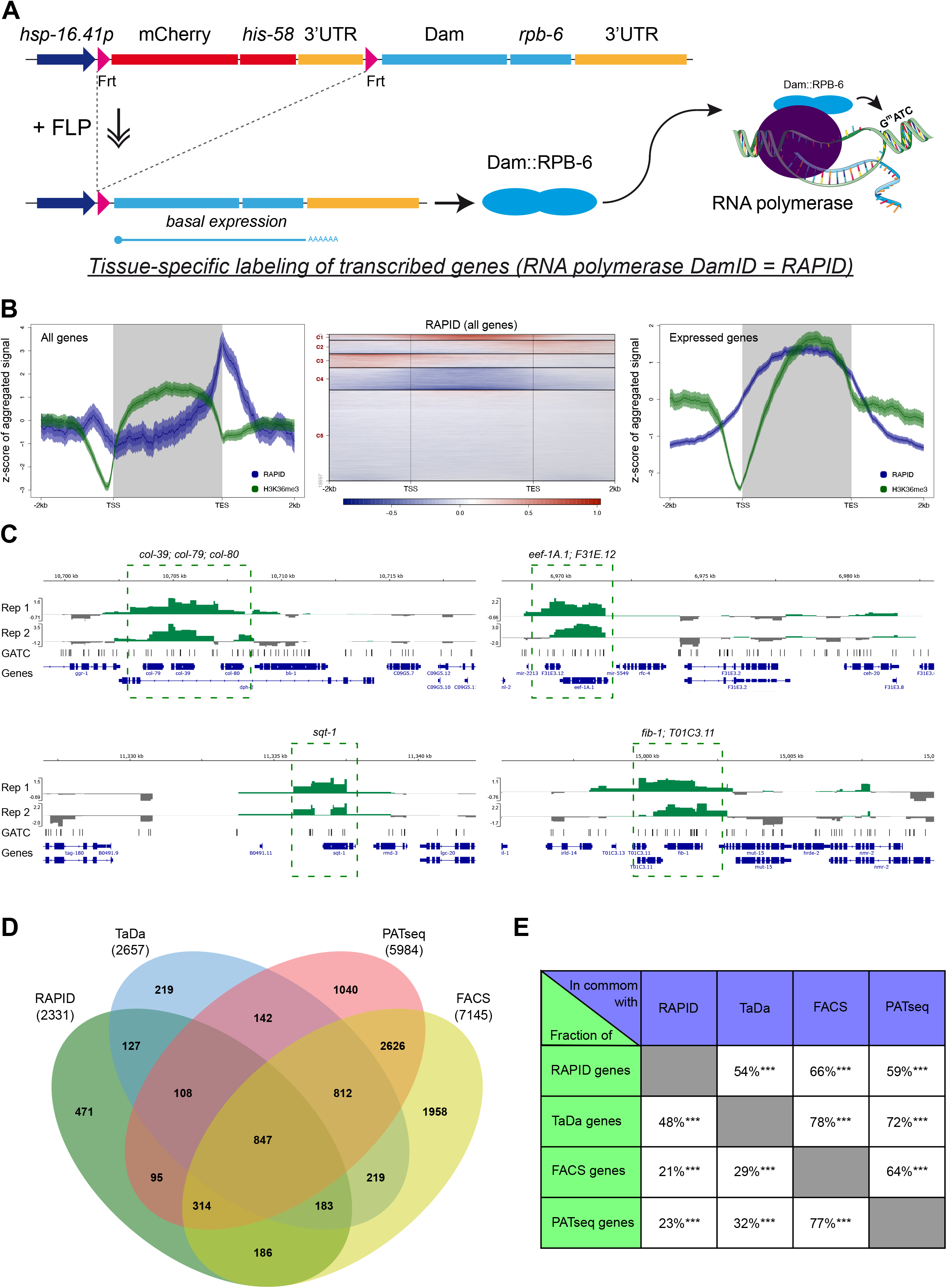
Tissue-specific identification of transcribed genes by RAPID. (A) Schematic representation of the RAPID method. Spatial control is provided by tissue-specific FLP expression that leads to excision of the *mCherry::his-58* cassette and basal transcription of *Dam::rpb-6* from the *hsp-16.41* promoter. Dam::RPB-6 incorporates into RNA polymerases and marks GATC sites in transcribed genes. A *GFP::Dam* transgene is used in parallel to control for differences in chromatin accessibility and unspecific methylation (not shown). (B-E) Data based on two replicas of L4s expressing either Dam::RPB-6 (strain BN1414) or GFP::Dam (strain BN561) in the hypodermis. The analysis was restricted to 19,975 protein coding genes annotated in WBCel235. (B) RAPID signal (blue) averaged in 10 bp bins across all genes (left graph) or across 2331 expressed genes (right graph) from 2 kb upstream of the transcriptional start site (TSS) to 2 kb downstream of the transcriptional end site (TES). The signal from TSS to TES was extrapolated to an arbitrary length of 3 kb for each gene. The distribution of the active chromatin mark H3K36me3 (Bannister *et al*. 2005) is included for comparison. The heatmap (middle graph) represents the individual genes from the left graph. (C) Representation of Dam::RPB-6/GFP::Dam ratios for individual replicas (Rep 1, Rep 2) at selected loci. Genes with significant RAPID signal are indicated (FDR<0.05). (D-E) Comparison of the 2331 transcribed genes identified by RAPID with three alternative methods for transcriptomic profiling (see main text for details). Numbers in parentheses indicate the number of protein coding genes reported in each study.

To evaluate the specificity of RAPID, we compared the list of 2331 transcribed genes to three datasets of genes reported to be expressed in the hypodermis (Blazie *et al*.2017; Kaletsky *et al*. 2018; Katsanos *et al*. 2021) (Figure 7C-D; Table S6). Our list overlapped significantly with each of the individual datasets (54-66% overlap; p<2.2e-16 Fisher’s exact test), and 1860 genes (80%) overlapped with at least one of the three datasets. These overlaps are comparable to those observed when comparing the three other datasets among each other (Figure 7D). Moreover, when exploring a small arbitrarily chosen subset of the 471 genes not overlapping with any of the three other datasets, at least 22/50 (44%) are reliably detected in hypodermal cells in single cell RNAseq performed on embryos (Packer *et al*. 2019) (Table S7). This argues that many of the genes identified uniquely by RAPID are expressed in the hypodermal lineage during development. We conclude that RAPID is an attractive and versatile method to identify transcribed genes in specific tissues with high confidence.

## Discussion

An increasing number of specific and efficient FLP and Cre drivers generated by the *C. elegans* community are facilitating a variety of precise functional assays to interrogate neural circuits (Davis *et al*. 2008; Schmitt *et al*. 2012; Konietzka *et al*.2020; Davis *et al*. 2021), cell proliferation and differentiation (Ruijtenberg and van den Heuvel 2015), gene expression programs (Gomez-Saldivar *et al*. 2020; Van de Walle *et al*. 2020) and chromatin organization (Cabianca *et al*. 2019; Harr *et al*.2020). In this study, we report four new FLP lines that reliably induce recombination specifically in either distal tip cells, the somatic gonad, coelomocytes or in epithelial Pn.p cells. The efficiency of recombination was ~100% in all animals evaluated for the four lines. We also demonstrate that similarly high recombination efficiency is achieved when FLP is expressed from single-copy transgenes on different autosomes. In the case of the *ckb-3p::FLP* line, recombination was observed already in the earliest progenitors Z1 and Z4, which implies that the line is suitable to target all cell types of the somatic gonad, including distal tip cells, gonadal sheath, spermatheca, spermatheca-uterine valve and the uterus (Hubbard and Greenstein 2005). In contrast, the activity of the *hlh-12p::FLP* line is restricted to the distal tip cell and may thus be useful to investigate its stem cell niche function (Kimble and Seidel 2008). Similarly, biological processes in coelomocytes are amenable to manipulation by the *unc-122p::FLP* line, whereas the *lin-31p::FLP* line induces recombination in the cells producing the vulva. We previously created a FLP line to induce recombination in intestinal cells based on the *elt-2* promoter. However, because this promoter can also drive expression in other tissues (Ruijtenberg and van den Heuvel 2015; Munoz-Jimenez *et al*. 2017), we developed here a novel line based on the *nhx-2* promoter. The new line is equally efficient to produce recombination in the intestine (Table 1) and with the advantage of not provoking recombination elsewhere. We also attempted to construct lines for specific neurons using the promoters of *gpa-14* and *unc-17*. In the case of *gpa-14*, we observed variable recombination rates in FLP-expressing cells (0, 1 or 2 alleles of the single-copy reporter). It seems that the behavior of the specific cells is reproducible between individuals, but we do not know if this is due to differences in FLP expression levels and/or the FRT sites being less accessible in certain cells. For the *unc-17p::FLP* line, we observed widespread recombination in cholinergic neurons as expected but also in the hypodermis. We speculate that this could be due to an inability of our construct to precisely reflect *unc-17* expression (Pereira *et al*. 2015), although we also note that two cholinergic neurons, PVNL and PVNR, are sister cells to two hypodermal cells (TLappa and TRappa). Thus, if *unc-17p::FLP* is activated already in the progenitor cells TLapp and TRapp, GFP::HIS-58 expression would be expected in both cell types.

We previously proposed that inserting a GFP KO cassette (with FRT sites in *gfp* introns 1 and 2) immediately after the start codon of a gene provides an informative GFP fusion protein and a conditional KO allele in a single step (Munoz-Jimenez *et al*. 2017). In this study we have further validated this approach by comparing strains that either carry the GFP KO cassette in the endogenous *baf-1* locus or a GFP cassette with a single FRT and a second FRT downstream of *baf-1*. When combined with an *nhx-2p::FLP* driver to KO *baf-1* specifically in the intestine, identical reductions of fertility were detected in the two strains. Furthermore, using the GFP KO cassette, we determined that conditional KO of *npp-2*/NUP85 in the hypodermis caused a dramatic lifespan shortening. We therefore conclude that insertion of the GFP KO cassette into an endogenous locus by CRISPR provides an attractive dual tool to study protein dynamics and KO phenotypes in a single workflow. A limitation of the approach is that the GFP KO cassette should be inserted in the 5’-end of the open reading frame. As with other gene tagging experiments, tests for hypomorph phenotypes should be performed.

Although gene KO is often considered an ultimate test for gene function, slow mRNA and protein turnover can delay the appearance of loss-of-function phenotypes. This is particularly relevant for organisms with rapid development and relatively short lifespan, such as *C. elegans*. Moreover, the irreversible nature of gene KO may be disadvantageous in some experimental designs. Targeted protein degradation is a relevant alternative and auxin-inducible degradation (AID) stands out as an attractive and popular option due to its fast kinetics and both inducible and reversible nature (Zhang *et al*. 2015). Concerns have been raised regarding auxin-independent depletion of AID-tagged proteins (see references in (Nance and Frokjaer-Jensen 2019)) and potential effects of auxin on endoplasmic reticulum stress (Bhoi *et al*. 2021), but recent optimization of AID tools may have overcome this (Hills-Muckey *et al*. 2021). We show here that GFPdeg, an alternative protein degradation system based on a GFP-binding nanobody (Wang *et al*. 2017) and adapted for FLP-mediated spatiotemporal control, can efficiently deplete 80% of a long-lived nuclear pore protein, MEL-28/ELYS, fused to GFP. By comparing the degradation of GFP::MEL-28 in intestinal cells using either the original, constitutive version of GFPdeg (Wang *et al*. 2017) or the heat inducible version of GFPdeg developed in this study, we conclude that they are equally efficient. Importantly, the GFPdeg system is immediately applicable to study the function of any GFP-tagged fusion protein. We chose to focus on stable nuclear pore proteins (Savas *et al*. 2012; Toyama *et al*. 2013) in our study as they are probably more challenging to deplete but we assume that higher depletion efficiencies will be observed for proteins with shorter half-lives. The original GFPdeg version (Wang *et al*. 2017) has the advantage that it involves only two components (GFP-tagged fusion protein and constitutive, tissue-specific GFPdeg) and does not require induction. In contrast, our FLP-based GFPdeg system includes three components (GFP-tagged fusion protein, tissue-specific FLP driver and heat-inducible, FRT-flanked GFPdeg) and requires one or two short 15 min exposures to 34°C. On the other hand, our inducible system allows to evaluate processes at any stage in development or during aging by raising the nematodes at standard temperatures until induction at the desired time point. This cannot be achieved with the constitutive system where the animal’s development potentially arrests once the concentration of the target protein is below a required threshold. We envision that the FLP-controlled GFPdeg system can be further modified to incorporate other inducers (Monsalve *et al*. 2019; Wurmthaler *et al*. 2019; Driesschaert *et al*. 2021) in situations where short heat pulses are incompatible with the experimental design.

Depending on the experimental setup, either site-specific gene KO or targeted protein degradation may be sufficient to elucidate gene function. However, bimodal control of protein levels may be particularly valuable for the study of long-lived proteins during early developmental stages as highlighted in this study. By combining the GFP KO cassette and the GFPdeg system, we have generated a protocol that provides two-fold control of tissue-specific protein depletion. On one hand, a specific FLP driver induces gene KO in a tissue of interest and thus prevents synthesis of mRNA from the target gene. On the other hand, the same FLP driver excises tissue-specifically the FRT-flanked *mCh::his-58* cassette from the GFPdeg construct, thus enabling heat-induced and rapid degradation of existing GFP-tagged target protein. We show that the combined KO and GFPdeg systems can deplete GFP::MEL-28 and GFP::NPP-2 to non-detectable levels, thus offering a highly efficient approach for spatiotemporal control of protein levels in a multicellular organism.

One of the strengths of recombinase-based genome engineering is its versatility: the possibility of using collections of stable FLP and Cre drivers to control a variety of different effectors through straight-forward genetic crosses (Hubbard 2014; Nance and Frokjaer-Jensen 2019; Driesschaert *et al*. 2021). The Cre/loxP system was recently utilized to generate tissue-specific transcription footprints by RNA polymerase DamID (RAPID) in *C. elegans* (Gomez-Saldivar *et al*. 2020). In RAPID and other DamID methods for transcriptional profiling *E. coli* Dam is fused to an RNA polymerase subunit and expressed at low levels to mark *in vivo* transcribed genes (Southall *et al*. 2013; Askjaer and Harr 2021). In this study we have widened the range of tissues immediately accessible to RAPID by adapting RAPID for FLP-mediated spatial control. We have validated the potential of RAPID in *C. elegans* by expressing Dam::RPB-6/POLR2F specifically in the hypodermal lineage through the use of a *dpy-7p::FLP* driver and identified 2331 transcribed genes. We compared our list with hypodermal data from another recent DamID-based (TaDa) study (Katsanos *et al*. 2021) and two RNAseq-based studies (Blazie *et al*. 2017; Kaletsky *et al*. 2018) and found a significant overlap in all cases (p<2.2e-16). We propose that the set of 847 protein coding genes identified in all four studies (Table S6) can be considered a high-confidence transcriptional signature of the hypodermis. A pilot comparison of genes found uniquely in RAPID (i.e. not confirmed by any of the three other methods) revealed a relevant overlap with genes detected in hypodermal cells by single-cell RNA-seq (Packer *et al*. 2019), suggesting that many of the unique RAPID genes are true positive hits.

Although the overlap between the gene lists identified in our RAPID experiment and the TaDa study is much higher than would be expected by chance, ~50% of the genes identified in one study were not found in the other. In both cases, Dam::RPB-6 was used to mark transcribed genes and samples were prepared at the L4 stage so how can the many unique genes be explained? An important difference is that Katsanos and co-workers used a mutated version of the *dpy-7* promoter (*“dpy7syn1”*) for TaDa in the hypodermis to avoid signal from seam cells (Katsanos *et al*. 2021), whereas we used a wild type *dpy-7p* sequence to drive FLP expression (Munoz-Jimenez *et al*. 2017). Katsanos and co-workers also determined the transcriptomic profile for seam cells and found that ~1/3 of the genes in their hypodermis and seam cell datasets are unique (Katsanos *et al*. 2021). Our RAPID gene list overlaps significantly with both the entire seam cell gene list (1013 overlapping genes; p<2.2e-16) and the unique seam cell gene list (106 overlapping genes; p=0.016), suggesting that part of the RAPID signal in our experiments is derived from seam cells (Figure S2). It should be noted that the lineages of the two cell types are interconnected because many of the daughter cells from dividing seam cell precursors fuse with the hypodermis during larval development. Differences in the protocols for preparation of deep sequencing libraries (sequences of adapters, PCR conditions, +/- removal of adapters and fragmentation, etc.) are also likely to introduce variation in the lists of identified genes. Finally, the strategies underlying tissue specificity is also different between RAPID and TaDa (Askjaer and Harr 2021). Spatial control of basal Dam::RPB-6 expression is achieved in RAPID by combining FLP or Cre-mediated genome recombination with a non-induced *hsp* promoter (Figure 7A), whereas TaDa is based on a tissue-specific promoter and translation re-initiation of a bicistronic mRNA containing an upstream open reading frame and *Dam::rpb-6* separated by one or several stop codons. TaDa is feasible with a single transgene whereas RAPID relies on a recombinase transgene and a *Dam::rpb-6* transgene typically integrated on separate chromosomes. Although this appears to favor TaDa versus RAPID, the growing list of stable and efficient FLP and Cre drivers implies that different tissues can be explored with RAPID through a simple cross to combine the recombinase and *Dam::rpb-6* transgenes. In contrast, TaDa requires the design, cloning and transgenesis for each new cell type to be analyzed.

Because the methylation on GATC by Dam is only erased when cells enter S phase and replicate their DNA, RAPID and TaDa provide the history of transcription during the life of the cell. PATseq and FACS produce instead a compilation of all mRNAs present in the cell at the time of harvest. Depending on the biological question to be addressed, a particular method might be most suitable. The PATseq and FACS studies report the largest hypodermal gene sets (Blazie *et al*. 2017; Kaletsky *et al*.2018), but they also suffer higher intrinsic risks of including false positive hits. For instance, FACS isolation of a particular cell type might include contaminants in the form of cells from other tissues. Similarly, in the PATseq protocol, mRNA from unlabeled cells might potentially co-purify with the mRNA from the target cells. Such false positives are difficult to control for in the FACS and PATseq protocols. In contrast, RAPID and TaDa are based on isolation of genomic DNA from the entire organism followed by amplification of methylated GATC fragments from a tissue of interest, library preparation and a statistical analysis to identify genes with a higher Dam::RPB-6/GFP::Dam methylation ratio. Importantly, although the signal from Dam::RPB-6 reflects RNA polymerase occupancy on transcribed genes rather than transcription itself, the signal intensity correlates with transcript abundance (Southall *et al*. 2013; Gomez-Saldivar *et al*. 2020; Katsanos *et al*. 2021). Furthermore, comparing Dam::RPB-6 methylation values across experimental conditions enables identification of differentially expressed genes (Widmer *et al*.2018).

We conclude that FLP-controlled RAPID has the capacity to identify transcribed genes in a tissue of interest with high confidence. We have tested the method in the hypodermis, but it is easily applied to other cell types for which an appropriate FLP driver is available. In particular, we envision that RAPID will be advantageous for cell types that are hard to isolate in an intact state, such as neurons (Gomez-Saldivar *et al*. 2020).

## Acknowledgments

We are grateful to Andrew Chisholm, Carmen Ortiz González, Michael Krieg, Montserrat Porta, Ngang Heok Tang, Patricia Regina Rider García and Sara Castaño Díaz for reagents and advice. We also wish to acknowledge Ana Miñan Fernández from the CABD Functional Genomics Facility for microinjections, Laura Tomás from the CABD Proteomics and Biochemistry Facility for recombinant proteins, Katherina García from the CABD Imaging Facility for technical assistance; the Caenorhabditis Genetics Center (CGC), which is funded by the National Institutes of Health (NIH) Office of Research Infrastructure Programs (P40 OD010440) and WormBase. This project was funded by the Spanish State Research Agency and the European Regional Development Fund (BFU2016-79313-P, CEX2020-001088-M and PID2019-105069GB-I00).

---

Supplementary Figure S1. Characterization of novel FLP drivers. Upon temperature induction of the *hsp-16.41* promoter, the reporter expresses red (mCh) or green (GFP) histone HIS-58 in cell lineages without or with FLP-mediated recombination, respectively. (A) Recombination activity by *lin-31p::FLP::SL2::mNG* is observed in the head (left panel) and central region (right panel) of BN1023 animals. Bright GFP::HIS-58 expression is observed in the nuclei of cells forming the vulva (filled arrows), whereas faint expression is detected in the hypodermis (open arrows; right panel). Presumably, the faint expression is derived from Pn.p cells that fuse with hyp7. (B) Recombination activity by *gpa-14p::FLP::SL2::mNG* is observed in the head (left panel) and tail (right panel) of BN1208 animals. Filled arrows point to nuclei expressing GFP::HIS-58 but not mCh::HIS-58 (i.e. recombination of both alleles); a double arrow points to a nucleus expressing both fluorescent proteins (i.e. recombination of one allele) and open arrows indicate cells that express diffuse mNG (and hence also FLP) but not GFP::HIS-58. (C) Expression of mNG from the *unc-17p::FLP::SL2::mNG* transgene overlaps with mKate2 signal from endogenously tagged *unc-17* (lower left panel) and recombination (GFP::HIS-58) is detected as expected in multiple neurons in the head (upper left panel) and ventral nerve cord (upper right panel) of BN1133 animals. Unexpectedly, GFP::HIS-58 is also observed in hypodermal nuclei (lower right panel). (D) Recombination activity by *hlh-12p::FLP::SL2::mTagBFP2* is observed in the distal tip cell of BN1319 animals. The distal tip cell is marked by co-expressed mTagBFP2. Scale bars 10 μm.

Supplementary Figure S2. Overlap between protein coding gene sets identified by TaDa in seam cells and hypodermis (Katsanos *et al*. 2021) and by RAPID in hypodermis (this study).

