## Supplementary Figures and Tables for "Expanded FLP toolbox for spatiotemporal protein degradation and transcriptomic profiling in *C. elegans*"

Supplementary Figure S1

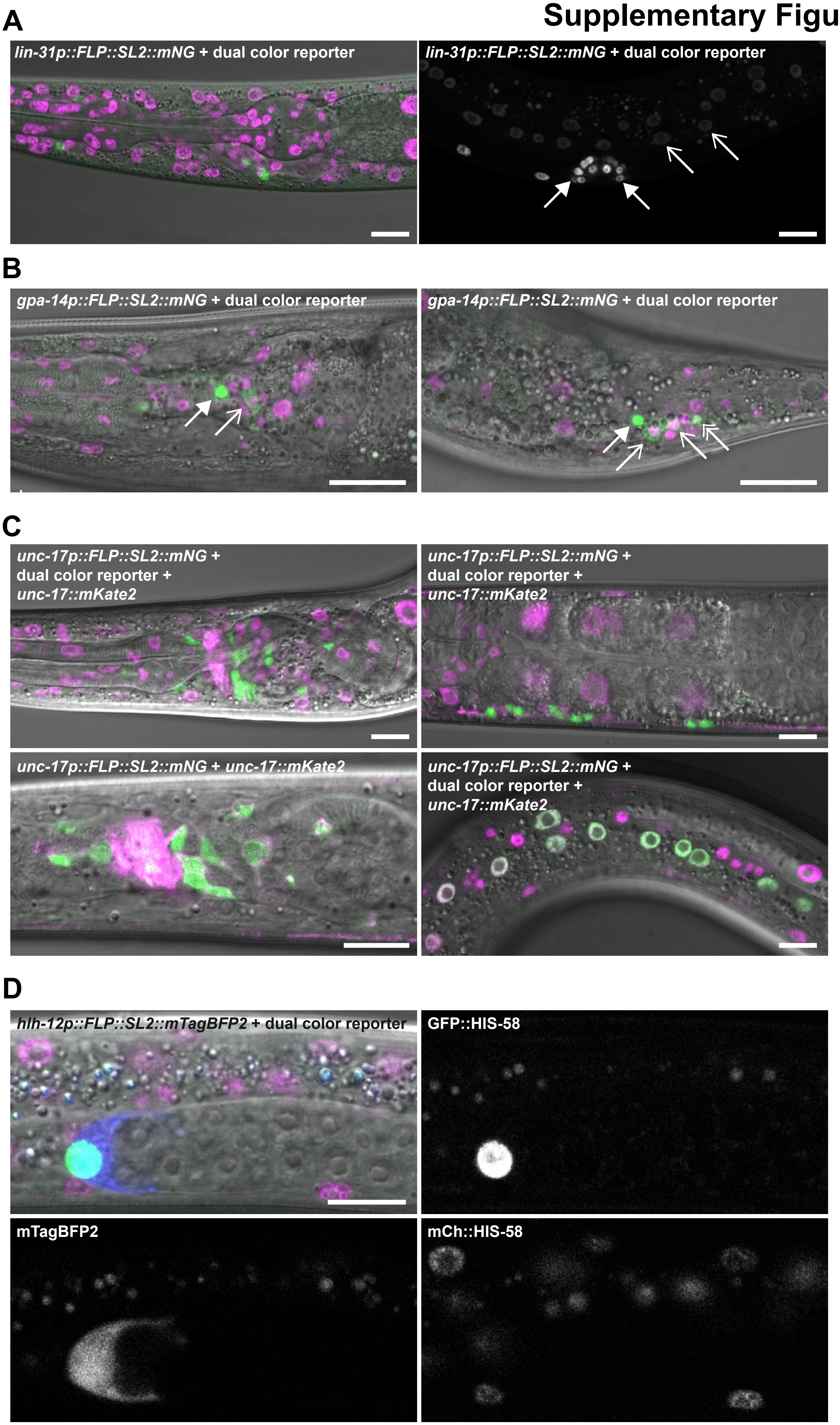

### Supplementary Figure S2

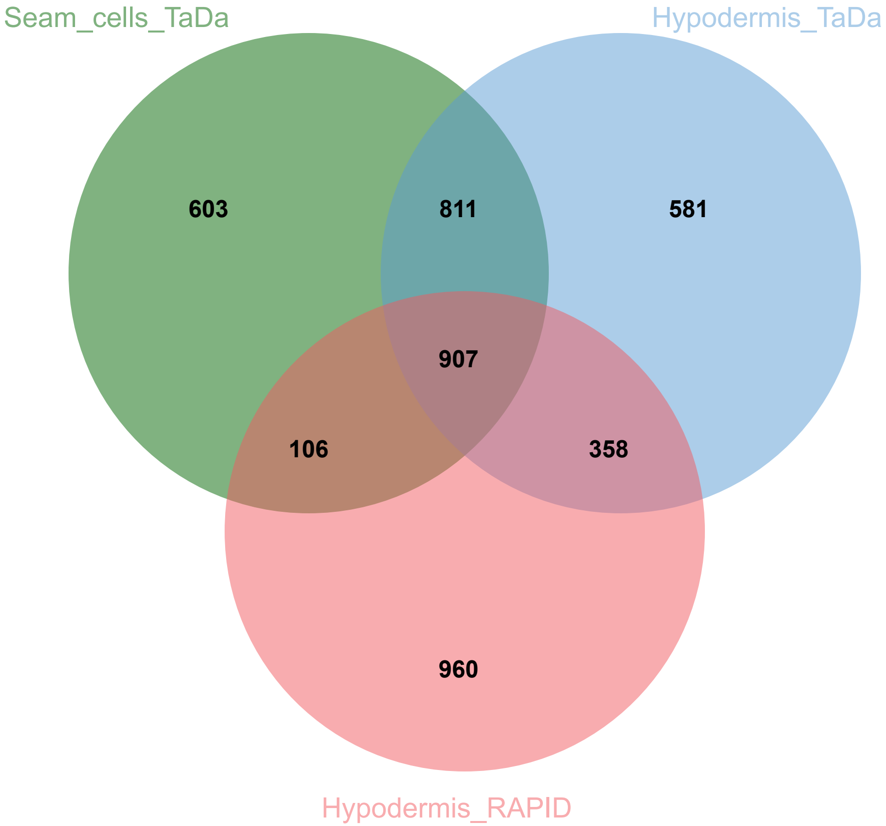

**Supplementary Table S1 Primers used in this study**

| Primer # | Primer sequence | Primer description |
|---|---|---|
| B1012 | ACATCAACCGTCGTATCTAAcgcgtgctgtctcatcctactttca | Forward primer to amplify gpd-2/-3 transsplicing sequence |
| B1364 | TCCTTGATGAGTTCGGACATgatgcgttgaagcagtttcc | Reverse primer to amplify gpd-2/-3 transsplicing sequence |
| B1365 | ggaaactgcttcaacgcatcATGTCCGAACTCATCAAGGA | Forward primer to amplify mTagBFP2 |
| B1366 | TGAAGGATATGCAGATACATacgcgTTAGTTGAGCTTGTGTCCGAG | Reverse primer to amplify mTagBFP2 |
| B961 | ttgATGCTTAACAGAAGTCGACA | baf-1 sgRNA; top oligo; for SapTrap vector; cuts at ATG |
| B962 | aacTGTCGACTTCTGTTAAGCAT | baf-1 sgRNA; bottom oligo; for SapTrap vector; cuts at ATG |
| B963 | TGGcgattttcgaaaatattttcaaattcaagaatgtaattattttgtgtttcagaaacc | baf-1 5'HA for SapTrap vector; top oligo; baf-1 promoter |
| B964 | CATggtttctgaaacacaaaataattacattcttgaatttgaaaatattttcgaaaatcg | baf-1 5'HA for SapTrap vector; bottom oligo; baf-1 promoter |
| B965 | acgTCGACTTCTGTTAAGCATCGTGAGTTCGTCGGAGAGCCAATGGGCGACAAAGAAGTC | baf-1 3'HA for SapTrap vector; top oligo; baf-1 ORF |
| B966 | tacGACTTCTTTGTCGCCCATTGGCTCTCCGACGAACTCACGATGCTTAACAGAAGTCGA | baf-1 3'HA for SapTrap vector; bottom oligo; baf-1 ORF |
| B1042 | gaGCTCTTCgTggtttcgcgatgcgcctat | mel-28 5'HA forward primer for #1397; mel-28 promoter |
| B1043 | gaGCTCTTCcCATtctgaaaaaaaataaacctttgc | mel-28 5'HA reverse primer for #1397; mel-28 promoter |
| B1044 | gaGCTCTTCaGGTaccggatcagctggat | mel-28 3'HA forward primer for #1397; linker and mel-28 5'end |
| B1045 | gaGCTCTTCcTACAGTCCGCATCGATCAAAGCA | mel-28 3'HA reverse primer for #1397; linker and mel-28 5'end |
| B1050 | ttgATCAAGGATACGAGTGTTGG | mel-28 sgRNA; top oligo; for SapTrap vector; cuts at ATG |
| B1051 | aacCCAACACTCGTATCCTTGAT | mel-28 sgRNA; bottom oligo; for SapTrap vector; cuts at ATG |
| B1231 | ttgttagtatctctttcagATGA | npp-2 sgRNA; top oligo; for SapTrap vector; cuts at ATG |
| B1232 | aacTCATctgaaagagatactaa | npp-2 sgRNA; bottom oligo; for SapTrap vector; cuts at ATG |
| B1233 | tggctaccgtgcttactatatttttaataccacttcacatgacattagtatctctttcag | npp-2 5'HA for SapTrap vector; top oligo; npp-2 promoter |
| B1234 | catctgaaagagatactaatgtcatgtgaagtggtattaaaaatatagtaagcacggtag | npp-2 5'HA for SapTrap vector; bottom oligo; npp-2 promoter |
| B1235 | acgACGGATGTCGCTCGAGAATTCGGGATAGCTGTCAGCGACTCCTCCACGGTCTTCGCC | npp-2 3'HA for SapTrap vector; top oligo; npp-2 ORF |
| B1236 | tacGGCGAAGACCGTGGAGGAGTCGCTGACAGCTATCCCGAATTCTCGAGCGACATCCGT | npp-2 3'HA for SapTrap vector; bottom oligo; npp-2 ORF |
| B1067 | IDT | trRNA |
| B1068 | GCTACCATAGGCACCACGAG | dpy-10 crRNA |
| B725 | CACTTGAACTTCAATACGGCAAGATGAGAATGACTGGAAACCGTACCGCATGCGGTGCCTA<br>TGGTAGCGGAGCTTCACATGGCTTCAGACCAACAGCCTAT | dpy-10 ssDNA repair template |
| B1312 | AATGGGGCATAGTCGCAAAG | baf-1 crRNA |
| B1313 | atttgcacttcgaaaaacaccatttgttccagtgtagtcgttttccgcttGAAGTTCCTATTCTCTAGAAAGTA<br>TAGGAACTTCtgcgactatgccccatttcaaattaatctgtttctgaaatatttctcata | baf-1 ssDNA repair template with FRT |
| B1314 | caccatttgttccagtgtagtcg | Forward primer to detect FRT insertion in baf-1 3'UTR |
| B1315 | tagtccaagaccacacgacaag | Reverse primer to detect FRT insertion in baf-1 3'UTR |
| B1333 | GACCTCGAGaaaaATGGATCAAGTCCAACTGGTG | Forward primer to amplify vhhGFP4::ZIF-1 |
| B1334 | tgcGCTAGCTTATTGTTGAATATTTATCGTGCAC | Reverse primer to amplify vhhGFP4::ZIF-1 |
| B1456 | catctctgaagaggatctggccggctcaagatctATGGCCGATGAGGACGATTATC | Forward primer to amplify rpb-6 |
| B1457 | atgcggagcttcgcgatgcgctagcCTACCAATCAGCGAGTTGAA | Reverse primer to amplify rpb-6 |

**Supplementary Table S2 Novel promoters for FLP expression used in this study**

| Promoter | Primers (F=forward; R=reverse) | Length [bp] | Sequence verified |
|---|---|---|---|
| *ckb-3* | F ctgcggccgCAGTCTACCAACATTCTCGAGCTC (NotI) | 1949 | full length |
| | R caggtaccGTTAATTTAGCAGCTTTTGAGAAATG (KpnI) | | |
| *gpa-14a* | F AATTAAAGTTTCATTTTTCACTTGTG (NotI*) | 3008 | >90% |
| | R TCGGCTGGGTACCATACACCTG (KpnI) | | |
| *hlh-12* | F gtgcggccgcAGCTTTAGAACATTAAAAAAAACTAATA (NotI) | 971 | full length |
| | R caggtaccTTTAATAAAATTGTGTAAGATGACGCTA (KpnI) | | |
| *lin-31* | F GAGCCGGCTGTAGCATgcggcCGCTAGTTGCCAAGAGTTGC (NotI) | 2932 | full length |
| | R gtCGAATTGTGGCATttttggtacCAGGGAATATGTATAGAGTTTTGCTTGG (KpnI) | | |
| *nhx-2* | F atgcggccgCAGTTTCAGCTTTGTGGCGAC (NotI) | 1978 | >90% |
| | R ctggtaccGATTTAATCACTGAAAATTATTTC (KpnI) | | |
| *unc-17* | F GAGCCGGCTGTAGCATgcggccgcTACACCAATCATTTCTCCCCT (NotI) | 3219 | >90% |
| | R gtCGAATTGTGGCATttttggtacCAAGAGATGCGGAAAATAGAAAGAC (KpnI) | | |
| *unc-122* | F actgcggccGCCTTTTGTAATGTTTTCCCGC (NotI) | 807 | full length |
| | R gtctgtacATTGTGAGCCCAATGAAGTAAAATTTC (BsrGI) | | |

* gpa-14a promoter was cloned by combining a 2.4 kb NotI/ClaI restriction fragment from pNP259 with a 0.6 kb ClaI/KpnI PCR fragment; inserted into NotI/KpnI of pBN338

Sequence data are available from the authors

**Supplementary Table S3 Plasmids used in this study**

| Plasmid | Promoter or gene | Use | Reference |
|---|---|---|---|
| pBN338 | dat-1p::FLP::SL2::mNG | MosSCI cxTi10882 IV | Muñoz-Jiménez et al. 2017 |
| pBN548 | dat-1p::FLP::SL2::mNG | MosSCI oxTi365 V | this study |
| pBN397 | ckb-3p::FLP::SL2::mNG | MosSCI cxTi10882 IV | this study |
| pBN527 | gpa-14p::FLP::SL2::mNG | MosSCI cxTi10882 IV | this study |
| pBN523 | hlh-12p::FLP::SL2::mNG | MosSCI cxTi10882 IV | this study |
| pBN532 | hlh-12p::FLP::SL2::mTagBFP2 | MosSCI cxTi10882 IV | this study |
| pBN549 | hlh-12p::FLP::SL2::mTagBFP2 | MosSCI oxTi365 V | this study |
| pBN336 | lin-31p::FLP::SL2::mNG | MosSCI cxTi10882 IV | this study |
| pBN472 | nhx-2p::FLP::SL2::mNG | MosSCI cxTi10882 IV | this study |
| pBN515 | unc-17p::FLP::SL2::mNG | MosSCI cxTi10882 IV | this study |
| pBN487 | unc-122p::FLP::SL2::mNG | MosSCI cxTi10882 IV | this study |
| pCFJ601 | eft-3p::Mos1 | MosSCI | Frøkjær-Jensen et al. 2012 |
| pBN8 | MosSCI destination vector | MosSCI | Rodenas et al. 2012 |
| pBN1 | lmn-1p::mCh::his-58 | Injection marker | Rodenas et al. 2012 |
| pCFJ90 | myo-2p::mCh | Injection marker | Frøkjær-Jensen et al. 2008 |
| pCFJ104 | myo-3p::mCh | Injection marker | Frøkjær-Jensen et al. 2008 |
| #1397 | mel-28p::gfp::mel-28 | PCR template | Goméz-Saldivar et al. 2016 |
| #1286 | eft-3p::Cas9 | CRISPR/Cas9 | Friedland et al. 2013 |
| pMLS252 | gfp & unc-119(+) | SapTrap donor vector | Schwartz & Jorgensen 2016 |
| pMLS328 | Cre | SapTrap Cre expression vector | Schwartz & Jorgensen 2016 |
| pMLS256 | SapTrap destination vector | SapTrap cloning | Schwartz & Jorgensen 2016 |
| pMLS288 | N-terminal connector | SapTrap donor vector | Schwartz & Jorgensen 2016 |
| pMLS291 | mCh & unc-119(+) | SapTrap donor vector | Schwartz & Jorgensen 2016 |
| pBN312 | g>f>p & unc-119(+) | SapTrap donor vector | Muñoz-Jiménez et al. 2017 |
| pBN477 | gf>p & unc-119(+) | SapTrap donor vector | this study |
| pBN309 | mCh::baf-1 | SapTrap targeting vector (pMLS256+pMLS288+pMLS291+B961/ B962+B963/B964+B965/B966) | this study |
| pBN493 | gf>p::baf-1 | SapTrap targeting vector (pMLS256+pMLS288+pBN477+B961/B 962+B963/B964+B965/B966) | this study |
| pBN351 | g>f>p::mel-28 | SapTrap targeting vector (see details in Materials and Methods) | this study |
| pBN433 | g>f>p::npp-2 | SapTrap targeting vector (pMLS256+pMLS288+pBN312+B1231/ B1232+B1233/B1234+B1235/B1236) | this study |
| pBN209 | hsp16.41p>mCh::his-58>dam::emr-1 | DamID; MosSCI ttTi5605 II | Cabianca et al. 2019 |
| pBN488 | hsp16.41p>mCh::his-58>vhhGFP4::zif-1 | GFPdeg; MosSCI ttTi5605 II | this study |
| pBN537 | hsp16.41p>mCh::his-58>dam::rpb-6 | RAPID; MosSCI ttTi5605 II | this study |

">" denotes an FRT site

**Supplementary Table S4 Strains used in this study**

| Strain | Promoter or gene | Genotype | Reference |
|---|---|---|---|
| BN578 | Fluorescent markers II & IV | bqSi189[pBN13(unc-119(+) lmn-1p::mCherry::his-58)] II; bqSi577[pBN306(unc- 119(+) myo-2p::GFP)] IV | Muñoz-Jiménez et al. 2017 |
| BN596 | Dual color reporter | bqSi294[pBN154(unc-119(+) hsp16.41p>mCh::his-58>gfp::his-58)] II; bqSi577[pBN306(unc-119(+) myo-2p::GFP)] IV | Macías-León & Askjaer 2018 |
| EG4322 | MosSCI host strain | ttTi5605 II; unc-119(ed9) III | Frøkjær-Jensen et al. 2008 |
| EG6700 | MosSCI host strain | unc-119(ed3) III; cxTi10882 IV; oxEx1579 | Frøkjær-Jensen et al. 2012 |
| EG8082 | MosSCI host strain | unc-119(ed3) III; oxTi365 V; oxEx1580 | Frøkjær-Jensen et al. 2014 |
| HT1593 | CRISPR/Cas9 injection strain | unc-119(ed3) III | CGC |
| BN852 | *ckb-3* | unc-119(ed3) III; bqSi852[pBN397(unc-119(+) ckb-3p::FLP::SL2::mNG)] IV | this study |
| BN853 | *ckb-3* | bqSi189[pBN13(unc-119(+) lmn-1p::mCherry::his-58)] II; bqSi852[pBN397(unc-119(+) ckb-3p::FLP::SL2::mNG)] IV | this study |
| BN854 | *ckb-3* | bqSi294[pBN154(unc-119(+) hsp16.41p>mCh::his-58>gfp::his-58)] II; bqSi852[pBN397(unc-119(+) ckb-3p::FLP::SL2::mNG)] IV | this study |
| BN1381 | *dat-1* | unc-119(ed3) III; bqSi1381[pBN548(unc-119(+) dat-1p::FLP::SL2::mNG)] V | this study |
| BN1384 | *dat-1* | bqSi189[pBN13(unc-119(+) lmn-1p::mCherry::his-58)] II; bqSi1381[pBN548(unc-119(+) dat-1p::FLP::SL2::mNG)] V | this study |
| BN1385 | *dat-1* | bqSi294[pBN154(unc-119(+) hsp16.41p>mCh::his-58>gfp::his-58)] II; bqSi1381[pBN548(unc-119(+) dat-1p::FLP::SL2::mNG)] V | this study |
| BN1205 | *gpa-14* | unc-119(ed3) III; bqSi1205[pBN527(unc-119(+) gpa-14p::FLP::SL2::mNG)] IV | this study |
| BN1207 | *gpa-14* | bqSi189[pBN13(unc-119(+) lmn-1p::mCherry::his-58)] II; bqSi1205[pBN527(unc-119(+) gpa-14p::FLP::SL2::mNG)] IV | this study |
| BN1208 | *gpa-14* | bqSi294[pBN154(unc-119(+) hsp16.41p>mCh::his-58>gfp::his-58)] II; bqSi1205[pBN527(unc-119(+) gpa-14p::FLP::SL2::mNG)] IV | this study |
| BN1201 | *hlh-12* | unc-119(ed3) III; bqSi1201[pBN523(unc-119(+) hlh-12p::FLP::SL2::mNG)] IV | this study |
| BN1203 | *hlh-12* | bqSi189[pBN13(unc-119(+) lmn-1p::mCherry::his-58)] II; bqSi1201[pBN523(unc-119(+) hlh-12p::FLP::SL2::mNG)] IV | this study |
| BN1204 | *hlh-12* | bqSi294[pBN154(unc-119(+) hsp16.41p>mCh::his-58>gfp::his-58)] II; bqSi1201[pBN523(unc-119(+) hlh-12p::FLP::SL2::mNG)] IV | this study |
| BN1308 | *hlh-12* | unc-119(ed3) III; bqSi1308[pBN532(unc-119(+) hlh-12p::FLP::SL2::mTagBFP2)] IV | this study |
| BN1318 | *hlh-12* | bqSi189[pBN13(unc-119(+) lmn-1p::mCherry::his-58)] II; bqSi1308[pBN532(unc-119(+) hlh-12p::FLP::SL2::mTagBFP2)] IV | this study |
| BN1319 | *hlh-12* | bqSi294[pBN154(unc-119(+) hsp16.41p>mCh::his-58>gfp::his-58)] II; bqSi1308[pBN532(unc-119(+) hlh-12p::FLP::SL2::mTagBFP2)] IV | this study |
| BN1405 | *hlh-12* | unc-119(ed3) III; bqSi1405[pBN549(unc-119(+) hlh-12p::FLP::SL2::mTagBFP2)] V | this study |
| BN1398 | *hlh-12* | bqSi189[pBN13(unc-119(+) lmn-1p::mCherry::his-58)] II; bqSi1405[pBN549(unc-119(+) hlh-12p::FLP::SL2::mTagBFP2)] V | this study |
| BN1399 | *hlh-12* | bqSi294[pBN154(unc-119(+) hsp16.41p>mCh::his-58>gfp::his-58)] II; bqSi1405[pBN549(unc-119(+) hlh-12p::FLP::SL2::mTagBFP2)] V | this study |
| BN1021 | *lin-31* | unc-119(ed3) III; bqSi1021[pBN336(unc-119(+) lin-31p::FLP::SL2::mNG)] IV | this study |

| | | | |
|---|---|---|---|
| BN1022 | *lin-31* | bqSi189[pBN13(unc-119(+) lmn-1p::mCherry::his-58)] II; bqSi1021[pBN336(unc-119(+) lin-31p::FLP::SL2::mNG)] IV | this study |
| BN1023 | *lin-31* | bqSi294[pBN154(unc-119(+) hsp16.41p>mCh::his-58>gfp::his-58)] II; bqSi1021[pBN336(unc-119(+) lin-31p::FLP::SL2::mNG)] IV | this study |
| BN997 | *nhx-2* | unc-119(ed3) III; bqSi997[pBN472(unc-119(+) nhx-2p::FLP::SL2::mNG)] IV | this study |
| BN998 | *nhx-2* | bqSi189[pBN13(unc-119(+) lmn-1p::mCherry::his-58)] II; bqSi997[pBN472(unc-119(+) nhx-2p::FLP::SL2::mNG)] IV | this study |
| BN999 | *nhx-2* | bqSi294[pBN154(unc-119(+) hsp16.41p>mCh::his-58>gfp::his-58)] II; bqSi997[pBN472(unc-119(+) nhx-2p::FLP::SL2::mNG)] IV | this study |
| BN1123 | *unc-17* | unc-119(ed3) III; bqSi1123[pBN515(unc-119(+) unc-17p::FLP::SL2::mNG)] IV. | this study |
| BN1124 | *unc-17* | bqSi189[pBN13(unc-119(+) lmn-1p::mCherry::his-58)] II; bqSi1123[pBN515(unc-119(+) unc-17p::FLP::SL2::mNG)] IV | this study |
| BN1125 | *unc-17* | bqSi294[pBN154(unc-119(+) hsp16.41p>mCh::his-58>gfp::his-58)] II; bqSi1123[pBN515(unc-119(+) unc-17p::FLP::SL2::mNG)] IV | this study |
| OH15568 | *unc-17* | unc-17(ot907[unc-17::mKate2::3xflag]) IV | Pereira et al. 2019 |
| BN1132 | *unc-17* | unc-17(ot907[unc-17::mKate2::3xflag]) bqSi1123[pBN515(unc-119(+) unc-17p::FLP::SL2::mNG)] IV | this study |
| BN1133 | *unc-17* | bqSi294[pBN154(unc-119(+) hsp16.41p>mCh::his-58>gfp::his-58)] II; unc-17(ot907[unc-17::mKate2::3xflag]) bqSi1123[pBN515(unc-119(+) unc-17p::FLP::SL2::mNG)] IV | this study |
| BN1027 | *unc-122* | unc-119(ed3) III; bqSi1027[pBN487(unc-119(+) unc-122p::FLP::SL2::mNG)] IV | this study |
| BN1028 | *unc-122* | bqSi189[pBN13(unc-119(+) lmn-1p::mCherry::his-58)] II; bqSi1027[pBN487(unc-119(+) unc-122p::FLP::SL2::mNG)] IV | this study |
| BN1029 | *unc-122* | bqSi294[pBN154(unc-119(+) hsp16.41p>mCh::his-58>gfp::his-58)] II; bqSi1027[pBN487(unc-119(+) unc-122p::FLP::SL2::mNG)] IV | this study |
| BN580 | *baf-1* | baf-1(bq12[g>f>p::baf-1]) III | Muñoz-Jiménez et al. 2017 |
| BN581 | *baf-1* | baf-1(bq13[mCh::baf-1]) III | this study |
| BN1164 | *baf-1* | baf-1(bq47[gf>p::baf-1]) III | this study |
| BN1215 | *baf-1* | baf-1(bq52[gf>p::baf-1>]) III | this study |
| BN1182 | *baf-1* | baf-1(bq13[mCh::baf-1])/baf-1(bq12[g>f>p::baf-1]) III; bqSi997[pBN472(unc-119(+) nhx-2p::FLP::SL2::mNG)] IV | this study |
| BN1223 | *baf-1* | baf-1(bq13[mCh::baf-1])/baf-1(bq52[gf>p::baf-1>]) III; bqSi997[pBN472(unc-119(+) nhx-2p::FLP::SL2::mNG)] IV | this study |
| BN1097 | GFPdeg hypodermis | bqSi1030[pBN488(unc-119(+) hsp16.41p>mCh::his-58>vhhGFP4::zif-1)] II; bqSi548[pBN266(unc-119(+) dpy-7p::FLP)] IV | this study |
| BN1117 | GFPdeg intestine | bqSi1030[pBN488(unc-119(+) hsp16.41p>mCh::his-58>vhhGFP4::zif-1)] II; bqSi508[pBN282(unc-119(+) elt-2p::FLP)] IV | this study |
| OD2768 | GFPdeg intestine | ltSi910[pOD2044/pSW378; elt-2p::vhhGFP4::ZIF-1::SL2::mCh::his; unc-119(+)]II | Wang et al. 2017 |
| BN452 | *mel-28* | bqSi189[pBN13(unc-119(+) lmn-1p::mCherry::his-58)] II; mel-28(bq5[gfp::mel-28]) III | Gómez-Saldivar et al. 2016 |
| BN902 | *mel-28* | mel-28(bq17[g>f>p::mel-28]) III | this study |
| BN793 | *mel-28* | bqSi189[pBN13(unc-119(+) lmn-1p::mCherry::his-58)] II; mel-28(bq17[g>f>p::mel-28]) III | this study |

| | | | |
|---|---|---|---|
| BN1121 | *mel-28* | bqSi189[pBN13(unc-119(+) lmn-1p::mCherry::his-58)] II; mel-28(bq17[g>f>p::mel-28]) unc-119(ed3) III; bqSi548[pBN266(unc-119(+) dpy-7p::FLP)] IV | this study |
| BN1122 | *mel-28* | bqSi1030[pBN488(unc-119(+) hsp16.41p>mCh::his-58>vhhGFP4::zif-1)] II; mel-28(bq17[g>f>p::mel-28]) unc-119(ed3) III; bqSi548[pBN266(unc-119(+) dpy-7p::FLP)] IV | this study |
| BN1084 | *mel-28* | bqSi1030[pBN488(unc-119(+) hsp16.41p>mCh::his-58>vhhGFP4::zif-1)] II; mel-28(bq5[gfp::mel-28]) III; bqSi548[pBN266(unc-119(+) dpy-7p::FLP)] IV | this study |
| BN1118 | *mel-28* | bqSi1030[pBN488(unc-119(+) hsp16.41p>mCh::his-58>vhhGFP4::zif-1)] II; mel-28(bq5[gfp::mel-28]) III; bqSi508[pBN282(unc-119(+) elt-2p::FLP)] IV | this study |
| BN746 | *mel-28* | ltSi910[pOD2044/pSW378; elt-2p::vhhGFP4::ZIF-1::SL2::mCh::his; unc-119(+)]II; mel-28(bq5[gfp::mel-28]) III | this study |
| BN1044 | *npp-2* | npp-2(bq38[g>f>p::npp-2]) I | this study |
| BN1082 | *npp-2* | npp-2(bq38[g>f>p::npp-2]) I; bqSi189[pBN13(unc-119(+) lmn-1p::mCherry::his-58)] II | this study |
| BN1111 | *npp-2* | npp-2(bq38[g>f>p::npp-2]) I; bqSi189[pBN13(unc-119(+) lmn-1p::mCherry::his-58)] II; bqSi548[pBN266(unc-119(+) dpy-7p::FLP)] IV | this study |
| BN1086 | *npp-2* | npp-2(bq38[g>f>p::npp-2]) I; bqSi1030[pBN488(unc-119(+) hsp16.41p>mCh::his-58>vhhGFP4::zif-1)] II; bqSi548[pBN266(unc-119(+) dpy-7p::FLP)] IV | this study |
| BN561 | RAPID GFP::Dam hypodermis | bqSi447[pBN181(unc-119(+) hsp16.41p>mCh::his-58>gfp::dam)] II; bqSi548[pBN266(unc-119(+) dpy-7p::FLP)] IV | this study |
| BN1414 | RAPID Dam::RPB-6 hypodermis | bqSi1411[pBN537(unc-119(+) hsp16.41p>mCh::his-58>dam::rpb-6)] II; bqSi548[pBN266(unc-119(+) dpy-7p::FLP)] IV | this study |

Note that crossed FLP strains may still carry unc-119(ed3) or unc-119(ed9) III.

">" denotes an FRT site

**Supplementary Table S5 Protein coding genes expressed in hypodermis determined by RAPID**

| WormBase Gene ID | Gene Name | Sequence Name | Score | GATCs (#) | FDR |
|---|---|---|---|---|---|
| WBGene00000004 | aat-3 | F52H2.2 | 0.6568508 | 18 | 8.97E-08 |
| WBGene00000006 | aat-5 | C55C2.5 | 0.14958567 | 12 | 4.50E-02 |
| WBGene00000017 | abf-6 | T22H6.7 | 0.4211109 | 4 | 1.78E-02 |
| WBGene00000041 | aco-2 | F54H12.1 | 0.53724818 | 20 | 5.00E-07 |
| WBGene00000044 | acr-5 | K03F8.2 | 0.19681885 | 9 | 3.44E-02 |
| WBGene00000057 | acr-18 | F28F8.1 | 0.51274259 | 27 | 1.74E-08 |
| WBGene00000074 | adm-2 | C04A11.4 | 0.1737565 | 27 | 3.24E-03 |
| WBGene00000075 | adm-4 | ZK154.7 | 0.52121724 | 15 | 1.43E-05 |
| WBGene00000080 | adr-2 | T20H4.4 | 0.26465595 | 11 | 8.32E-03 |
| WBGene00000082 | adt-1 | C02B4.1 | 0.19241558 | 30 | 9.92E-04 |
| WBGene00000083 | adt-2 | F08C6.1 | 0.33022598 | 32 | 2.14E-06 |
| WBGene00000084 | aex-1 | D2030.10 | 0.10877002 | 21 | 4.94E-02 |
| WBGene00000093 | agt-1 | Y62E10A.5 | 0.22581463 | 9 | 2.35E-02 |
| WBGene00000100 | ajm-1 | C25A11.4 | 0.29477645 | 75 | 2.35E-11 |
| WBGene00000107 | alh-1 | F54D8.3 | 0.2709541 | 17 | 1.49E-03 |
| WBGene00000112 | alh-6 | F56D12.1 | 0.21607035 | 10 | 2.18E-02 |
| WBGene00000114 | alh-8 | F13D12.4 | 0.60470093 | 11 | 4.02E-05 |
| WBGene00000115 | alh-9 | F01F1.6 | 1.05489119 | 13 | 2.80E-09 |
| WBGene00000122 | aly-3 | M18.7 | 0.20259616 | 7 | 4.60E-02 |
| WBGene00000146 | ape-1 | F46F3.4 | 0.48981435 | 34 | 8.71E-10 |
| WBGene00000149 | apl-1 | C42D8.8 | 0.33563814 | 19 | 1.64E-04 |
| WBGene00000161 | apa-2 | T20B5.1 | 0.55270991 | 14 | 1.40E-05 |
| WBGene00000165 | aps-3 | Y48G8AL.14 | 0.62002215 | 6 | 1.09E-03 |
| WBGene00000175 | aqp-7 | M02F4.8 | 0.78470849 | 15 | 6.12E-08 |
| WBGene00000221 | atf-4 | T04C10.4 | 0.50058324 | 6 | 3.36E-03 |
| WBGene00000223 | atf-7 | C07G2.2 | 0.41853971 | 28 | 3.22E-07 |
| WBGene00000225 | atgp-2 | C38C6.2 | 0.39079125 | 11 | 1.15E-03 |
| WBGene00000229 | atp-2 | C34E10.6 | 0.63457292 | 12 | 1.22E-05 |
| WBGene00000233 | avr-15 | R11G10.1 | 0.41904425 | 49 | 2.18E-11 |
| WBGene00000238 | bar-1 | C54D1.6 | 0.29958208 | 14 | 1.92E-03 |
| WBGene00000240 | pah-1 | K08F8.4 | 0.42043884 | 25 | 1.19E-06 |
| WBGene00000246 | bcc-1 | M7.3 | 0.29775966 | 30 | 1.54E-05 |
| WBGene00000253 | bli-3 | F56C11.1 | 0.17502814 | 25 | 4.17E-03 |
| WBGene00000254 | bli-4 | K04F10.4 | 0.21400197 | 55 | 2.94E-06 |
| WBGene00000256 | bli-6 | Y73B6BL.34 | 0.30809529 | 6 | 2.05E-02 |
| WBGene00000262 | bra-1 | F54B11.6 | 0.31554891 | 6 | 1.91E-02 |
| WBGene00000265 | brd-1 | K04C2.4 | 0.78180073 | 13 | 4.03E-07 |
| WBGene00000273 | brp-1 | Y79H2A.1 | 0.71932336 | 18 | 1.94E-08 |
| WBGene00000282 | cah-4 | R01E6.3 | 0.55925953 | 9 | 2.90E-04 |
| WBGene00000285 | cal-1 | C13C12.1 | 0.18835505 | 9 | 3.84E-02 |
| WBGene00000292 | cap-1 | D2024.6 | 0.40240288 | 9 | 2.29E-03 |
| WBGene00000293 | cap-2 | M106.5 | 0.18131902 | 13 | 2.24E-02 |
| WBGene00000295 | cat-1 | W01C8.6 | 0.46854488 | 16 | 2.54E-05 |
| WBGene00000298 | cat-4 | F32G8.6 | 0.16109248 | 10 | 4.82E-02 |
| WBGene00000366 | cbp-1 | R10E11.1 | 0.17169232 | 36 | 9.41E-04 |

| | | | | | |
|---|---|---|---|---|---|
| WBGene00000377 | cct-1 | T05C12.7 | 0.59526923 | 10 | 9.17E-05 |
| WBGene00000379 | cct-4 | K01C8.10 | 0.16945985 | 12 | 3.21E-02 |
| WBGene00000381 | cct-6 | F01F1.8 | 0.36111169 | 14 | 5.80E-04 |
| WBGene00000383 | cdc-14 | C17G10.4 | 0.33498154 | 11 | 2.76E-03 |
| WBGene00000399 | cdh-7 | R05H10.6 | 0.19827992 | 35 | 3.22E-04 |
| WBGene00000401 | cdh-9 | F59C12.1 | 0.16175436 | 25 | 6.49E-03 |
| WBGene00000402 | cdh-10 | C45G7.5 | 0.10135272 | 60 | 6.56E-03 |
| WBGene00000407 | cdk-5 | T27E9.3 | 0.4580441 | 7 | 3.03E-03 |
| WBGene00000412 | cdr-1 | F35E8.11 | 0.3351151 | 6 | 1.59E-02 |
| WBGene00000420 | ced-6 | F56D2.7 | 0.20029156 | 17 | 7.69E-03 |
| WBGene00000421 | ced-7 | C48B4.4 | 0.1064204 | 37 | 1.91E-02 |
| WBGene00000422 | ced-8 | F08F1.5 | 0.27828934 | 7 | 2.05E-02 |
| WBGene00000463 | ceh-43 | C28A5.4 | 0.41367639 | 14 | 2.09E-04 |
| WBGene00000482 | chd-3 | T14G8.1 | 0.20078756 | 29 | 8.55E-04 |
| WBGene00000509 | cka-1 | C28D4.2 | 0.2095334 | 20 | 3.48E-03 |
| WBGene00000512 | ckb-2 | B0285.9 | 0.29254893 | 7 | 1.77E-02 |
| WBGene00000526 | clc-5 | C01C10.4 | 0.25528986 | 5 | 4.33E-02 |
| WBGene00000528 | clh-1 | T27D12.2 | 0.48102478 | 31 | 6.40E-09 |
| WBGene00000542 | clp-1 | C06G4.2 | 0.22816147 | 27 | 4.62E-04 |
| WBGene00000548 | clr-1 | F56D1.4 | 0.36646766 | 32 | 4.65E-07 |
| WBGene00000552 | cmd-1 | T21H3.3 | 0.29654628 | 6 | 2.29E-02 |
| WBGene00000556 | cnc-2 | R09B5.3 | 0.78147798 | 2 | 9.19E-03 |
| WBGene00000559 | cnc-5 | R09B5.10 | 0.44371797 | 2 | 4.03E-02 |
| WBGene00000563 | cng-3 | F38E11.12 | 0.43060927 | 9 | 1.58E-03 |
| WBGene00000585 | cogc-2 | C06G3.10 | 0.28397673 | 11 | 6.15E-03 |
| WBGene00000594 | col-3 | T28C6.6 | 0.34857314 | 5 | 2.03E-02 |
| WBGene00000599 | col-10 | B0222.8 | 0.30704233 | 6 | 2.07E-02 |
| WBGene00000601 | col-12 | F15H10.1 | 0.16684149 | 13 | 2.92E-02 |
| WBGene00000602 | col-13 | F15H10.2 | 0.34552376 | 13 | 1.13E-03 |
| WBGene00000603 | col-14 | C46A5.3 | 0.45353166 | 6 | 5.23E-03 |
| WBGene00000606 | col-17 | F11G11.10 | 1.44009018 | 8 | 1.51E-08 |
| WBGene00000616 | col-39 | C09G5.4 | 1.83647062 | 5 | 1.11E-07 |
| WBGene00000618 | col-41 | T10B10.1 | 0.7976021 | 4 | 1.33E-03 |
| WBGene00000625 | col-48 | Y54E10BL.2 | 1.00669103 | 9 | 8.01E-07 |
| WBGene00000626 | col-49 | K09H9.3 | 0.58902585 | 6 | 1.46E-03 |
| WBGene00000631 | col-54 | F33D11.3 | 0.49981653 | 8 | 1.11E-03 |
| WBGene00000634 | col-58 | F26B1.4 | 0.38620916 | 6 | 9.85E-03 |
| WBGene00000647 | col-71 | Y49F6B.10 | 0.20620666 | 8 | 3.67E-02 |
| WBGene00000648 | col-72 | W09G10.1 | 0.1707333 | 12 | 3.14E-02 |
| WBGene00000649 | col-73 | F11G11.12 | 0.64176156 | 5 | 1.86E-03 |
| WBGene00000654 | col-78 | W07A12.5 | 0.39132369 | 9 | 2.65E-03 |
| WBGene00000655 | col-79 | C09G5.3 | 1.08793258 | 4 | 1.80E-04 |
| WBGene00000656 | col-80 | C09G5.5 | 0.33736215 | 3 | 4.51E-02 |
| WBGene00000657 | col-81 | F38A3.1 | 0.20804851 | 10 | 2.45E-02 |
| WBGene00000664 | col-89 | F17C8.2 | 0.45939758 | 5 | 8.22E-03 |
| WBGene00000665 | col-90 | C29E4.1 | 1.63304577 | 5 | 5.80E-07 |
| WBGene00000667 | col-92 | W05B2.6 | 0.39310214 | 11 | 1.11E-03 |
| WBGene00000668 | col-93 | W05B2.5 | 0.29745284 | 9 | 9.13E-03 |

| | | | | | |
|---|---|---|---|---|---|
| WBGene00000669 | col-94 | W05B2.1 | 0.46204153 | 10 | 6.27E-04 |
| WBGene00000671 | col-96 | Y41C4A.19 | 0.29001182 | 8 | 1.35E-02 |
| WBGene00000672 | col-97 | ZK1010.7 | 0.5500536 | 8 | 6.10E-04 |
| WBGene00000673 | col-98 | F14F7.1 | 0.83402066 | 4 | 1.03E-03 |
| WBGene00000674 | col-99 | F29C4.8 | 0.2883668 | 31 | 1.66E-05 |
| WBGene00000677 | col-103 | F56B3.1 | 1.15234882 | 6 | 7.33E-06 |
| WBGene00000680 | col-106 | Y77E11A.15 | 0.57373878 | 7 | 8.82E-04 |
| WBGene00000681 | col-107 | C34H4.4 | 0.59485376 | 5 | 2.73E-03 |
| WBGene00000685 | col-111 | F29B9.9 | 0.38559672 | 7 | 6.55E-03 |
| WBGene00000691 | col-117 | T28C6.4 | 0.42061313 | 5 | 1.13E-02 |
| WBGene00000696 | col-122 | T05A1.2 | 0.2612675 | 7 | 2.46E-02 |
| WBGene00000699 | col-125 | C29F4.1 | 0.57891761 | 4 | 5.99E-03 |
| WBGene00000702 | col-128 | F12F6.9 | 0.6956624 | 7 | 2.40E-04 |
| WBGene00000707 | col-133 | F52B11.4 | 0.8223426 | 4 | 1.12E-03 |
| WBGene00000712 | col-139 | F41F3.4 | 0.22332728 | 9 | 2.42E-02 |
| WBGene00000715 | col-142 | T15B7.4 | 0.232852 | 10 | 1.71E-02 |
| WBGene00000717 | col-144 | B0222.6 | 0.49600471 | 7 | 2.02E-03 |
| WBGene00000718 | col-145 | B0222.7 | 0.80147541 | 6 | 1.99E-04 |
| WBGene00000722 | col-149 | B0024.1 | 0.54518412 | 7 | 1.20E-03 |
| WBGene00000726 | col-153 | F17C11.3 | 0.27473529 | 10 | 9.35E-03 |
| WBGene00000732 | col-159 | F57B1.3 | 0.76964977 | 25 | 1.06E-11 |
| WBGene00000733 | col-160 | F57B1.4 | 1.2609465 | 10 | 6.19E-09 |
| WBGene00000739 | col-166 | T07H6.3 | 0.98509386 | 8 | 3.42E-06 |
| WBGene00000740 | col-167 | T10E10.2 | 0.34358214 | 7 | 1.02E-02 |
| WBGene00000742 | col-169 | T10E10.5 | 0.70589361 | 7 | 2.16E-04 |
| WBGene00000743 | col-170 | T10E10.6 | 0.26231498 | 6 | 3.16E-02 |
| WBGene00000746 | col-173 | F41C6.5 | 0.18532212 | 12 | 2.45E-02 |
| WBGene00000753 | col-180 | C44C10.1 | 0.25793946 | 9 | 1.54E-02 |
| WBGene00000754 | col-181 | W03G11.1 | 1.58774734 | 7 | 1.78E-08 |
| WBGene00000758 | col-185 | H06A10.2 | 0.57645915 | 4 | 6.09E-03 |
| WBGene00000761 | coq-1 | C24A11.9 | 0.374788 | 10 | 2.21E-03 |
| WBGene00000787 | cps-6 | C41D11.8 | 0.25125339 | 8 | 2.14E-02 |
| WBGene00000795 | crn-2 | CD4.2 | 0.90249927 | 6 | 7.68E-05 |
| WBGene00000802 | crt-1 | Y38A10A.5 | 1.13359152 | 11 | 1.00E-08 |
| WBGene00000818 | csn-6 | Y67H2A.6 | 0.18391522 | 10 | 3.47E-02 |
| WBGene00000831 | ctl-2 | Y54G11A.5 | 0.15356068 | 21 | 1.39E-02 |
| WBGene00000833 | cts-1 | T20G5.2 | 0.22957186 | 17 | 3.89E-03 |
| WBGene00000835 | cuc-1 | ZK652.11 | 0.61871323 | 2 | 1.87E-02 |
| WBGene00000837 | cul-2 | ZK520.4 | 0.37863309 | 14 | 4.12E-04 |
| WBGene00000871 | cye-1 | C37A2.4 | 0.66488648 | 13 | 3.38E-06 |
| WBGene00000876 | cyl-1 | C52E4.6 | 0.24722916 | 18 | 2.03E-03 |
| WBGene00000879 | cyn-3 | Y75B12B.5 | 0.44318051 | 6 | 5.77E-03 |
| WBGene00000880 | cyn-4 | F59E10.2 | 0.27224317 | 10 | 9.70E-03 |
| WBGene00000881 | cyn-5 | F31C3.1 | 0.31828359 | 5 | 2.59E-02 |
| WBGene00000883 | cyn-7 | Y75B12B.2 | 0.62036927 | 6 | 1.09E-03 |
| WBGene00000884 | cyn-8 | D1009.2 | 0.65794291 | 10 | 3.71E-05 |
| WBGene00000887 | cyn-11 | T01B7.4 | 0.4581502 | 5 | 8.30E-03 |
| WBGene00000894 | dab-1 | M110.5 | 0.36578379 | 29 | 1.54E-06 |

| | | | | | |
|---|---|---|---|---|---|
| WBGene00000898 | daf-2 | Y55D5A.5 | 0.11174486 | 86 | 4.74E-04 |
| WBGene00000901 | daf-5 | W01G7.1 | 0.14274782 | 61 | 2.40E-04 |
| WBGene00000912 | daf-16 | R13H8.1 | 0.12627463 | 80 | 1.63E-04 |
| WBGene00000915 | hsp-90 | C47E8.5 | 1.03194835 | 16 | 1.07E-10 |
| WBGene00000928 | dao-2 | M03A1.7 | 1.31638305 | 3 | 1.82E-04 |
| WBGene00000938 | dcp-66 | C26C6.5 | 0.1487566 | 20 | 1.79E-02 |
| WBGene00000950 | deg-1 | C47C12.6 | 0.15131412 | 40 | 1.52E-03 |
| WBGene00000951 | deg-3 | K03B8.9 | 0.27036327 | 12 | 5.81E-03 |
| WBGene00000963 | dhp-1 | R06C7.3 | 0.20183945 | 17 | 7.42E-03 |
| WBGene00000976 | dhs-13 | F36H9.3 | 0.38801308 | 23 | 8.03E-06 |
| WBGene00000984 | dhs-21 | R11D1.11 | 0.41291295 | 8 | 3.12E-03 |
| WBGene00000995 | die-1 | C18D1.1 | 0.65869538 | 26 | 2.00E-10 |
| WBGene00001005 | dlc-1 | T26A5.9 | 0.68584474 | 7 | 2.67E-04 |
| WBGene00001006 | dlg-1 | C25F6.2 | 0.13444281 | 21 | 2.39E-02 |
| WBGene00001008 | dlk-1 | F33E2.2 | 0.5241267 | 27 | 1.16E-08 |
| WBGene00001016 | dna-2 | F43G6.1 | 0.16140475 | 21 | 1.12E-02 |
| WBGene00001018 | dnc-2 | C28H8.12 | 0.47856155 | 3 | 2.03E-02 |
| WBGene00001020 | dnj-2 | B0035.2 | 0.24438786 | 8 | 2.33E-02 |
| WBGene00001023 | dnj-5 | C04A2.7 | 0.5290955 | 24 | 5.78E-08 |
| WBGene00001030 | dnj-12 | F39B2.10 | 0.15190459 | 12 | 4.32E-02 |
| WBGene00001038 | dnj-20 | T15H9.7 | 0.11067476 | 28 | 2.89E-02 |
| WBGene00001063 | dpy-1 | M01E10.2 | 0.10093512 | 27 | 4.39E-02 |
| WBGene00001064 | dpy-2 | T14B4.6 | 0.80510727 | 8 | 2.92E-05 |
| WBGene00001065 | dpy-3 | EGAP7.1 | 0.64533572 | 7 | 4.11E-04 |
| WBGene00001066 | dpy-4 | Y41E3.2 | 0.7459211 | 4 | 1.90E-03 |
| WBGene00001068 | dpy-6 | F16F9.2 | 0.16254331 | 11 | 4.13E-02 |
| WBGene00001069 | dpy-7 | F46C8.6 | 0.39624998 | 3 | 3.23E-02 |
| WBGene00001070 | dpy-8 | C31H2.2 | 1.03600278 | 8 | 1.86E-06 |
| WBGene00001072 | dpy-10 | T14B4.7 | 0.95897446 | 7 | 1.45E-05 |
| WBGene00001073 | dpy-11 | F46E10.9 | 0.14614309 | 15 | 3.39E-02 |
| WBGene00001074 | dpy-13 | F30B5.1 | 1.49400875 | 7 | 4.85E-08 |
| WBGene00001077 | dpy-18 | Y47D3B.10 | 0.36833099 | 13 | 7.46E-04 |
| WBGene00001078 | dpy-19 | F22B7.10 | 0.17928102 | 14 | 1.99E-02 |
| WBGene00001080 | dpy-21 | Y59A8B.1 | 0.08786366 | 43 | 3.68E-02 |
| WBGene00001116 | dyc-1 | C33G3.1 | 0.16316836 | 45 | 4.18E-04 |
| WBGene00001131 | dys-1 | F15D3.1 | 0.20614268 | 74 | 1.42E-07 |
| WBGene00001134 | eat-3 | D2013.5 | 0.2189504 | 27 | 6.42E-04 |
| WBGene00001149 | bcat-1 | K02A4.1 | 0.47047512 | 11 | 3.30E-04 |
| WBGene00001159 | eff-1 | C26D10.5 | 0.29116584 | 11 | 5.49E-03 |
| WBGene00001163 | efn-2 | C43F9.8 | 0.17003707 | 11 | 3.67E-02 |
| WBGene00001167 | eef-2 | F25H5.4 | 0.6433277 | 24 | 1.49E-09 |
| WBGene00001168 | eef-1A.1 | F31E3.5 | 1.75302131 | 10 | 5.11E-12 |
| WBGene00001169 | eef-1A.2 | R03G5.1 | 1.34098423 | 11 | 3.88E-10 |
| WBGene00001173 | egl-4 | F55A8.2 | 0.18218861 | 87 | 1.75E-07 |
| WBGene00001177 | egl-8 | B0348.4 | 0.10681971 | 55 | 5.92E-03 |
| WBGene00001179 | egl-10 | F28C1.2 | 0.13447983 | 32 | 8.06E-03 |
| WBGene00001184 | egl-15 | F58A3.2 | 0.22210318 | 25 | 8.71E-04 |
| WBGene00001194 | egl-27 | C04A2.3 | 0.09211356 | 34 | 4.37E-02 |

| | | | | | |
|---|---|---|---|---|---|
| WBGene00001196 | egl-30 | M01D7.7 | 0.25959193 | 23 | 4.18E-04 |
| WBGene00001209 | egl-45 | C27D11.1 | 0.19966421 | 25 | 1.84E-03 |
| WBGene00001210 | egl-46 | K11G9.4 | 0.2003184 | 11 | 2.28E-02 |
| WBGene00001226 | eif-3.C | T23D8.4 | 0.12087426 | 23 | 2.98E-02 |
| WBGene00001227 | eif-3.D | R08D7.3 | 0.61950991 | 11 | 3.18E-05 |
| WBGene00001229 | eif-3.F | D2013.7 | 0.25694792 | 7 | 2.58E-02 |
| WBGene00001233 | eif-3.K | T16G1.11 | 0.31938469 | 5 | 2.57E-02 |
| WBGene00001239 | elo-1 | F56H11.4 | 0.78262846 | 5 | 5.91E-04 |
| WBGene00001240 | elo-2 | F11E6.5 | 0.26265352 | 5 | 4.08E-02 |
| WBGene00001251 | elt-3 | K02B9.4 | 0.14377779 | 17 | 2.86E-02 |
| WBGene00001303 | sec-61.G | F32D8.6 | 0.85889582 | 3 | 2.39E-03 |
| WBGene00001325 | eor-2 | C44H4.7 | 0.13776427 | 15 | 4.03E-02 |
| WBGene00001329 | epn-1 | T04C10.2 | 0.15209607 | 12 | 4.31E-02 |
| WBGene00001330 | eps-8 | Y57G11C.24 | 0.20935873 | 39 | 8.95E-05 |
| WBGene00001368 | exc-7 | F35H8.5 | 0.2539703 | 14 | 4.66E-03 |
| WBGene00001393 | fat-1 | Y67H2A.8 | 0.15808803 | 11 | 4.43E-02 |
| WBGene00001396 | fat-4 | T13F2.1 | 0.30762422 | 10 | 5.82E-03 |
| WBGene00001398 | fat-6 | VZK822L.1 | 0.36419576 | 10 | 2.57E-03 |
| WBGene00001405 | fce-1 | C04F12.10 | 0.43613872 | 5 | 9.94E-03 |
| WBGene00001412 | fem-2 | T19C3.8 | 0.29244389 | 15 | 1.64E-03 |
| WBGene00001423 | fib-1 | T01C3.7 | 1.18891514 | 12 | 1.01E-09 |
| WBGene00001428 | fkb-3 | C05C8.3 | 0.51583228 | 6 | 2.91E-03 |
| WBGene00001429 | fkb-4 | ZC455.10 | 0.26824516 | 11 | 7.87E-03 |
| WBGene00001431 | fkb-6 | F31D4.3 | 0.79437607 | 13 | 3.20E-07 |
| WBGene00001443 | fli-1 | B0523.5 | 0.45553345 | 27 | 1.35E-07 |
| WBGene00001453 | flp-10 | T06C10.4 | 0.46712604 | 5 | 7.72E-03 |
| WBGene00001478 | fmo-3 | Y39A1A.19 | 0.23563911 | 8 | 2.58E-02 |
| WBGene00001479 | fmo-4 | F53F4.5 | 0.4434489 | 8 | 2.17E-03 |
| WBGene00001486 | frh-1 | F59G1.7 | 0.24529393 | 5 | 4.70E-02 |
| WBGene00001493 | frm-7 | C51F7.1 | 0.64673515 | 20 | 2.60E-08 |
| WBGene00001497 | fars-1 | T08B2.9 | 0.66911266 | 10 | 3.16E-05 |
| WBGene00001500 | ftn-1 | C54F6.14 | 0.62821994 | 5 | 2.08E-03 |
| WBGene00001502 | ftt-2 | F52D10.3 | 0.88307687 | 5 | 2.61E-04 |
| WBGene00001510 | fzr-1 | ZK1307.6 | 0.2148705 | 16 | 6.69E-03 |
| WBGene00001514 | xnd-1 | C05D2.5 | 0.16972455 | 13 | 2.77E-02 |
| WBGene00001515 | gap-1 | T24C12.2 | 0.3205107 | 16 | 6.57E-04 |
| WBGene00001516 | gap-2 | ZK899.8 | 0.07672906 | 95 | 1.76E-02 |
| WBGene00001527 | gcs-1 | F37B12.2 | 0.22554843 | 15 | 6.55E-03 |
| WBGene00001531 | gcy-4 | ZK970.5 | 0.18663817 | 16 | 1.24E-02 |
| WBGene00001532 | gcy-5 | ZK970.6 | 0.3383242 | 20 | 1.07E-04 |
| WBGene00001537 | gcy-11 | C30G4.3 | 0.29902918 | 32 | 7.94E-06 |
| WBGene00001555 | gcy-35 | T04D3.4 | 0.53740547 | 33 | 1.91E-10 |
| WBGene00001558 | gdi-1 | Y57G11C.10 | 0.51747854 | 10 | 2.82E-04 |
| WBGene00001559 | gei-1 | F45H7.2 | 0.19031429 | 57 | 1.13E-05 |
| WBGene00001560 | gei-3 | T22H6.6 | 0.17709199 | 52 | 6.38E-05 |
| WBGene00001562 | lin-66 | B0513.1 | 0.27005566 | 29 | 6.02E-05 |
| WBGene00001564 | icl-1 | C05E4.9 | 0.35243397 | 25 | 1.14E-05 |
| WBGene00001565 | gei-8 | C14B9.6 | 0.31213865 | 34 | 2.40E-06 |

| | | | | | |
|---|---|---|---|---|---|
| WBGene00001582 | gfi-2 | K02A11.1 | 0.44293208 | 23 | 1.48E-06 |
| WBGene00001588 | lgc-56 | F09C12.1 | 0.35648239 | 11 | 1.97E-03 |
| WBGene00001592 | glc-2 | F25F8.2 | 0.39976134 | 11 | 1.00E-03 |
| WBGene00001604 | gln-3 | Y105C5B.28 | 0.46916076 | 13 | 1.19E-04 |
| WBGene00001615 | glr-4 | C06A8.9 | 0.09998917 | 36 | 2.76E-02 |
| WBGene00001623 | glt-5 | Y53C12A.2 | 0.29484834 | 13 | 2.84E-03 |
| WBGene00001631 | gly-6 | H38K22.5 | 0.29339598 | 21 | 2.68E-04 |
| WBGene00001644 | gly-19 | F22D6.12 | 0.22406871 | 7 | 3.66E-02 |
| WBGene00001649 | gob-1 | H13N06.3 | 0.26353242 | 11 | 8.47E-03 |
| WBGene00001674 | gpa-12 | F18G5.3 | 0.18687653 | 15 | 1.46E-02 |
| WBGene00001682 | gpc-2 | F08B6.2 | 0.99597657 | 5 | 1.04E-04 |
| WBGene00001683 | gpd-1 | T09F3.3 | 0.31203336 | 10 | 5.46E-03 |
| WBGene00001684 | gpd-2 | K10B3.8 | 1.18786879 | 7 | 1.27E-06 |
| WBGene00001685 | gpd-3 | K10B3.7 | 0.68268078 | 7 | 2.76E-04 |
| WBGene00001690 | grd-1 | R08B4.1 | 0.34161157 | 25 | 1.63E-05 |
| WBGene00001707 | grh-1 | Y48G8AR.1 | 0.14959371 | 34 | 3.37E-03 |
| WBGene00001712 | grl-3 | K03B8.7 | 0.871347 | 2 | 6.21E-03 |
| WBGene00001713 | grl-4 | F42C5.7 | 0.55223173 | 6 | 2.07E-03 |
| WBGene00001720 | grl-11 | ZK512.9 | 0.39198334 | 3 | 3.31E-02 |
| WBGene00001725 | grl-16 | Y65B4BR.6 | 0.49652297 | 8 | 1.15E-03 |
| WBGene00001730 | grl-21 | ZC168.5 | 0.46040779 | 4 | 1.36E-02 |
| WBGene00001733 | grl-24 | F11E6.2 | 0.42237003 | 3 | 2.79E-02 |
| WBGene00001740 | gro-1 | ZC395.6 | 0.36049333 | 11 | 1.85E-03 |
| WBGene00001744 | gars-1 | T10F2.1 | 0.25082521 | 20 | 1.14E-03 |
| WBGene00001748 | gsp-2 | F56C9.1 | 0.67764217 | 15 | 5.62E-07 |
| WBGene00001753 | gst-5 | R03D7.6 | 0.3305275 | 5 | 2.35E-02 |
| WBGene00001756 | gst-8 | F11G11.1 | 0.36313806 | 8 | 5.65E-03 |
| WBGene00001760 | gst-12 | F37B1.2 | 0.65792098 | 7 | 3.59E-04 |
| WBGene00001768 | gst-20 | Y48E1B.10 | 0.4065857 | 11 | 8.98E-04 |
| WBGene00001774 | gst-26 | Y53F4B.29 | 0.46119455 | 5 | 8.10E-03 |
| WBGene00001778 | gst-30 | ZK546.11 | 0.18153445 | 11 | 3.06E-02 |
| WBGene00001781 | gst-33 | C02A12.1 | 0.39061191 | 7 | 6.21E-03 |
| WBGene00001784 | gst-36 | R07B1.4 | 0.49643273 | 5 | 6.08E-03 |
| WBGene00001788 | gst-40 | F56B3.10 | 0.80339277 | 6 | 1.95E-04 |
| WBGene00001843 | hgo-1 | W06D4.1 | 0.67567335 | 19 | 2.60E-08 |
| WBGene00001851 | hif-1 | F38A6.3 | 0.65520815 | 16 | 4.21E-07 |
| WBGene00001854 | hil-3 | F22F1.1 | 0.46445593 | 6 | 4.72E-03 |
| WBGene00001898 | his-24 | M163.3 | 0.63405301 | 6 | 9.58E-04 |
| WBGene00001955 | hlh-11 | F58A4.7 | 0.43689189 | 26 | 4.25E-07 |
| WBGene00001981 | hnd-1 | C44C10.8 | 0.17941482 | 9 | 4.32E-02 |
| WBGene00001993 | hpd-1 | T21C12.2 | 0.66720872 | 8 | 1.51E-04 |
| WBGene00001994 | hpk-1 | F20B6.8 | 0.19731999 | 26 | 1.66E-03 |
| WBGene00001997 | hpr-9 | Y39A1A.23 | 0.30378257 | 11 | 4.50E-03 |
| WBGene00002000 | hrpr-1 | F58D5.1 | 0.44486902 | 17 | 2.62E-05 |
| WBGene00002005 | hsp-1 | F26D10.3 | 1.30518565 | 21 | 1.03E-16 |
| WBGene00002007 | hsp-3 | C15H9.6 | 0.8043912 | 20 | 3.69E-10 |
| WBGene00002008 | hsp-4 | F43E2.8 | 0.83226409 | 15 | 2.29E-08 |
| WBGene00002010 | hsp-6 | C37H5.8 | 0.38246912 | 16 | 1.68E-04 |

| | | | | | |
|---|---|---|---|---|---|
| WBGene00002024 | hsp-43 | C14F11.5 | 0.26154888 | 11 | 8.74E-03 |
| WBGene00002025 | hsp-60 | Y22D7AL.5 | 0.54878939 | 24 | 3.07E-08 |
| WBGene00002026 | hsp-70 | C12C8.1 | 0.26260148 | 11 | 8.59E-03 |
| WBGene00002037 | hum-4 | F46C3.3 | 0.11450195 | 46 | 6.66E-03 |
| WBGene00002045 | icd-1 | C56C10.8 | 1.10251994 | 6 | 1.17E-05 |
| WBGene00002048 | ida-1 | B0244.2 | 0.15898143 | 35 | 1.95E-03 |
| WBGene00002051 | ifa-3 | F52E10.5 | 0.40669039 | 16 | 9.90E-05 |
| WBGene00002053 | ifb-1 | F10C1.2 | 0.43573568 | 28 | 1.70E-07 |
| WBGene00002065 | iff-2 | F54C9.1 | 0.54241462 | 8 | 6.68E-04 |
| WBGene00002070 | ile-1 | K07A1.8 | 0.39746215 | 8 | 3.76E-03 |
| WBGene00002074 | ima-3 | F32E10.4 | 0.3015497 | 8 | 1.18E-02 |
| WBGene00002077 | imb-3 | C53D5.6 | 0.48699328 | 16 | 1.70E-05 |
| WBGene00002078 | xpo-1 | ZK742.1 | 0.12227517 | 43 | 5.38E-03 |
| WBGene00002080 | xpo-3 | C49H3.10 | 0.17359952 | 29 | 2.42E-03 |
| WBGene00002087 | ins-4 | ZK75.1 | 0.72446799 | 5 | 9.49E-04 |
| WBGene00002088 | ins-5 | ZK84.3 | 0.21373495 | 10 | 2.26E-02 |
| WBGene00002092 | ins-9 | C06E2.8 | 0.88027929 | 4 | 7.52E-04 |
| WBGene00002100 | ins-17 | F56F3.6 | 0.89863342 | 4 | 6.62E-04 |
| WBGene00002120 | ins-37 | F08G2.6 | 0.30782284 | 6 | 2.06E-02 |
| WBGene00002127 | inx-5 | R09F10.4 | 0.16397152 | 11 | 4.04E-02 |
| WBGene00002134 | inx-12 | ZK770.3 | 0.56361529 | 13 | 2.14E-05 |
| WBGene00002135 | inx-13 | Y8G1A.2 | 0.19285018 | 13 | 1.82E-02 |
| WBGene00002163 | ist-1 | C54D1.3 | 0.0965859 | 38 | 2.92E-02 |
| WBGene00002181 | kal-1 | K03D10.1 | 0.12462918 | 34 | 1.03E-02 |
| WBGene00002189 | kin-1 | ZK909.2 | 0.08743461 | 78 | 9.42E-03 |
| WBGene00002192 | kin-4 | C10C6.1 | 0.20353278 | 58 | 3.55E-06 |
| WBGene00002200 | kin-16 | M176.7 | 0.35418885 | 14 | 6.64E-04 |
| WBGene00002214 | klc-1 | M7.2 | 0.40145901 | 11 | 9.73E-04 |
| WBGene00002215 | klc-2 | C18C4.10 | 0.42183125 | 14 | 1.78E-04 |
| WBGene00002218 | klp-6 | R144.1 | 0.21322852 | 20 | 3.15E-03 |
| WBGene00002240 | ksr-2 | F58D5.4 | 0.16639282 | 25 | 5.56E-03 |
| WBGene00002258 | lbp-6 | W02D3.5 | 0.64063994 | 5 | 1.88E-03 |
| WBGene00002262 | ldh-1 | F13D12.2 | 0.26862621 | 11 | 7.82E-03 |
| WBGene00002264 | lec-1 | W09H1.6 | 0.39685398 | 19 | 3.40E-05 |
| WBGene00002265 | lec-2 | F52H3.7 | 1.1774941 | 21 | 3.81E-15 |
| WBGene00002324 | let-49 | Y54E5B.3 | 0.80683539 | 7 | 7.35E-05 |
| WBGene00002363 | let-92 | F38H4.9 | 0.2324425 | 15 | 5.68E-03 |
| WBGene00002717 | let-526 | C01G8.9 | 0.30212783 | 43 | 2.32E-07 |
| WBGene00002827 | let-653 | C29E6.1 | 0.15490051 | 18 | 1.95E-02 |
| WBGene00002845 | let-711 | F57B9.2 | 0.22673049 | 48 | 5.36E-06 |
| WBGene00002879 | let-754 | C29E4.8 | 0.43229272 | 13 | 2.33E-04 |
| WBGene00002889 | let-765 | F20H11.2 | 0.26647499 | 36 | 1.08E-05 |
| WBGene00002891 | let-767 | C56G2.6 | 0.34523726 | 7 | 1.01E-02 |
| WBGene00002915 | let-805 | H19M22.2 | 0.52251955 | 128 | 2.34E-34 |
| WBGene00002976 | lev-9 | T07H6.5 | 0.25738113 | 20 | 9.55E-04 |
| WBGene00002990 | lin-1 | C37F5.1 | 0.25817688 | 34 | 2.66E-05 |
| WBGene00003009 | lin-23 | K10B2.1 | 0.2593425 | 14 | 4.20E-03 |
| WBGene00003012 | lin-26 | F18A1.2 | 0.80475759 | 9 | 1.14E-05 |

| | | | | | |
|---|---|---|---|---|---|
| WBGene00003018 | lin-32 | T14F9.5 | 0.32418589 | 3 | 4.85E-02 |
| WBGene00003034 | lin-49 | F42A9.2 | 0.15510119 | 18 | 1.94E-02 |
| WBGene00003037 | lin-54 | JC8.6 | 0.24880597 | 17 | 2.49E-03 |
| WBGene00003040 | lin-59 | T12F5.4 | 0.21320603 | 37 | 1.09E-04 |
| WBGene00003044 | lir-1 | F18A1.3 | 0.22062906 | 32 | 2.15E-04 |
| WBGene00003053 | lmp-1 | C03B1.12 | 0.29075812 | 6 | 2.42E-02 |
| WBGene00003055 | lon-1 | F48E8.1 | 0.38497437 | 11 | 1.26E-03 |
| WBGene00003057 | lon-3 | ZK836.1 | 1.48953432 | 5 | 1.87E-06 |
| WBGene00003063 | lpd-7 | R13A5.12 | 0.21304854 | 8 | 3.38E-02 |
| WBGene00003071 | lrp-1 | F29D11.1 | 0.43229867 | 100 | 3.13E-22 |
| WBGene00003078 | lsm-4 | F32A5.7 | 0.42062795 | 4 | 1.78E-02 |
| WBGene00003081 | lsm-7 | ZK593.7 | 0.55174759 | 5 | 3.87E-03 |
| WBGene00003083 | lst-1 | T22A3.3 | 0.26151338 | 12 | 6.75E-03 |
| WBGene00003084 | lst-2 | R160.7 | 0.41803181 | 19 | 1.97E-05 |
| WBGene00003089 | ltd-1 | K02C4.4 | 0.30929721 | 19 | 3.23E-04 |
| WBGene00003112 | mab-21 | F35G12.6 | 0.28800523 | 9 | 1.03E-02 |
| WBGene00003124 | mai-1 | K10B3.9 | 0.31497531 | 7 | 1.39E-02 |
| WBGene00003130 | map-2 | Y116A8A.9 | 0.13421276 | 26 | 1.47E-02 |
| WBGene00003132 | mat-1 | Y110A7A.17 | 0.23462454 | 17 | 3.46E-03 |
| WBGene00003144 | max-2 | Y38F1A.10 | 0.10883741 | 34 | 2.08E-02 |
| WBGene00003148 | mbf-1 | H21P03.1 | 0.40067538 | 7 | 5.58E-03 |
| WBGene00003151 | mca-1 | W09C2.3 | 0.53792315 | 31 | 6.27E-10 |
| WBGene00003161 | mdf-2 | Y69A2AR.30 | 0.46037346 | 5 | 8.16E-03 |
| WBGene00003162 | mdh-2 | F20H11.3 | 0.32415898 | 7 | 1.26E-02 |
| WBGene00003163 | mdl-1 | R03E9.1 | 0.30924888 | 8 | 1.07E-02 |
| WBGene00003171 | mec-7 | ZK154.3 | 0.2699841 | 10 | 1.00E-02 |
| WBGene00003172 | mec-8 | F46A9.6 | 0.57386481 | 10 | 1.25E-04 |
| WBGene00003178 | mec-17 | F57H12.7 | 0.1962881 | 10 | 2.90E-02 |
| WBGene00003186 | mek-2 | Y54E10BL.6 | 0.19245429 | 37 | 2.98E-04 |
| WBGene00003214 | mel-32 | C05D11.11 | 0.32544057 | 24 | 3.92E-05 |
| WBGene00003218 | mep-1 | M04B2.1 | 0.13211901 | 18 | 3.40E-02 |
| WBGene00003253 | mig-22 | PAR2.4 | 0.3022156 | 9 | 8.57E-03 |
| WBGene00003367 | mix-1 | M106.1 | 0.088397 | 38 | 4.38E-02 |
| WBGene00003372 | mlc-4 | C56G7.1 | 0.33934109 | 8 | 7.51E-03 |
| WBGene00003384 | moc-1 | T06H11.4 | 0.38628018 | 6 | 9.85E-03 |
| WBGene00003394 | mom-1 | T07H6.2 | 0.15201289 | 16 | 2.66E-02 |
| WBGene00003400 | dapk-1 | K12C11.4 | 0.23566977 | 78 | 3.50E-09 |
| WBGene00003476 | mtm-3 | T24A11.1 | 0.1909773 | 27 | 1.75E-03 |
| WBGene00003482 | mua-3 | K08E5.3 | 0.07979231 | 93 | 1.29E-02 |
| WBGene00003485 | mua-6 | W10G6.3 | 1.0120301 | 24 | 1.11E-14 |
| WBGene00003497 | mup-4 | K07D8.1 | 0.74974291 | 46 | 2.31E-19 |
| WBGene00003511 | mxl-3 | F46G10.6 | 0.44743686 | 7 | 3.39E-03 |
| WBGene00003517 | nac-1 | F31F6.6 | 0.38423847 | 7 | 6.64E-03 |
| WBGene00003522 | nas-3 | K06A4.1 | 0.3233851 | 11 | 3.31E-03 |
| WBGene00003528 | nas-9 | C37H5.9 | 0.24034135 | 18 | 2.41E-03 |
| WBGene00003562 | ncr-2 | F09G8.4 | 0.09640341 | 31 | 4.21E-02 |
| WBGene00003583 | ndx-6 | EEED8.8 | 0.26817282 | 7 | 2.29E-02 |
| WBGene00003588 | nex-1 | ZC155.1 | 0.81952853 | 11 | 1.38E-06 |

| | | | | | |
|---|---|---|---|---|---|
| WBGene00003589 | nex-2 | T07C4.9 | 0.30022297 | 18 | 5.55E-04 |
| WBGene00003593 | nfm-1 | F42A10.2 | 0.1863042 | 18 | 9.03E-03 |
| WBGene00003598 | nhl-2 | F26F4.7 | 0.27627456 | 21 | 4.35E-04 |
| WBGene00003600 | nhr-1 | R09G11.2 | 0.26263536 | 54 | 1.21E-07 |
| WBGene00003603 | nhr-4 | F32B6.1 | 0.26748231 | 8 | 1.77E-02 |
| WBGene00003610 | nhr-11 | ZC410.1 | 0.29886847 | 15 | 1.43E-03 |
| WBGene00003613 | nhr-14 | T01B10.4 | 0.25899543 | 12 | 7.05E-03 |
| WBGene00003615 | nhr-16 | T12C9.6 | 0.21241583 | 7 | 4.15E-02 |
| WBGene00003616 | nhr-17 | C02B4.2 | 0.2939294 | 10 | 7.09E-03 |
| WBGene00003618 | nhr-19 | E02H1.7 | 0.10763846 | 24 | 4.19E-02 |
| WBGene00003622 | nhr-23 | C01H6.5 | 0.11200838 | 22 | 4.20E-02 |
| WBGene00003625 | nhr-31 | C26B2.3 | 0.19238865 | 27 | 1.66E-03 |
| WBGene00003626 | nhr-32 | K08H2.8 | 0.22386341 | 11 | 1.58E-02 |
| WBGene00003627 | nhr-34 | F58G6.5 | 0.20105128 | 43 | 6.58E-05 |
| WBGene00003630 | nhr-40 | T03G6.2 | 0.16906493 | 36 | 1.06E-03 |
| WBGene00003633 | nhr-43 | C29E6.5 | 0.24388192 | 16 | 3.54E-03 |
| WBGene00003634 | nhr-44 | T19A5.4 | 0.20646866 | 9 | 3.03E-02 |
| WBGene00003636 | nhr-46 | C45E5.6 | 0.19051386 | 35 | 4.59E-04 |
| WBGene00003638 | nhr-48 | ZK662.3 | 0.1256007 | 24 | 2.36E-02 |
| WBGene00003643 | nhr-53 | K06B4.11 | 0.37982962 | 6 | 1.05E-02 |
| WBGene00003654 | nhr-64 | C45E1.1 | 0.2219402 | 17 | 4.65E-03 |
| WBGene00003656 | nhr-66 | T09A12.4 | 0.15125185 | 45 | 8.38E-04 |
| WBGene00003658 | nhr-68 | H12C20.3 | 0.44927089 | 10 | 7.54E-04 |
| WBGene00003660 | nhr-70 | Y51A2D.17 | 0.33299166 | 7 | 1.15E-02 |
| WBGene00003661 | nhr-71 | K11E4.5 | 0.20855571 | 19 | 4.32E-03 |
| WBGene00003674 | nhr-84 | T06C12.7 | 0.22195217 | 12 | 1.32E-02 |
| WBGene00003675 | nhr-85 | W05B5.3 | 0.36585815 | 19 | 7.54E-05 |
| WBGene00003681 | nhr-91 | Y15E3A.1 | 0.32910878 | 12 | 2.15E-03 |
| WBGene00003687 | nhr-97 | H27C11.1 | 0.20573108 | 27 | 1.03E-03 |
| WBGene00003691 | nhr-101 | H12C20.6 | 0.23692163 | 6 | 4.01E-02 |
| WBGene00003692 | nhr-102 | T06C12.6 | 0.71661011 | 9 | 3.65E-05 |
| WBGene00003694 | nhr-104 | R11E3.5 | 0.20520026 | 9 | 3.08E-02 |
| WBGene00003699 | nhr-109 | T12C9.5 | 0.35260562 | 7 | 9.31E-03 |
| WBGene00003701 | nhr-111 | F44G3.9 | 0.31400805 | 9 | 7.34E-03 |
| WBGene00003704 | nhr-114 | Y45G5AM.1 | 0.58265947 | 19 | 2.85E-07 |
| WBGene00003705 | nhr-115 | T27B7.4 | 0.32610054 | 14 | 1.15E-03 |
| WBGene00003709 | nhr-119 | K12H6.1 | 0.20139073 | 24 | 2.08E-03 |
| WBGene00003710 | nhr-120 | C25B8.6 | 0.2233073 | 15 | 6.86E-03 |
| WBGene00003727 | nhr-137 | C56E10.4 | 0.09672579 | 28 | 4.84E-02 |
| WBGene00003728 | nhr-138 | C28D4.9 | 0.16411803 | 11 | 4.03E-02 |
| WBGene00003729 | nhx-1 | B0395.1 | 0.17789805 | 11 | 3.24E-02 |
| WBGene00003731 | nhx-3 | C54F6.13 | 0.23329843 | 11 | 1.36E-02 |
| WBGene00003736 | nhx-9 | ZK822.3 | 0.21266739 | 14 | 1.04E-02 |
| WBGene00003746 | nlp-8 | D2005.2 | 0.49800928 | 10 | 3.73E-04 |
| WBGene00003763 | nlp-25 | Y43F8C.1 | 0.50007232 | 5 | 5.90E-03 |
| WBGene00003765 | nlp-27 | B0213.2 | 1.36447439 | 3 | 1.39E-04 |
| WBGene00003767 | nlp-29 | B0213.4 | 1.26309694 | 2 | 1.12E-03 |
| WBGene00003768 | nlp-30 | B0213.5 | 0.57144974 | 6 | 1.73E-03 |

| | | | | | |
|---|---|---|---|---|---|
| WBGene00003771 | nlp-33 | T19C4.7 | 0.62572292 | 3 | 8.88E-03 |
| WBGene00003776 | nmy-1 | F52B10.1 | 0.18969191 | 35 | 4.77E-04 |
| WBGene00003785 | nos-3 | Y53C12B.3 | 0.11927847 | 20 | 3.97E-02 |
| WBGene00003793 | npp-7 | T19B4.2 | 0.35431226 | 20 | 6.98E-05 |
| WBGene00003795 | npp-9 | F59A2.1 | 0.24847355 | 15 | 4.07E-03 |
| WBGene00003818 | nsf-1 | H15N14.2 | 0.14155043 | 16 | 3.35E-02 |
| WBGene00003821 | nst-1 | K01C8.9 | 0.34075479 | 7 | 1.06E-02 |
| WBGene00003822 | nsy-1 | F59A6.1 | 0.20739517 | 29 | 6.64E-04 |
| WBGene00003831 | nuo-1 | C09H10.3 | 0.17726318 | 10 | 3.82E-02 |
| WBGene00003837 | oat-1 | T01B11.7 | 0.11252708 | 20 | 4.77E-02 |
| WBGene00003844 | odc-1 | K11C4.4 | 0.22839157 | 14 | 7.67E-03 |
| WBGene00003847 | blmp-1 | F25D7.3 | 0.58908415 | 17 | 9.21E-07 |
| WBGene00003848 | odr-1 | R01E6.1 | 0.1462644 | 20 | 1.92E-02 |
| WBGene00003858 | ogt-1 | K04G7.3 | 0.23635968 | 26 | 4.32E-04 |
| WBGene00003862 | old-1 | C08H9.5 | 0.18410054 | 11 | 2.94E-02 |
| WBGene00003865 | oma-2 | ZC513.6 | 0.44437901 | 7 | 3.50E-03 |
| WBGene00003878 | pept-3 | F56F4.5 | 1.29044481 | 20 | 7.40E-16 |
| WBGene00003883 | osm-1 | T27B1.1 | 0.11824395 | 34 | 1.36E-02 |
| WBGene00003887 | osm-7 | T05D4.4 | 0.41839802 | 12 | 4.73E-04 |
| WBGene00003889 | osm-9 | B0212.5 | 0.13612872 | 22 | 2.06E-02 |
| WBGene00003891 | osm-11 | F11C7.5 | 1.18575998 | 5 | 2.22E-05 |
| WBGene00003902 | pab-1 | Y106G6H.2 | 0.75886431 | 10 | 8.66E-06 |
| WBGene00003903 | pab-2 | F18H3.3 | 0.3622649 | 12 | 1.22E-03 |
| WBGene00003915 | pan-1 | M88.6 | 0.1843868 | 20 | 6.85E-03 |
| WBGene00003916 | par-1 | H39E23.1 | 0.20282952 | 73 | 2.34E-07 |
| WBGene00003918 | par-3 | F54E7.3 | 0.07838644 | 51 | 4.99E-02 |
| WBGene00003928 | pas-7 | ZK945.2 | 0.27895368 | 4 | 4.73E-02 |
| WBGene00003936 | pat-12 | T17H7.4 | 0.41614331 | 77 | 8.18E-17 |
| WBGene00003950 | pbs-4 | T20F5.2 | 0.32441872 | 5 | 2.47E-02 |
| WBGene00003951 | pbs-5 | K05C4.1 | 0.31344183 | 11 | 3.87E-03 |
| WBGene00003962 | pdi-1 | C14B1.1 | 1.13679292 | 10 | 3.71E-08 |
| WBGene00003963 | pdi-2 | C07A12.4 | 0.78452194 | 12 | 9.58E-07 |
| WBGene00003964 | pdi-3 | H06O01.1 | 0.19641439 | 17 | 8.41E-03 |
| WBGene00003965 | pdk-1 | H42K12.1 | 0.16319795 | 20 | 1.21E-02 |
| WBGene00003968 | peb-1 | T14F9.4 | 0.28485829 | 27 | 6.07E-05 |
| WBGene00003993 | pgl-2 | B0523.3 | 0.21201303 | 12 | 1.56E-02 |
| WBGene00003995 | pgp-1 | K08E7.9 | 0.12423277 | 25 | 2.26E-02 |
| WBGene00004004 | pgp-10 | C54D1.1 | 0.16615018 | 26 | 4.88E-03 |
| WBGene00004014 | phb-1 | Y37E3.9 | 0.90082358 | 5 | 2.26E-04 |
| WBGene00004023 | pho-4 | T16D1.2 | 0.75030539 | 5 | 7.69E-04 |
| WBGene00004033 | pkc-2 | E01H11.1 | 0.12061666 | 55 | 2.22E-03 |
| WBGene00004035 | pkd-2 | Y73F8A.1 | 0.17531013 | 19 | 1.02E-02 |
| WBGene00004038 | plc-3 | T01E8.3 | 0.18057625 | 29 | 1.85E-03 |
| WBGene00004043 | plk-2 | Y71F9B.7 | 0.20013637 | 12 | 1.91E-02 |
| WBGene00004095 | pqe-1 | F52C9.8 | 0.22931413 | 32 | 1.49E-04 |
| WBGene00004108 | pqn-18 | C34E7.1 | 0.20058131 | 36 | 2.41E-04 |
| WBGene00004109 | stam-1 | C34G6.7 | 0.69064651 | 12 | 4.70E-06 |
| WBGene00004110 | pqn-20 | C37A2.2 | 0.15339764 | 31 | 4.11E-03 |

| | | | | | |
|---|---|---|---|---|---|
| WBGene00004120 | pqn-32 | F29C12.1 | 0.2656526 | 10 | 1.07E-02 |
| WBGene00004131 | tent-5 | F55A12.9 | 0.18620055 | 36 | 4.75E-04 |
| WBGene00004132 | ifet-1 | F56F3.1 | 0.47428155 | 15 | 3.79E-05 |
| WBGene00004133 | abu-13 | F57B9.9 | 0.25922928 | 9 | 1.51E-02 |
| WBGene00004143 | pqn-59 | R119.4 | 0.79413243 | 18 | 3.11E-09 |
| WBGene00004164 | pqn-83 | Y39E4B.3 | 0.17294525 | 23 | 6.00E-03 |
| WBGene00004172 | pqn-92 | Y75B8A.27 | 0.26315799 | 10 | 1.11E-02 |
| WBGene00004175 | pqn-96 | ZK1236.6 | 0.371982 | 5 | 1.68E-02 |
| WBGene00004182 | prk-1 | C06E8.3 | 0.19412438 | 59 | 6.02E-06 |
| WBGene00004183 | prk-2 | F45H7.4 | 0.28964884 | 13 | 3.12E-03 |
| WBGene00004189 | pars-1 | T20H4.3 | 0.22591242 | 7 | 3.59E-02 |
| WBGene00004197 | prx-12 | F08B12.2 | 0.30550238 | 4 | 3.94E-02 |
| WBGene00004204 | swsn-4 | F01G4.1 | 0.19348074 | 27 | 1.60E-03 |
| WBGene00004210 | ptc-3 | Y110A2AL.8 | 0.9462131 | 30 | 1.11E-16 |
| WBGene00004216 | ptr-1 | C24B5.3 | 0.2239362 | 16 | 5.48E-03 |
| WBGene00004217 | ptr-2 | C32E8.8 | 0.40519638 | 18 | 4.25E-05 |
| WBGene00004218 | ptr-3 | C41D7.2 | 0.18339049 | 18 | 9.70E-03 |
| WBGene00004219 | ptr-4 | C45B2.7 | 0.29572063 | 21 | 2.51E-04 |
| WBGene00004222 | ptr-8 | F44F4.4 | 0.27328573 | 10 | 9.55E-03 |
| WBGene00004225 | ptr-11 | F56C11.2 | 0.13263832 | 24 | 1.88E-02 |
| WBGene00004226 | ptr-12 | K07A3.2 | 0.33623886 | 23 | 3.95E-05 |
| WBGene00004227 | ptr-13 | K07C10.1 | 0.45481072 | 17 | 2.08E-05 |
| WBGene00004228 | ptr-14 | R09H10.4 | 0.44250148 | 18 | 1.71E-05 |
| WBGene00004231 | ptr-17 | Y18D10A.7 | 0.15733162 | 12 | 3.94E-02 |
| WBGene00004232 | ptr-18 | Y38F1A.3 | 0.2003211 | 33 | 4.21E-04 |
| WBGene00004237 | ptr-23 | ZK270.1 | 0.56918202 | 21 | 1.11E-07 |
| WBGene00004245 | puf-9 | W06B11.2 | 0.19459623 | 16 | 1.04E-02 |
| WBGene00004256 | pxn-1 | ZK994.3 | 0.82428413 | 31 | 5.26E-15 |
| WBGene00004258 | pyc-1 | D2023.2 | 0.46480425 | 18 | 9.88E-06 |
| WBGene00004264 | qua-1 | T05C12.10 | 0.47668411 | 33 | 2.65E-09 |
| WBGene00004272 | rab-8 | D1037.4 | 0.64530053 | 5 | 1.81E-03 |
| WBGene00004279 | rab-21 | T01B7.3 | 0.28848989 | 6 | 2.47E-02 |
| WBGene00004300 | ram-2 | F38A3.2 | 0.60292059 | 6 | 1.28E-03 |
| WBGene00004310 | ras-1 | C44C11.1 | 0.14012872 | 15 | 3.84E-02 |
| WBGene00004312 | rba-1 | K07A1.11 | 0.3455508 | 7 | 1.00E-02 |
| WBGene00004313 | rbc-1 | F54E4.1 | 0.13611281 | 67 | 2.21E-04 |
| WBGene00004321 | rcan-1 | F54E7.7 | 0.24894747 | 6 | 3.58E-02 |
| WBGene00004326 | rde-4 | T20G5.11 | 0.32386811 | 6 | 1.77E-02 |
| WBGene00004336 | ret-1 | W06A7.3 | 0.09869111 | 64 | 6.50E-03 |
| WBGene00004358 | rhr-1 | F08F3.3 | 0.16803396 | 13 | 2.86E-02 |
| WBGene00004359 | rhr-2 | B0240.1 | 0.35102875 | 7 | 9.46E-03 |
| WBGene00004369 | rig-1 | K09E2.4 | 0.22006263 | 28 | 5.02E-04 |
| WBGene00004387 | rnp-4 | R07E5.14 | 0.49881466 | 7 | 1.96E-03 |
| WBGene00004395 | rol-3 | C16D9.2 | 0.63052099 | 37 | 1.88E-13 |
| WBGene00004397 | rol-6 | T01B7.7 | 0.45543274 | 3 | 2.32E-02 |
| WBGene00004398 | rol-8 | ZK1290.3 | 0.85380576 | 8 | 1.63E-05 |
| WBGene00004408 | rla-0 | F25H2.10 | 0.97873083 | 9 | 1.16E-06 |
| WBGene00004409 | rla-1 | Y37E3.7 | 0.9360114 | 4 | 5.12E-04 |

| | | | | | |
|---|---|---|---|---|---|
| WBGene00004410 | rla-2 | Y62E10A.1 | 0.60971046 | 2 | 1.95E-02 |
| WBGene00004412 | rpl-1 | Y71F9AL.13 | 1.41788925 | 8 | 1.97E-08 |
| WBGene00004414 | rpl-3 | F13B10.2 | 0.80211661 | 9 | 1.18E-05 |
| WBGene00004415 | rpl-4 | B0041.4 | 0.89090716 | 3 | 1.99E-03 |
| WBGene00004416 | rpl-5 | F54C9.5 | 0.67279358 | 8 | 1.41E-04 |
| WBGene00004417 | rpl-6 | R151.3 | 0.64957278 | 6 | 8.28E-04 |
| WBGene00004418 | rpl-7 | F53G12.10 | 0.42848034 | 8 | 2.60E-03 |
| WBGene00004419 | rpl-7A | Y24D9A.4 | 0.89451469 | 8 | 1.01E-05 |
| WBGene00004420 | rpl-9 | R13A5.8 | 0.77782984 | 7 | 1.00E-04 |
| WBGene00004421 | rpl-10 | F10B5.1 | 1.0915165 | 4 | 1.75E-04 |
| WBGene00004422 | rpl-11.1 | T22F3.4 | 0.29865532 | 4 | 4.13E-02 |
| WBGene00004423 | rpl-11.2 | F07D10.1 | 0.7004427 | 6 | 5.13E-04 |
| WBGene00004424 | rpl-12 | JC8.3 | 1.04304905 | 11 | 4.15E-08 |
| WBGene00004425 | rpl-13 | C32E8.2 | 0.66710992 | 4 | 3.26E-03 |
| WBGene00004426 | rpl-14 | C04F12.4 | 0.7704244 | 4 | 1.60E-03 |
| WBGene00004427 | rpl-15 | K11H12.2 | 0.26780781 | 6 | 3.00E-02 |
| WBGene00004428 | rpl-16 | M01F1.2 | 1.0938809 | 6 | 1.27E-05 |
| WBGene00004429 | rpl-17 | Y48G8AL.8 | 0.34885847 | 4 | 2.92E-02 |
| WBGene00004430 | rpl-18 | Y45F10D.12 | 0.6745853 | 5 | 1.42E-03 |
| WBGene00004431 | rpl-19 | C09D4.5 | 0.8038339 | 3 | 3.26E-03 |
| WBGene00004432 | rpl-20 | E04A4.8 | 1.63894594 | 4 | 4.04E-06 |
| WBGene00004434 | rpl-22 | C27A2.2 | 1.07033777 | 3 | 7.26E-04 |
| WBGene00004436 | rpl-24.1 | D1007.12 | 0.76152233 | 4 | 1.70E-03 |
| WBGene00004437 | rpl-24.2 | C03D6.8 | 0.8664415 | 3 | 2.29E-03 |
| WBGene00004444 | rpl-30 | Y106G6H.3 | 1.04998379 | 2 | 2.84E-03 |
| WBGene00004445 | rpl-31 | W09C5.6 | 0.75160584 | 5 | 7.61E-04 |
| WBGene00004446 | rpl-32 | T24B8.1 | 1.18539722 | 4 | 9.19E-05 |
| WBGene00004447 | rpl-33 | F10E7.7 | 1.08388862 | 4 | 1.85E-04 |
| WBGene00004449 | rpl-35 | ZK652.4 | 0.44018854 | 3 | 2.53E-02 |
| WBGene00004450 | rpl-36 | F37C12.4 | 1.04458216 | 3 | 8.40E-04 |
| WBGene00004452 | rpl-38 | C06B8.8 | 1.56679734 | 3 | 4.43E-05 |
| WBGene00004456 | rpl-43 | Y48B6A.2 | 0.90829124 | 3 | 1.81E-03 |
| WBGene00004458 | rpn-1 | T22D1.9 | 0.33113411 | 23 | 4.62E-05 |
| WBGene00004462 | rpn-6.1 | F57B9.10 | 0.16024926 | 18 | 1.71E-02 |
| WBGene00004464 | rpn-8 | R12E2.3 | 0.52493321 | 9 | 4.56E-04 |
| WBGene00004465 | rpn-9 | T06D8.8 | 0.47052215 | 8 | 1.57E-03 |
| WBGene00004469 | rps-0 | B0393.1 | 0.39238387 | 8 | 3.99E-03 |
| WBGene00004470 | rps-1 | F56F3.5 | 0.88810926 | 9 | 3.82E-06 |
| WBGene00004471 | rps-2 | C49H3.11 | 0.88411084 | 9 | 4.02E-06 |
| WBGene00004472 | rps-3 | C23G10.3 | 0.52251529 | 4 | 8.84E-03 |
| WBGene00004473 | rps-4 | Y43B11AR.4 | 0.87040516 | 2 | 6.23E-03 |
| WBGene00004474 | rps-5 | T05E11.1 | 0.91932228 | 4 | 5.74E-04 |
| WBGene00004475 | rps-6 | Y71A12B.1 | 1.18787861 | 6 | 5.25E-06 |
| WBGene00004477 | rps-8 | F42C5.8 | 0.31982589 | 3 | 4.97E-02 |
| WBGene00004480 | rps-11 | F40F11.1 | 0.76625279 | 3 | 4.03E-03 |
| WBGene00004482 | rps-13 | C16A3.9 | 1.5920075 | 6 | 1.18E-07 |
| WBGene00004483 | rps-14 | F37C12.9 | 0.6282291 | 3 | 8.76E-03 |
| WBGene00004485 | rps-16 | T01C3.6 | 0.66940566 | 3 | 6.95E-03 |

| | | | | | |
|---|---|---|---|---|---|
| WBGene00004487 | rps-18 | Y57G11C.16 | 1.49557464 | 4 | 1.08E-05 |
| WBGene00004488 | rps-19 | T05F1.3 | 0.56711935 | 6 | 1.80E-03 |
| WBGene00004489 | rps-20 | Y105E8A.16 | 0.36183206 | 4 | 2.67E-02 |
| WBGene00004490 | rps-21 | F37C12.11 | 1.25481799 | 2 | 1.16E-03 |
| WBGene00004493 | rps-24 | T07A9.11 | 0.32915646 | 4 | 3.35E-02 |
| WBGene00004494 | rps-25 | K02B2.5 | 1.02964955 | 2 | 3.10E-03 |
| WBGene00004495 | rps-26 | F39B2.6 | 0.34631184 | 3 | 4.28E-02 |
| WBGene00004499 | rps-30 | C26F1.4 | 0.79564147 | 6 | 2.10E-04 |
| WBGene00004704 | rsp-7 | D2089.1 | 0.53936931 | 26 | 1.23E-08 |
| WBGene00004736 | sca-1 | K11D9.2 | 0.31763504 | 39 | 3.63E-07 |
| WBGene00004745 | sdc-1 | F52E10.1 | 0.13092672 | 28 | 1.36E-02 |
| WBGene00004751 | sea-2 | K10G6.3 | 0.19544361 | 71 | 6.64E-07 |
| WBGene00004755 | sec-24.1 | F12F6.6 | 0.15520163 | 28 | 5.55E-03 |
| WBGene00004760 | sel-2 | F10F2.1 | 0.11180515 | 46 | 7.82E-03 |
| WBGene00004768 | sel-11 | F55A11.3 | 0.30914158 | 10 | 5.69E-03 |
| WBGene00004776 | ser-1 | F59C12.2 | 0.13592777 | 37 | 4.59E-03 |
| WBGene00004777 | ser-2 | C02D4.2 | 0.14848191 | 38 | 2.22E-03 |
| WBGene00004783 | seu-1 | Y73B6BL.5 | 0.11441139 | 23 | 3.63E-02 |
| WBGene00004826 | skr-20 | R12H7.5 | 0.36886716 | 4 | 2.55E-02 |
| WBGene00004827 | skr-21 | K08H2.1 | 0.41255801 | 5 | 1.20E-02 |
| WBGene00004854 | slt-1 | F40E10.4 | 0.13870398 | 21 | 2.12E-02 |
| WBGene00004856 | sma-2 | ZK370.2 | 0.26820512 | 29 | 6.46E-05 |
| WBGene00004859 | sma-5 | W06B3.2 | 0.16053717 | 20 | 1.30E-02 |
| WBGene00004862 | sma-9 | T05A10.1 | 0.29397153 | 49 | 6.07E-08 |
| WBGene00004882 | smg-4 | F46B6.3 | 0.28787122 | 4 | 4.45E-02 |
| WBGene00004900 | snf-1 | W03G9.1 | 0.24589165 | 13 | 6.93E-03 |
| WBGene00004902 | snf-3 | T13B5.1 | 0.32791842 | 14 | 1.11E-03 |
| WBGene00004911 | snf-12 | T25B6.7 | 0.62113732 | 17 | 4.38E-07 |
| WBGene00004929 | soc-2 | AC7.2 | 0.08048612 | 57 | 3.61E-02 |
| WBGene00004930 | sod-1 | C15F1.7 | 0.2977551 | 8 | 1.23E-02 |
| WBGene00004945 | sop-2 | C50E10.4 | 0.17563248 | 25 | 4.09E-03 |
| WBGene00004947 | sos-1 | T28F12.3 | 0.140458 | 105 | 2.68E-06 |
| WBGene00004963 | spe-9 | C17D12.6 | 0.70652063 | 5 | 1.10E-03 |
| WBGene00004969 | spe-15 | F47G6.4 | 0.09908833 | 46 | 1.67E-02 |
| WBGene00004980 | spk-1 | B0464.5 | 0.27306259 | 28 | 7.05E-05 |
| WBGene00004985 | spo-11 | T05E11.4 | 0.16721059 | 12 | 3.34E-02 |
| WBGene00004998 | spp-13 | F08F1.6 | 0.61996258 | 2 | 1.86E-02 |
| WBGene00005001 | spp-16 | F32D8.9 | 0.69053832 | 4 | 2.78E-03 |
| WBGene00005004 | spp-19 | K04A8.9 | 0.38222704 | 5 | 1.54E-02 |
| WBGene00005008 | spr-3 | C07A12.5 | 0.18969409 | 9 | 3.77E-02 |
| WBGene00005016 | sqt-1 | B0491.2 | 0.81629656 | 6 | 1.73E-04 |
| WBGene00005018 | sqt-3 | F23H12.4 | 1.23564786 | 6 | 3.35E-06 |
| WBGene00005048 | sra-22 | F28C12.7 | 0.28235155 | 5 | 3.48E-02 |
| WBGene00005078 | src-2 | F49B2.5 | 0.25852416 | 26 | 2.01E-04 |
| WBGene00005121 | srd-43 | R04D3.9 | 0.25166406 | 7 | 2.73E-02 |
| WBGene00005122 | srd-44 | F17A2.8 | 0.36901777 | 4 | 2.54E-02 |
| WBGene00005123 | srd-45 | F17A2.7 | 0.35753484 | 5 | 1.88E-02 |
| WBGene00005126 | srd-48 | F17A2.9 | 0.60239358 | 5 | 2.56E-03 |

| | | | | | |
|---|---|---|---|---|---|
| WBGene00005133 | srd-55 | K02A2.2 | 0.28953284 | 8 | 1.36E-02 |
| WBGene00005227 | srh-1 | T11F9.18 | 0.20424943 | 7 | 4.52E-02 |
| WBGene00005241 | srh-16 | F55C5.9 | 0.16129002 | 11 | 4.21E-02 |
| WBGene00005448 | srh-241 | F37B4.6 | 0.27418065 | 5 | 3.72E-02 |
| WBGene00005450 | srh-244 | C03G6.9 | 0.36434442 | 3 | 3.87E-02 |
| WBGene00005477 | srh-271 | W02H5.6 | 0.16065521 | 10 | 4.85E-02 |
| WBGene00005648 | srp-7 | F20D6.4 | 0.18166202 | 18 | 1.01E-02 |
| WBGene00005663 | sars-1 | C47E12.1 | 0.31713413 | 9 | 7.04E-03 |
| WBGene00005727 | srv-16 | Y105C5B.10 | 1.4227429 | 6 | 5.77E-07 |
| WBGene00006044 | ssp-16 | T27A3.3 | 0.26413297 | 5 | 4.03E-02 |
| WBGene00006059 | stc-1 | F54C9.2 | 0.39908875 | 8 | 3.68E-03 |
| WBGene00006064 | sto-2 | F32A6.5 | 0.24146884 | 36 | 3.52E-05 |
| WBGene00006066 | sto-4 | Y71H9A.3 | 0.4127974 | 9 | 2.00E-03 |
| WBGene00006072 | str-4 | C50B6.10 | 0.23595728 | 7 | 3.23E-02 |
| WBGene00006097 | str-31 | C54F6.10 | 0.52094832 | 5 | 4.98E-03 |
| WBGene00006106 | str-41 | R13D7.1 | 0.24576577 | 6 | 3.69E-02 |
| WBGene00006111 | str-46 | C31B8.6 | 0.85054051 | 5 | 3.40E-04 |
| WBGene00006121 | str-56 | T06C12.3 | 0.45214763 | 4 | 1.43E-02 |
| WBGene00006147 | str-88 | R08H2.2 | 0.16564494 | 11 | 3.93E-02 |
| WBGene00006220 | str-176 | T01B11.5 | 0.46190679 | 10 | 6.28E-04 |
| WBGene00006236 | str-200 | F20E11.4 | 0.29568564 | 4 | 4.22E-02 |
| WBGene00006310 | sul-3 | C54D2.4 | 0.17703684 | 13 | 2.43E-02 |
| WBGene00006331 | sup-26 | R10E4.2 | 0.23096793 | 11 | 1.41E-02 |
| WBGene00006342 | sup-37 | C01B7.1 | 0.21010593 | 25 | 1.30E-03 |
| WBGene00006352 | sur-6 | F26E4.1 | 0.20111684 | 16 | 9.05E-03 |
| WBGene00006353 | sur-7 | F01G12.2 | 0.8238189 | 8 | 2.34E-05 |
| WBGene00006366 | sym-1 | C44H4.3 | 0.48450872 | 9 | 7.77E-04 |
| WBGene00006369 | sym-4 | R03E1.1 | 0.39596857 | 16 | 1.25E-04 |
| WBGene00006373 | syx-5 | F55A11.2 | 0.20556231 | 11 | 2.10E-02 |
| WBGene00006380 | tab-1 | F31E8.3 | 0.44279741 | 9 | 1.35E-03 |
| WBGene00006407 | lim-9 | F25H5.1 | 0.18071119 | 88 | 1.77E-07 |
| WBGene00006418 | spl-2 | B0222.4 | 0.34691139 | 10 | 3.30E-03 |
| WBGene00006431 | tag-52 | C02F12.4 | 0.39244826 | 11 | 1.12E-03 |
| WBGene00006439 | ant-1.1 | T27E9.1 | 1.12803664 | 7 | 2.40E-06 |
| WBGene00006444 | shn-1 | C33B4.3 | 0.48292501 | 22 | 7.41E-07 |
| WBGene00006450 | tag-77 | C28C12.10 | 0.20086279 | 26 | 1.47E-03 |
| WBGene00006462 | svh-5 | C33A11.4 | 0.12472411 | 24 | 2.43E-02 |
| WBGene00006472 | tag-123 | F08F1.7 | 0.41054884 | 16 | 9.09E-05 |
| WBGene00006483 | dgk-4 | F42A9.1 | 0.17413265 | 46 | 1.90E-04 |
| WBGene00006493 | ivns-1 | R09A8.3 | 0.19917528 | 26 | 1.56E-03 |
| WBGene00006504 | kcc-1 | R13A1.2 | 0.22103836 | 29 | 3.94E-04 |
| WBGene00006514 | tdp-1 | F44G4.4 | 0.55839066 | 7 | 1.04E-03 |
| WBGene00006517 | madd-3 | E02H4.3 | 0.1092094 | 40 | 1.37E-02 |
| WBGene00006518 | bckd-1B | F27D4.5 | 0.25062352 | 11 | 1.04E-02 |
| WBGene00006523 | tam-1 | F26G5.9 | 0.26277008 | 27 | 1.34E-04 |
| WBGene00006528 | tba-1 | F26E4.8 | 0.20084248 | 13 | 1.57E-02 |
| WBGene00006529 | tba-2 | C47B2.3 | 0.27354143 | 12 | 5.51E-03 |
| WBGene00006533 | tba-7 | T28D6.2 | 0.4366057 | 10 | 9.05E-04 |

| | | | | | |
|---|---|---|---|---|---|
| WBGene00006536 | tbb-1 | K01G5.7 | 0.53410584 | 12 | 6.66E-05 |
| WBGene00006537 | tbb-2 | C36E8.5 | 0.54794938 | 11 | 9.78E-05 |
| WBGene00006543 | tbx-2 | F21H11.3 | 0.25630031 | 14 | 4.46E-03 |
| WBGene00006544 | tbx-7 | ZK328.8 | 0.2458263 | 9 | 1.80E-02 |
| WBGene00006546 | tbx-9 | T07C4.6 | 0.2165319 | 8 | 3.24E-02 |
| WBGene00006565 | tfg-1 | Y63D3A.5 | 0.43585424 | 12 | 3.52E-04 |
| WBGene00006570 | tig-2 | F39G3.8 | 0.29457907 | 12 | 3.86E-03 |
| WBGene00006580 | tlp-1 | T23G4.1 | 0.37826306 | 4 | 2.39E-02 |
| WBGene00006588 | tnt-3 | C14F5.3 | 0.18136903 | 30 | 1.54E-03 |
| WBGene00006591 | toh-1 | T24A11.3 | 0.35402347 | 10 | 2.98E-03 |
| WBGene00006592 | dpy-31 | R151.5 | 0.23633818 | 14 | 6.57E-03 |
| WBGene00006593 | tol-1 | C07F11.1 | 0.0910145 | 42 | 3.23E-02 |
| WBGene00006594 | tom-1 | M01A10.2 | 0.08995689 | 49 | 2.53E-02 |
| WBGene00006595 | top-1 | M01E5.5 | 0.40602245 | 19 | 2.68E-05 |
| WBGene00006599 | tpa-1 | B0545.1 | 0.27886973 | 32 | 1.85E-05 |
| WBGene00006601 | tpi-1 | Y17G7B.7 | 0.14934407 | 13 | 4.01E-02 |
| WBGene00006603 | tps-2 | F19H8.1 | 0.14825352 | 36 | 2.84E-03 |
| WBGene00006605 | tra-2 | C15F1.3 | 0.12766851 | 37 | 6.84E-03 |
| WBGene00006611 | tre-5 | C23H3.7 | 0.26769727 | 13 | 4.66E-03 |
| WBGene00006626 | tsn-1 | F10G7.2 | 0.55327334 | 13 | 2.58E-05 |
| WBGene00006638 | tsp-12 | T14G10.6 | 0.2414908 | 7 | 3.04E-02 |
| WBGene00006646 | tsp-20 | B0198.1 | 0.22950324 | 11 | 1.44E-02 |
| WBGene00006649 | tth-1 | F08F1.8 | 0.76493873 | 4 | 1.66E-03 |
| WBGene00006656 | twk-1 | F21C3.1 | 0.23582684 | 18 | 2.69E-03 |
| WBGene00006660 | twk-5 | B0334.2 | 0.60471024 | 5 | 2.52E-03 |
| WBGene00006665 | twk-10 | K04A8.4 | 0.21089266 | 25 | 1.27E-03 |
| WBGene00006668 | twk-13 | R04F11.4 | 0.11969164 | 37 | 1.01E-02 |
| WBGene00006674 | twk-21 | T01B4.1 | 0.68202308 | 17 | 1.06E-07 |
| WBGene00006678 | twk-25 | M04B2.5 | 0.63051234 | 15 | 1.49E-06 |
| WBGene00006686 | twk-34 | K06B4.12 | 0.24046505 | 11 | 1.22E-02 |
| WBGene00006699 | uba-1 | C47E12.5 | 0.23011829 | 20 | 1.99E-03 |
| WBGene00006702 | ubc-3 | Y71G12B.15 | 0.48193933 | 10 | 4.71E-04 |
| WBGene00006704 | ubc-7 | F58A4.10 | 0.22381281 | 8 | 2.97E-02 |
| WBGene00006706 | ubc-9 | F29B9.6 | 0.26935644 | 5 | 3.86E-02 |
| WBGene00006718 | ubc-23 | C28G1.1 | 0.52658581 | 10 | 2.47E-04 |
| WBGene00006721 | ubh-1 | F46E10.8 | 0.30359138 | 6 | 2.14E-02 |
| WBGene00006722 | ubh-2 | Y40G12A.2 | 0.23906294 | 5 | 4.95E-02 |
| WBGene00006725 | ubl-1 | H06I04.4 | 0.80694806 | 3 | 3.20E-03 |
| WBGene00006727 | ubq-1 | F25B5.4 | 0.37682367 | 33 | 2.01E-07 |
| WBGene00006731 | uev-2 | F56D2.4 | 0.63757046 | 6 | 9.27E-04 |
| WBGene00006732 | uev-3 | F26H9.7 | 0.22286679 | 6 | 4.58E-02 |
| WBGene00006736 | ulp-1 | T10F2.3 | 0.21054364 | 22 | 2.29E-03 |
| WBGene00006739 | ulp-4 | C41C4.6 | 0.57701546 | 16 | 2.35E-06 |
| WBGene00006761 | unc-24 | F57H12.2 | 0.25657403 | 11 | 9.45E-03 |
| WBGene00006769 | unc-33 | Y37E11C.1 | 0.52956296 | 22 | 1.87E-07 |
| WBGene00006771 | tln-1 | Y71G12B.11 | 0.36663344 | 76 | 1.59E-14 |
| WBGene00006773 | unc-37 | W02D3.9 | 0.20188962 | 16 | 8.90E-03 |
| WBGene00006777 | unc-41 | C27H6.1 | 0.1490853 | 46 | 8.46E-04 |

| | | | | | |
|---|---|---|---|---|---|
| WBGene00006779 | unc-43 | K11E8.1 | 0.10958732 | 69 | 1.89E-03 |
| WBGene00006781 | unc-45 | F30H5.1 | 0.11979256 | 24 | 2.84E-02 |
| WBGene00006786 | unc-51 | Y60A3A.1 | 0.46797705 | 29 | 3.07E-08 |
| WBGene00006787 | unc-52 | ZC101.2 | 0.35107854 | 138 | 7.46E-24 |
| WBGene00006792 | unc-58 | T06H11.1 | 0.09495904 | 58 | 1.19E-02 |
| WBGene00006794 | unc-60 | C38C3.5 | 0.27727001 | 15 | 2.24E-03 |
| WBGene00006803 | unc-70 | K11C4.3 | 0.36621244 | 55 | 5.98E-11 |
| WBGene00006815 | unc-83 | W01A11.3 | 0.27172686 | 67 | 1.89E-09 |
| WBGene00006831 | unc-104 | C52E12.2 | 0.1621094 | 64 | 3.52E-05 |
| WBGene00006833 | unc-108 | F53F10.4 | 0.2987514 | 14 | 1.95E-03 |
| WBGene00006839 | unc-115 | F09B9.2 | 0.19271985 | 15 | 1.29E-02 |
| WBGene00006840 | unc-116 | R05D3.7 | 0.15127583 | 14 | 3.44E-02 |
| WBGene00006853 | unc-130 | C47G2.2 | 0.4193601 | 8 | 2.89E-03 |
| WBGene00006861 | cal-5 | C24H10.5 | 0.43323672 | 8 | 2.45E-03 |
| WBGene00006862 | pnk-4 | C42D8.3 | 0.17285081 | 14 | 2.26E-02 |
| WBGene00006876 | vab-10 | ZK1151.1 | 0.26811245 | 247 | 3.67E-30 |
| WBGene00006882 | vab-19 | T22D2.1 | 0.35658861 | 30 | 1.50E-06 |
| WBGene00006887 | vav-1 | C35B8.2 | 0.11634003 | 40 | 9.42E-03 |
| WBGene00006890 | vem-1 | K07E3.8 | 0.48513087 | 4 | 1.14E-02 |
| WBGene00006910 | vha-1 | R10E11.8 | 0.8991375 | 5 | 2.29E-04 |
| WBGene00006911 | vha-2 | R10E11.2 | 0.42902038 | 3 | 2.69E-02 |
| WBGene00006914 | vha-5 | F35H10.4 | 0.28985731 | 15 | 1.73E-03 |
| WBGene00006916 | vha-7 | C26H9A.1 | 0.17341953 | 27 | 3.28E-03 |
| WBGene00006917 | vha-8 | C17H12.14 | 0.56685955 | 4 | 6.51E-03 |
| WBGene00006919 | vha-10 | F46F11.5 | 0.66164326 | 4 | 3.39E-03 |
| WBGene00006920 | vha-11 | Y38F2AL.3 | 0.41225848 | 15 | 1.37E-04 |
| WBGene00006921 | vha-12 | F20B6.2 | 0.48754928 | 10 | 4.34E-04 |
| WBGene00006923 | vhp-1 | F08B1.1 | 0.20494784 | 51 | 1.18E-05 |
| WBGene00006924 | vig-1 | F56D12.5 | 0.51074496 | 11 | 1.75E-04 |
| WBGene00006934 | vps-54 | T21C9.2 | 0.16349972 | 43 | 5.37E-04 |
| WBGene00006941 | wnk-1 | C46C2.1 | 0.59011942 | 48 | 8.24E-16 |
| WBGene00006946 | prx-10 | C34E10.4 | 0.17301616 | 13 | 2.61E-02 |
| WBGene00006950 | wrt-4 | ZK678.5 | 0.40053211 | 16 | 1.13E-04 |
| WBGene00006954 | wrt-8 | C29F3.2 | 0.40405099 | 10 | 1.45E-03 |
| WBGene00006955 | wrt-9 | B0344.2 | 0.13575684 | 41 | 3.10E-03 |
| WBGene00006956 | wrt-10 | ZK1290.8 | 0.72750121 | 9 | 3.17E-05 |
| WBGene00006958 | wve-1 | R06C1.3 | 0.32254262 | 14 | 1.23E-03 |
| WBGene00006959 | xbp-1 | R74.3 | 0.72832679 | 10 | 1.35E-05 |
| WBGene00006964 | xrn-2 | Y48B6A.3 | 0.13619509 | 38 | 4.09E-03 |
| WBGene00006976 | zhp-3 | K02B12.8 | 0.90856313 | 6 | 7.26E-05 |
| WBGene00006987 | zmp-1 | EGAP1.3 | 0.29071607 | 21 | 2.89E-04 |
| WBGene00006996 | zyg-11 | C08B11.1 | 0.18132125 | 10 | 3.60E-02 |
| WBGene00007012 | mdt-4 | ZK546.13 | 0.60801434 | 10 | 7.63E-05 |
| WBGene00007013 | mdt-8 | Y62F5A.1 | 0.0838907 | 64 | 2.20E-02 |
| WBGene00007016 | mdt-15 | R12B2.5 | 0.41957057 | 26 | 7.72E-07 |
| WBGene00007030 | epc-1 | Y111B2A.11 | 0.25588836 | 21 | 7.74E-04 |
| WBGene00007047 | wts-1 | T20F10.1 | 0.25914093 | 18 | 1.52E-03 |
| WBGene00007053 | chd-7 | T04D1.4 | 0.17750482 | 54 | 4.57E-05 |

| | | | | | |
|---|---|---|---|---|---|
| WBGene00007058 | dmd-6 | F13G11.1 | 0.4178388 | 34 | 2.16E-08 |
| WBGene00007064 | rga-9 | 2RSSE.1 | 0.1493907 | 25 | 9.79E-03 |
| WBGene00007071 | AC3.5 | AC3.5 | 0.71329849 | 21 | 1.89E-09 |
| WBGene00007105 | znf-207 | B0035.1 | 0.63558346 | 8 | 2.20E-04 |
| WBGene00007115 | B0198.2 | B0198.2 | 0.21250137 | 14 | 1.04E-02 |
| WBGene00007137 | lcmt-1 | B0285.4 | 0.27328255 | 8 | 1.65E-02 |
| WBGene00007140 | cyp-29A4 | B0331.1 | 0.55970037 | 10 | 1.53E-04 |
| WBGene00007142 | ttr-18 | B0334.1 | 0.6663503 | 6 | 7.07E-04 |
| WBGene00007143 | B0334.3 | B0334.3 | 0.38162262 | 16 | 1.72E-04 |
| WBGene00007150 | acly-2 | B0365.1 | 0.17929928 | 18 | 1.07E-02 |
| WBGene00007178 | B0457.2 | B0457.2 | 0.43058709 | 11 | 6.16E-04 |
| WBGene00007180 | B0457.6 | B0457.6 | 0.16043382 | 10 | 4.87E-02 |
| WBGene00007189 | pigm-1 | B0491.1 | 0.40518152 | 9 | 2.21E-03 |
| WBGene00007202 | B0564.2 | B0564.2 | 0.38065175 | 18 | 7.75E-05 |
| WBGene00007203 | best-1 | B0564.3 | 0.55295082 | 4 | 7.17E-03 |
| WBGene00007207 | fask-1 | B0564.7 | 0.15823557 | 15 | 2.64E-02 |
| WBGene00007214 | C01A2.2 | C01A2.2 | 0.19597819 | 9 | 3.47E-02 |
| WBGene00007227 | C01G6.5 | C01G6.5 | 0.38081903 | 19 | 5.13E-05 |
| WBGene00007256 | swsn-9 | C01H6.7 | 0.34275209 | 12 | 1.70E-03 |
| WBGene00007262 | jud-4 | C02D4.1 | 0.40677641 | 7 | 5.22E-03 |
| WBGene00007264 | C02F4.4 | C02F4.4 | 0.57370485 | 5 | 3.24E-03 |
| WBGene00007270 | rei-1 | C03C10.4 | 0.68208605 | 6 | 6.10E-04 |
| WBGene00007275 | C03D6.1 | C03D6.1 | 0.54734871 | 10 | 1.83E-04 |
| WBGene00007286 | C04A11.2 | C04A11.2 | 0.43271045 | 10 | 9.57E-04 |
| WBGene00007287 | gck-4 | C04A11.3 | 0.16462203 | 17 | 1.76E-02 |
| WBGene00007288 | C04A11.5 | C04A11.5 | 0.69844458 | 3 | 5.90E-03 |
| WBGene00007299 | C04F12.5 | C04F12.5 | 1.20823903 | 2 | 1.42E-03 |
| WBGene00007317 | C05A9.2 | C05A9.2 | 0.25421682 | 9 | 1.61E-02 |
| WBGene00007325 | C05C9.1 | C05C9.1 | 0.97773837 | 11 | 1.16E-07 |
| WBGene00007343 | C05E7.1 | C05E7.1 | 1.31164309 | 4 | 3.85E-05 |
| WBGene00007344 | C05E7.2 | C05E7.2 | 0.39141738 | 3 | 3.32E-02 |
| WBGene00007352 | cdc-48.1 | C06A1.1 | 0.47288244 | 21 | 1.68E-06 |
| WBGene00007363 | stdh-1 | C06B3.4 | 0.47069883 | 5 | 7.50E-03 |
| WBGene00007365 | C06B3.6 | C06B3.6 | 0.38134476 | 4 | 2.34E-02 |
| WBGene00007366 | C06B3.7 | C06B3.7 | 0.18920669 | 8 | 4.49E-02 |
| WBGene00007372 | C06B8.7 | C06B8.7 | 0.24033614 | 73 | 7.00E-09 |
| WBGene00007384 | swt-3 | C06G8.1 | 0.37407479 | 10 | 2.23E-03 |
| WBGene00007391 | C06H2.7 | C06H2.7 | 0.16341303 | 10 | 4.66E-02 |
| WBGene00007395 | dcar-1 | C06H5.7 | 0.40688732 | 18 | 4.08E-05 |
| WBGene00007396 | rnp-9 | C07A4.1 | 0.25185864 | 9 | 1.66E-02 |
| WBGene00007397 | C07A4.2 | C07A4.2 | 0.75880307 | 7 | 1.23E-04 |
| WBGene00007398 | C07A4.3 | C07A4.3 | 0.29218413 | 5 | 3.21E-02 |
| WBGene00007401 | C07A9.5 | C07A9.5 | 0.25399945 | 12 | 7.67E-03 |
| WBGene00007403 | set-3 | C07A9.7 | 0.17647209 | 14 | 2.11E-02 |
| WBGene00007405 | C07A9.9 | C07A9.9 | 0.23205358 | 11 | 1.39E-02 |
| WBGene00007406 | C07A9.10 | C07A9.10 | 0.16052163 | 10 | 4.86E-02 |
| WBGene00007409 | C07B5.4 | C07B5.4 | 0.10773609 | 24 | 4.18E-02 |
| WBGene00007436 | C08B11.9 | C08B11.9 | 0.27984899 | 6 | 2.68E-02 |

| | | | | | |
|---|---|---|---|---|---|
| WBGene00007444 | C08F8.2 | C08F8.2 | 0.13747567 | 22 | 1.98E-02 |
| WBGene00007449 | C08F8.9 | C08F8.9 | 0.37845985 | 4 | 2.38E-02 |
| WBGene00007463 | vgln-1 | C08H9.2 | 0.27386708 | 34 | 1.32E-05 |
| WBGene00007465 | chil-1 | C08H9.4 | 0.17523785 | 18 | 1.18E-02 |
| WBGene00007474 | C08H9.15 | C08H9.15 | 0.36306065 | 4 | 2.65E-02 |
| WBGene00007479 | C09F9.2 | C09F9.2 | 0.23597763 | 35 | 5.71E-05 |
| WBGene00007488 | dph-2 | C09G5.2 | 0.44293654 | 40 | 3.80E-10 |
| WBGene00007492 | C09G9.1 | C09G9.1 | 0.27233097 | 15 | 2.49E-03 |
| WBGene00007514 | catp-8 | C10C6.6 | 0.37725809 | 19 | 5.62E-05 |
| WBGene00007529 | C11H1.3 | C11H1.3 | 0.24195527 | 12 | 9.40E-03 |
| WBGene00007533 | cbl-1 | C12C8.2 | 0.44379513 | 7 | 3.52E-03 |
| WBGene00007545 | C13B4.1 | C13B4.1 | 0.23795412 | 24 | 6.46E-04 |
| WBGene00007546 | nhr-153 | C13C4.1 | 0.38295818 | 9 | 2.96E-03 |
| WBGene00007548 | C13C4.4 | C13C4.4 | 0.20975933 | 7 | 4.27E-02 |
| WBGene00007557 | C14A4.6 | C14A4.6 | 0.31892164 | 8 | 9.58E-03 |
| WBGene00007560 | C14A4.9 | C14A4.9 | 0.25690936 | 5 | 4.28E-02 |
| WBGene00007562 | C14A4.12 | C14A4.12 | 0.26309177 | 16 | 2.32E-03 |
| WBGene00007586 | ril-2 | C14C10.3 | 0.78673966 | 15 | 5.87E-08 |
| WBGene00007587 | mma-1 | C14C10.4 | 0.13471182 | 23 | 1.95E-02 |
| WBGene00007592 | C14H10.2 | C14H10.2 | 0.40232365 | 31 | 1.59E-07 |
| WBGene00007601 | C15C6.2 | C15C6.2 | 0.49789488 | 8 | 1.14E-03 |
| WBGene00007630 | har-1 | C16C10.11 | 0.54874403 | 3 | 1.37E-02 |
| WBGene00007633 | C16D2.1 | C16D2.1 | 0.44757465 | 6 | 5.53E-03 |
| WBGene00007639 | C17D12.5 | C17D12.5 | 0.43241818 | 3 | 2.64E-02 |
| WBGene00007651 | nra-3 | C17G1.4 | 0.33136317 | 20 | 1.30E-04 |
| WBGene00007675 | C18D4.6 | C18D4.6 | 0.31443232 | 13 | 1.99E-03 |
| WBGene00007679 | C18D11.1 | C18D11.1 | 0.69487809 | 9 | 4.86E-05 |
| WBGene00007680 | maa-1 | C18D11.2 | 0.2249683 | 8 | 2.93E-02 |
| WBGene00007711 | lid-1 | C25A1.12 | 0.25232157 | 10 | 1.29E-02 |
| WBGene00007723 | C25F9.2 | C25F9.2 | 0.1996702 | 15 | 1.12E-02 |
| WBGene00007736 | igdb-2 | C25G4.10 | 0.15663015 | 38 | 1.48E-03 |
| WBGene00007753 | C27A7.1 | C27A7.1 | 0.15662171 | 18 | 1.87E-02 |
| WBGene00007756 | C27A7.5 | C27A7.5 | 0.44107489 | 24 | 9.68E-07 |
| WBGene00007759 | C27A7.9 | C27A7.9 | 0.53996653 | 2 | 2.65E-02 |
| WBGene00007764 | C27B7.7 | C27B7.7 | 0.1273354 | 27 | 1.71E-02 |
| WBGene00007778 | C27D8.2 | C27D8.2 | 0.20435055 | 16 | 8.43E-03 |
| WBGene00007779 | C27D8.3 | C27D8.3 | 0.30626415 | 17 | 6.56E-04 |
| WBGene00007787 | C27H6.8 | C27H6.8 | 0.26256707 | 12 | 6.63E-03 |
| WBGene00007791 | C28A5.6 | C28A5.6 | 0.29193331 | 22 | 2.08E-04 |
| WBGene00007804 | C29F3.3 | C29F3.3 | 0.68759101 | 6 | 5.79E-04 |
| WBGene00007810 | C29F7.1 | C29F7.1 | 0.26275136 | 21 | 6.37E-04 |
| WBGene00007836 | C31C9.2 | C31C9.2 | 0.66180464 | 11 | 1.64E-05 |
| WBGene00007841 | C31C9.7 | C31C9.7 | 0.7265996 | 5 | 9.33E-04 |
| WBGene00007848 | cytb-5.1 | C31E10.7 | 0.31474913 | 6 | 1.93E-02 |
| WBGene00007849 | tbc-19 | C31E10.8 | 0.18868626 | 25 | 2.65E-03 |
| WBGene00007856 | C31H5.5 | C31H5.5 | 0.51344685 | 7 | 1.68E-03 |
| WBGene00007861 | C32A9.1 | C32A9.1 | 0.25618321 | 6 | 3.35E-02 |
| WBGene00007877 | nfki-1 | C33A11.1 | 0.09637162 | 32 | 4.01E-02 |

| | | | | | |
|---|---|---|---|---|---|
| WBGene00007878 | C33A11.2 | C33A11.2 | 0.22030539 | 17 | 4.83E-03 |
| WBGene00007882 | C33A12.3 | C33A12.3 | 0.33908363 | 8 | 7.53E-03 |
| WBGene00007885 | ugt-21 | C33A12.6 | 0.19649941 | 13 | 1.70E-02 |
| WBGene00007886 | ethe-1 | C33A12.7 | 0.30477939 | 6 | 2.12E-02 |
| WBGene00007890 | C33A12.19 | C33A12.19 | 0.33287075 | 4 | 3.26E-02 |
| WBGene00007893 | C33B4.5 | C33B4.5 | 0.37332879 | 3 | 3.68E-02 |
| WBGene00007894 | C33D3.3 | C33D3.3 | 0.46299539 | 5 | 7.98E-03 |
| WBGene00007901 | C33D9.8 | C33D9.8 | 0.29775838 | 7 | 1.67E-02 |
| WBGene00007903 | lgc-21 | C33G3.3 | 0.26761462 | 13 | 4.67E-03 |
| WBGene00007913 | cyp-36A1 | C34B7.3 | 0.19848051 | 17 | 8.02E-03 |
| WBGene00007927 | C34C12.8 | C34C12.8 | 0.25346681 | 6 | 3.43E-02 |
| WBGene00007934 | C34E7.4 | C34E7.4 | 0.59706481 | 14 | 5.89E-06 |
| WBGene00007947 | C35A5.3 | C35A5.3 | 0.37352209 | 5 | 1.65E-02 |
| WBGene00007975 | gls-1 | C36B1.8 | 0.17988106 | 15 | 1.69E-02 |
| WBGene00007977 | gska-3 | C36B1.10 | 0.33051196 | 7 | 1.18E-02 |
| WBGene00007982 | pxd-1 | C36E8.3 | 0.20406377 | 11 | 2.15E-02 |
| WBGene00007983 | C36E8.4 | C36E8.4 | 0.22233624 | 12 | 1.31E-02 |
| WBGene00007988 | best-8 | C37A5.1 | 0.39060732 | 12 | 7.58E-04 |
| WBGene00007989 | fipr-22 | C37A5.2 | 0.48826243 | 6 | 3.77E-03 |
| WBGene00007991 | C37A5.7 | C37A5.7 | 0.16686893 | 12 | 3.36E-02 |
| WBGene00007994 | C37E2.2 | C37E2.2 | 0.18591785 | 8 | 4.67E-02 |
| WBGene00007996 | C38C6.3 | C38C6.3 | 0.92229559 | 8 | 7.23E-06 |
| WBGene00007997 | sre-13 | C38C6.4 | 0.47323536 | 5 | 7.34E-03 |
| WBGene00007999 | tag-297 | C38C6.6 | 0.70359987 | 12 | 3.77E-06 |
| WBGene00008032 | C39E9.8 | C39E9.8 | 0.30357646 | 15 | 1.30E-03 |
| WBGene00008050 | chst-1 | C41C4.1 | 0.53974726 | 4 | 7.85E-03 |
| WBGene00008052 | ctns-1 | C41C4.7 | 0.24036407 | 6 | 3.88E-02 |
| WBGene00008054 | C41C4.9 | C41C4.9 | 0.36682387 | 5 | 1.75E-02 |
| WBGene00008084 | C44C10.3 | C44C10.3 | 0.23272929 | 8 | 2.67E-02 |
| WBGene00008089 | C44C10.9 | C44C10.9 | 0.26920878 | 11 | 7.75E-03 |
| WBGene00008094 | C44H4.4 | C44H4.4 | 0.12964324 | 51 | 1.70E-03 |
| WBGene00008100 | C44H9.5 | C44H9.5 | 0.15894367 | 12 | 3.84E-02 |
| WBGene00008111 | C46C2.3 | C46C2.3 | 0.21186302 | 10 | 2.32E-02 |
| WBGene00008113 | C46C2.5 | C46C2.5 | 0.67358189 | 3 | 6.78E-03 |
| WBGene00008117 | gsr-1 | C46F11.2 | 0.41566808 | 12 | 4.96E-04 |
| WBGene00008132 | gale-1 | C47B2.6 | 0.16165225 | 25 | 6.51E-03 |
| WBGene00008144 | gasr-8 | C47E8.6 | 0.34887554 | 10 | 3.21E-03 |
| WBGene00008151 | rrp-1 | C47E12.7 | 0.41402072 | 8 | 3.08E-03 |
| WBGene00008156 | C47E12.13 | C47E12.13 | 0.22219767 | 6 | 4.60E-02 |
| WBGene00008165 | C47G2.4 | C47G2.4 | 0.23893987 | 11 | 1.25E-02 |
| WBGene00008183 | rin-1 | C48G7.3 | 0.09188634 | 87 | 4.02E-03 |
| WBGene00008195 | ceh-88 | C49C3.5 | 0.15926038 | 12 | 3.82E-02 |
| WBGene00008196 | C49C3.6 | C49C3.6 | 0.73060257 | 3 | 4.92E-03 |
| WBGene00008197 | C49C3.7 | C49C3.7 | 0.50449941 | 9 | 5.97E-04 |
| WBGene00008205 | sams-1 | C49F5.1 | 0.63565991 | 12 | 1.19E-05 |
| WBGene00008210 | C49F5.6 | C49F5.6 | 0.20920851 | 10 | 2.41E-02 |
| WBGene00008212 | C49F5.8 | C49F5.8 | 0.26988319 | 7 | 2.25E-02 |
| WBGene00008215 | C49F8.3 | C49F8.3 | 0.36077981 | 6 | 1.25E-02 |

| | | | | | |
|---|---|---|---|---|---|
| WBGene00008218 | nasp-2 | C50B6.2 | 0.28504808 | 5 | 3.40E-02 |
| WBGene00008222 | C50B6.9 | C50B6.9 | 0.19938296 | 7 | 4.76E-02 |
| WBGene00008224 | slrp-1 | C50B8.1 | 0.4336001 | 5 | 1.01E-02 |
| WBGene00008233 | C50F4.8 | C50F4.8 | 0.33478569 | 8 | 7.93E-03 |
| WBGene00008260 | C52G5.2 | C52G5.2 | 0.87156383 | 7 | 3.69E-05 |
| WBGene00008270 | C53A5.13 | C53A5.13 | 0.37142146 | 19 | 6.54E-05 |
| WBGene00008277 | mltn-12 | C53B4.8 | 0.14596549 | 21 | 1.73E-02 |
| WBGene00008279 | C53C7.3 | C53C7.3 | 0.5162729 | 11 | 1.61E-04 |
| WBGene00008320 | slc-25A29 | C54G10.4 | 0.4812637 | 15 | 3.28E-05 |
| WBGene00008331 | ttll-5 | C55A6.2 | 0.39000512 | 16 | 1.43E-04 |
| WBGene00008332 | C55A6.3 | C55A6.3 | 0.67932037 | 5 | 1.37E-03 |
| WBGene00008333 | C55A6.4 | C55A6.4 | 0.55499841 | 7 | 1.08E-03 |
| WBGene00008336 | C55A6.7 | C55A6.7 | 0.50576029 | 10 | 3.34E-04 |
| WBGene00008362 | cfim-2 | D1046.1 | 0.62438842 | 13 | 7.07E-06 |
| WBGene00008395 | D1086.8 | D1086.8 | 0.24382513 | 6 | 3.76E-02 |
| WBGene00008404 | wbp-2 | D2013.6 | 0.32520342 | 4 | 3.44E-02 |
| WBGene00008412 | D2030.2 | D2030.2 | 0.13854328 | 14 | 4.40E-02 |
| WBGene00008414 | D2030.4 | D2030.4 | 0.54454555 | 3 | 1.40E-02 |
| WBGene00008419 | wdr-23 | D2030.9 | 0.12958119 | 26 | 1.73E-02 |
| WBGene00008433 | marc-2 | D2089.2 | 0.77463606 | 5 | 6.31E-04 |
| WBGene00008443 | pde-3 | E01F3.1 | 0.10754797 | 46 | 1.01E-02 |
| WBGene00008496 | F01D5.6 | F01D5.6 | 0.43682197 | 10 | 9.02E-04 |
| WBGene00008505 | F01G4.6 | F01G4.6 | 0.91244386 | 7 | 2.38E-05 |
| WBGene00008506 | tkt-1 | F01G10.1 | 0.1584483 | 15 | 2.63E-02 |
| WBGene00008511 | F01G10.9 | F01G10.9 | 0.37628384 | 18 | 8.63E-05 |
| WBGene00008512 | F01G10.10 | F01G10.10 | 0.15843345 | 14 | 2.99E-02 |
| WBGene00008514 | mrpl-44 | F02A9.4 | 0.285035 | 18 | 8.05E-04 |
| WBGene00008516 | F02C12.2 | F02C12.2 | 0.52061143 | 4 | 8.95E-03 |
| WBGene00008521 | F02D8.4 | F02D8.4 | 0.36306676 | 12 | 1.21E-03 |
| WBGene00008526 | F02D10.6 | F02D10.6 | 0.23403067 | 7 | 3.29E-02 |
| WBGene00008549 | din-1 | F07A11.6 | 0.21120864 | 73 | 1.07E-07 |
| WBGene00008555 | F07C6.4 | F07C6.4 | 0.10666779 | 65 | 3.16E-03 |
| WBGene00008570 | kcnl-2 | F08A10.1 | 0.10053984 | 82 | 2.03E-03 |
| WBGene00008579 | F08G2.8 | F08G2.8 | 0.24521539 | 10 | 1.43E-02 |
| WBGene00008601 | mob-2 | F09A5.4 | 0.12409597 | 36 | 8.85E-03 |
| WBGene00008605 | mlt-9 | F09B12.1 | 0.13264878 | 14 | 4.94E-02 |
| WBGene00008621 | F09C8.1 | F09C8.1 | 0.17539355 | 14 | 2.15E-02 |
| WBGene00008622 | F09C8.2 | F09C8.2 | 0.19612349 | 20 | 4.99E-03 |
| WBGene00008623 | spe-43 | F09E8.1 | 0.28618146 | 7 | 1.89E-02 |
| WBGene00008644 | F10C2.3 | F10C2.3 | 0.13354363 | 19 | 2.98E-02 |
| WBGene00008652 | F10D11.6 | F10D11.6 | 0.41433886 | 15 | 1.31E-04 |
| WBGene00008664 | nubp-1 | F10G8.6 | 0.29736252 | 5 | 3.08E-02 |
| WBGene00008669 | acs-14 | F11A3.1 | 0.41090264 | 12 | 5.37E-04 |
| WBGene00008679 | F11A5.13 | F11A5.13 | 0.67234526 | 5 | 1.45E-03 |
| WBGene00008683 | repo-1 | F11A10.2 | 0.26710187 | 8 | 1.78E-02 |
| WBGene00008687 | F11A10.6 | F11A10.6 | 0.23952594 | 8 | 2.47E-02 |
| WBGene00008688 | rbm-34 | F11A10.7 | 0.77231506 | 5 | 6.43E-04 |
| WBGene00008706 | gba-3 | F11E6.1 | 0.23702057 | 16 | 4.11E-03 |

| | | | | | |
|---|---|---|---|---|---|
| WBGene00008707 | F11E6.3 | F11E6.3 | 0.59985 | 6 | 1.32E-03 |
| WBGene00008712 | F11E6.9 | F11E6.9 | 0.90918636 | 5 | 2.11E-04 |
| WBGene00008714 | F11F1.1 | F11F1.1 | 0.16350988 | 13 | 3.10E-02 |
| WBGene00008735 | F13D2.1 | F13D2.1 | 0.11670572 | 73 | 7.42E-04 |
| WBGene00008743 | F13D12.9 | F13D12.9 | 0.32197386 | 18 | 3.26E-04 |
| WBGene00008746 | F13E6.2 | F13E6.2 | 0.13387421 | 15 | 4.37E-02 |
| WBGene00008752 | fipr-15 | F13E9.3 | 0.69403008 | 2 | 1.35E-02 |
| WBGene00008765 | ttx-7 | F13G3.5 | 0.23413687 | 8 | 2.63E-02 |
| WBGene00008769 | mrpl-13 | F13G3.11 | 0.36703878 | 5 | 1.74E-02 |
| WBGene00008779 | F14B4.1 | F14B4.1 | 0.77361845 | 18 | 5.15E-09 |
| WBGene00008780 | hxk-1 | F14B4.2 | 0.31027945 | 31 | 6.81E-06 |
| WBGene00008781 | rpoa-2 | F14B4.3 | 0.12525756 | 18 | 4.02E-02 |
| WBGene00008803 | lips-10 | F14E5.5 | 0.15116634 | 18 | 2.13E-02 |
| WBGene00008848 | inft-2 | F15B9.4 | 0.30646617 | 17 | 6.53E-04 |
| WBGene00008856 | F15D3.6 | F15D3.6 | 0.26422836 | 14 | 3.82E-03 |
| WBGene00008857 | timm-23 | F15D3.7 | 1.14788348 | 9 | 1.25E-07 |
| WBGene00008858 | pals-1 | F15D3.8 | 0.84965163 | 9 | 6.33E-06 |
| WBGene00008876 | F16A11.1 | F16A11.1 | 0.11386445 | 28 | 2.57E-02 |
| WBGene00008877 | rtcb-1 | F16A11.2 | 0.48270399 | 7 | 2.33E-03 |
| WBGene00008878 | ppfr-1 | F16A11.3 | 0.089933 | 82 | 6.19E-03 |
| WBGene00008882 | F16B12.6 | F16B12.6 | 0.19346941 | 26 | 1.90E-03 |
| WBGene00008886 | F16C3.2 | F16C3.2 | 0.1599177 | 10 | 4.90E-02 |
| WBGene00008920 | eef-1G | F17C11.9 | 0.27881572 | 8 | 1.54E-02 |
| WBGene00008926 | F17H10.2 | F17H10.2 | 0.52597105 | 10 | 2.49E-04 |
| WBGene00008927 | snx-17 | F17H10.3 | 0.28734238 | 15 | 1.82E-03 |
| WBGene00008940 | F18H3.4 | F18H3.4 | 0.14994388 | 18 | 2.20E-02 |
| WBGene00008952 | F19C6.3 | F19C6.3 | 0.43844489 | 14 | 1.29E-04 |
| WBGene00008959 | F19H6.4 | F19H6.4 | 0.63786049 | 8 | 2.14E-04 |
| WBGene00008963 | F19H8.2 | F19H8.2 | 0.40258221 | 2 | 4.82E-02 |
| WBGene00008964 | mltn-9 | F19H8.4 | 0.46405274 | 27 | 9.95E-08 |
| WBGene00008972 | F20C5.7 | F20C5.7 | 0.23665775 | 9 | 2.03E-02 |
| WBGene00008975 | F20D1.3 | F20D1.3 | 0.37230181 | 12 | 1.03E-03 |
| WBGene00008976 | plp-2 | F20D1.4 | 0.50271943 | 6 | 3.29E-03 |
| WBGene00008993 | F21A3.3 | F21A3.3 | 0.2931205 | 4 | 4.29E-02 |
| WBGene00008997 | lgc-31 | F21A3.7 | 0.27032472 | 24 | 2.29E-04 |
| WBGene00009002 | hint-1 | F21C3.3 | 0.75064858 | 4 | 1.84E-03 |
| WBGene00009013 | mrps-33 | F21D5.8 | 0.376221 | 3 | 3.62E-02 |
| WBGene00009025 | phf-34 | F21G4.4 | 0.23449871 | 8 | 2.62E-02 |
| WBGene00009028 | F21H7.2 | F21H7.2 | 0.46389651 | 9 | 1.02E-03 |
| WBGene00009050 | F22D6.2 | F22D6.2 | 0.29911625 | 7 | 1.65E-02 |
| WBGene00009054 | F22D6.9 | F22D6.9 | 0.28499842 | 11 | 6.05E-03 |
| WBGene00009057 | cept-1 | F22E10.5 | 0.19562359 | 11 | 2.46E-02 |
| WBGene00009060 | F22E12.3 | F22E12.3 | 0.16264201 | 12 | 3.60E-02 |
| WBGene00009064 | F22G12.4 | F22G12.4 | 0.42367778 | 18 | 2.70E-05 |
| WBGene00009081 | F23B12.4 | F23B12.4 | 0.17627223 | 19 | 9.92E-03 |
| WBGene00009082 | dlat-1 | F23B12.5 | 0.26450016 | 9 | 1.41E-02 |
| WBGene00009087 | F23D12.3 | F23D12.3 | 0.37892777 | 4 | 2.38E-02 |
| WBGene00009093 | F23H12.3 | F23H12.3 | 0.20653186 | 9 | 3.02E-02 |

| | | | | | |
|---|---|---|---|---|---|
| WBGene00009112 | tag-353 | F25D7.2 | 0.24034449 | 7 | 3.08E-02 |
| WBGene00009119 | ndk-1 | F25H2.5 | 0.39178466 | 4 | 2.17E-02 |
| WBGene00009126 | pyk-1 | F25H5.3 | 0.36052335 | 27 | 4.05E-06 |
| WBGene00009133 | bed-3 | F25H8.6 | 0.26646421 | 26 | 1.53E-04 |
| WBGene00009158 | F26E4.3 | F26E4.3 | 0.35823737 | 13 | 8.97E-04 |
| WBGene00009159 | F26E4.4 | F26E4.4 | 0.23501069 | 9 | 2.08E-02 |
| WBGene00009160 | F26E4.5 | F26E4.5 | 0.19804446 | 9 | 3.38E-02 |
| WBGene00009177 | F26H9.5 | F26H9.5 | 0.32478848 | 8 | 8.93E-03 |
| WBGene00009178 | uggt-2 | F26H9.8 | 0.33495498 | 33 | 1.23E-06 |
| WBGene00009218 | acs-20 | F28D1.9 | 0.16454338 | 19 | 1.34E-02 |
| WBGene00009232 | nkat-1 | F28H6.3 | 0.33593108 | 6 | 1.58E-02 |
| WBGene00009233 | F28H6.4 | F28H6.4 | 0.12519804 | 23 | 2.61E-02 |
| WBGene00009239 | irld-6 | F28H7.6 | 0.20578038 | 11 | 2.10E-02 |
| WBGene00009243 | F29C6.1 | F29C6.1 | 0.23732957 | 7 | 3.18E-02 |
| WBGene00009261 | F30A10.2 | F30A10.2 | 0.33780303 | 7 | 1.09E-02 |
| WBGene00009299 | F31E9.6 | F31E9.6 | 0.18555255 | 9 | 3.99E-02 |
| WBGene00009306 | maph-1.1 | F32A7.5 | 0.42733058 | 22 | 3.82E-06 |
| WBGene00009308 | F32A11.3 | F32A11.3 | 0.34917643 | 8 | 6.68E-03 |
| WBGene00009328 | F32D8.3 | F32D8.3 | 0.4145861 | 3 | 2.92E-02 |
| WBGene00009334 | F32D8.12 | F32D8.12 | 0.13726395 | 14 | 4.52E-02 |
| WBGene00009339 | F32G8.3 | F32G8.3 | 0.50324374 | 5 | 5.75E-03 |
| WBGene00009340 | best-14 | F32G8.4 | 0.26062082 | 8 | 1.92E-02 |
| WBGene00009341 | thoc-3 | F32H2.4 | 0.27902517 | 10 | 8.79E-03 |
| WBGene00009342 | fasn-1 | F32H2.5 | 1.01750224 | 60 | 1.19E-33 |
| WBGene00009349 | F32H5.3 | F32H5.3 | 0.73089667 | 4 | 2.10E-03 |
| WBGene00009355 | F33A8.7 | F33A8.7 | 0.21842153 | 7 | 3.89E-02 |
| WBGene00009358 | F33C8.4 | F33C8.4 | 0.6657752 | 3 | 7.09E-03 |
| WBGene00009367 | F33H2.3 | F33H2.3 | 0.52383627 | 5 | 4.86E-03 |
| WBGene00009380 | F34H10.3 | F34H10.3 | 0.57287267 | 4 | 6.25E-03 |
| WBGene00009382 | F34H10.5 | F34H10.5 | 0.62700092 | 4 | 4.30E-03 |
| WBGene00009386 | tag-290 | F35B12.6 | 0.22208292 | 8 | 3.04E-02 |
| WBGene00009405 | F35C11.6 | F35C11.6 | 0.43273294 | 6 | 6.36E-03 |
| WBGene00009439 | mlcd-1 | F35G12.1 | 0.23112078 | 6 | 4.23E-02 |
| WBGene00009447 | F35H8.2 | F35H8.2 | 0.31385543 | 7 | 1.41E-02 |
| WBGene00009450 | ugt-58 | F35H8.6 | 0.42612784 | 5 | 1.08E-02 |
| WBGene00009452 | F36A2.2 | F36A2.2 | 0.1993513 | 12 | 1.94E-02 |
| WBGene00009453 | F36A2.3 | F36A2.3 | 0.27786126 | 19 | 7.26E-04 |
| WBGene00009482 | F36G3.1 | F36G3.1 | 0.10848638 | 27 | 3.35E-02 |
| WBGene00009499 | ent-6 | F36H2.2 | 0.27240975 | 12 | 5.61E-03 |
| WBGene00009504 | F37B12.1 | F37B12.1 | 0.52827531 | 7 | 1.43E-03 |
| WBGene00009528 | F38A6.4 | F38A6.4 | 0.33611119 | 9 | 5.49E-03 |
| WBGene00009532 | ccch-1 | F38B7.1 | 0.17856633 | 43 | 2.31E-04 |
| WBGene00009542 | copb-2 | F38E11.5 | 0.18509434 | 26 | 2.54E-03 |
| WBGene00009552 | piki-1 | F39B1.1 | 0.722907 | 36 | 4.99E-15 |
| WBGene00009584 | drap-1 | F40F9.7 | 0.26969242 | 19 | 8.96E-04 |
| WBGene00009587 | mig-38 | F40F11.2 | 0.29080402 | 41 | 7.87E-07 |
| WBGene00009618 | F41E7.2 | F41E7.2 | 0.21698195 | 19 | 3.48E-03 |
| WBGene00009619 | npr-6 | F41E7.3 | 0.17284883 | 13 | 2.62E-02 |

| | | | | | |
|---|---|---|---|---|---|
| WBGene00009621 | fipr-21 | F41E7.5 | 0.55048485 | 2 | 2.53E-02 |
| WBGene00009622 | F41E7.6 | F41E7.6 | 0.34889967 | 9 | 4.64E-03 |
| WBGene00009623 | F41E7.7 | F41E7.7 | 0.20704199 | 8 | 3.63E-02 |
| WBGene00009625 | F41E7.9 | F41E7.9 | 0.46396518 | 10 | 6.10E-04 |
| WBGene00009626 | F42A8.1 | F42A8.1 | 0.36869533 | 20 | 4.73E-05 |
| WBGene00009628 | tatn-1 | F42D1.2 | 1.4279472 | 10 | 5.56E-10 |
| WBGene00009635 | F42F12.3 | F42F12.3 | 0.56223461 | 8 | 5.27E-04 |
| WBGene00009645 | F42G10.1 | F42G10.1 | 0.28814432 | 17 | 9.99E-04 |
| WBGene00009652 | F43D2.3 | F43D2.3 | 0.48722023 | 7 | 2.22E-03 |
| WBGene00009658 | pap-3 | F43G6.5 | 0.35727436 | 16 | 2.93E-04 |
| WBGene00009661 | patr-1 | F43G6.9 | 0.39438342 | 17 | 8.48E-05 |
| WBGene00009674 | nucb-1 | F44A6.1 | 0.19767513 | 14 | 1.39E-02 |
| WBGene00009678 | magu-4 | F44D12.1 | 0.1090502 | 24 | 4.01E-02 |
| WBGene00009686 | ent-7 | F44D12.9 | 0.17625301 | 11 | 3.33E-02 |
| WBGene00009712 | F44G4.2 | F44G4.2 | 0.61924062 | 4 | 4.54E-03 |
| WBGene00009717 | dep-1 | F44G4.8 | 0.08659193 | 39 | 4.61E-02 |
| WBGene00009722 | F45D3.2 | F45D3.2 | 1.15426148 | 5 | 2.86E-05 |
| WBGene00009723 | F45D3.3 | F45D3.3 | 1.15389527 | 4 | 1.14E-04 |
| WBGene00009724 | F45D3.4 | F45D3.4 | 1.2739064 | 3 | 2.31E-04 |
| WBGene00009726 | tbc-13 | F45E6.3 | 0.11698025 | 20 | 4.23E-02 |
| WBGene00009728 | sdz-19 | F45E6.6 | 0.33789979 | 3 | 4.49E-02 |
| WBGene00009759 | ttr-12 | F46B3.4 | 0.23932182 | 5 | 4.94E-02 |
| WBGene00009772 | ztf-7 | F46B6.7 | 0.18403129 | 17 | 1.12E-02 |
| WBGene00009783 | rer-1 | F46C5.8 | 0.31851481 | 6 | 1.86E-02 |
| WBGene00009787 | F46F2.3 | F46F2.3 | 0.91507127 | 3 | 1.74E-03 |
| WBGene00009789 | F46F2.5 | F46F2.5 | 0.20473938 | 13 | 1.47E-02 |
| WBGene00009811 | F47B8.10 | F47B8.10 | 0.14489179 | 16 | 3.11E-02 |
| WBGene00009812 | suca-1 | F47B10.1 | 0.20067369 | 12 | 1.89E-02 |
| WBGene00009813 | haly-1 | F47B10.2 | 0.54788136 | 14 | 1.53E-05 |
| WBGene00009816 | F47B10.5 | F47B10.5 | 0.21090475 | 7 | 4.21E-02 |
| WBGene00009829 | tmed-10 | F47G9.1 | 0.22912127 | 10 | 1.81E-02 |
| WBGene00009830 | cutl-18 | F47G9.3 | 0.43164495 | 11 | 6.06E-04 |
| WBGene00009850 | F48F7.5 | F48F7.5 | 0.39037702 | 4 | 2.20E-02 |
| WBGene00009867 | F49C5.4 | F49C5.4 | 0.69071998 | 22 | 1.61E-09 |
| WBGene00009868 | F49C5.5 | F49C5.5 | 0.38048958 | 5 | 1.56E-02 |
| WBGene00009878 | F49C12.9 | F49C12.9 | 0.30888568 | 9 | 7.85E-03 |
| WBGene00009888 | F49E2.5 | F49E2.5 | 0.18402485 | 35 | 6.18E-04 |
| WBGene00009918 | gcsh-2 | F52A8.5 | 0.43157502 | 6 | 6.43E-03 |
| WBGene00009925 | F52B11.2 | F52B11.2 | 0.4801234 | 8 | 1.40E-03 |
| WBGene00009926 | noah-2 | F52B11.3 | 0.59256264 | 25 | 3.86E-09 |
| WBGene00009929 | abts-2 | F52D10.1 | 0.12618815 | 16 | 4.69E-02 |
| WBGene00009935 | F52E10.4 | F52E10.4 | 0.15857895 | 13 | 3.39E-02 |
| WBGene00009944 | mtcu-2 | F52H3.2 | 0.22337699 | 8 | 2.99E-02 |
| WBGene00009947 | ttc-36 | F52H3.5 | 1.00776837 | 5 | 9.44E-05 |
| WBGene00009980 | F53F1.2 | F53F1.2 | 0.29097483 | 4 | 4.35E-02 |
| WBGene00009982 | F53F1.4 | F53F1.4 | 1.2484115 | 2 | 1.19E-03 |
| WBGene00009984 | F53F1.6 | F53F1.6 | 0.51881521 | 5 | 5.07E-03 |
| WBGene00010001 | F53F8.4 | F53F8.4 | 1.19370692 | 6 | 4.97E-06 |

| | | | | | |
|---|---|---|---|---|---|
| WBGene00010012 | saeg-1 | F53H10.2 | 0.10820192 | 37 | 1.75E-02 |
| WBGene00010018 | fbxa-69 | F54B8.3 | 0.26054981 | 6 | 3.21E-02 |
| WBGene00010037 | F54C8.4 | F54C8.4 | 0.34456476 | 6 | 1.46E-02 |
| WBGene00010052 | F54D5.7 | F54D5.7 | 0.23352556 | 15 | 5.55E-03 |
| WBGene00010063 | dhrs-4 | F54F3.4 | 0.26495446 | 5 | 4.01E-02 |
| WBGene00010079 | F55A11.6 | F55A11.6 | 0.33121454 | 20 | 1.30E-04 |
| WBGene00010080 | F55A11.7 | F55A11.7 | 0.23804288 | 12 | 1.00E-02 |
| WBGene00010082 | F55A11.11 | F55A11.11 | 0.54692754 | 6 | 2.17E-03 |
| WBGene00010091 | ssp-35 | F55C5.1 | 0.32856304 | 5 | 2.39E-02 |
| WBGene00010115 | aakb-1 | F55F3.1 | 0.20044674 | 14 | 1.32E-02 |
| WBGene00010117 | nkb-3 | F55F3.3 | 0.17910765 | 12 | 2.73E-02 |
| WBGene00010135 | F55H12.4 | F55H12.4 | 0.20200131 | 7 | 4.63E-02 |
| WBGene00010144 | F56C4.1 | F56C4.1 | 0.34948486 | 13 | 1.05E-03 |
| WBGene00010155 | F56F3.4 | F56F3.4 | 0.94373729 | 7 | 1.71E-05 |
| WBGene00010170 | F56H6.9 | F56H6.9 | 0.26720174 | 13 | 4.70E-03 |
| WBGene00010173 | F56H6.13 | F56H6.13 | 0.19471555 | 9 | 3.53E-02 |
| WBGene00010184 | F57A10.2 | F57A10.2 | 0.90652501 | 3 | 1.83E-03 |
| WBGene00010205 | F57F5.3 | F57F5.3 | 0.34117254 | 8 | 7.35E-03 |
| WBGene00010221 | F57G12.1 | F57G12.1 | 0.38460337 | 9 | 2.90E-03 |
| WBGene00010224 | F58A3.4 | F58A3.4 | 0.32902841 | 5 | 2.38E-02 |
| WBGene00010232 | F58B3.6 | F58B3.6 | 0.27279631 | 7 | 2.18E-02 |
| WBGene00010251 | sta-2 | F58E6.1 | 0.16887578 | 10 | 4.31E-02 |
| WBGene00010260 | ddx-17 | F58E10.3 | 1.02876443 | 13 | 4.50E-09 |
| WBGene00010279 | letm-1 | F58G11.1 | 0.32499102 | 17 | 4.25E-04 |
| WBGene00010289 | F58H1.6 | F58H1.6 | 0.36564393 | 9 | 3.72E-03 |
| WBGene00010290 | F58H1.7 | F58H1.7 | 0.24731407 | 7 | 2.86E-02 |
| WBGene00010291 | F58H10.1 | F58H10.1 | 0.9259519 | 4 | 5.49E-04 |
| WBGene00010315 | F59B2.13 | F59B2.13 | 0.21392088 | 8 | 3.35E-02 |
| WBGene00010317 | idh-1 | F59B8.2 | 0.72596136 | 11 | 6.00E-06 |
| WBGene00010326 | F59C6.5 | F59C6.5 | 0.19048406 | 8 | 4.42E-02 |
| WBGene00010356 | famk-1 | H03A11.1 | 0.35997213 | 18 | 1.29E-04 |
| WBGene00010359 | mam-6 | H03G16.2 | 0.28916932 | 7 | 1.83E-02 |
| WBGene00010387 | H12D21.11 | H12D21.11 | 0.35286229 | 4 | 2.84E-02 |
| WBGene00010409 | H21P03.2 | H21P03.2 | 0.43440593 | 8 | 2.42E-03 |
| WBGene00010418 | H27A22.1 | H27A22.1 | 0.22791494 | 12 | 1.19E-02 |
| WBGene00010419 | atp-1 | H28O16.1 | 0.30360465 | 13 | 2.42E-03 |
| WBGene00010425 | lpin-1 | H37A05.1 | 0.51114834 | 19 | 1.79E-06 |
| WBGene00010433 | H40L08.1 | H40L08.1 | 0.29949376 | 14 | 1.92E-03 |
| WBGene00010434 | H40L08.3 | H40L08.3 | 0.16729795 | 21 | 9.45E-03 |
| WBGene00010450 | metl-18 | K01A11.2 | 0.2733705 | 6 | 2.85E-02 |
| WBGene00010454 | K01B6.3 | K01B6.3 | 0.30047432 | 15 | 1.39E-03 |
| WBGene00010456 | K01C8.1 | K01C8.1 | 0.26089824 | 15 | 3.15E-03 |
| WBGene00010478 | K01G5.5 | K01G5.5 | 0.25816429 | 17 | 2.00E-03 |
| WBGene00010486 | K01H12.4 | K01H12.4 | 0.26365184 | 7 | 2.40E-02 |
| WBGene00010493 | meg-2 | K02B9.2 | 0.29357062 | 16 | 1.19E-03 |
| WBGene00010501 | K02C4.2 | K02C4.2 | 0.42104373 | 3 | 2.81E-02 |
| WBGene00010502 | K02C4.3 | K02C4.3 | 0.14305955 | 23 | 1.51E-02 |
| WBGene00010503 | K02C4.5 | K02C4.5 | 0.48950763 | 13 | 8.23E-05 |

| | | | | | |
|---|---|---|---|---|---|
| WBGene00010525 | K03B8.8 | K03B8.8 | 0.41354967 | 5 | 1.19E-02 |
| WBGene00010556 | rack-1 | K04D7.1 | 0.99195207 | 15 | 8.37E-10 |
| WBGene00010567 | K04G2.9 | K04G2.9 | 1.01008845 | 6 | 2.79E-05 |
| WBGene00010594 | swt-5 | K06A4.4 | 0.34899183 | 14 | 7.34E-04 |
| WBGene00010599 | K06B4.4 | K06B4.4 | 0.24250008 | 8 | 2.38E-02 |
| WBGene00010606 | cyp-13B2 | K06G5.2 | 0.36257454 | 7 | 8.37E-03 |
| WBGene00010619 | K07A1.15 | K07A1.15 | 0.29531649 | 5 | 3.13E-02 |
| WBGene00010621 | egg-6 | K07A12.2 | 0.31595999 | 27 | 1.99E-05 |
| WBGene00010625 | K07C5.2 | K07C5.2 | 0.3481279 | 7 | 9.76E-03 |
| WBGene00010627 | nol-56 | K07C5.4 | 0.59891328 | 14 | 5.68E-06 |
| WBGene00010630 | ttll-15 | K07C5.7 | 0.34665442 | 11 | 2.30E-03 |
| WBGene00010661 | tyr-2 | K08E3.1 | 0.75232225 | 14 | 2.87E-07 |
| WBGene00010664 | dbn-1 | K08E3.4 | 0.46898059 | 13 | 1.20E-04 |
| WBGene00010665 | rml-1 | K08E3.5 | 0.33685814 | 20 | 1.12E-04 |
| WBGene00010666 | K08E4.2 | K08E4.2 | 0.34072997 | 5 | 2.16E-02 |
| WBGene00010667 | K08E4.3 | K08E4.3 | 0.20296977 | 10 | 2.63E-02 |
| WBGene00010673 | K08E7.5 | K08E7.5 | 0.11881371 | 20 | 4.02E-02 |
| WBGene00010677 | gtbp-1 | K08F4.2 | 0.7087009 | 12 | 3.46E-06 |
| WBGene00010682 | K08F8.5 | K08F8.5 | 0.34736113 | 9 | 4.73E-03 |
| WBGene00010690 | K08H2.3 | K08H2.3 | 0.73892296 | 6 | 3.58E-04 |
| WBGene00010697 | uda-1 | K08H10.4 | 0.47466408 | 13 | 1.08E-04 |
| WBGene00010700 | nipi-3 | K09A9.1 | 0.34113617 | 13 | 1.22E-03 |
| WBGene00010704 | K09A11.1 | K09A11.1 | 0.62476805 | 8 | 2.50E-04 |
| WBGene00010705 | cyp-14A1 | K09A11.2 | 0.84176214 | 8 | 1.89E-05 |
| WBGene00010719 | K09E4.1 | K09E4.1 | 0.14049218 | 15 | 3.81E-02 |
| WBGene00010723 | cpg-7 | K09E4.6 | 0.4863799 | 6 | 3.84E-03 |
| WBGene00010730 | ensa-1 | K10C3.2 | 0.30878957 | 8 | 1.08E-02 |
| WBGene00010731 | K10C3.4 | K10C3.4 | 0.20598027 | 31 | 4.81E-04 |
| WBGene00010755 | vglu-2 | K10G9.1 | 0.52318914 | 13 | 4.46E-05 |
| WBGene00010758 | vnut-1 | K10H10.1 | 0.21975722 | 12 | 1.37E-02 |
| WBGene00010759 | cysl-2 | K10H10.2 | 0.47479261 | 8 | 1.49E-03 |
| WBGene00010760 | K10H10.4 | K10H10.4 | 0.66566251 | 5 | 1.53E-03 |
| WBGene00010761 | K10H10.5 | K10H10.5 | 0.64032774 | 2 | 1.71E-02 |
| WBGene00010766 | mrps-27 | K11B4.1 | 0.1924518 | 10 | 3.07E-02 |
| WBGene00010768 | K11D2.1 | K11D2.1 | 0.15820539 | 39 | 1.21E-03 |
| WBGene00010770 | K11D2.4 | K11D2.4 | 0.12595891 | 19 | 3.62E-02 |
| WBGene00010773 | K11E4.1 | K11E4.1 | 1.24314797 | 8 | 1.58E-07 |
| WBGene00010776 | pix-1 | K11E4.4 | 0.60707789 | 26 | 1.19E-09 |
| WBGene00010778 | gpdh-2 | K11H3.1 | 0.3545798 | 18 | 1.47E-04 |
| WBGene00010808 | sepa-1 | M01E5.6 | 0.28130255 | 10 | 8.51E-03 |
| WBGene00010809 | lias-1 | M01F1.3 | 0.35825193 | 7 | 8.76E-03 |
| WBGene00010827 | txt-19 | M02B1.2 | 0.19058751 | 10 | 3.15E-02 |
| WBGene00010833 | M03B6.1 | M03B6.1 | 0.39696398 | 6 | 8.90E-03 |
| WBGene00010834 | mct-3 | M03B6.2 | 0.24049352 | 17 | 3.02E-03 |
| WBGene00010837 | M03B6.5 | M03B6.5 | 0.46513538 | 2 | 3.67E-02 |
| WBGene00010847 | M04B2.4 | M04B2.4 | 0.18619209 | 11 | 2.85E-02 |
| WBGene00010867 | ifbp-1 | M04G12.1 | 0.21037235 | 38 | 1.03E-04 |
| WBGene00010868 | somi-1 | M04G12.4 | 0.15140105 | 19 | 1.88E-02 |

| | | | | | |
|---|---|---|---|---|---|
| WBGene00010871 | M05B5.3 | M05B5.3 | 0.47232172 | 15 | 3.95E-05 |
| WBGene00010883 | M7.7 | M7.7 | 0.59389645 | 15 | 3.18E-06 |
| WBGene00010889 | M18.3 | M18.3 | 0.28326936 | 15 | 1.98E-03 |
| WBGene00010896 | snu-13 | M28.5 | 0.70931192 | 6 | 4.72E-04 |
| WBGene00010937 | M163.5 | M163.5 | 0.21122418 | 9 | 2.84E-02 |
| WBGene00010941 | gss-1 | M176.2 | 0.1832892 | 11 | 2.98E-02 |
| WBGene00010947 | M176.11 | M176.11 | 0.36569929 | 5 | 1.76E-02 |
| WBGene00010988 | metr-1 | R03D7.1 | 0.35576349 | 32 | 7.29E-07 |
| WBGene00011003 | R04B5.5 | R04B5.5 | 0.54692594 | 7 | 1.17E-03 |
| WBGene00011011 | R04D3.3 | R04D3.3 | 0.30898665 | 7 | 1.48E-02 |
| WBGene00011013 | srxa-8 | R04D3.10 | 1.50889789 | 5 | 1.59E-06 |
| WBGene00011015 | R04F11.2 | R04F11.2 | 0.29554902 | 5 | 3.12E-02 |
| WBGene00011017 | R04F11.5 | R04F11.5 | 0.39718522 | 3 | 3.22E-02 |
| WBGene00011030 | R05D7.5 | R05D7.5 | 0.18722677 | 11 | 2.80E-02 |
| WBGene00011042 | R05H10.1 | R05H10.1 | 0.25296306 | 7 | 2.69E-02 |
| WBGene00011056 | R06B9.5 | R06B9.5 | 0.32828936 | 4 | 3.37E-02 |
| WBGene00011059 | R06C1.4 | R06C1.4 | 1.37616007 | 5 | 4.70E-06 |
| WBGene00011071 | R06F6.8 | R06F6.8 | 0.13416421 | 28 | 1.21E-02 |
| WBGene00011074 | R07A4.3 | R07A4.3 | 0.17165377 | 12 | 3.09E-02 |
| WBGene00011088 | kmo-2 | R07B7.4 | 0.38461184 | 12 | 8.39E-04 |
| WBGene00011108 | R07E3.7 | R07E3.7 | 0.14932214 | 25 | 9.82E-03 |
| WBGene00011110 | prdx-3 | R07E5.2 | 0.58068693 | 8 | 4.23E-04 |
| WBGene00011114 | R07E5.6 | R07E5.6 | 0.26193758 | 15 | 3.08E-03 |
| WBGene00011116 | pdcd-2 | R07E5.10 | 0.71441773 | 13 | 1.37E-06 |
| WBGene00011117 | R07E5.11 | R07E5.11 | 0.48531521 | 5 | 6.66E-03 |
| WBGene00011128 | adk-1 | R07H5.8 | 0.6877382 | 11 | 1.09E-05 |
| WBGene00011140 | R08B4.4 | R08B4.4 | 0.17468642 | 11 | 3.41E-02 |
| WBGene00011146 | pde-2 | R08D7.6 | 0.20006901 | 32 | 5.10E-04 |
| WBGene00011159 | chil-17 | R09D1.3 | 0.43041244 | 8 | 2.54E-03 |
| WBGene00011171 | R09E10.1 | R09E10.1 | 0.19615695 | 7 | 4.93E-02 |
| WBGene00011173 | acs-18 | R09E10.3 | 0.24272107 | 14 | 5.80E-03 |
| WBGene00011177 | R09E10.8 | R09E10.8 | 0.37839569 | 4 | 2.38E-02 |
| WBGene00011183 | R09H10.6 | R09H10.6 | 0.43607067 | 7 | 3.82E-03 |
| WBGene00011185 | R10D12.1 | R10D12.1 | 0.45134847 | 10 | 7.32E-04 |
| WBGene00011216 | usp-46 | R10E11.3 | 0.16245811 | 11 | 4.13E-02 |
| WBGene00011230 | nud-2 | R11A5.2 | 0.20095892 | 10 | 2.71E-02 |
| WBGene00011232 | pck-2 | R11A5.4 | 0.37913189 | 23 | 1.06E-05 |
| WBGene00011234 | R11A5.6 | R11A5.6 | 0.45951647 | 2 | 3.76E-02 |
| WBGene00011235 | suro-1 | R11A5.7 | 0.32120924 | 16 | 6.47E-04 |
| WBGene00011239 | pges-2 | R11A8.5 | 0.56936711 | 7 | 9.24E-04 |
| WBGene00011263 | R13H4.5 | R13H4.5 | 0.20379466 | 8 | 3.77E-02 |
| WBGene00011273 | R53.4 | R53.4 | 0.44061093 | 4 | 1.55E-02 |
| WBGene00011278 | R74.2 | R74.2 | 0.35658602 | 6 | 1.30E-02 |
| WBGene00011292 | allo-1 | R102.5 | 0.15124549 | 11 | 4.93E-02 |
| WBGene00011300 | R107.5 | R107.5 | 0.25921627 | 10 | 1.17E-02 |
| WBGene00011304 | mnk-1 | R166.5 | 0.2600615 | 51 | 3.12E-07 |
| WBGene00011305 | hrg-9 | R186.1 | 0.42415168 | 6 | 6.90E-03 |
| WBGene00011325 | T01C3.11 | T01C3.11 | 0.42865735 | 3 | 2.69E-02 |

| | | | | | |
|---|---|---|---|---|---|
| WBGene00011337 | T01G1.2 | T01G1.2 | 0.48281196 | 9 | 7.95E-04 |
| WBGene00011339 | T01G5.1 | T01G5.1 | 0.47533237 | 15 | 3.71E-05 |
| WBGene00011348 | T01H3.2 | T01H3.2 | 0.17091647 | 18 | 1.32E-02 |
| WBGene00011393 | T03D8.6 | T03D8.6 | 0.26518399 | 18 | 1.31E-03 |
| WBGene00011398 | qdpr-1 | T03F6.1 | 0.78200157 | 4 | 1.48E-03 |
| WBGene00011404 | T03F7.7 | T03F7.7 | 0.76438047 | 5 | 6.86E-04 |
| WBGene00011411 | sel-13 | T04A8.10 | 0.50511019 | 4 | 9.96E-03 |
| WBGene00011415 | him-18 | T04A8.15 | 0.18553827 | 14 | 1.77E-02 |
| WBGene00011433 | pde-1 | T04D3.3 | 0.12102521 | 22 | 3.22E-02 |
| WBGene00011434 | T04D3.5 | T04D3.5 | 0.69476835 | 6 | 5.42E-04 |
| WBGene00011435 | T04D3.8 | T04D3.8 | 0.43759171 | 2 | 4.14E-02 |
| WBGene00011436 | T04F3.1 | T04F3.1 | 0.35629593 | 59 | 2.67E-11 |
| WBGene00011440 | sfxn-1.5 | T04F8.1 | 0.27611435 | 8 | 1.59E-02 |
| WBGene00011451 | T04H1.5 | T04H1.5 | 0.40187455 | 7 | 5.50E-03 |
| WBGene00011480 | enpl-1 | T05E11.3 | 0.1438983 | 18 | 2.55E-02 |
| WBGene00011481 | imp-2 | T05E11.5 | 0.28806933 | 9 | 1.03E-02 |
| WBGene00011482 | pigk-1 | T05E11.6 | 0.37022641 | 5 | 1.70E-02 |
| WBGene00011497 | T05F1.13 | T05F1.13 | 0.25171378 | 10 | 1.30E-02 |
| WBGene00011522 | srap-1 | T06D8.1 | 0.80396817 | 102 | 1.25E-43 |
| WBGene00011527 | cchl-1 | T06D8.6 | 0.34752811 | 8 | 6.81E-03 |
| WBGene00011529 | T06D8.9 | T06D8.9 | 0.1910771 | 14 | 1.59E-02 |
| WBGene00011530 | T06D8.10 | T06D8.10 | 0.3195519 | 39 | 3.29E-07 |
| WBGene00011540 | T06E6.10 | T06E6.10 | 0.32172665 | 4 | 3.52E-02 |
| WBGene00011554 | T07A5.1 | T07A5.1 | 0.21403724 | 7 | 4.08E-02 |
| WBGene00011589 | T07F10.3 | T07F10.3 | 0.23255082 | 13 | 8.83E-03 |
| WBGene00011615 | lsd-1 | T08D10.2 | 0.16419591 | 18 | 1.55E-02 |
| WBGene00011635 | mdmh-35 | T09A5.7 | 0.78044785 | 6 | 2.42E-04 |
| WBGene00011643 | slc-17.9 | T09B9.2 | 0.25218306 | 12 | 7.91E-03 |
| WBGene00011662 | T09F3.2 | T09F3.2 | 0.53344225 | 7 | 1.35E-03 |
| WBGene00011678 | T10B9.9 | T10B9.9 | 0.82400108 | 6 | 1.61E-04 |
| WBGene00011679 | ucr-2.2 | T10B10.2 | 0.44492212 | 10 | 8.03E-04 |
| WBGene00011694 | T10G3.3 | T10G3.3 | 0.81578225 | 11 | 1.47E-06 |
| WBGene00011698 | cyp-34A1 | T10H4.10 | 0.31205761 | 10 | 5.46E-03 |
| WBGene00011729 | set-16 | T12D8.1 | 0.15376046 | 53 | 2.71E-04 |
| WBGene00011735 | hip-1 | T12D8.8 | 0.19703824 | 36 | 2.85E-04 |
| WBGene00011736 | T12D8.9 | T12D8.9 | 0.20189663 | 21 | 3.56E-03 |
| WBGene00011737 | sqst-1 | T12G3.1 | 0.4964285 | 19 | 2.62E-06 |
| WBGene00011738 | T12G3.2 | T12G3.2 | 0.11154101 | 38 | 1.39E-02 |
| WBGene00011751 | T13F3.4 | T13F3.4 | 0.30753624 | 5 | 2.83E-02 |
| WBGene00011756 | ctg-2 | T13H5.2 | 0.28003587 | 10 | 8.66E-03 |
| WBGene00011760 | T13H5.6 | T13H5.6 | 0.1336171 | 20 | 2.70E-02 |
| WBGene00011763 | T14B1.1 | T14B1.1 | 0.30402583 | 13 | 2.41E-03 |
| WBGene00011768 | oac-46 | T14D7.2 | 0.48164419 | 14 | 5.56E-05 |
| WBGene00011770 | T14G8.2 | T14G8.2 | 0.26544065 | 6 | 3.07E-02 |
| WBGene00011773 | ttr-53 | T14G10.3 | 0.32389155 | 4 | 3.47E-02 |
| WBGene00011775 | copg-1 | T14G10.5 | 0.13190198 | 23 | 2.12E-02 |
| WBGene00011776 | pigs-1 | T14G10.7 | 0.22678438 | 10 | 1.87E-02 |
| WBGene00011777 | T14G10.8 | T14G10.8 | 0.2684118 | 5 | 3.89E-02 |

| | | | | | |
|---|---|---|---|---|---|
| WBGene00011814 | gtf-2H2C | T16H12.4 | 0.25152121 | 8 | 2.14E-02 |
| WBGene00011820 | T18D3.1 | T18D3.1 | 0.17085496 | 17 | 1.52E-02 |
| WBGene00011828 | nepr-1 | T19A6.3 | 0.23545523 | 8 | 2.59E-02 |
| WBGene00011831 | T19B10.2 | T19B10.2 | 0.70700476 | 9 | 4.15E-05 |
| WBGene00011832 | bgal-1 | T19B10.3 | 0.25714282 | 16 | 2.64E-03 |
| WBGene00011849 | T19H5.4 | T19H5.4 | 0.80018332 | 4 | 1.31E-03 |
| WBGene00011850 | T20B3.1 | T20B3.1 | 0.40548774 | 15 | 1.58E-04 |
| WBGene00011867 | chc-1 | T20G5.1 | 0.09807609 | 37 | 2.86E-02 |
| WBGene00011871 | T20G5.9 | T20G5.9 | 0.32222673 | 6 | 1.80E-02 |
| WBGene00011877 | T21B4.3 | T21B4.3 | 0.23527936 | 8 | 2.59E-02 |
| WBGene00011883 | mrpl-50 | T21B10.1 | 0.33384612 | 8 | 8.02E-03 |
| WBGene00011884 | enol-1 | T21B10.2 | 0.40796544 | 14 | 2.33E-04 |
| WBGene00011885 | T21B10.3 | T21B10.3 | 0.11342607 | 24 | 3.48E-02 |
| WBGene00011891 | del-6 | T21C9.3 | 0.35337791 | 15 | 4.64E-04 |
| WBGene00011895 | T21C9.9 | T21C9.9 | 0.37878094 | 8 | 4.69E-03 |
| WBGene00011899 | nlp-68 | T21C12.3 | 0.31872995 | 4 | 3.60E-02 |
| WBGene00011908 | pash-1 | T22A3.5 | 0.12125107 | 21 | 3.47E-02 |
| WBGene00011935 | scrm-1 | T22H2.5 | 0.21946437 | 6 | 4.72E-02 |
| WBGene00011953 | ppm-2 | T23F11.1 | 0.25391604 | 11 | 9.85E-03 |
| WBGene00011959 | nyn-1 | T23G4.3 | 0.26193565 | 11 | 8.68E-03 |
| WBGene00011976 | chmp-7 | T24B8.2 | 0.65246788 | 10 | 4.02E-05 |
| WBGene00011977 | T24B8.3 | T24B8.3 | 0.55680952 | 8 | 5.63E-04 |
| WBGene00012004 | dyrb-1 | T24H10.6 | 0.33884554 | 3 | 4.47E-02 |
| WBGene00012005 | jun-1 | T24H10.7 | 0.1539598 | 46 | 6.33E-04 |
| WBGene00012007 | T25B9.1 | T25B9.1 | 0.45751153 | 7 | 3.04E-03 |
| WBGene00012033 | T26C5.3 | T26C5.3 | 0.23609427 | 31 | 1.41E-04 |
| WBGene00012085 | T27D12.1 | T27D12.1 | 0.6916428 | 12 | 4.62E-06 |
| WBGene00012106 | T27F6.7 | T27F6.7 | 0.21664146 | 11 | 1.77E-02 |
| WBGene00012107 | T27F6.8 | T27F6.8 | 0.91727094 | 10 | 8.81E-07 |
| WBGene00012118 | grsp-2 | T28C6.1 | 0.63428414 | 18 | 1.56E-07 |
| WBGene00012120 | T28C6.5 | T28C6.5 | 0.59689612 | 7 | 6.89E-04 |
| WBGene00012149 | VF13D12L.3 | VF13D12L.3 | 0.19242935 | 32 | 7.04E-04 |
| WBGene00012152 | cnc-10 | VK10D6R.1 | 0.83944245 | 3 | 2.67E-03 |
| WBGene00012169 | ttbk-4 | W01B6.2 | 0.19595079 | 11 | 2.44E-02 |
| WBGene00012178 | glb-27 | W01C9.5 | 0.12005321 | 22 | 3.31E-02 |
| WBGene00012180 | W01D2.3 | W01D2.3 | 0.35886485 | 12 | 1.30E-03 |
| WBGene00012186 | mlt-11 | W01F3.3 | 0.82106449 | 49 | 1.83E-22 |
| WBGene00012219 | W03C9.1 | W03C9.1 | 0.78072364 | 9 | 1.57E-05 |
| WBGene00012220 | W03C9.2 | W03C9.2 | 0.26368783 | 8 | 1.85E-02 |
| WBGene00012222 | W03C9.6 | W03C9.6 | 0.57150305 | 21 | 1.04E-07 |
| WBGene00012224 | W03G11.2 | W03G11.2 | 0.42046273 | 5 | 1.13E-02 |
| WBGene00012256 | lpr-5 | W04G3.2 | 0.25456893 | 5 | 4.36E-02 |
| WBGene00012261 | lpr-3 | W04G3.8 | 0.79814093 | 8 | 3.17E-05 |
| WBGene00012272 | dhc-4 | W05B2.4 | 0.17647323 | 92 | 1.55E-07 |
| WBGene00012317 | ztf-6 | W06H12.1 | 0.23917372 | 18 | 2.48E-03 |
| WBGene00012322 | W07A12.4 | W07A12.4 | 0.12009651 | 21 | 3.59E-02 |
| WBGene00012324 | rhy-1 | W07A12.7 | 0.19462515 | 10 | 2.97E-02 |
| WBGene00012330 | zip-3 | W07G1.3 | 0.15282608 | 15 | 2.95E-02 |

| | | | | | |
|---|---|---|---|---|---|
| WBGene00012336 | W07G4.2 | W07G4.2 | 0.46469406 | 6 | 4.71E-03 |
| WBGene00012337 | scyl-1 | W07G4.3 | 0.2393071 | 23 | 7.79E-04 |
| WBGene00012343 | casc-3 | W08E3.2 | 0.56702243 | 17 | 1.54E-06 |
| WBGene00012347 | W08G11.3 | W08G11.3 | 0.17549239 | 17 | 1.37E-02 |
| WBGene00012348 | pptr-1 | W08G11.4 | 0.20832279 | 16 | 7.73E-03 |
| WBGene00012351 | W09C5.1 | W09C5.1 | 0.28765137 | 5 | 3.33E-02 |
| WBGene00012358 | W09D6.5 | W09D6.5 | 0.22412937 | 6 | 4.52E-02 |
| WBGene00012376 | nduf-7 | W10D5.2 | 0.12908784 | 17 | 4.02E-02 |
| WBGene00012397 | Y6E2A.1 | Y6E2A.1 | 0.26231506 | 5 | 4.09E-02 |
| WBGene00012405 | ztf-25 | Y6G8.3 | 0.5218557 | 9 | 4.75E-04 |
| WBGene00012408 | Y7A5A.2 | Y7A5A.2 | 0.28022898 | 4 | 4.69E-02 |
| WBGene00012429 | Y11D7A.5 | Y11D7A.5 | 0.69351998 | 7 | 2.46E-04 |
| WBGene00012436 | flh-3 | Y11D7A.13 | 0.23558624 | 12 | 1.05E-02 |
| WBGene00012440 | nlp-53 | Y12A6A.2 | 0.23100253 | 6 | 4.24E-02 |
| WBGene00012443 | Y15E3A.4 | Y15E3A.4 | 0.35440946 | 13 | 9.62E-04 |
| WBGene00012463 | nadk-2 | Y17G7B.10 | 0.22056149 | 43 | 2.21E-05 |
| WBGene00012471 | Y17G7B.20 | Y17G7B.20 | 0.49499539 | 10 | 3.90E-04 |
| WBGene00012472 | Y17G7B.21 | Y17G7B.21 | 0.61885785 | 8 | 2.69E-04 |
| WBGene00012483 | coa-5 | Y18D10A.16 | 0.25672958 | 6 | 3.33E-02 |
| WBGene00012484 | car-1 | Y18D10A.17 | 0.44632643 | 7 | 3.43E-03 |
| WBGene00012535 | Y37A1A.2 | Y37A1A.2 | 0.27154815 | 13 | 4.34E-03 |
| WBGene00012540 | Y37A1B.7 | Y37A1B.7 | 0.16093805 | 10 | 4.83E-02 |
| WBGene00012546 | Y37D8A.4 | Y37D8A.4 | 0.61067386 | 7 | 5.95E-04 |
| WBGene00012553 | cox-5A | Y37D8A.14 | 0.41414713 | 4 | 1.86E-02 |
| WBGene00012592 | Y38E10A.14 | Y38E10A.14 | 0.51971433 | 14 | 2.65E-05 |
| WBGene00012593 | nspe-7 | Y38E10A.15 | 0.48690048 | 3 | 1.94E-02 |
| WBGene00012607 | Y38F1A.4 | Y38F1A.4 | 0.43417559 | 10 | 9.37E-04 |
| WBGene00012670 | cpg-23 | Y39B6A.8 | 0.29115331 | 8 | 1.33E-02 |
| WBGene00012710 | Y39C12A.9 | Y39C12A.9 | 0.64714246 | 5 | 1.78E-03 |
| WBGene00012717 | Y39E4B.6 | Y39E4B.6 | 0.35397685 | 17 | 2.17E-04 |
| WBGene00012721 | Y39E4B.13 | Y39E4B.13 | 0.46326518 | 7 | 2.86E-03 |
| WBGene00012754 | Y41C4A.7 | Y41C4A.7 | 0.42616694 | 5 | 1.08E-02 |
| WBGene00012762 | Y41E3.1 | Y41E3.1 | 0.17964781 | 21 | 6.67E-03 |
| WBGene00012763 | atln-2 | Y41E3.3 | 0.11930107 | 26 | 2.46E-02 |
| WBGene00012768 | eef-1B.2 | Y41E3.10 | 0.46161682 | 23 | 8.35E-07 |
| WBGene00012769 | hrpu-1 | Y41E3.11 | 0.29760666 | 41 | 5.47E-07 |
| WBGene00012791 | Y43D4A.5 | Y43D4A.5 | 0.29450693 | 40 | 8.72E-07 |
| WBGene00012796 | Y43F4A.1 | Y43F4A.1 | 0.26680261 | 16 | 2.14E-03 |
| WBGene00012886 | Y45F10D.6 | Y45F10D.6 | 0.34542668 | 3 | 4.31E-02 |
| WBGene00012917 | Y46G5A.29 | Y46G5A.29 | 0.3255335 | 18 | 2.99E-04 |
| WBGene00012942 | Y47D3B.6 | Y47D3B.6 | 0.54258375 | 4 | 7.70E-03 |
| WBGene00012947 | Y47H9C.1 | Y47H9C.1 | 0.30207354 | 4 | 4.03E-02 |
| WBGene00012966 | exos-1 | Y48A6B.5 | 0.26835337 | 6 | 2.98E-02 |
| WBGene00012972 | rsa-2 | Y48A6B.11 | 0.26740194 | 22 | 4.28E-04 |
| WBGene00012978 | Y48B6A.1 | Y48B6A.1 | 0.24673976 | 14 | 5.37E-03 |
| WBGene00012999 | rpoa-1 | Y48E1A.1 | 0.1485982 | 71 | 4.73E-05 |
| WBGene00013007 | Y48E1B.8 | Y48E1B.8 | 0.29073076 | 8 | 1.34E-02 |
| WBGene00013008 | clec-146 | Y48E1B.9 | 0.4611895 | 5 | 8.10E-03 |

| | | | | | |
|---|---|---|---|---|---|
| WBGene00013013 | clec-145 | Y48E1B.16 | 0.32657505 | 7 | 1.23E-02 |
| WBGene00013025 | vha-13 | Y49A3A.2 | 0.48996643 | 17 | 9.21E-06 |
| WBGene00013034 | tat-1 | Y49E10.11 | 0.09769964 | 30 | 4.21E-02 |
| WBGene00013106 | set-26 | Y51H4A.12 | 0.10508517 | 33 | 2.61E-02 |
| WBGene00013128 | dxbp-1 | Y52B11A.9 | 0.18799576 | 12 | 2.35E-02 |
| WBGene00013139 | Y53C12A.3 | Y53C12A.3 | 0.45413977 | 6 | 5.20E-03 |
| WBGene00013140 | mop-25.2 | Y53C12A.4 | 0.2673258 | 10 | 1.04E-02 |
| WBGene00013146 | Y53C12B.7 | Y53C12B.7 | 0.20397706 | 7 | 4.54E-02 |
| WBGene00013187 | npr-34 | Y54E2A.1 | 0.17510098 | 14 | 2.16E-02 |
| WBGene00013192 | Y54E2A.7 | Y54E2A.7 | 0.30565078 | 8 | 1.12E-02 |
| WBGene00013217 | ddl-3 | Y54G11A.8 | 0.18402215 | 8 | 4.78E-02 |
| WBGene00013242 | Y56A3A.30 | Y56A3A.30 | 0.23200809 | 23 | 9.76E-04 |
| WBGene00013244 | Y56A3A.33 | Y56A3A.33 | 0.14534256 | 16 | 3.08E-02 |
| WBGene00013255 | tric-1B.1 | Y57A10A.10 | 0.24698591 | 40 | 1.04E-05 |
| WBGene00013260 | rsr-2 | Y57A10A.19 | 0.21887617 | 18 | 4.07E-03 |
| WBGene00013266 | Y57A10A.26 | Y57A10A.26 | 0.48214798 | 22 | 7.58E-07 |
| WBGene00013287 | Y57A10C.9 | Y57A10C.9 | 0.37569292 | 10 | 2.18E-03 |
| WBGene00013297 | Y57G11B.5 | Y57G11B.5 | 0.27778759 | 7 | 2.07E-02 |
| WBGene00013301 | Y57G11C.3 | Y57G11C.3 | 0.27873739 | 7 | 2.05E-02 |
| WBGene00013302 | vti-1 | Y57G11C.4 | 0.41463638 | 8 | 3.06E-03 |
| WBGene00013304 | ptp-5.2 | Y57G11C.6 | 0.19181586 | 15 | 1.32E-02 |
| WBGene00013311 | sec-61.A | Y57G11C.15 | 1.14465322 | 15 | 3.54E-11 |
| WBGene00013312 | hhat-2 | Y57G11C.17 | 0.43013718 | 7 | 4.07E-03 |
| WBGene00013324 | mrps-7 | Y57G11C.34 | 0.31670681 | 7 | 1.36E-02 |
| WBGene00013352 | lon-8 | Y59A8B.20 | 0.24035739 | 14 | 6.08E-03 |
| WBGene00013385 | Y62E10A.19 | Y62E10A.19 | 0.28974467 | 13 | 3.12E-03 |
| WBGene00013418 | Y65A5A.1 | Y65A5A.1 | 1.78789374 | 6 | 1.86E-08 |
| WBGene00013436 | Y66D12A.10 | Y66D12A.10 | 0.36429482 | 7 | 8.22E-03 |
| WBGene00013438 | ztf-29 | Y66D12A.12 | 0.93572371 | 11 | 2.23E-07 |
| WBGene00013439 | Y66D12A.13 | Y66D12A.13 | 0.21342367 | 12 | 1.52E-02 |
| WBGene00013458 | clc-8 | Y67A10A.9 | 0.17171885 | 24 | 5.39E-03 |
| WBGene00013461 | Y67H2A.2 | Y67H2A.2 | 0.2184439 | 27 | 6.54E-04 |
| WBGene00013462 | micu-1 | Y67H2A.4 | 0.43768067 | 9 | 1.44E-03 |
| WBGene00013463 | kdp-1 | Y67H2A.5 | 0.5012604 | 6 | 3.34E-03 |
| WBGene00013489 | col-42 | Y69H2.14 | 1.46580895 | 6 | 3.85E-07 |
| WBGene00013499 | Y70G10A.3 | Y70G10A.3 | 0.50803992 | 17 | 6.05E-06 |
| WBGene00013507 | Y71A12B.10 | Y71A12B.10 | 0.26836584 | 11 | 7.85E-03 |
| WBGene00013523 | Y73F8A.14 | Y73F8A.14 | 0.35713434 | 3 | 4.03E-02 |
| WBGene00013531 | Y73F8A.26 | Y73F8A.26 | 0.23113954 | 6 | 4.23E-02 |
| WBGene00013543 | Y75B8A.6 | Y75B8A.6 | 0.13732821 | 13 | 5.00E-02 |
| WBGene00013556 | Y75B8A.23 | Y75B8A.23 | 0.70035446 | 3 | 5.83E-03 |
| WBGene00013560 | zip-12 | Y75B8A.29 | 0.34541525 | 10 | 3.37E-03 |
| WBGene00013568 | Y75B12B.3 | Y75B12B.3 | 0.56343441 | 2 | 2.39E-02 |
| WBGene00013573 | Y75B12B.11 | Y75B12B.11 | 0.97550781 | 3 | 1.24E-03 |
| WBGene00013575 | acs-5 | Y76A2B.3 | 0.23948278 | 13 | 7.79E-03 |
| WBGene00013577 | Y76A2B.5 | Y76A2B.5 | 0.3172785 | 5 | 2.62E-02 |
| WBGene00013578 | scav-2 | Y76A2B.6 | 0.34540353 | 14 | 7.87E-04 |
| WBGene00013591 | gcn-2 | Y81G3A.3 | 0.161584 | 37 | 1.33E-03 |

| | | | | | |
|---|---|---|---|---|---|
| WBGene00013599 | eipr-1 | Y87G2A.11 | 0.12856466 | 15 | 4.88E-02 |
| WBGene00013639 | Y105C5A.15 | Y105C5A.15 | 0.0932073 | 56 | 1.48E-02 |
| WBGene00013650 | Y105C5B.9 | Y105C5B.9 | 2.39104581 | 8 | 1.82E-13 |
| WBGene00013672 | catp-1 | Y105E8A.12 | 0.39777052 | 66 | 5.65E-14 |
| WBGene00013688 | riok-2 | Y105E8B.3 | 0.22436614 | 12 | 1.27E-02 |
| WBGene00013709 | csnk-1 | Y106G6E.6 | 0.13255822 | 17 | 3.71E-02 |
| WBGene00013717 | madf-10 | Y106G6H.4 | 0.76840752 | 6 | 2.71E-04 |
| WBGene00013736 | gtf-2A2 | Y111B2A.13 | 0.45990977 | 5 | 8.19E-03 |
| WBGene00013766 | prmt-1 | Y113G7B.17 | 0.37945323 | 9 | 3.10E-03 |
| WBGene00013858 | ssp-34 | ZC168.6 | 1.41189185 | 4 | 1.93E-05 |
| WBGene00013859 | ZC247.1 | ZC247.1 | 0.22213466 | 82 | 5.87E-09 |
| WBGene00013866 | cbs-1 | ZC373.1 | 0.61114475 | 20 | 6.80E-08 |
| WBGene00013871 | gnrr-3 | ZC374.1 | 0.43709596 | 12 | 3.45E-04 |
| WBGene00013878 | atfs-1 | ZC376.7 | 0.43773291 | 17 | 3.10E-05 |
| WBGene00013883 | npr-13 | ZC412.1 | 0.32970077 | 17 | 3.81E-04 |
| WBGene00013886 | ZC412.5 | ZC412.5 | 0.32282876 | 4 | 3.50E-02 |
| WBGene00013902 | ZC455.1 | ZC455.1 | 0.23822308 | 32 | 1.02E-04 |
| WBGene00013920 | pssy-1 | ZC506.3 | 0.22889239 | 15 | 6.11E-03 |
| WBGene00013923 | ghi-1 | ZK20.1 | 0.32916164 | 7 | 1.19E-02 |
| WBGene00013924 | rad-23 | ZK20.3 | 0.38661116 | 5 | 1.49E-02 |
| WBGene00013926 | nep-1 | ZK20.6 | 0.14343292 | 21 | 1.85E-02 |
| WBGene00013969 | tep-1 | ZK337.1 | 0.39589863 | 25 | 2.68E-06 |
| WBGene00013980 | dos-1 | ZK507.4 | 0.2781954 | 5 | 3.60E-02 |
| WBGene00013985 | sec-16A.1 | ZK512.5 | 0.16182075 | 28 | 4.34E-03 |
| WBGene00013998 | gtf-2E1 | ZK550.4 | 0.42218528 | 8 | 2.80E-03 |
| WBGene00014001 | pyk-2 | ZK593.1 | 0.15431506 | 12 | 4.15E-02 |
| WBGene00014012 | riok-3 | ZK632.3 | 0.26099113 | 16 | 2.43E-03 |
| WBGene00014025 | asna-1 | ZK637.5 | 0.33968234 | 7 | 1.07E-02 |
| WBGene00014046 | clec-60 | ZK666.6 | 0.25477835 | 6 | 3.39E-02 |
| WBGene00014051 | spv-1 | ZK669.1 | 0.12145907 | 50 | 3.16E-03 |
| WBGene00014052 | ZK669.2 | ZK669.2 | 0.34797287 | 9 | 4.69E-03 |
| WBGene00014058 | ZK673.2 | ZK673.2 | 0.35572085 | 8 | 6.18E-03 |
| WBGene00014070 | srxa-9 | ZK678.4 | 0.32683824 | 11 | 3.14E-03 |
| WBGene00014075 | dhhc-4 | ZK757.4 | 0.25140831 | 15 | 3.83E-03 |
| WBGene00014087 | ZK809.5 | ZK809.5 | 0.12087056 | 22 | 3.23E-02 |
| WBGene00014089 | ZK822.1 | ZK822.1 | 0.15676561 | 19 | 1.64E-02 |
| WBGene00014091 | ZK822.4 | ZK822.4 | 0.7209805 | 8 | 7.96E-05 |
| WBGene00014092 | ZK822.5 | ZK822.5 | 0.17572535 | 14 | 2.14E-02 |
| WBGene00014093 | ZK829.1 | ZK829.1 | 0.2192285 | 13 | 1.13E-02 |
| WBGene00014095 | gdh-1 | ZK829.4 | 0.64809647 | 9 | 9.01E-05 |
| WBGene00014098 | ogdh-2 | ZK836.2 | 0.30964753 | 43 | 1.52E-07 |
| WBGene00014103 | best-26 | ZK849.5 | 0.2567679 | 14 | 4.42E-03 |
| WBGene00014115 | gld-4 | ZK858.1 | 0.15165532 | 14 | 3.41E-02 |
| WBGene00014128 | ZK892.4 | ZK892.4 | 0.24081278 | 7 | 3.06E-02 |
| WBGene00014129 | ZK892.5 | ZK892.5 | 0.59044082 | 4 | 5.53E-03 |
| WBGene00014139 | nstp-5 | ZK896.9 | 0.18989445 | 12 | 2.27E-02 |
| WBGene00014148 | ZK909.3 | ZK909.3 | 0.27650088 | 8 | 1.59E-02 |
| WBGene00014171 | nep-26 | ZK970.1 | 0.10971118 | 55 | 4.82E-03 |

| | | | | | |
|---|---|---|---|---|---|
| WBGene00014172 | clpp-1 | ZK970.2 | 0.5748688 | 13 | 1.74E-05 |
| WBGene00014173 | ZK970.7 | ZK970.7 | 0.61742461 | 4 | 4.60E-03 |
| WBGene00014202 | mmcm-1 | ZK1058.1 | 0.24935682 | 22 | 7.29E-04 |
| WBGene00014235 | ZK1225.1 | ZK1225.1 | 0.25877946 | 8 | 1.96E-02 |
| WBGene00014246 | ZK1307.3 | ZK1307.3 | 0.87479408 | 7 | 3.56E-05 |
| WBGene00014258 | ZK1320.9 | ZK1320.9 | 0.3248985 | 7 | 1.25E-02 |
| WBGene00014262 | ZK1321.4 | ZK1321.4 | 0.48603728 | 14 | 5.10E-05 |
| WBGene00014300 | D2023.1 | D2023.1 | 0.28428035 | 36 | 4.68E-06 |
| WBGene00014698 | C37A5.3 | C37A5.3 | 0.8686257 | 3 | 2.26E-03 |
| WBGene00014699 | C37A5.5 | C37A5.5 | 0.58945908 | 1 | 4.16E-02 |
| WBGene00014965 | Y106G6D.5 | Y106G6D.5 | 0.23419743 | 19 | 2.23E-03 |
| WBGene00015002 | B0034.1 | B0034.1 | 0.345102 | 15 | 5.51E-04 |
| WBGene00015007 | ain-2 | B0041.2 | 0.24045256 | 29 | 1.87E-04 |
| WBGene00015008 | lmd-2 | B0041.3 | 0.35366209 | 5 | 1.95E-02 |
| WBGene00015021 | nfs-1 | B0205.6 | 0.34131775 | 11 | 2.50E-03 |
| WBGene00015040 | cyp-34A5 | B0213.10 | 0.22161641 | 11 | 1.63E-02 |
| WBGene00015044 | cyp-34A9 | B0213.15 | 0.45528019 | 8 | 1.89E-03 |
| WBGene00015046 | nlp-34 | B0213.17 | 0.5578963 | 2 | 2.45E-02 |
| WBGene00015056 | B0222.5 | B0222.5 | 0.24036189 | 8 | 2.44E-02 |
| WBGene00015059 | B0228.1 | B0228.1 | 0.51619436 | 4 | 9.23E-03 |
| WBGene00015064 | B0228.7 | B0228.7 | 0.57108567 | 6 | 1.73E-03 |
| WBGene00015138 | B0310.2 | B0310.2 | 0.43801849 | 14 | 1.30E-04 |
| WBGene00015141 | ugt-46 | B0310.5 | 0.16154484 | 12 | 3.67E-02 |
| WBGene00015143 | rbm-26 | B0336.3 | 0.35426056 | 22 | 3.30E-05 |
| WBGene00015147 | B0336.7 | B0336.7 | 0.31051735 | 20 | 2.28E-04 |
| WBGene00015148 | hpo-28 | B0336.11 | 0.1990699 | 14 | 1.36E-02 |
| WBGene00015156 | cwf-19L2 | B0361.2 | 0.20608581 | 68 | 4.44E-07 |
| WBGene00015163 | B0361.9 | B0361.9 | 0.79900618 | 5 | 5.17E-04 |
| WBGene00015178 | B0416.2 | B0416.2 | 0.32765654 | 5 | 2.40E-02 |
| WBGene00015207 | luc-7L | B0495.8 | 0.45739103 | 14 | 8.91E-05 |
| WBGene00015208 | B0495.9 | B0495.9 | 0.36657697 | 4 | 2.59E-02 |
| WBGene00015235 | cdc-26 | B0511.9 | 0.24881342 | 8 | 2.21E-02 |
| WBGene00015285 | gmeb-1 | C01B12.2 | 0.74313922 | 8 | 6.11E-05 |
| WBGene00015291 | C01B12.8 | C01B12.8 | 0.3394089 | 3 | 4.45E-02 |
| WBGene00015293 | C01C4.3 | C01C4.3 | 0.13478942 | 46 | 1.99E-03 |
| WBGene00015294 | C01C10.2 | C01C10.2 | 0.26128289 | 6 | 3.19E-02 |
| WBGene00015320 | C02B8.1 | C02B8.1 | 0.31760132 | 6 | 1.88E-02 |
| WBGene00015332 | tyr-1 | C02C2.1 | 0.50198013 | 10 | 3.52E-04 |
| WBGene00015335 | acdh-6 | C02D5.1 | 0.19966733 | 12 | 1.92E-02 |
| WBGene00015348 | C02F5.5 | C02F5.5 | 0.40935401 | 5 | 1.24E-02 |
| WBGene00015357 | C02F12.8 | C02F12.8 | 0.20165791 | 12 | 1.86E-02 |
| WBGene00015373 | C03B1.2 | C03B1.2 | 0.69596797 | 10 | 2.15E-05 |
| WBGene00015410 | tbc-6 | C04A2.1 | 0.5268908 | 14 | 2.31E-05 |
| WBGene00015430 | C04E7.3 | C04E7.3 | 0.16881748 | 37 | 9.35E-04 |
| WBGene00015435 | C04E12.5 | C04E12.5 | 0.28960559 | 12 | 4.19E-03 |
| WBGene00015458 | C04G6.10 | C04G6.10 | 0.97666965 | 3 | 1.23E-03 |
| WBGene00015464 | C05C8.7 | C05C8.7 | 0.6605271 | 13 | 3.66E-06 |
| WBGene00015467 | basl-1 | C05D2.3 | 0.2358738 | 16 | 4.22E-03 |

| | | | | | |
|---|---|---|---|---|---|
| WBGene00015470 | wdr-37 | C05D2.10 | 0.24946992 | 13 | 6.49E-03 |
| WBGene00015474 | C05D9.7 | C05D9.7 | 0.22729853 | 8 | 2.85E-02 |
| WBGene00015484 | atgl-1 | C05D11.7 | 0.39774193 | 10 | 1.59E-03 |
| WBGene00015496 | C05E11.7 | C05E11.7 | 0.19219669 | 15 | 1.31E-02 |
| WBGene00015508 | mvb-12 | C06A6.3 | 0.41505844 | 5 | 1.18E-02 |
| WBGene00015512 | mthf-1 | C06A8.1 | 0.42590235 | 24 | 1.57E-06 |
| WBGene00015514 | nlp-77 | C06A8.3 | 0.52976633 | 3 | 1.52E-02 |
| WBGene00015529 | C06E2.5 | C06E2.5 | 0.8305938 | 3 | 2.80E-03 |
| WBGene00015545 | C06G1.1 | C06G1.1 | 0.42499068 | 11 | 6.73E-04 |
| WBGene00015547 | ain-1 | C06G1.4 | 0.62252351 | 17 | 4.24E-07 |
| WBGene00015558 | C06G4.4 | C06G4.4 | 0.74503222 | 6 | 3.38E-04 |
| WBGene00015561 | C07A12.7 | C07A12.7 | 0.20615631 | 12 | 1.72E-02 |
| WBGene00015615 | fbxc-48 | C08G5.2 | 0.2520382 | 8 | 2.12E-02 |
| WBGene00015636 | C09D4.6 | C09D4.6 | 0.3338526 | 4 | 3.24E-02 |
| WBGene00015646 | mlt-10 | C09E8.3 | 0.39716995 | 17 | 7.94E-05 |
| WBGene00015647 | C09F5.1 | C09F5.1 | 0.1372736 | 16 | 3.68E-02 |
| WBGene00015676 | mct-6 | C10E2.6 | 0.12383377 | 18 | 4.17E-02 |
| WBGene00015684 | C10G8.8 | C10G8.8 | 0.23039228 | 16 | 4.76E-03 |
| WBGene00015740 | C13E3.1 | C13E3.1 | 0.20660104 | 20 | 3.76E-03 |
| WBGene00015741 | C13F10.1 | C13F10.1 | 0.25356542 | 7 | 2.67E-02 |
| WBGene00015753 | C14B9.3 | C14B9.3 | 0.3284516 | 6 | 1.70E-02 |
| WBGene00015759 | C14C6.5 | C14C6.5 | 0.28977409 | 5 | 3.27E-02 |
| WBGene00015769 | C14C11.7 | C14C11.7 | 0.87311755 | 5 | 2.83E-04 |
| WBGene00015781 | rml-3 | C14F11.6 | 0.4641939 | 4 | 1.32E-02 |
| WBGene00015786 | C15B12.4 | C15B12.4 | 0.26673221 | 15 | 2.79E-03 |
| WBGene00015795 | C15F1.2 | C15F1.2 | 0.33110743 | 34 | 1.03E-06 |
| WBGene00015796 | C15F1.5 | C15F1.5 | 0.18081049 | 12 | 2.65E-02 |
| WBGene00015802 | kynu-1 | C15H9.7 | 0.26808419 | 10 | 1.03E-02 |
| WBGene00015809 | znf-622 | C16A3.4 | 0.23889958 | 8 | 2.48E-02 |
| WBGene00015841 | skpo-2 | C16C8.2 | 0.51635122 | 31 | 1.51E-09 |
| WBGene00015858 | C16D9.3 | C16D9.3 | 0.33391625 | 6 | 1.61E-02 |
| WBGene00015859 | C16D9.4 | C16D9.4 | 0.34800002 | 6 | 1.41E-02 |
| WBGene00015860 | C16D9.5 | C16D9.5 | 0.29643797 | 12 | 3.74E-03 |
| WBGene00015861 | C16D9.6 | C16D9.6 | 0.23346186 | 9 | 2.12E-02 |
| WBGene00015866 | C16E9.2 | C16E9.2 | 0.16850257 | 9 | 4.99E-02 |
| WBGene00015887 | C17C3.1 | C17C3.1 | 0.45159992 | 13 | 1.64E-04 |
| WBGene00015896 | C17C3.15 | C17C3.15 | 0.99446648 | 6 | 3.24E-05 |
| WBGene00015915 | C17G10.1 | C17G10.1 | 0.18118961 | 9 | 4.22E-02 |
| WBGene00015919 | C17G10.7 | C17G10.7 | 0.54897678 | 5 | 3.96E-03 |
| WBGene00015926 | C17H11.6 | C17H11.6 | 0.17113949 | 42 | 4.05E-04 |
| WBGene00015943 | tiar-1 | C18A3.5 | 0.42578316 | 11 | 6.65E-04 |
| WBGene00015993 | C18H7.1 | C18H7.1 | 0.84533563 | 14 | 4.70E-08 |
| WBGene00015994 | C18H7.4 | C18H7.4 | 0.32266871 | 7 | 1.28E-02 |
| WBGene00016006 | fln-2 | C23F12.1 | 0.25557702 | 96 | 5.24E-12 |
| WBGene00016011 | C23G10.2 | C23G10.2 | 0.40670767 | 3 | 3.05E-02 |
| WBGene00016013 | ugt-66 | C23G10.6 | 0.45999539 | 14 | 8.47E-05 |
| WBGene00016018 | C23H3.2 | C23H3.2 | 0.3484635 | 8 | 6.73E-03 |
| WBGene00016020 | sptl-1 | C23H3.4 | 0.11107885 | 29 | 2.66E-02 |

| | | | | | |
|---|---|---|---|---|---|
| WBGene00016067 | clc-7 | C24H10.1 | 0.33428936 | 18 | 2.41E-04 |
| WBGene00016069 | C24H10.3 | C24H10.3 | 0.28349047 | 5 | 3.44E-02 |
| WBGene00016081 | C25A6.1 | C25A6.1 | 0.65957873 | 11 | 1.70E-05 |
| WBGene00016093 | srsx-34 | C25E10.3 | 0.22542595 | 22 | 1.48E-03 |
| WBGene00016094 | C25E10.4 | C25E10.4 | 0.2980538 | 11 | 4.93E-03 |
| WBGene00016095 | C25E10.5 | C25E10.5 | 0.31636322 | 7 | 1.37E-02 |
| WBGene00016103 | dpyd-1 | C25F6.3 | 0.45407814 | 18 | 1.28E-05 |
| WBGene00016104 | ddr-1 | C25F6.4 | 0.30280265 | 15 | 1.32E-03 |
| WBGene00016105 | C25F6.6 | C25F6.6 | 1.05648577 | 4 | 2.23E-04 |
| WBGene00016119 | C25H3.10 | C25H3.10 | 0.1924269 | 8 | 4.32E-02 |
| WBGene00016133 | C26B9.3 | C26B9.3 | 0.21268008 | 7 | 4.14E-02 |
| WBGene00016136 | C26B9.7 | C26B9.7 | 0.595023 | 2 | 2.08E-02 |
| WBGene00016148 | C26F1.3 | C26F1.3 | 0.25352415 | 10 | 1.27E-02 |
| WBGene00016156 | ari-1.3 | C27A12.6 | 0.24873618 | 5 | 4.57E-02 |
| WBGene00016181 | C28C12.11 | C28C12.11 | 0.3715566 | 6 | 1.13E-02 |
| WBGene00016187 | C28G1.2 | C28G1.2 | 0.52639375 | 6 | 2.64E-03 |
| WBGene00016188 | sec-15 | C28G1.3 | 0.18476059 | 16 | 1.30E-02 |
| WBGene00016195 | erd-2.2 | C28H8.4 | 0.28979154 | 7 | 1.82E-02 |
| WBGene00016196 | C28H8.5 | C28H8.5 | 0.31024938 | 5 | 2.77E-02 |
| WBGene00016201 | tdo-2 | C28H8.11 | 0.63869831 | 6 | 9.17E-04 |
| WBGene00016208 | C29E4.11 | C29E4.11 | 0.42492011 | 6 | 6.85E-03 |
| WBGene00016264 | C30F12.5 | C30F12.5 | 0.17939382 | 20 | 7.84E-03 |
| WBGene00016269 | C30G4.4 | C30G4.4 | 0.1112618 | 32 | 2.14E-02 |
| WBGene00016281 | pals-32 | C31B8.4 | 0.1933228 | 9 | 3.60E-02 |
| WBGene00016283 | zmp-3 | C31B8.8 | 0.25199197 | 13 | 6.20E-03 |
| WBGene00016292 | tbc-7 | C31H2.1 | 0.24211439 | 25 | 4.48E-04 |
| WBGene00016320 | C32E8.1 | C32E8.1 | 0.8290869 | 4 | 1.07E-03 |
| WBGene00016325 | C32E8.9 | C32E8.9 | 0.20509262 | 10 | 2.55E-02 |
| WBGene00016331 | obr-4 | C32F10.1 | 0.24145723 | 15 | 4.71E-03 |
| WBGene00016355 | lact-5 | C33F10.7 | 0.42427618 | 11 | 6.81E-04 |
| WBGene00016403 | grsp-3 | C34D4.11 | 0.54195045 | 4 | 7.73E-03 |
| WBGene00016408 | prmt-5 | C34E10.5 | 0.21676358 | 16 | 6.42E-03 |
| WBGene00016415 | ampd-1 | C34F11.3 | 0.34713252 | 21 | 5.87E-05 |
| WBGene00016417 | C34F11.8 | C34F11.8 | 0.42322548 | 4 | 1.75E-02 |
| WBGene00016419 | tyr-4 | C34G6.2 | 0.172419 | 18 | 1.27E-02 |
| WBGene00016421 | cdc-7 | C34G6.5 | 0.18068259 | 12 | 2.66E-02 |
| WBGene00016422 | noah-1 | C34G6.6 | 1.06803029 | 21 | 8.40E-14 |
| WBGene00016428 | dmsr-7 | C35A11.1 | 0.2754563 | 6 | 2.79E-02 |
| WBGene00016446 | C35D10.10 | C35D10.10 | 0.4072062 | 6 | 8.09E-03 |
| WBGene00016461 | C35E7.9 | C35E7.9 | 0.59505606 | 6 | 1.38E-03 |
| WBGene00016493 | C37A2.7 | C37A2.7 | 0.47895321 | 3 | 2.03E-02 |
| WBGene00016495 | C37C3.1 | C37C3.1 | 0.18440494 | 13 | 2.12E-02 |
| WBGene00016496 | C37C3.2 | C37C3.2 | 0.61524388 | 7 | 5.66E-04 |
| WBGene00016499 | txt-5 | C37C3.7 | 0.27315975 | 6 | 2.85E-02 |
| WBGene00016500 | memo-1 | C37C3.8 | 0.19175402 | 16 | 1.11E-02 |
| WBGene00016505 | ttr-33 | C37C3.13 | 0.6871353 | 5 | 1.29E-03 |
| WBGene00016507 | abhd-5.2 | C37H5.3 | 0.18196545 | 15 | 1.62E-02 |
| WBGene00016508 | C37H5.5 | C37H5.5 | 0.2435019 | 27 | 2.67E-04 |

| | | | | | |
|---|---|---|---|---|---|
| WBGene00016509 | adss-1 | C37H5.6 | 0.19423932 | 22 | 3.71E-03 |
| WBGene00016524 | C39B5.6 | C39B5.6 | 0.14873414 | 16 | 2.86E-02 |
| WBGene00016529 | C39D10.1 | C39D10.1 | 0.30714253 | 4 | 3.90E-02 |
| WBGene00016533 | C39D10.6 | C39D10.6 | 0.33850819 | 10 | 3.73E-03 |
| WBGene00016535 | C39D10.8 | C39D10.8 | 0.58787712 | 2 | 2.14E-02 |
| WBGene00016559 | C41A3.2 | C41A3.2 | 0.2942368 | 20 | 3.53E-04 |
| WBGene00016576 | lido-9 | C41H7.6 | 0.3458057 | 8 | 6.95E-03 |
| WBGene00016596 | C42D4.3 | C42D4.3 | 0.22397973 | 6 | 4.53E-02 |
| WBGene00016611 | bicd-1 | C43G2.2 | 0.1866807 | 16 | 1.24E-02 |
| WBGene00016613 | best-9 | C43G2.4 | 0.20119457 | 14 | 1.30E-02 |
| WBGene00016630 | acer-1 | C44B7.10 | 0.6294413 | 10 | 5.60E-05 |
| WBGene00016644 | abhd-3.2 | C44C1.5 | 0.24247987 | 8 | 2.38E-02 |
| WBGene00016645 | C44C1.6 | C44C1.6 | 0.53953395 | 3 | 1.44E-02 |
| WBGene00016653 | ssb-1 | C44E4.4 | 0.32401528 | 6 | 1.77E-02 |
| WBGene00016655 | acbp-1 | C44E4.6 | 0.34439223 | 5 | 2.10E-02 |
| WBGene00016657 | C44E12.1 | C44E12.1 | 0.40443687 | 7 | 5.36E-03 |
| WBGene00016659 | C45B2.2 | C45B2.2 | 0.56868422 | 3 | 1.22E-02 |
| WBGene00016661 | C45B2.6 | C45B2.6 | 0.1469421 | 15 | 3.34E-02 |
| WBGene00016665 | chil-11 | C45E5.2 | 0.59880376 | 6 | 1.34E-03 |
| WBGene00016671 | C45G7.4 | C45G7.4 | 0.44721753 | 13 | 1.78E-04 |
| WBGene00016728 | C46H11.2 | C46H11.2 | 0.36257372 | 7 | 8.37E-03 |
| WBGene00016739 | pitr-1 | C48A7.2 | 0.36247665 | 17 | 1.78E-04 |
| WBGene00016740 | C48B6.2 | C48B6.2 | 0.62484576 | 10 | 5.99E-05 |
| WBGene00016741 | C48B6.3 | C48B6.3 | 0.61467774 | 6 | 1.15E-03 |
| WBGene00016749 | C48E7.1 | C48E7.1 | 0.28389428 | 11 | 6.15E-03 |
| WBGene00016750 | let-611 | C48E7.2 | 0.49760721 | 12 | 1.24E-04 |
| WBGene00016767 | C49C8.3 | C49C8.3 | 0.52239413 | 5 | 4.92E-03 |
| WBGene00016788 | C49G7.10 | C49G7.10 | 0.44748315 | 7 | 3.39E-03 |
| WBGene00016790 | C49H3.3 | C49H3.3 | 0.54921355 | 3 | 1.37E-02 |
| WBGene00016794 | C49H3.9 | C49H3.9 | 0.1785186 | 12 | 2.75E-02 |
| WBGene00016860 | cyp-33C9 | C50H11.15 | 0.20983787 | 8 | 3.51E-02 |
| WBGene00016889 | lst-6 | C52E12.4 | 0.19842559 | 30 | 7.82E-04 |
| WBGene00016894 | C53B7.3 | C53B7.3 | 0.25442619 | 7 | 2.65E-02 |
| WBGene00016907 | C53H9.2 | C53H9.2 | 0.43446197 | 9 | 1.50E-03 |
| WBGene00016914 | C54D1.7 | C54D1.7 | 0.270822 | 7 | 2.23E-02 |
| WBGene00016916 | C54D2.2 | C54D2.2 | 0.65335073 | 3 | 7.60E-03 |
| WBGene00016927 | nhr-172 | C54F6.9 | 0.34075129 | 6 | 1.51E-02 |
| WBGene00016956 | C55C3.6 | C55C3.6 | 0.40667228 | 7 | 5.23E-03 |
| WBGene00016976 | C56E10.3 | C56E10.3 | 0.34168619 | 21 | 6.85E-05 |
| WBGene00016981 | rpn-13 | C56G2.7 | 0.59488508 | 11 | 4.69E-05 |
| WBGene00016982 | C56G2.9 | C56G2.9 | 0.29078263 | 6 | 2.42E-02 |
| WBGene00016987 | CC8.2 | CC8.2 | 0.29629543 | 6 | 2.29E-02 |
| WBGene00016992 | zhit-1 | CD4.7 | 0.66223118 | 4 | 3.38E-03 |
| WBGene00016993 | CD4.8 | CD4.8 | 0.15061656 | 14 | 3.48E-02 |
| WBGene00016997 | cebp-1 | D1005.3 | 0.69713715 | 5 | 1.19E-03 |
| WBGene00017012 | acs-22 | D1009.1 | 0.68031779 | 24 | 4.56E-10 |
| WBGene00017022 | D1022.3 | D1022.3 | 0.2746845 | 4 | 4.87E-02 |
| WBGene00017023 | D1022.4 | D1022.4 | 0.24918545 | 5 | 4.56E-02 |

| | | | | | |
|---|---|---|---|---|---|
| WBGene00017039 | trk-1 | D1073.1 | 0.43502025 | 23 | 1.89E-06 |
| WBGene00017046 | utx-1 | D2021.1 | 0.13442288 | 21 | 2.39E-02 |
| WBGene00017082 | DC2.5 | DC2.5 | 0.3453026 | 13 | 1.13E-03 |
| WBGene00017088 | akir-1 | E01A2.6 | 0.34820289 | 6 | 1.41E-02 |
| WBGene00017108 | cyp-43A1 | E03E2.1 | 0.63825334 | 10 | 4.93E-05 |
| WBGene00017121 | cyc-2.1 | E04A4.7 | 0.26961137 | 5 | 3.86E-02 |
| WBGene00017122 | pcrg-1 | E04F6.2 | 0.39983096 | 8 | 3.65E-03 |
| WBGene00017123 | maoc-1 | E04F6.3 | 0.45695215 | 9 | 1.12E-03 |
| WBGene00017124 | E04F6.4 | E04F6.4 | 0.35486842 | 13 | 9.54E-04 |
| WBGene00017125 | acdh-12 | E04F6.5 | 0.46190504 | 11 | 3.77E-04 |
| WBGene00017126 | E04F6.6 | E04F6.6 | 0.21597471 | 8 | 3.26E-02 |
| WBGene00017160 | F01F1.3 | F01F1.3 | 0.25741416 | 12 | 7.24E-03 |
| WBGene00017166 | aldo-2 | F01F1.12 | 0.84511677 | 8 | 1.81E-05 |
| WBGene00017168 | F01F1.14 | F01F1.14 | 1.26421702 | 5 | 1.17E-05 |
| WBGene00017217 | F07F6.4 | F07F6.4 | 0.2574261 | 9 | 1.55E-02 |
| WBGene00017220 | F07F6.8 | F07F6.8 | 0.63672266 | 7 | 4.50E-04 |
| WBGene00017258 | F08F1.4 | F08F1.4 | 0.37174779 | 3 | 3.71E-02 |
| WBGene00017262 | F08F3.4 | F08F3.4 | 0.23153541 | 10 | 1.74E-02 |
| WBGene00017272 | F08F8.7 | F08F8.7 | 0.21953618 | 6 | 4.72E-02 |
| WBGene00017283 | F09E5.3 | F09E5.3 | 0.23940018 | 7 | 3.11E-02 |
| WBGene00017293 | bmy-1 | F09E5.17 | 0.57551374 | 8 | 4.50E-04 |
| WBGene00017301 | hach-1 | F09F7.4 | 0.45768083 | 11 | 4.03E-04 |
| WBGene00017302 | F09F7.5 | F09F7.5 | 0.22352592 | 25 | 8.31E-04 |
| WBGene00017326 | dmd-5 | F10C1.5 | 0.61655785 | 4 | 4.62E-03 |
| WBGene00017329 | ugt-39 | F10D2.2 | 0.19954054 | 8 | 3.97E-02 |
| WBGene00017340 | F10D7.3 | F10D7.3 | 0.31081812 | 6 | 2.00E-02 |
| WBGene00017385 | F11G11.5 | F11G11.5 | 0.28010716 | 8 | 1.52E-02 |
| WBGene00017390 | F12A10.1 | F12A10.1 | 0.6484191 | 2 | 1.65E-02 |
| WBGene00017396 | suex-1 | F12A10.7 | 0.8713721 | 7 | 3.69E-05 |
| WBGene00017401 | F12D9.2 | F12D9.2 | 0.77709144 | 4 | 1.53E-03 |
| WBGene00017428 | F13D11.3 | F13D11.3 | 0.4559397 | 4 | 1.40E-02 |
| WBGene00017430 | bcl-11 | F13H6.1 | 0.27551494 | 25 | 1.47E-04 |
| WBGene00017438 | F13H8.5 | F13H8.5 | 0.44731968 | 11 | 4.74E-04 |
| WBGene00017463 | F14D12.1 | F14D12.1 | 0.45644642 | 18 | 1.21E-05 |
| WBGene00017471 | F14H12.3 | F14H12.3 | 0.29425104 | 8 | 1.28E-02 |
| WBGene00017483 | lgc-22 | F15E6.2 | 0.50023026 | 10 | 3.61E-04 |
| WBGene00017506 | F16B4.4 | F16B4.4 | 1.00611321 | 4 | 3.16E-04 |
| WBGene00017524 | F16G10.9 | F16G10.9 | 0.38583182 | 3 | 3.43E-02 |
| WBGene00017536 | F17A9.4 | F17A9.4 | 0.32303257 | 5 | 2.50E-02 |
| WBGene00017559 | mpz-3 | F18C5.4 | 0.65916801 | 9 | 7.79E-05 |
| WBGene00017560 | F18C5.5 | F18C5.5 | 1.05327928 | 5 | 6.52E-05 |
| WBGene00017565 | ddo-2 | F18E3.7 | 0.22200905 | 7 | 3.74E-02 |
| WBGene00017568 | F18E9.1 | F18E9.1 | 0.3234337 | 7 | 1.27E-02 |
| WBGene00017569 | F18E9.3 | F18E9.3 | 0.21634818 | 10 | 2.17E-02 |
| WBGene00017570 | F18E9.4 | F18E9.4 | 0.27016783 | 7 | 2.24E-02 |
| WBGene00017580 | lgc-4 | F18G5.4 | 0.2962975 | 16 | 1.12E-03 |
| WBGene00017582 | F18G5.6 | F18G5.6 | 0.26083978 | 8 | 1.91E-02 |
| WBGene00017629 | F20B6.5 | F20B6.5 | 0.34189756 | 13 | 1.21E-03 |

| | | | | | |
|---|---|---|---|---|---|
| WBGene00017638 | F20D6.9 | F20D6.9 | 0.3448512 | 3 | 4.32E-02 |
| WBGene00017650 | F21A9.1 | F21A9.1 | 0.31248394 | 13 | 2.06E-03 |
| WBGene00017654 | F21C10.4 | F21C10.4 | 0.34759922 | 4 | 2.95E-02 |
| WBGene00017655 | F21C10.5 | F21C10.5 | 0.94852967 | 2 | 4.43E-03 |
| WBGene00017693 | flp-23 | F22B7.2 | 1.03514298 | 6 | 2.21E-05 |
| WBGene00017694 | F22B7.3 | F22B7.3 | 0.28912078 | 8 | 1.37E-02 |
| WBGene00017698 | F22B7.9 | F22B7.9 | 2.0368982 | 10 | 8.50E-14 |
| WBGene00017700 | F22D3.4 | F22D3.4 | 0.22248061 | 14 | 8.60E-03 |
| WBGene00017707 | F22E5.8 | F22E5.8 | 0.28002226 | 4 | 4.70E-02 |
| WBGene00017714 | F22F1.2 | F22F1.2 | 0.91030848 | 9 | 2.85E-06 |
| WBGene00017722 | F22F7.4 | F22F7.4 | 0.26256758 | 7 | 2.43E-02 |
| WBGene00017735 | did-2 | F23C8.6 | 0.49956658 | 5 | 5.93E-03 |
| WBGene00017743 | F23F1.2 | F23F1.2 | 0.39545057 | 6 | 9.03E-03 |
| WBGene00017757 | bra-2 | F23H11.1 | 0.35849173 | 8 | 5.98E-03 |
| WBGene00017765 | gcst-1 | F25B4.1 | 0.24415821 | 11 | 1.15E-02 |
| WBGene00017770 | F25B4.7 | F25B4.7 | 0.19999113 | 7 | 4.73E-02 |
| WBGene00017771 | F25B4.8 | F25B4.8 | 0.36926933 | 3 | 3.76E-02 |
| WBGene00017772 | clec-1 | F25B4.9 | 1.09655318 | 4 | 1.69E-04 |
| WBGene00017781 | F25E2.3 | F25E2.3 | 0.20568456 | 7 | 4.46E-02 |
| WBGene00017789 | F25E5.8 | F25E5.8 | 0.35070513 | 5 | 1.99E-02 |
| WBGene00017794 | mltn-13 | F25F6.1 | 1.00945502 | 16 | 1.76E-10 |
| WBGene00017795 | F25F8.1 | F25F8.1 | 0.14303574 | 25 | 1.21E-02 |
| WBGene00017834 | F26F12.3 | F26F12.3 | 0.10699783 | 29 | 3.11E-02 |
| WBGene00017838 | F26G1.2 | F26G1.2 | 0.88925874 | 5 | 2.48E-04 |
| WBGene00017840 | ttm-2 | F26G1.4 | 0.16959737 | 11 | 3.70E-02 |
| WBGene00017841 | F26G1.5 | F26G1.5 | 0.94378125 | 3 | 1.48E-03 |
| WBGene00017864 | pcca-1 | F27D9.5 | 0.27391212 | 10 | 9.46E-03 |
| WBGene00017885 | dip-2 | F28B3.4 | 0.13442995 | 54 | 9.21E-04 |
| WBGene00017904 | lim-8 | F28F5.3 | 0.3716604 | 40 | 1.56E-08 |
| WBGene00017922 | F29B9.7 | F29B9.7 | 0.40511145 | 4 | 1.98E-02 |
| WBGene00017923 | F29B9.8 | F29B9.8 | 1.1163677 | 14 | 2.41E-10 |
| WBGene00017954 | F31E8.4 | F31E8.4 | 0.49417742 | 4 | 1.07E-02 |
| WBGene00017968 | skpo-3 | F32A5.2 | 0.14044812 | 19 | 2.50E-02 |
| WBGene00017971 | F32A5.8 | F32A5.8 | 0.57208906 | 4 | 6.28E-03 |
| WBGene00017979 | F32B5.6 | F32B5.6 | 0.3485055 | 13 | 1.07E-03 |
| WBGene00017981 | figl-1 | F32D1.1 | 0.15945987 | 13 | 3.34E-02 |
| WBGene00017982 | hpo-18 | F32D1.2 | 0.47629967 | 4 | 1.21E-02 |
| WBGene00017984 | gmpr-1 | F32D1.5 | 0.2265675 | 7 | 3.57E-02 |
| WBGene00017991 | clec-180 | F32E10.3 | 0.38777987 | 19 | 4.29E-05 |
| WBGene00017998 | F33D4.6 | F33D4.6 | 0.1451852 | 13 | 4.33E-02 |
| WBGene00018003 | F33D11.6 | F33D11.6 | 0.18369886 | 8 | 4.80E-02 |
| WBGene00018005 | nlp-62 | F33D11.8 | 0.31797086 | 6 | 1.87E-02 |
| WBGene00018006 | gpaa-1 | F33D11.9 | 0.21396882 | 15 | 8.33E-03 |
| WBGene00018009 | dhhc-3 | F33D11.12 | 0.27655545 | 11 | 6.90E-03 |
| WBGene00018014 | wdr-83 | F33G12.2 | 0.48187552 | 4 | 1.17E-02 |
| WBGene00018015 | F33G12.3 | F33G12.3 | 1.10612943 | 5 | 4.24E-05 |
| WBGene00018031 | F35B3.4 | F35B3.4 | 0.30938374 | 5 | 2.79E-02 |
| WBGene00018039 | F35D2.1 | F35D2.1 | 0.13680993 | 15 | 4.12E-02 |

| | | | | | |
|---|---|---|---|---|---|
| WBGene00018070 | F35H10.5 | F35H10.5 | 1.04809982 | 4 | 2.37E-04 |
| WBGene00018073 | F35H10.10 | F35H10.10 | 0.24990028 | 34 | 3.85E-05 |
| WBGene00018111 | F36H9.2 | F36H9.2 | 0.2908969 | 18 | 6.98E-04 |
| WBGene00018131 | F37A4.1 | F37A4.1 | 0.2933559 | 10 | 7.15E-03 |
| WBGene00018144 | F37C4.4 | F37C4.4 | 0.40954822 | 11 | 8.57E-04 |
| WBGene00018145 | F37C4.5 | F37C4.5 | 0.53073535 | 14 | 2.14E-05 |
| WBGene00018204 | F39F10.4 | F39F10.4 | 0.27925455 | 5 | 3.57E-02 |
| WBGene00018206 | ugt-61 | F39G3.1 | 0.26800549 | 8 | 1.76E-02 |
| WBGene00018226 | F40B5.2 | F40B5.2 | 0.47216569 | 8 | 1.54E-03 |
| WBGene00018232 | F40E3.5 | F40E3.5 | 0.18256693 | 33 | 9.09E-04 |
| WBGene00018267 | F41C3.1 | F41C3.1 | 0.61811106 | 5 | 2.26E-03 |
| WBGene00018268 | F41C3.2 | F41C3.2 | 0.96542486 | 12 | 4.47E-08 |
| WBGene00018281 | tbc-18 | F41D9.1 | 0.26062081 | 16 | 2.45E-03 |
| WBGene00018282 | F41D9.2 | F41D9.2 | 1.07495262 | 10 | 9.05E-08 |
| WBGene00018283 | sulp-3 | F41D9.5 | 0.3264048 | 13 | 1.60E-03 |
| WBGene00018293 | F41E6.12 | F41E6.12 | 0.27447589 | 6 | 2.82E-02 |
| WBGene00018294 | atg-18 | F41E6.13 | 0.30534488 | 15 | 1.25E-03 |
| WBGene00018297 | F41F3.3 | F41F3.3 | 0.67849401 | 2 | 1.44E-02 |
| WBGene00018302 | F41G3.6 | F41G3.6 | 0.47366137 | 9 | 8.96E-04 |
| WBGene00018321 | sand-1 | F41H10.11 | 0.14951137 | 13 | 4.00E-02 |
| WBGene00018330 | elks-1 | F42A6.9 | 0.20502793 | 29 | 7.27E-04 |
| WBGene00018339 | abcf-3 | F42A10.1 | 0.17075464 | 12 | 3.14E-02 |
| WBGene00018349 | F42C5.9 | F42C5.9 | 0.28360762 | 8 | 1.46E-02 |
| WBGene00018373 | F43B10.1 | F43B10.1 | 0.14799606 | 71 | 5.00E-05 |
| WBGene00018395 | mtch-1 | F43E2.7 | 0.27669858 | 5 | 3.64E-02 |
| WBGene00018400 | F43H9.4 | F43H9.4 | 0.41255349 | 5 | 1.20E-02 |
| WBGene00018402 | kvs-3 | F44A2.2 | 0.19526597 | 8 | 4.18E-02 |
| WBGene00018403 | F44A2.3 | F44A2.3 | 0.25448157 | 14 | 4.62E-03 |
| WBGene00018405 | F44A2.5 | F44A2.5 | 0.11750006 | 19 | 4.50E-02 |
| WBGene00018418 | F44E2.4 | F44E2.4 | 0.2200055 | 22 | 1.73E-03 |
| WBGene00018425 | F44E7.3 | F44E7.3 | 0.30084051 | 4 | 4.07E-02 |
| WBGene00018430 | nhr-142 | F44E7.8 | 0.18426365 | 20 | 6.88E-03 |
| WBGene00018472 | nep-16 | F45E4.7 | 0.18586662 | 28 | 1.78E-03 |
| WBGene00018482 | F45F2.10 | F45F2.10 | 0.16498039 | 23 | 7.67E-03 |
| WBGene00018486 | glb-16 | F46C8.7 | 0.356007 | 8 | 6.16E-03 |
| WBGene00018487 | F46C8.8 | F46C8.8 | 0.53006576 | 12 | 7.14E-05 |
| WBGene00018488 | acs-1 | F46E10.1 | 0.14384134 | 26 | 1.05E-02 |
| WBGene00018514 | twnk-1 | F46G11.1 | 0.14373889 | 25 | 1.18E-02 |
| WBGene00018515 | F46G11.2 | F46G11.2 | 0.25050579 | 14 | 4.99E-03 |
| WBGene00018518 | F46H5.2 | F46H5.2 | 0.67446559 | 13 | 2.84E-06 |
| WBGene00018532 | F47B7.1 | F47B7.1 | 0.66077996 | 5 | 1.59E-03 |
| WBGene00018565 | F47E1.1 | F47E1.1 | 0.58459311 | 5 | 2.97E-03 |
| WBGene00018566 | F47E1.2 | F47E1.2 | 0.36821108 | 14 | 5.05E-04 |
| WBGene00018571 | F47F2.3 | F47F2.3 | 0.35732757 | 6 | 1.29E-02 |
| WBGene00018576 | F47G3.1 | F47G3.1 | 0.23226187 | 15 | 5.70E-03 |
| WBGene00018586 | ubxn-3 | F48A11.5 | 0.36577862 | 17 | 1.65E-04 |
| WBGene00018587 | F48B9.1 | F48B9.1 | 0.3665237 | 12 | 1.14E-03 |
| WBGene00018607 | F48E3.8 | F48E3.8 | 0.08602907 | 44 | 3.93E-02 |

| | | | | | |
|---|---|---|---|---|---|
| WBGene00018638 | F49E8.7 | F49E8.7 | 0.23810653 | 13 | 7.98E-03 |
| WBGene00018657 | acl-4 | F49H12.6 | 0.18664092 | 17 | 1.06E-02 |
| WBGene00018675 | F52C9.5 | F52C9.5 | 0.36530844 | 16 | 2.46E-04 |
| WBGene00018700 | lmd-3 | F52E1.13 | 0.11044803 | 77 | 1.01E-03 |
| WBGene00018702 | F52E4.5 | F52E4.5 | 0.33801003 | 4 | 3.15E-02 |
| WBGene00018703 | sec-3 | F52E4.7 | 0.15991917 | 11 | 4.30E-02 |
| WBGene00018710 | F52G3.1 | F52G3.1 | 0.18690781 | 15 | 1.46E-02 |
| WBGene00018719 | ccd-5 | F52H2.7 | 0.23070783 | 29 | 2.72E-04 |
| WBGene00018738 | swip-10 | F53B1.6 | 0.29829545 | 18 | 5.82E-04 |
| WBGene00018739 | F53B1.8 | F53B1.8 | 0.1394302 | 27 | 1.11E-02 |
| WBGene00018740 | tra-4 | F53B3.1 | 0.15091185 | 13 | 3.90E-02 |
| WBGene00018744 | F53B3.6 | F53B3.6 | 0.39136151 | 13 | 4.91E-04 |
| WBGene00018745 | F53C3.1 | F53C3.1 | 0.27029086 | 5 | 3.84E-02 |
| WBGene00018748 | F53C3.4 | F53C3.4 | 0.27094769 | 7 | 2.22E-02 |
| WBGene00018750 | F53C3.6 | F53C3.6 | 0.20964172 | 11 | 1.97E-02 |
| WBGene00018764 | F53F10.2 | F53F10.2 | 0.29615845 | 15 | 1.52E-03 |
| WBGene00018786 | hmbx-1 | F54A5.1 | 0.28200687 | 13 | 3.59E-03 |
| WBGene00018789 | F54C1.1 | F54C1.1 | 0.27735181 | 17 | 1.28E-03 |
| WBGene00018802 | F54D8.6 | F54D8.6 | 0.16497869 | 23 | 7.67E-03 |
| WBGene00018804 | lido-15 | F54D10.3 | 0.33279062 | 8 | 8.12E-03 |
| WBGene00018811 | pmt-2 | F54D11.1 | 0.69405992 | 21 | 3.25E-09 |
| WBGene00018842 | F54H12.2 | F54H12.2 | 0.20919137 | 8 | 3.54E-02 |
| WBGene00018844 | F54H12.4 | F54H12.4 | 0.23261001 | 6 | 4.18E-02 |
| WBGene00018846 | eef-1B.1 | F54H12.6 | 0.4675012 | 11 | 3.46E-04 |
| WBGene00018847 | marc-6 | F55A3.1 | 0.17151801 | 18 | 1.30E-02 |
| WBGene00018869 | rfip-1 | F55C12.1 | 0.18550609 | 25 | 2.94E-03 |
| WBGene00018949 | acbp-4 | F56C9.5 | 0.52961151 | 6 | 2.56E-03 |
| WBGene00018953 | F56C9.10 | F56C9.10 | 0.10785979 | 22 | 4.75E-02 |
| WBGene00018969 | F56D3.1 | F56D3.1 | 0.35406582 | 9 | 4.33E-03 |
| WBGene00018974 | fcho-1 | F56D12.6 | 0.1493321 | 25 | 9.81E-03 |
| WBGene00018985 | F56F10.2 | F56F10.2 | 0.19161764 | 10 | 3.10E-02 |
| WBGene00018987 | lgl-1 | F56F10.4 | 0.15177716 | 19 | 1.86E-02 |
| WBGene00018990 | klf-1 | F56F11.3 | 0.44806503 | 13 | 1.75E-04 |
| WBGene00018996 | F57B9.1 | F57B9.1 | 0.43228465 | 9 | 1.55E-03 |
| WBGene00019007 | ucr-11 | F57B10.14 | 0.35334089 | 4 | 2.83E-02 |
| WBGene00019011 | F57C9.4 | F57C9.4 | 0.13264989 | 17 | 3.70E-02 |
| WBGene00019047 | F58E2.3 | F58E2.3 | 0.21629 | 8 | 3.25E-02 |
| WBGene00019070 | F58H7.5 | F58H7.5 | 0.25940357 | 9 | 1.51E-02 |
| WBGene00019074 | cpd-1 | F59A3.1 | 0.1022876 | 32 | 3.12E-02 |
| WBGene00019081 | F59A3.8 | F59A3.8 | 0.15802237 | 15 | 2.65E-02 |
| WBGene00019102 | F59C12.3 | F59C12.3 | 0.57638005 | 16 | 2.38E-06 |
| WBGene00019109 | F59E11.2 | F59E11.2 | 0.20767112 | 9 | 2.98E-02 |
| WBGene00019111 | F59E11.5 | F59E11.5 | 0.4257089 | 9 | 1.69E-03 |
| WBGene00019113 | F59E11.7 | F59E11.7 | 0.20058212 | 7 | 4.70E-02 |
| WBGene00019122 | F59E12.6 | F59E12.6 | 0.14252204 | 18 | 2.64E-02 |
| WBGene00019148 | H03E18.1 | H03E18.1 | 0.25959379 | 27 | 1.50E-04 |
| WBGene00019185 | H10E21.5 | H10E21.5 | 0.20846251 | 14 | 1.13E-02 |
| WBGene00019204 | H14N18.4 | H14N18.4 | 0.25420635 | 22 | 6.32E-04 |

| | | | | | |
|---|---|---|---|---|---|
| WBGene00019211 | H18N23.2 | H18N23.2 | 0.11086538 | 51 | 5.86E-03 |
| WBGene00019236 | H23N18.5 | H23N18.5 | 0.90566212 | 3 | 1.84E-03 |
| WBGene00019249 | hrpa-2 | H28G03.1 | 0.20085645 | 11 | 2.26E-02 |
| WBGene00019265 | H35N09.1 | H35N09.1 | 0.56947628 | 11 | 6.98E-05 |
| WBGene00019268 | H41C03.1 | H41C03.1 | 0.17484737 | 10 | 3.95E-02 |
| WBGene00019287 | K01A12.3 | K01A12.3 | 1.3487468 | 15 | 5.17E-13 |
| WBGene00019290 | K02A2.5 | K02A2.5 | 0.8350862 | 8 | 2.04E-05 |
| WBGene00019294 | K02A6.3 | K02A6.3 | 0.22942109 | 10 | 1.80E-02 |
| WBGene00019297 | K02B2.6 | K02B2.6 | 0.68652421 | 5 | 1.29E-03 |
| WBGene00019298 | pnp-1 | K02D7.1 | 0.13897477 | 19 | 2.59E-02 |
| WBGene00019300 | swt-1 | K02D7.5 | 0.2126734 | 13 | 1.27E-02 |
| WBGene00019304 | K02D10.4 | K02D10.4 | 0.60903938 | 6 | 1.21E-03 |
| WBGene00019317 | K02E10.1 | K02E10.1 | 0.25840967 | 14 | 4.28E-03 |
| WBGene00019322 | ahcy-1 | K02F2.2 | 1.4693582 | 9 | 1.81E-09 |
| WBGene00019326 | K02F3.2 | K02F3.2 | 0.1106902 | 25 | 3.55E-02 |
| WBGene00019327 | zip-2 | K02F3.4 | 0.97200971 | 11 | 1.26E-07 |
| WBGene00019331 | dos-3 | K02F3.8 | 0.25071884 | 12 | 8.11E-03 |
| WBGene00019365 | rei-2 | K03E6.7 | 0.17439036 | 10 | 3.98E-02 |
| WBGene00019380 | K04C2.2 | K04C2.2 | 0.52832135 | 15 | 1.24E-05 |
| WBGene00019382 | K04C2.5 | K04C2.5 | 1.11082569 | 4 | 1.54E-04 |
| WBGene00019408 | K05F1.6 | K05F1.6 | 0.14155562 | 18 | 2.70E-02 |
| WBGene00019457 | vms-1 | K06H7.3 | 0.19633673 | 7 | 4.92E-02 |
| WBGene00019464 | K07B1.4 | K07B1.4 | 0.24888854 | 8 | 2.21E-02 |
| WBGene00019465 | acl-14 | K07B1.5 | 0.15519966 | 15 | 2.81E-02 |
| WBGene00019466 | tos-1 | K07B1.6 | 0.52489261 | 19 | 1.26E-06 |
| WBGene00019487 | ephx-1 | K07D4.7 | 0.22751417 | 25 | 7.28E-04 |
| WBGene00019489 | K07E1.1 | K07E1.1 | 0.71905988 | 5 | 9.92E-04 |
| WBGene00019490 | K07E3.1 | K07E3.1 | 0.28350273 | 22 | 2.66E-04 |
| WBGene00019492 | K07E3.4 | K07E3.4 | 0.46115572 | 9 | 1.06E-03 |
| WBGene00019503 | tbce-1 | K07H8.1 | 0.3291643 | 18 | 2.73E-04 |
| WBGene00019507 | K07H8.7 | K07H8.7 | 0.24601332 | 6 | 3.68E-02 |
| WBGene00019509 | K07H8.9 | K07H8.9 | 0.65832133 | 10 | 3.69E-05 |
| WBGene00019510 | nucl-1 | K07H8.10 | 0.26993933 | 9 | 1.31E-02 |
| WBGene00019518 | K08B5.1 | K08B5.1 | 0.5573021 | 9 | 2.98E-04 |
| WBGene00019519 | K08B5.2 | K08B5.2 | 0.964181 | 8 | 4.39E-06 |
| WBGene00019537 | K08D12.3 | K08D12.3 | 0.30006116 | 5 | 3.01E-02 |
| WBGene00019540 | K08D12.6 | K08D12.6 | 1.16218426 | 11 | 6.40E-09 |
| WBGene00019571 | K09E2.2 | K09E2.2 | 0.30270174 | 20 | 2.81E-04 |
| WBGene00019572 | K09E2.3 | K09E2.3 | 0.41977946 | 12 | 4.62E-04 |
| WBGene00019578 | K09E3.7 | K09E3.7 | 0.13675612 | 14 | 4.56E-02 |
| WBGene00019630 | emb-1 | K10D2.4 | 0.37853418 | 4 | 2.38E-02 |
| WBGene00019633 | K10D2.7 | K10D2.7 | 0.59451431 | 3 | 1.06E-02 |
| WBGene00019648 | K11D12.8 | K11D12.8 | 0.41132312 | 9 | 2.04E-03 |
| WBGene00019656 | slc-25A10 | K11G12.5 | 0.46628406 | 10 | 5.90E-04 |
| WBGene00019658 | K11H12.1 | K11H12.1 | 0.44607262 | 4 | 1.50E-02 |
| WBGene00019669 | K12B6.4 | K12B6.4 | 0.32678135 | 3 | 4.78E-02 |
| WBGene00019693 | ostd-1 | M01A10.3 | 0.67242354 | 4 | 3.15E-03 |
| WBGene00019697 | M01B12.4 | M01B12.4 | 0.13760115 | 24 | 1.61E-02 |

| | | | | | |
|---|---|---|---|---|---|
| WBGene00019700 | btb-11 | M01D1.3 | 0.48790605 | 8 | 1.28E-03 |
| WBGene00019719 | M01H9.3 | M01H9.3 | 0.18618765 | 23 | 3.99E-03 |
| WBGene00019726 | bris-1 | M02B7.5 | 0.13472631 | 23 | 1.95E-02 |
| WBGene00019754 | M03E7.2 | M03E7.2 | 0.29983414 | 5 | 3.02E-02 |
| WBGene00019760 | calu-1 | M03F4.7 | 0.56733157 | 13 | 2.00E-05 |
| WBGene00019761 | M03F8.1 | M03F8.1 | 0.18905367 | 11 | 2.72E-02 |
| WBGene00019780 | M60.4 | M60.4 | 0.16218932 | 11 | 4.15E-02 |
| WBGene00019819 | aass-1 | R02D3.1 | 0.5063136 | 35 | 2.37E-10 |
| WBGene00019823 | fnta-1 | R02D3.5 | 0.2190525 | 11 | 1.70E-02 |
| WBGene00019825 | R02D3.8 | R02D3.8 | 0.51223365 | 4 | 9.49E-03 |
| WBGene00019827 | mop-25.1 | R02E12.2 | 0.26612648 | 25 | 2.01E-04 |
| WBGene00019831 | R02F2.1 | R02F2.1 | 0.18344088 | 18 | 9.69E-03 |
| WBGene00019832 | osg-1 | R02F2.2 | 0.09214494 | 47 | 2.42E-02 |
| WBGene00019841 | R02F11.3 | R02F11.3 | 0.35042498 | 17 | 2.35E-04 |
| WBGene00019843 | R03E9.2 | R03E9.2 | 0.17308883 | 9 | 4.70E-02 |
| WBGene00019846 | gpx-7 | R03G5.5 | 0.39463699 | 2 | 5.00E-02 |
| WBGene00019855 | R03H10.2 | R03H10.2 | 0.12471227 | 30 | 1.45E-02 |
| WBGene00019858 | R03H10.6 | R03H10.6 | 0.17563459 | 9 | 4.54E-02 |
| WBGene00019871 | R04E5.8 | R04E5.8 | 0.29306372 | 22 | 2.01E-04 |
| WBGene00019875 | mca-2 | R05C11.3 | 0.37531945 | 38 | 2.88E-08 |
| WBGene00019895 | aagr-2 | R05F9.12 | 0.16614857 | 41 | 6.11E-04 |
| WBGene00019900 | vdac-1 | R05G6.7 | 0.30440525 | 8 | 1.14E-02 |
| WBGene00019908 | R05H11.2 | R05H11.2 | 0.82493431 | 8 | 2.31E-05 |
| WBGene00019925 | fbxc-31 | R07C3.9 | 0.20465172 | 7 | 4.50E-02 |
| WBGene00019946 | chat-1 | R08C7.2 | 0.28755629 | 4 | 4.46E-02 |
| WBGene00019947 | htz-1 | R08C7.3 | 0.37737547 | 4 | 2.40E-02 |
| WBGene00019955 | R08C7.12 | R08C7.12 | 0.24912197 | 26 | 2.78E-04 |
| WBGene00019961 | R08E5.1 | R08E5.1 | 0.55727633 | 8 | 5.60E-04 |
| WBGene00019967 | cyp-33C8 | R08F11.3 | 0.31569719 | 11 | 3.74E-03 |
| WBGene00019979 | R09B5.11 | R09B5.11 | 0.67664025 | 9 | 6.19E-05 |
| WBGene00019980 | chil-14 | R09B5.12 | 0.28868797 | 5 | 3.30E-02 |
| WBGene00020007 | R11F4.1 | R11F4.1 | 0.53152651 | 15 | 1.16E-05 |
| WBGene00020031 | suco-1 | R12E2.2 | 0.26506124 | 12 | 6.36E-03 |
| WBGene00020036 | R12E2.11 | R12E2.11 | 0.29746009 | 5 | 3.07E-02 |
| WBGene00020039 | R12E2.14 | R12E2.14 | 0.65814255 | 3 | 7.40E-03 |
| WBGene00020052 | R13A5.10 | R13A5.10 | 0.25362729 | 7 | 2.67E-02 |
| WBGene00020092 | pcf-11 | R144.2 | 0.48046251 | 22 | 7.97E-07 |
| WBGene00020096 | R144.6 | R144.6 | 0.30266356 | 6 | 2.16E-02 |
| WBGene00020097 | larp-1 | R144.7 | 0.28354352 | 27 | 6.36E-05 |
| WBGene00020112 | pfd-5 | R151.9 | 1.45588596 | 3 | 8.28E-05 |
| WBGene00020116 | moa-1 | R155.2 | 0.11805472 | 22 | 3.51E-02 |
| WBGene00020131 | gcy-28 | T01A4.1 | 0.12304461 | 79 | 2.47E-04 |
| WBGene00020135 | aakg-4 | T01B6.3 | 0.2430974 | 17 | 2.84E-03 |
| WBGene00020136 | nlp-45 | T01B6.4 | 0.6009966 | 4 | 5.15E-03 |
| WBGene00020139 | eppl-1 | T01B11.2 | 0.24938433 | 10 | 1.35E-02 |
| WBGene00020160 | igcm-3 | T02C5.3 | 0.31219493 | 19 | 3.00E-04 |
| WBGene00020169 | mmaa-1 | T02G5.13 | 0.6058274 | 12 | 1.98E-05 |
| WBGene00020181 | T02H6.11 | T02H6.11 | 0.91464191 | 6 | 6.85E-05 |

| | | | | | |
|---|---|---|---|---|---|
| WBGene00020195 | T03G6.3 | T03G6.3 | 0.40567122 | 18 | 4.20E-05 |
| WBGene00020217 | T04G9.6 | T04G9.6 | 0.31545511 | 13 | 1.95E-03 |
| WBGene00020219 | T05A7.1 | T05A7.1 | 0.23986706 | 6 | 3.90E-02 |
| WBGene00020231 | T05A8.5 | T05A8.5 | 0.1560151 | 24 | 8.91E-03 |
| WBGene00020237 | phat-4 | T05B4.3 | 0.37775763 | 3 | 3.59E-02 |
| WBGene00020250 | T05C1.3 | T05C1.3 | 0.27892233 | 6 | 2.70E-02 |
| WBGene00020269 | erfa-1 | T05H4.6 | 0.42449764 | 8 | 2.72E-03 |
| WBGene00020274 | T05H4.11 | T05H4.11 | 0.53857494 | 4 | 7.91E-03 |
| WBGene00020275 | atp-4 | T05H4.12 | 0.59066631 | 5 | 2.82E-03 |
| WBGene00020276 | T05H4.15 | T05H4.15 | 0.20972514 | 10 | 2.39E-02 |
| WBGene00020284 | mel-46 | T06A10.1 | 0.10892073 | 32 | 2.36E-02 |
| WBGene00020300 | ssna-1 | T07A9.13 | 0.72456933 | 5 | 9.48E-04 |
| WBGene00020313 | T07E3.2 | T07E3.2 | 0.43404183 | 9 | 1.51E-03 |
| WBGene00020315 | T07E3.4 | T07E3.4 | 0.59003956 | 10 | 9.89E-05 |
| WBGene00020335 | ttr-30 | T08A9.2 | 0.26588903 | 6 | 3.05E-02 |
| WBGene00020339 | ttr-59 | T08A9.11 | 0.73120191 | 3 | 4.90E-03 |
| WBGene00020346 | rbm-5 | T08B2.5 | 0.31648675 | 21 | 1.40E-04 |
| WBGene00020347 | ech-1.2 | T08B2.7 | 0.34369942 | 16 | 3.95E-04 |
| WBGene00020348 | mrpl-23 | T08B2.8 | 0.73907072 | 5 | 8.43E-04 |
| WBGene00020366 | acdh-10 | T08G2.3 | 0.42424724 | 9 | 1.72E-03 |
| WBGene00020391 | cct-7 | T10B5.5 | 0.24996834 | 16 | 3.10E-03 |
| WBGene00020411 | T10E9.1 | T10E9.1 | 0.23099083 | 9 | 2.19E-02 |
| WBGene00020425 | syx-18 | T10H9.3 | 0.56250221 | 8 | 5.26E-04 |
| WBGene00020446 | T12B3.3 | T12B3.3 | 0.31236851 | 7 | 1.43E-02 |
| WBGene00020460 | nhr-273 | T12C9.1 | 0.22703066 | 7 | 3.55E-02 |
| WBGene00020478 | T13C2.2 | T13C2.2 | 0.17401717 | 21 | 7.82E-03 |
| WBGene00020479 | T13C2.3 | T13C2.3 | 0.18292498 | 9 | 4.13E-02 |
| WBGene00020483 | T13C5.2 | T13C5.2 | 0.47935562 | 6 | 4.10E-03 |
| WBGene00020484 | T13C5.3 | T13C5.3 | 0.32202647 | 9 | 6.61E-03 |
| WBGene00020486 | T13C5.6 | T13C5.6 | 0.36685842 | 6 | 1.18E-02 |
| WBGene00020497 | T14A8.2 | T14A8.2 | 0.57907179 | 5 | 3.10E-03 |
| WBGene00020507 | vha-15 | T14F9.1 | 0.44403204 | 9 | 1.32E-03 |
| WBGene00020530 | T15B12.1 | T15B12.1 | 0.3273454 | 25 | 2.63E-05 |
| WBGene00020546 | zig-13 | T17A3.10 | 0.22440916 | 12 | 1.27E-02 |
| WBGene00020550 | T17H7.1 | T17H7.1 | 0.86974695 | 9 | 4.86E-06 |
| WBGene00020555 | nhr-219 | T19A5.5 | 0.48704222 | 4 | 1.13E-02 |
| WBGene00020556 | pamn-1 | T19B4.1 | 0.26910914 | 14 | 3.47E-03 |
| WBGene00020566 | ttr-7 | T19C3.9 | 0.3144036 | 4 | 3.71E-02 |
| WBGene00020587 | ugt-9 | T19H12.1 | 0.20542431 | 14 | 1.20E-02 |
| WBGene00020604 | T20B12.7 | T20B12.7 | 0.38251374 | 5 | 1.54E-02 |
| WBGene00020625 | mrrf-1 | T20F5.3 | 0.41628232 | 4 | 1.84E-02 |
| WBGene00020626 | T20F5.4 | T20F5.4 | 0.35478169 | 9 | 4.29E-03 |
| WBGene00020629 | T20F5.7 | T20F5.7 | 0.36544524 | 7 | 8.12E-03 |
| WBGene00020636 | T20H4.5 | T20H4.5 | 0.73639017 | 4 | 2.03E-03 |
| WBGene00020647 | pqbp-1.1 | T21D12.3 | 0.37711646 | 9 | 3.20E-03 |
| WBGene00020658 | argn-1 | T21F4.1 | 0.7832438 | 7 | 9.45E-05 |
| WBGene00020660 | T21G5.2 | T21G5.2 | 0.49810103 | 2 | 3.18E-02 |
| WBGene00020670 | T22B2.6 | T22B2.6 | 0.39370046 | 4 | 2.15E-02 |

| | | | | | |
|---|---|---|---|---|---|
| WBGene00020673 | T22B7.4 | T22B7.4 | 0.23523412 | 9 | 2.07E-02 |
| WBGene00020693 | lron-8 | T22E7.1 | 0.14783451 | 17 | 2.60E-02 |
| WBGene00020696 | pygl-1 | T22F3.3 | 0.43968184 | 44 | 6.55E-11 |
| WBGene00020716 | phf-30 | T23B12.1 | 0.31317996 | 4 | 3.74E-02 |
| WBGene00020717 | mrpl-4 | T23B12.2 | 0.58486065 | 4 | 5.75E-03 |
| WBGene00020718 | mrps-2 | T23B12.3 | 0.29621362 | 6 | 2.30E-02 |
| WBGene00020719 | natc-1 | T23B12.4 | 0.16745947 | 19 | 1.25E-02 |
| WBGene00020720 | T23B12.5 | T23B12.5 | 0.37618173 | 5 | 1.62E-02 |
| WBGene00020781 | T24H7.2 | T24H7.2 | 0.52839321 | 9 | 4.36E-04 |
| WBGene00020797 | T25D3.3 | T25D3.3 | 0.21554152 | 14 | 9.85E-03 |
| WBGene00020822 | T26A5.6 | T26A5.6 | 0.29744225 | 11 | 4.98E-03 |
| WBGene00020824 | T26A8.1 | T26A8.1 | 0.37005823 | 15 | 3.28E-04 |
| WBGene00020825 | tmem-231 | T26A8.2 | 0.39201205 | 4 | 2.17E-02 |
| WBGene00020827 | T26A8.4 | T26A8.4 | 0.27746632 | 22 | 3.18E-04 |
| WBGene00020830 | T26C11.4 | T26C11.4 | 0.15997052 | 16 | 2.24E-02 |
| WBGene00020831 | T26C12.1 | T26C12.1 | 0.35734968 | 17 | 2.00E-04 |
| WBGene00020846 | T27A10.6 | T27A10.6 | 0.14763075 | 20 | 1.85E-02 |
| WBGene00020850 | nhr-226 | T27B7.3 | 0.18140534 | 14 | 1.91E-02 |
| WBGene00020851 | nhr-227 | T27B7.5 | 0.16273481 | 10 | 4.71E-02 |
| WBGene00020865 | T27E4.7 | T27E4.7 | 0.41708003 | 4 | 1.83E-02 |
| WBGene00020884 | T28B4.1 | T28B4.1 | 0.21608393 | 26 | 8.70E-04 |
| WBGene00020891 | cest-24 | T28C12.4 | 0.14644009 | 31 | 5.46E-03 |
| WBGene00020893 | T28C12.6 | T28C12.6 | 0.9330177 | 10 | 7.02E-07 |
| WBGene00020901 | T28F2.2 | T28F2.2 | 0.48836275 | 3 | 1.93E-02 |
| WBGene00020908 | VC5.2 | VC5.2 | 0.15020614 | 15 | 3.12E-02 |
| WBGene00020912 | W01A11.7 | W01A11.7 | 0.56257787 | 8 | 5.25E-04 |
| WBGene00020914 | sulp-6 | W01B11.2 | 0.182425 | 22 | 5.26E-03 |
| WBGene00020915 | nol-58 | W01B11.3 | 0.26498947 | 13 | 4.89E-03 |
| WBGene00020917 | W01B11.6 | W01B11.6 | 0.61578399 | 4 | 4.65E-03 |
| WBGene00020919 | set-19 | W01C8.3 | 0.16287017 | 19 | 1.40E-02 |
| WBGene00020923 | W02B3.3 | W02B3.3 | 0.55853902 | 3 | 1.30E-02 |
| WBGene00020930 | hlh-30 | W02C12.3 | 0.26793711 | 77 | 1.82E-10 |
| WBGene00020931 | cytb-5.2 | W02D3.1 | 0.88356233 | 3 | 2.08E-03 |
| WBGene00020937 | W02D3.12 | W02D3.12 | 0.27420644 | 4 | 4.89E-02 |
| WBGene00020947 | W02F12.2 | W02F12.2 | 0.25863857 | 6 | 3.27E-02 |
| WBGene00020948 | era-1 | W02F12.3 | 0.4225224 | 11 | 7.00E-04 |
| WBGene00020949 | W02F12.4 | W02F12.4 | 0.20856768 | 7 | 4.32E-02 |
| WBGene00020950 | dlst-1 | W02F12.5 | 0.32528315 | 9 | 6.33E-03 |
| WBGene00020951 | sna-1 | W02F12.6 | 0.70453565 | 6 | 4.94E-04 |
| WBGene00020952 | kel-8 | W02G9.2 | 0.10531085 | 50 | 8.99E-03 |
| WBGene00020957 | W02H5.2 | W02H5.2 | 0.15288264 | 19 | 1.81E-02 |
| WBGene00020984 | W03D8.1 | W03D8.1 | 0.56220491 | 8 | 5.28E-04 |
| WBGene00021043 | pck-1 | W05G11.6 | 0.6980869 | 25 | 1.15E-10 |
| WBGene00021047 | zfp-3 | W05H7.4 | 0.18907193 | 18 | 8.44E-03 |
| WBGene00021049 | W05H9.2 | W05H9.2 | 0.20273469 | 18 | 6.04E-03 |
| WBGene00021068 | W06H8.6 | W06H8.6 | 0.53099242 | 18 | 1.95E-06 |
| WBGene00021073 | nsun-1 | W07E6.1 | 0.26780375 | 13 | 4.65E-03 |
| WBGene00021076 | W07E6.5 | W07E6.5 | 0.29391052 | 12 | 3.90E-03 |

| | | | | | |
|---|---|---|---|---|---|
| WBGene00021079 | W08A12.2 | W08A12.2 | 0.31515913 | 5 | 2.66E-02 |
| WBGene00021080 | W08A12.3 | W08A12.3 | 0.52041133 | 18 | 2.53E-06 |
| WBGene00021087 | W08E12.6 | W08E12.6 | 0.27759466 | 9 | 1.19E-02 |
| WBGene00021088 | W08E12.7 | W08E12.7 | 0.29675788 | 12 | 3.72E-03 |
| WBGene00021095 | mlt-8 | W08F4.6 | 0.19994636 | 14 | 1.33E-02 |
| WBGene00021117 | clec-126 | W09G10.5 | 1.73970114 | 6 | 2.93E-08 |
| WBGene00021118 | clec-125 | W09G10.6 | 0.56959172 | 24 | 1.58E-08 |
| WBGene00021128 | W10C8.5 | W10C8.5 | 0.42771214 | 11 | 6.45E-04 |
| WBGene00021134 | W10G11.1 | W10G11.1 | 0.2871616 | 4 | 4.47E-02 |
| WBGene00021135 | W10G11.2 | W10G11.2 | 0.74571728 | 4 | 1.90E-03 |
| WBGene00021136 | W10G11.3 | W10G11.3 | 1.28558476 | 3 | 2.16E-04 |
| WBGene00021137 | W10G11.4 | W10G11.4 | 1.05453032 | 2 | 2.78E-03 |
| WBGene00021138 | clec-127 | W10G11.5 | 1.05639278 | 3 | 7.86E-04 |
| WBGene00021143 | clec-132 | W10G11.13 | 0.22352576 | 6 | 4.55E-02 |
| WBGene00021146 | lgc-45 | W10G11.16 | 0.74010095 | 40 | 7.13E-17 |
| WBGene00021149 | dnc-3 | W10G11.20 | 0.81393769 | 8 | 2.63E-05 |
| WBGene00021156 | Y4C6B.2 | Y4C6B.2 | 0.69420526 | 14 | 8.90E-07 |
| WBGene00021157 | Y4C6B.3 | Y4C6B.3 | 0.59786981 | 9 | 1.75E-04 |
| WBGene00021170 | arx-4 | Y6D11A.2 | 0.14161114 | 13 | 4.62E-02 |
| WBGene00021178 | Y9C9A.8 | Y9C9A.8 | 0.36907769 | 4 | 2.54E-02 |
| WBGene00021213 | Y18H1A.9 | Y18H1A.9 | 0.20882793 | 45 | 2.90E-05 |
| WBGene00021248 | Y22D7AL.10 | Y22D7AL.10 | 0.58305467 | 5 | 3.00E-03 |
| WBGene00021281 | ell-1 | Y24D9A.1 | 0.08404293 | 60 | 2.50E-02 |
| WBGene00021285 | Y24D9A.7 | Y24D9A.7 | 0.46164607 | 2 | 3.73E-02 |
| WBGene00021292 | copb-1 | Y25C1A.5 | 0.1344772 | 22 | 2.16E-02 |
| WBGene00021343 | cutl-23 | Y37B11A.1 | 0.38432005 | 20 | 3.10E-05 |
| WBGene00021348 | moag-4 | Y37E3.4 | 0.5517624 | 5 | 3.87E-03 |
| WBGene00021349 | arl-13 | Y37E3.5 | 0.1807385 | 13 | 2.27E-02 |
| WBGene00021350 | Y37E3.8 | Y37E3.8 | 1.00784557 | 4 | 3.12E-04 |
| WBGene00021352 | pcyt-2.1 | Y37E3.11 | 0.13945029 | 22 | 1.87E-02 |
| WBGene00021369 | siah-1 | Y37E11AR.2 | 0.14854079 | 17 | 2.56E-02 |
| WBGene00021371 | nape-1 | Y37E11AR.4 | 0.20281063 | 13 | 1.52E-02 |
| WBGene00021389 | Y38A10A.2 | Y38A10A.2 | 0.29291595 | 18 | 6.64E-04 |
| WBGene00021415 | nsy-4 | Y38F2AL.1 | 0.23417165 | 13 | 8.58E-03 |
| WBGene00021420 | trap-3 | Y38F2AR.2 | 1.31368642 | 7 | 3.31E-07 |
| WBGene00021427 | sec-61.B | Y38F2AR.9 | 0.75349066 | 2 | 1.04E-02 |
| WBGene00021430 | Y38F2AR.12 | Y38F2AR.12 | 0.24902338 | 47 | 1.71E-06 |
| WBGene00021460 | zwl-1 | Y39G10AR.2 | 0.10049856 | 26 | 4.71E-02 |
| WBGene00021469 | Y39G10AR.11 | Y39G10AR.11 | 0.28719212 | 25 | 9.99E-05 |
| WBGene00021486 | lbp-9 | Y40B10A.1 | 0.47948526 | 3 | 2.02E-02 |
| WBGene00021487 | comt-3 | Y40B10A.2 | 0.1448693 | 12 | 4.87E-02 |
| WBGene00021500 | Y40C7B.4 | Y40C7B.4 | 0.46922365 | 5 | 7.59E-03 |
| WBGene00021502 | Y40D12A.1 | Y40D12A.1 | 0.12952961 | 26 | 1.73E-02 |
| WBGene00021522 | nhr-274 | Y41D4B.21 | 0.14575281 | 14 | 3.83E-02 |
| WBGene00021533 | Y42G9A.3 | Y42G9A.3 | 0.54078639 | 8 | 6.81E-04 |
| WBGene00021535 | wht-7 | Y42G9A.6 | 0.29433667 | 26 | 5.84E-05 |
| WBGene00021540 | Y42H9AR.5 | Y42H9AR.5 | 0.28373249 | 5 | 3.44E-02 |
| WBGene00021544 | Y43B11AR.3 | Y43B11AR.3 | 0.44044421 | 7 | 3.65E-03 |

| | | | | | |
|---|---|---|---|---|---|
| WBGene00021556 | Y45G5AM.3 | Y45G5AM.3 | 0.21262085 | 8 | 3.40E-02 |
| WBGene00021561 | Y45G5AM.9 | Y45G5AM.9 | 0.16282996 | 50 | 2.18E-04 |
| WBGene00021562 | nuo-5 | Y45G12B.1 | 0.10485527 | 28 | 3.58E-02 |
| WBGene00021597 | spsb-1 | Y46E12BL.4 | 0.17768845 | 80 | 8.44E-07 |
| WBGene00021633 | Y47G6A.3 | Y47G6A.3 | 0.25287621 | 8 | 2.10E-02 |
| WBGene00021645 | Y47G6A.19 | Y47G6A.19 | 0.23143085 | 21 | 1.54E-03 |
| WBGene00021646 | Y47G6A.21 | Y47G6A.21 | 0.19487235 | 46 | 5.51E-05 |
| WBGene00021690 | Y48G8AL.12 | Y48G8AL.12 | 0.41178372 | 5 | 1.21E-02 |
| WBGene00021781 | Y51H7C.3 | Y51H7C.3 | 0.47630068 | 6 | 4.22E-03 |
| WBGene00021785 | Y51H7C.7 | Y51H7C.7 | 0.27812506 | 5 | 3.60E-02 |
| WBGene00021816 | Y53G8AR.9 | Y53G8AR.9 | 0.15919653 | 14 | 2.95E-02 |
| WBGene00021835 | Y54E10A.17 | Y54E10A.17 | 0.37058637 | 14 | 4.82E-04 |
| WBGene00021845 | rpb-7 | Y54E10BR.6 | 0.18637469 | 9 | 3.94E-02 |
| WBGene00021863 | Y54F10BM.9 | Y54F10BM.9 | 0.10754783 | 26 | 3.69E-02 |
| WBGene00021871 | dml-1 | Y54G2A.5 | 0.51480662 | 9 | 5.21E-04 |
| WBGene00021883 | Y54G2A.18 | Y54G2A.18 | 0.70825675 | 5 | 1.08E-03 |
| WBGene00021890 | Y54G2A.26 | Y54G2A.26 | 0.11766658 | 30 | 1.91E-02 |
| WBGene00021908 | Y55B1AR.4 | Y55B1AR.4 | 0.25660864 | 5 | 4.29E-02 |
| WBGene00021909 | Y55B1BL.1 | Y55B1BL.1 | 0.33702895 | 10 | 3.81E-03 |
| WBGene00021929 | dcap-1 | Y55F3AM.12 | 0.28095858 | 7 | 2.00E-02 |
| WBGene00021960 | tmem-258 | Y57E12AM.1 | 0.89144008 | 5 | 2.44E-04 |
| WBGene00021975 | Y58A7A.1 | Y58A7A.1 | 0.47045112 | 8 | 1.57E-03 |
| WBGene00022010 | catp-7 | Y59H11AR.2 | 0.13765021 | 35 | 5.18E-03 |
| WBGene00022012 | Y59H11AR.4 | Y59H11AR.4 | 0.28553578 | 7 | 1.90E-02 |
| WBGene00022033 | Y65B4BL.1 | Y65B4BL.1 | 0.94615291 | 9 | 1.78E-06 |
| WBGene00022042 | icd-2 | Y65B4BR.5 | 0.60685562 | 5 | 2.47E-03 |
| WBGene00022059 | Y67D8A.2 | Y67D8A.2 | 0.16924042 | 72 | 6.26E-06 |
| WBGene00022089 | Y69A2AR.18 | Y69A2AR.18 | 0.52876358 | 5 | 4.67E-03 |
| WBGene00022100 | Y69A2AR.31 | Y69A2AR.31 | 0.27829019 | 39 | 2.69E-06 |
| WBGene00022103 | cdh-12 | Y71D11A.1 | 0.12257319 | 81 | 2.19E-04 |
| WBGene00022114 | Y71F9AL.9 | Y71F9AL.9 | 0.82106947 | 9 | 9.23E-06 |
| WBGene00022119 | copa-1 | Y71F9AL.17 | 0.11790989 | 22 | 3.53E-02 |
| WBGene00022122 | trap-1 | Y71F9AM.6 | 0.59303117 | 6 | 1.41E-03 |
| WBGene00022125 | Y71F9B.1 | Y71F9B.1 | 0.47934851 | 6 | 4.10E-03 |
| WBGene00022127 | yop-1 | Y71F9B.3 | 0.25572983 | 9 | 1.58E-02 |
| WBGene00022129 | lron-11 | Y71F9B.8 | 0.29201411 | 34 | 5.89E-06 |
| WBGene00022139 | tub-2 | Y71G12A.3 | 0.42422812 | 42 | 4.00E-10 |
| WBGene00022158 | Y71G12B.23 | Y71G12B.23 | 0.32707253 | 8 | 8.69E-03 |
| WBGene00022160 | Y71G12B.25 | Y71G12B.25 | 0.24717668 | 11 | 1.09E-02 |
| WBGene00022231 | tyr-6 | Y73B6BL.1 | 0.52168816 | 20 | 7.61E-07 |
| WBGene00022233 | ipla-6 | Y73B6BL.4 | 0.30954488 | 12 | 2.99E-03 |
| WBGene00022282 | Y74C10AR.2 | Y74C10AR.2 | 0.45416987 | 4 | 1.41E-02 |
| WBGene00022307 | nadk-1 | Y77E11A.2 | 0.12210955 | 18 | 4.35E-02 |
| WBGene00022315 | Y81B9A.1 | Y81B9A.1 | 0.19086259 | 13 | 1.89E-02 |
| WBGene00022336 | Y82E9BR.3 | Y82E9BR.3 | 1.06573907 | 6 | 1.66E-05 |
| WBGene00022358 | Y92H12A.2 | Y92H12A.2 | 0.16854135 | 45 | 3.05E-04 |
| WBGene00022368 | Y92H12BR.2 | Y92H12BR.2 | 0.40005755 | 5 | 1.33E-02 |
| WBGene00022386 | Y95B8A.6 | Y95B8A.6 | 0.56487558 | 26 | 5.11E-09 |

| | | | | | |
|---|---|---|---|---|---|
| WBGene00022390 | him-19 | Y95B8A.11 | 0.18784118 | 15 | 1.43E-02 |
| WBGene00022478 | fbxa-35 | Y119D3A.3 | 0.24951275 | 6 | 3.56E-02 |
| WBGene00022479 | fbxa-36 | Y119D3A.4 | 0.33396976 | 5 | 2.28E-02 |
| WBGene00022517 | ZC123.1 | ZC123.1 | 0.3519016 | 22 | 3.54E-05 |
| WBGene00022532 | ZC155.4 | ZC155.4 | 0.24854246 | 8 | 2.21E-02 |
| WBGene00022533 | cutl-19 | ZC155.5 | 0.23864953 | 9 | 1.98E-02 |
| WBGene00022538 | ZC190.4 | ZC190.4 | 0.58990237 | 20 | 1.21E-07 |
| WBGene00022539 | ZC190.5 | ZC190.5 | 0.34931242 | 4 | 2.91E-02 |
| WBGene00022554 | duxl-1 | ZC204.2 | 0.13543802 | 16 | 3.83E-02 |
| WBGene00022580 | iglr-2 | ZC262.3 | 0.17756268 | 11 | 3.26E-02 |
| WBGene00022583 | mrps-18A | ZC262.8 | 0.59731499 | 3 | 1.04E-02 |
| WBGene00022596 | ZC395.4 | ZC395.4 | 0.5760547 | 3 | 1.17E-02 |
| WBGene00022598 | ztf-8 | ZC395.8 | 0.19566547 | 21 | 4.24E-03 |
| WBGene00022599 | daf-41 | ZC395.10 | 0.45376598 | 5 | 8.61E-03 |
| WBGene00022613 | sek-3 | ZC449.3 | 0.37751399 | 12 | 9.46E-04 |
| WBGene00022629 | algn-12 | ZC513.5 | 0.20037407 | 12 | 1.90E-02 |
| WBGene00022668 | ZK154.6 | ZK154.6 | 0.2934096 | 11 | 5.30E-03 |
| WBGene00022669 | ZK177.1 | ZK177.1 | 0.17973299 | 10 | 3.68E-02 |
| WBGene00022678 | sar-1 | ZK180.4 | 0.43707885 | 9 | 1.45E-03 |
| WBGene00022679 | ZK180.5 | ZK180.5 | 0.7817689 | 7 | 9.60E-05 |
| WBGene00022683 | ZK185.3 | ZK185.3 | 0.24799022 | 5 | 4.60E-02 |
| WBGene00022721 | ugtp-1 | ZK370.7 | 0.15658355 | 11 | 4.53E-02 |
| WBGene00022722 | tomm-70 | ZK370.8 | 0.19818244 | 14 | 1.38E-02 |
| WBGene00022743 | mlt-7 | ZK430.8 | 0.4812267 | 30 | 1.08E-08 |
| WBGene00022749 | ZK484.3 | ZK484.3 | 0.63719509 | 5 | 1.93E-03 |
| WBGene00022750 | tres-1 | ZK484.4 | 0.11261927 | 30 | 2.33E-02 |
| WBGene00022761 | ZK546.4 | ZK546.4 | 0.83809567 | 3 | 2.69E-03 |
| WBGene00022766 | cblc-1 | ZK546.17 | 0.2877896 | 12 | 4.33E-03 |
| WBGene00022767 | ZK563.2 | ZK563.2 | 0.16178429 | 15 | 2.45E-02 |
| WBGene00022780 | ZK622.1 | ZK622.1 | 0.80592438 | 10 | 4.39E-06 |
| WBGene00022781 | pmt-1 | ZK622.3 | 0.90013753 | 17 | 6.71E-10 |
| WBGene00022782 | ZK622.4 | ZK622.4 | 1.36562881 | 6 | 9.87E-07 |
| WBGene00022788 | slc-17.7 | ZK682.2 | 0.36618753 | 7 | 8.05E-03 |
| WBGene00022792 | ZK686.2 | ZK686.2 | 0.25914134 | 12 | 7.03E-03 |
| WBGene00022795 | ZK686.5 | ZK686.5 | 0.41485998 | 5 | 1.18E-02 |
| WBGene00022797 | best-24 | ZK688.2 | 0.43080566 | 11 | 6.14E-04 |
| WBGene00022800 | ZK688.5 | ZK688.5 | 0.49658343 | 27 | 3.11E-08 |
| WBGene00022801 | pcp-5 | ZK688.6 | 0.3547129 | 17 | 2.13E-04 |
| WBGene00022816 | fbn-1 | ZK783.1 | 0.38582325 | 82 | 2.05E-16 |
| WBGene00022817 | upp-1 | ZK783.2 | 1.33513979 | 10 | 2.12E-09 |
| WBGene00022826 | ZK816.3 | ZK816.3 | 0.98117276 | 1 | 1.23E-02 |
| WBGene00022827 | ZK816.4 | ZK816.4 | 0.69915715 | 7 | 2.32E-04 |
| WBGene00022850 | ZK1127.3 | ZK1127.3 | 0.43821283 | 6 | 6.04E-03 |
| WBGene00022855 | tcer-1 | ZK1127.9 | 0.15627505 | 13 | 3.54E-02 |
| WBGene00022856 | cth-2 | ZK1127.10 | 0.46843894 | 6 | 4.55E-03 |
| WBGene00022861 | dve-1 | ZK1193.5 | 0.2697366 | 21 | 5.23E-04 |
| WBGene00022884 | ZK1248.17 | ZK1248.17 | 0.77605003 | 2 | 9.42E-03 |
| WBGene00022892 | ZK1290.11 | ZK1290.11 | 0.31398876 | 4 | 3.72E-02 |

| | | | | | |
|---|---|---|---|---|---|
| WBGene00023068 | rpl-41.2 | F54D7.7 | 0.53765287 | 3 | 1.46E-02 |
| WBGene00023415 | F16C3.3 | F16C3.3 | 0.52163569 | 12 | 8.23E-05 |
| WBGene00023417 | swm-1 | C25E10.9 | 0.50927947 | 2 | 3.03E-02 |
| WBGene00023418 | R06F6.12 | R06F6.12 | 0.14385068 | 13 | 4.44E-02 |
| WBGene00023500 | C11H1.9 | C11H1.9 | 0.30998078 | 4 | 3.82E-02 |
| WBGene00044009 | ZK822.6 | ZK822.6 | 0.44277455 | 6 | 5.79E-03 |
| WBGene00044019 | C55A6.12 | C55A6.12 | 0.28166719 | 11 | 6.37E-03 |
| WBGene00044031 | F49C5.9 | F49C5.9 | 0.34317966 | 4 | 3.04E-02 |
| WBGene00044061 | tbc-12 | R11B5.1 | 0.49651626 | 18 | 4.55E-06 |
| WBGene00044107 | F58G6.9 | F58G6.9 | 0.31178957 | 4 | 3.77E-02 |
| WBGene00044109 | K02E11.10 | K02E11.10 | 1.18149304 | 5 | 2.29E-05 |
| WBGene00044133 | C44C10.12 | C44C10.12 | 0.58048204 | 4 | 5.93E-03 |
| WBGene00044180 | C47E8.11 | C47E8.11 | 0.78660147 | 3 | 3.59E-03 |
| WBGene00044191 | F43D2.6 | F43D2.6 | 0.19457189 | 11 | 2.50E-02 |
| WBGene00044194 | F47B8.14 | F47B8.14 | 0.79311598 | 5 | 5.43E-04 |
| WBGene00044243 | F16C3.4 | F16C3.4 | 0.41212313 | 5 | 1.21E-02 |
| WBGene00044258 | Y57G11C.51 | Y57G11C.51 | 0.59282216 | 5 | 2.77E-03 |
| WBGene00044263 | Y105C5A.26 | Y105C5A.26 | 0.42865091 | 2 | 4.30E-02 |
| WBGene00044264 | Y105E8A.32 | Y105E8A.32 | 0.28772891 | 6 | 2.49E-02 |
| WBGene00044294 | C01B10.11 | C01B10.11 | 0.69685268 | 8 | 1.06E-04 |
| WBGene00044326 | tag-322 | C33H5.10 | 0.28118946 | 5 | 3.51E-02 |
| WBGene00044344 | mrpl-39 | Y46H3A.7 | 0.25117408 | 9 | 1.68E-02 |
| WBGene00044381 | K10G6.5 | K10G6.5 | 0.40504015 | 5 | 1.28E-02 |
| WBGene00044391 | C25H3.15 | C25H3.15 | 0.41063314 | 3 | 2.98E-02 |
| WBGene00044393 | ZK1248.20 | ZK1248.20 | 0.54585398 | 4 | 7.52E-03 |
| WBGene00044464 | Y77E11A.16 | Y77E11A.16 | 0.17391042 | 10 | 4.01E-02 |
| WBGene00044471 | F56D6.9 | F56D6.9 | 0.28538632 | 4 | 4.53E-02 |
| WBGene00044510 | K07H8.11 | K07H8.11 | 0.25442937 | 11 | 9.77E-03 |
| WBGene00044534 | H23N18.6 | H23N18.6 | 0.18357908 | 8 | 4.80E-02 |
| WBGene00044542 | CD4.11 | CD4.11 | 0.48458686 | 20 | 2.07E-06 |
| WBGene00044569 | C25F6.8 | C25F6.8 | 0.4199174 | 4 | 1.79E-02 |
| WBGene00044570 | T22B7.8 | T22B7.8 | 0.22307884 | 17 | 4.53E-03 |
| WBGene00044577 | F43E12.1 | F43E12.1 | 0.39465672 | 2 | 5.00E-02 |
| WBGene00044611 | T27A3.8 | T27A3.8 | 0.34774183 | 3 | 4.25E-02 |
| WBGene00044612 | Y57E12AR.1 | Y57E12AR.1 | 0.22807803 | 6 | 4.36E-02 |
| WBGene00044621 | bus-5 | F53B1.4 | 0.59587435 | 9 | 1.79E-04 |
| WBGene00044630 | bus-17 | ZK678.8 | 0.13029373 | 21 | 2.69E-02 |
| WBGene00044634 | C29E4.14 | C29E4.14 | 0.31350411 | 5 | 2.70E-02 |
| WBGene00044636 | B0393.9 | B0393.9 | 0.38165431 | 7 | 6.83E-03 |
| WBGene00044667 | ZK622.5 | ZK622.5 | 0.1981335 | 8 | 4.04E-02 |
| WBGene00044675 | fbxa-76 | Y119D3B.22 | 0.48668357 | 6 | 3.83E-03 |
| WBGene00044707 | pals-15 | F22G12.7 | 0.30928599 | 4 | 3.84E-02 |
| WBGene00044732 | T03G11.10 | T03G11.10 | 0.37207082 | 10 | 2.30E-03 |
| WBGene00044767 | T23B7.3 | T23B7.3 | 0.41197025 | 11 | 8.26E-04 |
| WBGene00044789 | T07A9.14 | T07A9.14 | 0.4667373 | 4 | 1.30E-02 |
| WBGene00044799 | ZK686.6 | ZK686.6 | 0.31462521 | 8 | 1.01E-02 |
| WBGene00044916 | F40F8.11 | F40F8.11 | 0.23684287 | 8 | 2.55E-02 |
| WBGene00044917 | F40F8.12 | F40F8.12 | 0.28374463 | 4 | 4.58E-02 |

| | | | | | |
|---|---|---|---|---|---|
| WBGene00045013 | F08A7.1 | F08A7.1 | 0.42022146 | 8 | 2.86E-03 |
| WBGene00045061 | Y57G11B.8 | Y57G11B.8 | 0.94225242 | 3 | 1.49E-03 |
| WBGene00045063 | Y37A1A.4 | Y37A1A.4 | 0.0879214 | 57 | 2.09E-02 |
| WBGene00045183 | T28C6.10 | T28C6.10 | 0.42393121 | 5 | 1.10E-02 |
| WBGene00045245 | ttr-34 | F09A5.9 | 0.42211844 | 3 | 2.80E-02 |
| WBGene00045246 | C29E4.15 | C29E4.15 | 0.45251369 | 13 | 1.61E-04 |
| WBGene00045253 | F46C3.6 | F46C3.6 | 0.51663542 | 4 | 9.20E-03 |
| WBGene00045272 | F59C12.4 | F59C12.4 | 0.58609707 | 4 | 5.70E-03 |
| WBGene00045277 | ZK742.6 | ZK742.6 | 0.42926569 | 4 | 1.68E-02 |
| WBGene00045308 | C37H5.14 | C37H5.14 | 0.31514755 | 10 | 5.22E-03 |
| WBGene00045383 | sup-46 | C25A1.4 | 0.35235329 | 13 | 9.98E-04 |
| WBGene00045399 | Y47G6A.33 | Y47G6A.33 | 0.43808402 | 8 | 2.32E-03 |
| WBGene00045418 | F15B9.10 | F15B9.10 | 0.23581342 | 11 | 1.31E-02 |
| WBGene00045433 | F49D11.10 | F49D11.10 | 0.16774902 | 24 | 6.12E-03 |
| WBGene00045475 | T07A5.7 | T07A5.7 | 0.13913129 | 14 | 4.35E-02 |
| WBGene00045488 | F57B1.9 | F57B1.9 | 0.3916224 | 8 | 4.03E-03 |
| WBGene00050943 | ZC412.10 | ZC412.10 | 1.55905748 | 7 | 2.42E-08 |
| WBGene00050968 | R11G10.4 | R11G10.4 | 1.06159136 | 1 | 9.55E-03 |
| WBGene00077453 | Y62F5A.12 | Y62F5A.12 | 0.32840588 | 4 | 3.36E-02 |
| WBGene00077489 | C04G6.13 | C04G6.13 | 0.28465619 | 7 | 1.92E-02 |
| WBGene00077536 | F38B7.10 | F38B7.10 | 0.36700843 | 4 | 2.58E-02 |
| WBGene00077548 | F21C3.7 | F21C3.7 | 2.0456782 | 5 | 2.01E-08 |
| WBGene00077585 | T01G5.8 | T01G5.8 | 1.15338529 | 2 | 1.81E-03 |
| WBGene00077690 | mtp-18 | T13C5.8 | 0.45893591 | 5 | 8.25E-03 |
| WBGene00077693 | T04C12.11 | T04C12.11 | 0.34405843 | 3 | 4.34E-02 |
| WBGene00077696 | marc-1 | F58E6.12 | 0.31990621 | 4 | 3.57E-02 |
| WBGene00077704 | F12A10.9 | F12A10.9 | 0.37485557 | 4 | 2.44E-02 |
| WBGene00077714 | R102.11 | R102.11 | 0.42551388 | 5 | 1.08E-02 |
| WBGene00077786 | F53F8.7 | F53F8.7 | 0.18875485 | 9 | 3.82E-02 |
| WBGene00164968 | K08H2.10 | K08H2.10 | 0.74110148 | 12 | 2.00E-06 |
| WBGene00185014 | F17C8.9 | F17C8.9 | 0.6012323 | 6 | 1.30E-03 |
| WBGene00189933 | F38B7.11 | F38B7.11 | 0.57802579 | 5 | 3.13E-03 |
| WBGene00194704 | T07H3.8 | T07H3.8 | 0.38892364 | 4 | 2.22E-02 |
| WBGene00194708 | Y36E3A.2 | Y36E3A.2 | 0.38160025 | 3 | 3.51E-02 |
| WBGene00194733 | Y41C4A.21 | Y41C4A.21 | 0.19826784 | 7 | 4.82E-02 |
| WBGene00194734 | T04A6.4 | T04A6.4 | 0.50736386 | 3 | 1.73E-02 |
| WBGene00194742 | F37C12.21 | F37C12.21 | 0.68615496 | 3 | 6.32E-03 |
| WBGene00194835 | Y17G9B.11 | Y17G9B.11 | 0.54953454 | 3 | 1.36E-02 |
| WBGene00194867 | W09C2.10 | W09C2.10 | 0.86403933 | 2 | 6.41E-03 |
| WBGene00195146 | F54D5.17 | F54D5.17 | 0.4652566 | 3 | 2.19E-02 |
| WBGene00195208 | W02H3.3 | W02H3.3 | 0.40864182 | 5 | 1.24E-02 |
| WBGene00195212 | F09B12.7 | F09B12.7 | 0.21664641 | 7 | 3.96E-02 |
| WBGene00195246 | ZK546.19 | ZK546.19 | 0.56821897 | 3 | 1.23E-02 |
| WBGene00206373 | C27A2.12 | C27A2.12 | 0.79923385 | 6 | 2.03E-04 |
| WBGene00206479 | C44C11.6 | C44C11.6 | 0.41965567 | 4 | 1.79E-02 |
| WBGene00206484 | C31H2.14 | C31H2.14 | 0.90923733 | 2 | 5.26E-03 |
| WBGene00206487 | best-19 | T21D12.15 | 0.29006004 | 13 | 3.10E-03 |
| WBGene00206516 | F53E4.2 | F53E4.2 | 0.39553753 | 2 | 4.98E-02 |

| | | | | | |
|---|---|---|---|---|---|
| WBGene00206529 | C04A2.15 | C04A2.15 | 1.41609795 | 3 | 1.04E-04 |
| WBGene00219317 | F31E3.12 | F31E3.12 | 1.0902314 | 4 | 1.77E-04 |
| WBGene00219376 | C28G1.10 | C28G1.10 | 0.82621454 | 2 | 7.56E-03 |
| WBGene00220261 | ZC434.10 | ZC434.10 | 0.38262132 | 4 | 2.32E-02 |
| WBGene00235095 | B0035.21 | B0035.21 | 0.4834202 | 2 | 3.39E-02 |
| WBGene00235102 | dib-1 | Y54G2A.75 | 0.98615075 | 5 | 1.13E-04 |
| WBGene00235158 | Y37E3.30 | Y37E3.30 | 0.44209294 | 6 | 5.83E-03 |
| WBGene00235332 | K04A8.20 | K04A8.20 | 0.72846922 | 1 | 2.70E-02 |
| WBGene00269421 | R06F6.14 | R06F6.14 | 0.33172326 | 7 | 1.16E-02 |
| WBGene00271777 | Y48G8AR.8 | Y48G8AR.8 | 0.4943166 | 11 | 2.27E-04 |
| WBGene00271798 | F54D5.24 | F54D5.24 | 0.16543866 | 11 | 3.94E-02 |
| WBGene00271822 | F11G11.15 | F11G11.15 | 0.95829604 | 3 | 1.36E-03 |
| WBGene00302974 | Y65B4BM.3 | Y65B4BM.3 | 0.71148208 | 11 | 7.53E-06 |
| WBGene00302980 | F53E2.2 | F53E2.2 | 0.6247686 | 10 | 5.99E-05 |
| WBGene00302993 | T14B4.19 | T14B4.19 | 0.3377733 | 9 | 5.37E-03 |
| WBGene00303000 | C37C3.18 | C37C3.18 | 0.44091149 | 7 | 3.63E-03 |
| WBGene00303011 | F17C11.26 | F17C11.26 | 0.31184224 | 6 | 1.98E-02 |
| WBGene00303048 | F55C5.17 | F55C5.17 | 1.19348859 | 1 | 6.33E-03 |
| WBGene00303057 | F33D11.17 | F33D11.17 | 0.73030749 | 1 | 2.68E-02 |
| WBGene00303079 | T09F3.7 | T09F3.7 | 0.72320818 | 1 | 2.74E-02 |
| WBGene00303088 | Y44E3A.7 | Y44E3A.7 | 0.52296397 | 2 | 2.85E-02 |
| WBGene00303093 | Y55B1BL.2 | Y55B1BL.2 | 0.79808368 | 1 | 2.17E-02 |
| WBGene00303105 | R07E5.19 | R07E5.19 | 0.60937293 | 5 | 2.42E-03 |
| WBGene00303423 | F10D7.11 | F10D7.11 | 0.36186033 | 5 | 1.82E-02 |
| WBGene00304794 | H40L08.9 | H40L08.9 | 0.60878092 | 3 | 9.77E-03 |
| WBGene00304808 | F47E1.20 | F47E1.20 | 0.4300134 | 17 | 3.70E-05 |
| WBGene00304824 | Y7A5A.25 | Y7A5A.25 | 1.00879243 | 3 | 1.03E-03 |
| WBGene00304992 | F35C12.7 | F35C12.7 | 0.38978454 | 9 | 2.71E-03 |
| WBGene00305157 | T24H7.9 | T24H7.9 | 0.40599182 | 5 | 1.27E-02 |
| WBGene00305173 | H12C20.7 | H12C20.7 | 0.23283996 | 7 | 3.34E-02 |

**Supplementary Table S6 Unique and overlapping sets of protein coding genes expressed in hypodermis determined by RAPID, TaDa, PATseq and FACS**

| RAPID | TaDa | PATseq | FACS | RAPID\|TaDa | RAPID\|PATseq | RAPID\|FACS | TaDa\|PATseq | TaDa\|FACS | PATseq\|FACS | RAPID\|TaDa\|PATseq | RAPID\|TaDa\|FACS | RAPID\|PATseq\|FACS | TaDa\|PATseq\|FACS | RAPID\|TaDa\|PATseq\|FACS |
|---|---|---|---|---|---|---|---|---|---|---|---|---|---|---|
| WBGene00000006 | WBGene00000239 | WBGene00000034 | WBGene00000001 | WBGene00000057 | WBGene00000262 | WBGene00000017 | WBGene00000039 | WBGene00000073 | WBGene00000018 | WBGene00000399 | WBGene00000004 | WBGene00000074 | WBGene00000040 | WBGene00000041 |
| WBGene00000044 | WBGene00000453 | WBGene00000145 | WBGene00000010 | WBGene00000082 | WBGene00000285 | WBGene00000165 | WBGene00000189 | WBGene00000090 | WBGene00000036 | WBGene00000402 | WBGene00000225 | WBGene00000112 | WBGene00000064 | WBGene00000080 |
| WBGene00000075 | WBGene00000525 | WBGene00000180 | WBGene00000020 | WBGene00000093 | WBGene00000835 | WBGene00000238 | WBGene00000220 | WBGene00000106 | WBGene00000063 | WBGene00000594 | WBGene00000233 | WBGene00000122 | WBGene00000101 | WBGene00000083 |
| WBGene00000084 | WBGene00000623 | WBGene00000244 | WBGene00000022 | WBGene00000463 | WBGene00001429 | WBGene00000482 | WBGene00000624 | WBGene00000179 | WBGene00000065 | WBGene00000618 | WBGene00000246 | WBGene00000292 | WBGene00000105 | WBGene00000100 |
| WBGene00000295 | WBGene00000714 | WBGene00000280 | WBGene00000070 | WBGene00000664 | WBGene00001500 | WBGene00000671 | WBGene00000641 | WBGene00000187 | WBGene00000066 | WBGene00000625 | WBGene00000265 | WBGene00000379 | WBGene00000109 | WBGene00000107 |
| WBGene00000401 | WBGene00000724 | WBGene00000284 | WBGene00000072 | WBGene00000758 | WBGene00001560 | WBGene00000818 | WBGene00000650 | WBGene00000226 | WBGene00000067 | WBGene00000631 | WBGene00000420 | WBGene00000383 | WBGene00000110 | WBGene00000114 |
| WBGene00000412 | WBGene00000763 | WBGene00000296 | WBGene00000079 | WBGene00001020 | WBGene00003583 | WBGene00001116 | WBGene00000659 | WBGene00000236 | WBGene00000068 | WBGene00000634 | WBGene00000548 | WBGene00000509 | WBGene00000117 | WBGene00000115 |
| WBGene00000422 | WBGene00000878 | WBGene00000301 | WBGene00000086 | WBGene00001707 | WBGene00003615 | WBGene00001412 | WBGene00000663 | WBGene00000271 | WBGene00000081 | WBGene00000655 | WBGene00000995 | WBGene00000526 | WBGene00000123 | WBGene00000146 |
| WBGene00000512 | WBGene00000902 | WBGene00000393 | WBGene00000089 | WBGene00001788 | WBGene00003626 | WBGene00001623 | WBGene00000666 | WBGene00000459 | WBGene00000088 | WBGene00000665 | WBGene00001068 | WBGene00000556 | WBGene00000142 | WBGene00000149 |
| WBGene00000559 | WBGene00000905 | WBGene00000439 | WBGene00000099 | WBGene00001997 | WBGene00003674 | WBGene00001753 | WBGene00000689 | WBGene00000493 | WBGene00000092 | WBGene00000685 | WBGene00001233 | WBGene00000585 | WBGene00000151 | WBGene00000161 |
| WBGene00000563 | WBGene00000933 | WBGene00000444 | WBGene00000103 | WBGene00002088 | WBGene00003710 | WBGene00001760 | WBGene00000694 | WBGene00000537 | WBGene00000095 | WBGene00000691 | WBGene00001368 | WBGene00000601 | WBGene00000156 | WBGene00000175 |
| WBGene00000648 | WBGene00001057 | WBGene00000455 | WBGene00000136 | WBGene00002120 | WBGene00003763 | WBGene00002026 | WBGene00000695 | WBGene00000549 | WBGene00000097 | WBGene00000702 | WBGene00001478 | WBGene00000602 | WBGene00000163 | WBGene00000221 |
| WBGene00000654 | WBGene00001135 | WBGene00000469 | WBGene00000138 | WBGene00002181 | WBGene00004133 | WBGene00002087 | WBGene00000697 | WBGene00000687 | WBGene00000098 | WBGene00000743 | WBGene00001497 | WBGene00000726 | WBGene00000170 | WBGene00000223 |
| WBGene00000674 | WBGene00001284 | WBGene00000483 | WBGene00000147 | WBGene00002827 | WBGene00004827 | WBGene00002200 | WBGene00000704 | WBGene00000723 | WBGene00000102 | WBGene00000746 | WBGene00001515 | WBGene00000761 | WBGene00000176 | WBGene00000229 |
| WBGene00000884 | WBGene00001547 | WBGene00000511 | WBGene00000159 | WBGene00003562 | WBGene00004854 | WBGene00002240 | WBGene00000727 | WBGene00000981 | WBGene00000111 | WBGene00000753 | WBGene00001615 | WBGene00000787 | WBGene00000183 | WBGene00000240 |
| WBGene00000950 | WBGene00001700 | WBGene00000516 | WBGene00000164 | WBGene00003692 | WBGene00006044 | WBGene00002324 | WBGene00000730 | WBGene00001043 | WBGene00000113 | WBGene00000795 | WBGene00001674 | WBGene00000837 | WBGene00000197 | WBGene00000253 |
| WBGene00000951 | WBGene00001984 | WBGene00000560 | WBGene00000168 | WBGene00003878 | WBGene00006310 | WBGene00003037 | WBGene00000738 | WBGene00001058 | WBGene00000116 | WBGene00001063 | WBGene00001768 | WBGene00000880 | WBGene00000207 | WBGene00000254 |
| WBGene00001078 | WBGene00002227 | WBGene00000593 | WBGene00000188 | WBGene00004963 | WBGene00006431 | WBGene00003078 | WBGene00000778 | WBGene00001148 | WBGene00000118 | WBGene00001064 | WBGene00001774 | WBGene00000984 | WBGene00000231 | WBGene00000256 |
| WBGene00001210 | WBGene00002248 | WBGene00000596 | WBGene00000195 | WBGene00004985 | WBGene00006580 | WBGene00003081 | WBGene00000992 | WBGene00001215 | WBGene00000120 | WBGene00001065 | WBGene00001898 | WBGene00001005 | WBGene00000249 | WBGene00000273 |
| WBGene00001486 | WBGene00002252 | WBGene00000613 | WBGene00000202 | WBGene00005001 | WBGene00006976 | WBGene00003171 | WBGene00001071 | WBGene00001241 | WBGene00000121 | WBGene00001069 | WBGene00002100 | WBGene00001016 | WBGene00000251 | WBGene00000282 |
| WBGene00001531 | WBGene00003035 | WBGene00000614 | WBGene00000203 | WBGene00005004 | WBGene00007115 | WBGene00003384 | WBGene00001075 | WBGene00001538 | WBGene00000140 | WBGene00001555 | WBGene00002214 | WBGene00001018 | WBGene00000252 | WBGene00000293 |
| WBGene00001532 | WBGene00003545 | WBGene00000617 | WBGene00000211 | WBGene00005078 | WBGene00007202 | WBGene00003522 | WBGene00001076 | WBGene00001580 | WBGene00000150 | WBGene00001690 | WBGene00003089 | WBGene00001179 | WBGene00000276 | WBGene00000298 |
| WBGene00001537 | WBGene00003548 | WBGene00000620 | WBGene00000227 | WBGene00005727 | WBGene00007203 | WBGene00003589 | WBGene00001079 | WBGene00001620 | WBGene00000155 | WBGene00001730 | WBGene00003603 | WBGene00001229 | WBGene00000369 | WBGene00000366 |
| WBGene00001644 | WBGene00003552 | WBGene00000621 | WBGene00000235 | WBGene00006533 | WBGene00007214 | WBGene00003627 | WBGene00001480 | WBGene00001680 | WBGene00000157 | WBGene00003057 | WBGene00003616 | WBGene00001240 | WBGene00000371 | WBGene00000377 |
| WBGene00001712 | WBGene00003650 | WBGene00000628 | WBGene00000237 | WBGene00006674 | WBGene00007344 | WBGene00003654 | WBGene00001570 | WBGene00001775 | WBGene00000158 | WBGene00003186 | WBGene00003630 | WBGene00001325 | WBGene00000376 | WBGene00000381 |
| WBGene00001720 | WBGene00003706 | WBGene00000700 | WBGene00000250 | WBGene00006853 | WBGene00007363 | WBGene00003660 | WBGene00001691 | WBGene00001909 | WBGene00000160 | WBGene00003675 | WBGene00003709 | WBGene00001453 | WBGene00000380 | WBGene00000407 |
| WBGene00001733 | WBGene00003740 | WBGene00000701 | WBGene00000269 | WBGene00006954 | WBGene00007396 | WBGene00003837 | WBGene00001695 | WBGene00001915 | WBGene00000162 | WBGene00003731 | WBGene00004023 | WBGene00001514 | WBGene00000382 | WBGene00000421 |
| WBGene00001740 | WBGene00004059 | WBGene00000719 | WBGene00000281 | WBGene00007325 | WBGene00007436 | WBGene00003965 | WBGene00001719 | WBGene00001979 | WBGene00000169 | WBGene00003844 | WBGene00004038 | WBGene00001516 | WBGene00000425 | WBGene00000528 |
| WBGene00001756 | WBGene00004061 | WBGene00000720 | WBGene00000368 | WBGene00007401 | WBGene00007557 | WBGene00004312 | WBGene00001805 | WBGene00002001 | WBGene00000178 | WBGene00003862 | WBGene00004225 | WBGene00001682 | WBGene00000441 | WBGene00000542 |
| WBGene00001778 | WBGene00004127 | WBGene00000729 | WBGene00000373 | WBGene00007723 | WBGene00007562 | WBGene00004489 | WBGene00002039 | WBGene00002173 | WBGene00000181 | WBGene00004004 | WBGene00004226 | WBGene00001683 | WBGene00000443 | WBGene00000552 |
| WBGene00001781 | WBGene00004229 | WBGene00000734 | WBGene00000375 | WBGene00007778 | WBGene00007633 | WBGene00004490 | WBGene00002184 | WBGene00002583 | WBGene00000182 | WBGene00004216 | WBGene00004279 | WBGene00001784 | WBGene00000475 | WBGene00000599 |
| WBGene00001981 | WBGene00004904 | WBGene00000741 | WBGene00000386 | WBGene00007894 | WBGene00007675 | WBGene00004776 | WBGene00002282 | WBGene00003015 | WBGene00000184 | WBGene00004218 | WBGene00004310 | WBGene00001854 | WBGene00000479 | WBGene00000603 |
| WBGene00002051 | WBGene00005012 | WBGene00000749 | WBGene00000391 | WBGene00007997 | WBGene00008113 | WBGene00004856 | WBGene00003888 | WBGene00003024 | WBGene00000186 | WBGene00004219 | WBGene00004412 | WBGene00002048 | WBGene00000502 | WBGene00000606 |
| WBGene00002092 | WBGene00005196 | WBGene00000756 | WBGene00000408 | WBGene00008433 | WBGene00008222 | WBGene00004882 | WBGene00004185 | WBGene00003104 | WBGene00000192 | WBGene00004228 | WBGene00004414 | WBGene00002192 | WBGene00000508 | WBGene00000616 |
| WBGene00002218 | WBGene00005393 | WBGene00000766 | WBGene00000413 | WBGene00008511 | WBGene00008687 | WBGene00004969 | WBGene00004224 | WBGene00003107 | WBGene00000196 | WBGene00004232 | WBGene00004415 | WBGene00002215 | WBGene00000534 | WBGene00000626 |
| WBGene00002976 | WBGene00005416 | WBGene00000768 | WBGene00000418 | WBGene00008779 | WBGene00008712 | WBGene00006059 | WBGene00004281 | WBGene00003407 | WBGene00000198 | WBGene00004359 | WBGene00004416 | WBGene00002990 | WBGene00000547 | WBGene00000647 |
| WBGene00003018 | WBGene00005713 | WBGene00000798 | WBGene00000423 | WBGene00008858 | WBGene00008976 | WBGene00006418 | WBGene00004771 | WBGene00003429 | WBGene00000199 | WBGene00004395 | WBGene00004417 | WBGene00003053 | WBGene00000550 | WBGene00000649 |
| WBGene00003394 | WBGene00006016 | WBGene00000830 | WBGene00000438 | WBGene00009093 | WBGene00009025 | WBGene00006517 | WBGene00006473 | WBGene00003478 | WBGene00000200 | WBGene00004998 | WBGene00004418 | WBGene00003083 | WBGene00000608 | WBGene00000656 |
| WBGene00003634 | WBGene00006324 | WBGene00000851 | WBGene00000458 | WBGene00009447 | WBGene00009299 | WBGene00006570 | WBGene00006948 | WBGene00003480 | WBGene00000201 | WBGene00006732 | WBGene00004419 | WBGene00003084 | WBGene00000609 | WBGene00000657 |
| WBGene00003643 | WBGene00006509 | WBGene00000857 | WBGene00000498 | WBGene00009504 | WBGene00009355 | WBGene00006605 | WBGene00006952 | WBGene00003526 | WBGene00000204 | WBGene00006955 | WBGene00004420 | WBGene00003130 | WBGene00000615 | WBGene00000667 |
| WBGene00003694 | WBGene00006619 | WBGene00000888 | WBGene00000507 | WBGene00009652 | WBGene00009635 | WBGene00006646 | WBGene00007149 | WBGene00003580 | WBGene00000205 | WBGene00007299 | WBGene00004421 | WBGene00003148 | WBGene00000636 | WBGene00000668 |
| WBGene00003699 | WBGene00006865 | WBGene00000923 | WBGene00000519 | WBGene00009759 | WBGene00009645 | WBGene00006668 | WBGene00007170 | WBGene00003602 | WBGene00000206 | WBGene00007343 | WBGene00004422 | WBGene00003172 | WBGene00000639 | WBGene00000669 |
| WBGene00003701 | WBGene00007104 | WBGene00000930 | WBGene00000530 | WBGene00009867 | WBGene00009686 | WBGene00006736 | WBGene00007242 | WBGene00003609 | WBGene00000208 | WBGene00007479 | WBGene00004423 | WBGene00003253 | WBGene00000653 | WBGene00000672 |
| WBGene00003705 | WBGene00007146 | WBGene00000932 | WBGene00000532 | WBGene00010082 | WBGene00009816 | WBGene00006777 | WBGene00007434 | WBGene00003621 | WBGene00000209 | WBGene00007546 | WBGene00004424 | WBGene00003372 | WBGene00000678 | WBGene00000673 |
| WBGene00003729 | WBGene00007464 | WBGene00000965 | WBGene00000533 | WBGene00010155 | WBGene00009850 | WBGene00006887 | WBGene00007600 | WBGene00003637 | WBGene00000210 | WBGene00007560 | WBGene00004425 | WBGene00003517 | WBGene00000683 | WBGene00000677 |
| WBGene00003848 | WBGene00007583 | WBGene00000968 | WBGene00000553 | WBGene00010184 | WBGene00010290 | WBGene00006946 | WBGene00007823 | WBGene00003834 | WBGene00000214 | WBGene00007804 | WBGene00004426 | WBGene00003593 | WBGene00000693 | WBGene00000680 |
| WBGene00003883 | WBGene00007632 | WBGene00000973 | WBGene00000564 | WBGene00010501 | WBGene00010567 | WBGene00006958 | WBGene00007974 | WBGene00003851 | WBGene00000215 | WBGene00007988 | WBGene00004427 | WBGene00003618 | WBGene00000698 | WBGene00000681 |
| WBGene00003889 | WBGene00007744 | WBGene00000979 | WBGene00000565 | WBGene00010594 | WBGene00010619 | WBGene00007047 | WBGene00008655 | WBGene00003876 | WBGene00000216 | WBGene00007989 | WBGene00004428 | WBGene00003661 | WBGene00000703 | WBGene00000696 |
| WBGene00003993 | WBGene00007891 | WBGene00000990 | WBGene00000591 | WBGene00010705 | WBGene00010625 | WBGene00007105 | WBGene00008667 | WBGene00003922 | WBGene00000217 | WBGene00007996 | WBGene00004429 | WBGene00003691 | WBGene00000711 | WBGene00000699 |
| WBGene00003995 | WBGene00008150 | WBGene00001022 | WBGene00000592 | WBGene00010719 | WBGene00010758 | WBGene00007207 | WBGene00008742 | WBGene00004039 | WBGene00000218 | WBGene00008260 | WBGene00004430 | WBGene00003727 | WBGene00000713 | WBGene00000707 |
| WBGene00004035 | WBGene00008221 | WBGene00001040 | WBGene00000708 | WBGene00010768 | WBGene00010847 | WBGene00007317 | WBGene00008799 | WBGene00004140 | WBGene00000219 | WBGene00008496 | WBGene00004431 | WBGene00003746 | WBGene00000728 | WBGene00000712 |
| WBGene00004197 | WBGene00008232 | WBGene00001046 | WBGene00000765 | WBGene00010883 | WBGene00010947 | WBGene00007391 | WBGene00008912 | WBGene00004221 | WBGene00000222 | WBGene00008512 | WBGene00004432 | WBGene00003765 | WBGene00000735 | WBGene00000715 |
| WBGene00004231 | WBGene00008236 | WBGene00001432 | WBGene00000770 | WBGene00011013 | WBGene00011056 | WBGene00007403 | WBGene00009041 | WBGene00004234 | WBGene00000224 | WBGene00008605 | WBGene00004434 | WBGene00003767 | WBGene00000748 | WBGene00000717 |
| WBGene00004745 | WBGene00008423 | WBGene00001433 | WBGene00000772 | WBGene00011117 | WBGene00011108 | WBGene00007533 | WBGene00009331 | WBGene00004319 | WBGene00000228 | WBGene00009013 | WBGene00004436 | WBGene00003768 | WBGene00000751 | WBGene00000718 |
| WBGene00004777 | WBGene00008497 | WBGene00001442 | WBGene00000780 | WBGene00011325 | WBGene00011114 | WBGene00007711 | WBGene00009653 | WBGene00004324 | WBGene00000230 | WBGene00009133 | WBGene00004437 | WBGene00003865 | WBGene00000767 | WBGene00000722 |
| WBGene00005048 | WBGene00008556 | WBGene00001451 | WBGene00000803 | WBGene00011434 | WBGene00011171 | WBGene00007753 | WBGene00009879 | WBGene00004347 | WBGene00000245 | WBGene00009261 | WBGene00004444 | WBGene00003950 | WBGene00000776 | WBGene00000732 |
| WBGene00005121 | WBGene00008583 | WBGene00001454 | WBGene00000804 | WBGene00011482 | WBGene00011635 | WBGene00007756 | WBGene00009904 | WBGene00004378 | WBGene00000247 | WBGene00009528 | WBGene00004445 | WBGene00004033 | WBGene00000788 | WBGene00000733 |
| WBGene00005122 | WBGene00008740 | WBGene00001455 | WBGene00000815 | WBGene00011678 | WBGene00012429 | WBGene00007787 | WBGene00010267 | WBGene00004440 | WBGene00000248 | WBGene00010630 | WBGene00004446 | WBGene00004043 | WBGene00000800 | WBGene00000739 |
| WBGene00005123 | WBGene00008783 | WBGene00001475 | WBGene00000816 | WBGene00011976 | WBGene00012535 | WBGene00007791 | WBGene00010285 | WBGene00004441 | WBGene00000257 | WBGene00010673 | WBGene00004447 | WBGene00004108 | WBGene00000843 | WBGene00000740 |
| WBGene00005126 | WBGene00008891 | WBGene00001492 | WBGene00000817 | WBGene00012120 | WBGene00013128 | WBGene00007849 | WBGene00010468 | WBGene00004443 | WBGene00000263 | WBGene00010723 | WBGene00004449 | WBGene00004182 | WBGene00000854 | WBGene00000742 |
| WBGene00005133 | WBGene00009107 | WBGene00001539 | WBGene00000839 | WBGene00012180 | WBGene00013297 | WBGene00007856 | WBGene00010583 | WBGene00004453 | WBGene00000268 | WBGene00010760 | WBGene00004450 | WBGene00004204 | WBGene00000882 | WBGene00000754 |
| WBGene00005227 | WBGene00009108 | WBGene00001590 | WBGene00000863 | WBGene00012222 | WBGene00013980 | WBGene00007878 | WBGene00011095 | WBGene00004476 | WBGene00000274 | WBGene00011339 | WBGene00004452 | WBGene00004321 | WBGene00000897 | WBGene00000802 |
| WBGene00005241 | WBGene00009123 | WBGene00001613 | WBGene00000866 | WBGene00012272 | WBGene00014698 | WBGene00008151 | WBGene00011107 | WBGene00004478 | WBGene00000275 | WBGene00011530 | WBGene00004456 | WBGene00004326 | WBGene00000900 | WBGene00000831 |
| WBGene00005448 | WBGene00009287 | WBGene00001617 | WBGene00000872 | WBGene00012397 | WBGene00015348 | WBGene00008165 | WBGene00011201 | WBGene00004479 | WBGene00000277 | WBGene00011777 | WBGene00004470 | WBGene00004369 | WBGene00000908 | WBGene00000833 |
| WBGene00005450 | WBGene00009315 | WBGene00001636 | WBGene00000875 | WBGene00012607 | WBGene00015741 | WBGene00008333 | WBGene00011307 | WBGene00004484 | WBGene00000286 | WBGene00011895 | WBGene00004471 | WBGene00004387 | WBGene00000914 | WBGene00000871 |

| | | | | | | | | | | | | | | |
|---|---|---|---|---|---|---|---|---|---|---|---|---|---|---|
| WBGene00005477 | WBGene00009335 | WBGene00001669 | WBGene00000885 | WBGene00012670 | WBGene00015753 | WBGene00008336 | WBGene00011548 | WBGene00004486 | WBGene00000288 | WBGene00012540 | WBGene00004472 | WBGene00004462 | WBGene00000941 | WBGene00000876 |
| WBGene00006064 | WBGene00009360 | WBGene00001671 | WBGene00000889 | WBGene00012721 | WBGene00015781 | WBGene00008443 | WBGene00011590 | WBGene00004491 | WBGene00000289 | WBGene00012796 | WBGene00004473 | WBGene00004826 | WBGene00000962 | WBGene00000879 |
| WBGene00006072 | WBGene00009649 | WBGene00001693 | WBGene00000891 | WBGene00012763 | WBGene00015861 | WBGene00008621 | WBGene00011631 | WBGene00004492 | WBGene00000294 | WBGene00012917 | WBGene00004474 | WBGene00004929 | WBGene00000975 | WBGene00000881 |
| WBGene00006097 | WBGene00009713 | WBGene00001702 | WBGene00000892 | WBGene00013312 | WBGene00016136 | WBGene00008735 | WBGene00011747 | WBGene00004496 | WBGene00000370 | WBGene00013385 | WBGene00004475 | WBGene00005008 | WBGene00001029 | WBGene00000883 |
| WBGene00006106 | WBGene00009997 | WBGene00001724 | WBGene00000903 | WBGene00013556 | WBGene00016181 | WBGene00008769 | WBGene00011938 | WBGene00004498 | WBGene00000378 | WBGene00013523 | WBGene00004477 | WBGene00005648 | WBGene00001031 | WBGene00000887 |
| WBGene00006111 | WBGene00010108 | WBGene00001728 | WBGene00000904 | WBGene00013573 | WBGene00016281 | WBGene00008877 | WBGene00012032 | WBGene00004699 | WBGene00000387 | WBGene00013717 | WBGene00004480 | WBGene00006331 | WBGene00001032 | WBGene00000894 |
| WBGene00006121 | WBGene00010222 | WBGene00001732 | WBGene00000913 | WBGene00013858 | WBGene00016403 | WBGene00008878 | WBGene00012034 | WBGene00004781 | WBGene00000390 | WBGene00013883 | WBGene00004482 | WBGene00006450 | WBGene00001034 | WBGene00000898 |
| WBGene00006147 | WBGene00010455 | WBGene00001734 | WBGene00000917 | WBGene00014070 | WBGene00017168 | WBGene00008882 | WBGene00012255 | WBGene00004917 | WBGene00000392 | WBGene00015059 | WBGene00004483 | WBGene00006493 | WBGene00001039 | WBGene00000901 |
| WBGene00006236 | WBGene00010460 | WBGene00001769 | WBGene00000920 | WBGene00014246 | WBGene00017340 | WBGene00008963 | WBGene00012257 | WBGene00006058 | WBGene00000396 | WBGene00015335 | WBGene00004485 | WBGene00006504 | WBGene00001042 | WBGene00000912 |
| WBGene00006380 | WBGene00010467 | WBGene00001783 | WBGene00000939 | WBGene00015636 | WBGene00017524 | WBGene00009028 | WBGene00012275 | WBGene00006397 | WBGene00000403 | WBGene00015646 | WBGene00004487 | WBGene00006514 | WBGene00001047 | WBGene00000915 |
| WBGene00006544 | WBGene00010469 | WBGene00001791 | WBGene00000940 | WBGene00015859 | WBGene00017654 | WBGene00009087 | WBGene00012433 | WBGene00006443 | WBGene00000405 | WBGene00015795 | WBGene00004488 | WBGene00006588 | WBGene00001059 | WBGene00000928 |
| WBGene00006546 | WBGene00010566 | WBGene00001814 | WBGene00000967 | WBGene00015993 | WBGene00018430 | WBGene00009159 | WBGene00012648 | WBGene00006466 | WBGene00000409 | WBGene00015841 | WBGene00004493 | WBGene00006594 | WBGene00001067 | WBGene00000938 |
| WBGene00006665 | WBGene00010683 | WBGene00001816 | WBGene00000982 | WBGene00016081 | WBGene00018486 | WBGene00009233 | WBGene00012777 | WBGene00006641 | WBGene00000410 | WBGene00016013 | WBGene00004494 | WBGene00006601 | WBGene00001081 | WBGene00000963 |
| WBGene00006718 | WBGene00010714 | WBGene00001819 | WBGene00001001 | WBGene00016094 | WBGene00019633 | WBGene00009328 | WBGene00013173 | WBGene00006971 | WBGene00000411 | WBGene00016421 | WBGene00004495 | WBGene00006638 | WBGene00001086 | WBGene00000976 |
| WBGene00006731 | WBGene00011141 | WBGene00001858 | WBGene00001002 | WBGene00016428 | WBGene00019658 | WBGene00009382 | WBGene00013216 | WBGene00006988 | WBGene00000414 | WBGene00016740 | WBGene00004499 | WBGene00006649 | WBGene00001088 | WBGene00001006 |
| WBGene00006761 | WBGene00011184 | WBGene00001910 | WBGene00001007 | WBGene00016461 | WBGene00020136 | WBGene00009789 | WBGene00013284 | WBGene00007008 | WBGene00000415 | WBGene00017396 | WBGene00004704 | WBGene00006660 | WBGene00001089 | WBGene00001008 |
| WBGene00006781 | WBGene00011241 | WBGene00001975 | WBGene00001027 | WBGene00017039 | WBGene00020250 | WBGene00009878 | WBGene00013496 | WBGene00007015 | WBGene00000416 | WBGene00017560 | WBGene00006543 | WBGene00006704 | WBGene00001093 | WBGene00001023 |
| WBGene00006792 | WBGene00011345 | WBGene00001991 | WBGene00001033 | WBGene00017108 | WBGene00020546 | WBGene00010091 | WBGene00013592 | WBGene00007100 | WBGene00000417 | WBGene00018039 | WBGene00006565 | WBGene00006794 | WBGene00001094 | WBGene00001030 |
| WBGene00006987 | WBGene00011454 | WBGene00002018 | WBGene00001045 | WBGene00017122 | WBGene00020660 | WBGene00010315 | WBGene00013891 | WBGene00007108 | WBGene00000419 | WBGene00018515 | WBGene00006611 | WBGene00006839 | WBGene00001130 | WBGene00001038 |
| WBGene00007140 | WBGene00011524 | WBGene00002019 | WBGene00001048 | WBGene00017428 | WBGene00020673 | WBGene00010418 | WBGene00014009 | WBGene00007116 | WBGene00000424 | WBGene00018985 | WBGene00006656 | WBGene00006862 | WBGene00001137 | WBGene00001066 |
| WBGene00007262 | WBGene00011563 | WBGene00002020 | WBGene00001055 | WBGene00017650 | WBGene00020716 | WBGene00010486 | WBGene00014030 | WBGene00007236 | WBGene00000426 | WBGene00019113 | WBGene00006678 | WBGene00006916 | WBGene00001160 | WBGene00001070 |
| WBGene00007286 | WBGene00011673 | WBGene00002031 | WBGene00001056 | WBGene00017794 | WBGene00020717 | WBGene00011011 | WBGene00014245 | WBGene00007267 | WBGene00000437 | WBGene00019236 | WBGene00006706 | WBGene00007013 | WBGene00001178 | WBGene00001072 |
| WBGene00007366 | WBGene00011715 | WBGene00002103 | WBGene00001060 | WBGene00018005 | WBGene00020797 | WBGene00011216 | WBGene00015094 | WBGene00007412 | WBGene00000464 | WBGene00019290 | WBGene00006721 | WBGene00007189 | WBGene00001182 | WBGene00001073 |
| WBGene00007372 | WBGene00011796 | WBGene00002118 | WBGene00001061 | WBGene00018281 | WBGene00021502 | WBGene00011234 | WBGene00015200 | WBGene00007504 | WBGene00000465 | WBGene00019382 | WBGene00006722 | WBGene00007256 | WBGene00001208 | WBGene00001074 |
| WBGene00007397 | WBGene00011801 | WBGene00002132 | WBGene00001085 | WBGene00018587 | WBGene00022598 | WBGene00011440 | WBGene00015300 | WBGene00007573 | WBGene00000466 | WBGene00019908 | WBGene00006831 | WBGene00007288 | WBGene00001225 | WBGene00001077 |
| WBGene00007398 | WBGene00011913 | WBGene00002393 | WBGene00001090 | WBGene00018675 | WBGene00044471 | WBGene00011554 | WBGene00015541 | WBGene00007623 | WBGene00000467 | WBGene00020039 | WBGene00006964 | WBGene00007395 | WBGene00001228 | WBGene00001080 |
| WBGene00007405 | WBGene00011956 | WBGene00002804 | WBGene00001092 | WBGene00018949 | WBGene00045013 | WBGene00011589 | WBGene00015806 | WBGene00007718 | WBGene00000472 | WBGene00020626 | WBGene00007064 | WBGene00007529 | WBGene00001234 | WBGene00001131 |
| WBGene00007406 | WBGene00011968 | WBGene00002977 | WBGene00001099 | WBGene00019081 | WBGene00045253 | WBGene00011736 | WBGene00016172 | WBGene00007917 | WBGene00000473 | WBGene00020912 | WBGene00007180 | WBGene00007587 | WBGene00001262 | WBGene00001134 |
| WBGene00007465 | WBGene00012199 | WBGene00002991 | WBGene00001102 | WBGene00019287 | WBGene00045277 | WBGene00011776 | WBGene00016329 | WBGene00007944 | WBGene00000474 | WBGene00021134 | WBGene00007287 | WBGene00007592 | WBGene00001336 | WBGene00001149 |
| WBGene00007639 | WBGene00012237 | WBGene00003041 | WBGene00001145 | WBGene00019503 | WBGene00195246 | WBGene00011871 | WBGene00016628 | WBGene00008166 | WBGene00000480 | WBGene00021343 | WBGene00007444 | WBGene00007601 | WBGene00001340 | WBGene00001159 |
| WBGene00007759 | WBGene00012321 | WBGene00003046 | WBGene00001151 | WBGene00019519 | | WBGene00011908 | WBGene00016805 | WBGene00008405 | WBGene00000496 | WBGene00021389 | WBGene00007488 | WBGene00007630 | WBGene00001358 | WBGene00001163 |
| WBGene00007764 | WBGene00012477 | WBGene00003074 | WBGene00001156 | WBGene00019571 | | WBGene00011935 | WBGene00016934 | WBGene00008429 | WBGene00000500 | WBGene00021415 | WBGene00007548 | WBGene00007882 | WBGene00001387 | WBGene00001167 |
| WBGene00007810 | WBGene00012547 | WBGene00003077 | WBGene00001157 | WBGene00019700 | | WBGene00012220 | WBGene00016970 | WBGene00008441 | WBGene00000506 | WBGene00021690 | WBGene00007680 | WBGene00007890 | WBGene00001395 | WBGene00001168 |
| WBGene00007861 | WBGene00012561 | WBGene00003238 | WBGene00001186 | WBGene00019761 | | WBGene00012322 | WBGene00017307 | WBGene00008565 | WBGene00000515 | WBGene00022012 | WBGene00008094 | WBGene00007913 | WBGene00001411 | WBGene00001169 |
| WBGene00007885 | WBGene00012578 | WBGene00003245 | WBGene00001190 | WBGene00020195 | | WBGene00012483 | WBGene00017357 | WBGene00008642 | WBGene00000517 | WBGene00022554 | WBGene00008320 | WBGene00007927 | WBGene00001430 | WBGene00001173 |
| WBGene00007886 | WBGene00012609 | WBGene00003385 | WBGene00001207 | WBGene00020550 | | WBGene00012592 | WBGene00017716 | WBGene00008720 | WBGene00000527 | WBGene00022613 | WBGene00008331 | WBGene00007975 | WBGene00001441 | WBGene00001177 |
| WBGene00007893 | WBGene00012627 | WBGene00003425 | WBGene00001214 | WBGene00020625 | | WBGene00012593 | WBGene00017769 | WBGene00009188 | WBGene00000529 | WBGene00022816 | WBGene00008669 | WBGene00008052 | WBGene00001488 | WBGene00001184 |
| WBGene00007901 | WBGene00012784 | WBGene00003434 | WBGene00001224 | WBGene00020893 | | WBGene00012972 | WBGene00017807 | WBGene00009285 | WBGene00000535 | WBGene00044109 | WBGene00008714 | WBGene00008100 | WBGene00001491 | WBGene00001194 |
| WBGene00007977 | WBGene00012952 | WBGene00003446 | WBGene00001236 | WBGene00021117 | | WBGene00013140 | WBGene00017830 | WBGene00009314 | WBGene00000536 | WBGene00044344 | WBGene00008848 | WBGene00008132 | WBGene00001496 | WBGene00001196 |
| WBGene00007982 | WBGene00012976 | WBGene00003448 | WBGene00001246 | WBGene00021118 | | WBGene00013242 | WBGene00017901 | WBGene00009364 | WBGene00000545 | WBGene00044612 | WBGene00008927 | WBGene00008183 | WBGene00001498 | WBGene00001209 |
| WBGene00007983 | WBGene00013026 | WBGene00003449 | WBGene00001249 | WBGene00021135 | | WBGene00013301 | WBGene00017921 | WBGene00009451 | WBGene00000554 | WBGene00045183 | WBGene00008997 | WBGene00008195 | WBGene00001501 | WBGene00001226 |
| WBGene00007991 | WBGene00013055 | WBGene00003451 | WBGene00001281 | WBGene00021136 | | WBGene00013302 | WBGene00018723 | WBGene00009769 | WBGene00000557 | WBGene00050943 | WBGene00009002 | WBGene00008196 | WBGene00001503 | WBGene00001227 |
| WBGene00007994 | WBGene00013332 | WBGene00003466 | WBGene00001332 | WBGene00021146 | | WBGene00013324 | WBGene00018871 | WBGene00009843 | WBGene00000558 | WBGene00206373 | WBGene00009057 | WBGene00008197 | WBGene00001520 | WBGene00001239 |
| WBGene00008050 | WBGene00013375 | WBGene00003468 | WBGene00001352 | WBGene00021213 | | WBGene00013998 | WBGene00018997 | WBGene00009976 | WBGene00000566 | | WBGene00009064 | WBGene00008218 | WBGene00001561 | WBGene00001251 |
| WBGene00008054 | WBGene00013388 | WBGene00003471 | WBGene00001365 | WBGene00021285 | | WBGene00014052 | WBGene00019021 | WBGene00009977 | WBGene00000567 | | WBGene00009452 | WBGene00008332 | WBGene00001574 | WBGene00001303 |
| WBGene00008084 | WBGene00013521 | WBGene00003525 | WBGene00001445 | WBGene00021544 | | WBGene00014087 | WBGene00019060 | WBGene00010015 | WBGene00000584 | | WBGene00009499 | WBGene00008395 | WBGene00001584 | WBGene00001329 |
| WBGene00008111 | WBGene00013622 | WBGene00003568 | WBGene00001447 | WBGene00022539 | | WBGene00014091 | WBGene00019154 | WBGene00010038 | WBGene00000597 | | WBGene00009722 | WBGene00008412 | WBGene00001596 | WBGene00001330 |
| WBGene00008144 | WBGene00014116 | WBGene00003645 | WBGene00001448 | WBGene00022761 | | WBGene00014139 | WBGene00019184 | WBGene00010042 | WBGene00000611 | | WBGene00009925 | WBGene00008414 | WBGene00001628 | WBGene00001393 |
| WBGene00008156 | WBGene00014149 | WBGene00003682 | WBGene00001450 | WBGene00022780 | | WBGene00014258 | WBGene00019467 | WBGene00010045 | WBGene00000638 | | WBGene00009944 | WBGene00008514 | WBGene00001633 | WBGene00001396 |
| WBGene00008210 | WBGene00015093 | WBGene00003690 | WBGene00001456 | WBGene00022782 | | WBGene00015147 | WBGene00019520 | WBGene00010268 | WBGene00000670 | | WBGene00010001 | WBGene00008555 | WBGene00001635 | WBGene00001398 |
| WBGene00008212 | WBGene00015231 | WBGene00003712 | WBGene00001460 | WBGene00022884 | | WBGene00015148 | WBGene00019765 | WBGene00010322 | WBGene00000675 | | WBGene00010012 | WBGene00008601 | WBGene00001645 | WBGene00001405 |
| WBGene00008277 | WBGene00015466 | WBGene00003722 | WBGene00001463 | WBGene00044180 | | WBGene00015178 | WBGene00019834 | WBGene00010370 | WBGene00000692 | | WBGene00010063 | WBGene00008765 | WBGene00001651 | WBGene00001423 |
| WBGene00008279 | WBGene00015488 | WBGene00003735 | WBGene00001477 | WBGene00044194 | | WBGene00015293 | WBGene00020006 | WBGene00010911 | WBGene00000716 | | WBGene00010080 | WBGene00008780 | WBGene00001694 | WBGene00001428 |
| WBGene00008516 | WBGene00015516 | WBGene00003755 | WBGene00001481 | WBGene00044799 | | WBGene00015357 | WBGene00020281 | WBGene00010924 | WBGene00000731 | | WBGene00010232 | WBGene00008781 | WBGene00001703 | WBGene00001431 |
| WBGene00008526 | WBGene00015519 | WBGene00003778 | WBGene00001489 | WBGene00077536 | | WBGene00015529 | WBGene00020396 | WBGene00010976 | WBGene00000752 | | WBGene00010433 | WBGene00008856 | WBGene00001714 | WBGene00001443 |
| WBGene00008570 | WBGene00015523 | WBGene00003859 | WBGene00001494 | WBGene00077548 | | WBGene00015561 | WBGene00020560 | WBGene00011051 | WBGene00000757 | | WBGene00010773 | WBGene00008876 | WBGene00001716 | WBGene00001479 |
| WBGene00008579 | WBGene00015855 | WBGene00003939 | WBGene00001518 | WBGene00164968 | | WBGene00015786 | WBGene00020567 | WBGene00011055 | WBGene00000764 | | WBGene00010776 | WBGene00008886 | WBGene00001747 | WBGene00001493 |
| WBGene00008623 | WBGene00015968 | WBGene00003954 | WBGene00001522 | WBGene00194742 | | WBGene00015915 | WBGene00020646 | WBGene00011302 | WBGene00000773 | | WBGene00010827 | WBGene00008993 | WBGene00001755 | WBGene00001502 |
| WBGene00008644 | WBGene00016040 | WBGene00003957 | WBGene00001526 | WBGene00206484 | | WBGene00016104 | WBGene00020672 | WBGene00011344 | WBGene00000775 | | WBGene00011088 | WBGene00009243 | WBGene00001824 | WBGene00001510 |
| WBGene00008679 | WBGene00016132 | WBGene00003977 | WBGene00001563 | WBGene00206529 | | WBGene00016156 | WBGene00021627 | WBGene00011823 | WBGene00000777 | | WBGene00011185 | WBGene00009341 | WBGene00001840 | WBGene00001527 |
| WBGene00008706 | WBGene00016364 | WBGene00003991 | WBGene00001568 | WBGene00219317 | | WBGene00016188 | WBGene00021653 | WBGene00011888 | WBGene00000779 | | WBGene00011239 | WBGene00009380 | WBGene00001972 | WBGene00001558 |
| WBGene00008746 | WBGene00016492 | WBGene00004003 | WBGene00001571 | WBGene00220261 | | WBGene00016283 | WBGene00021850 | WBGene00011972 | WBGene00000781 | | WBGene00011337 | WBGene00009439 | WBGene00001999 | WBGene00001559 |
| WBGene00008752 | WBGene00016527 | WBGene00004174 | WBGene00001609 | | | WBGene00016446 | WBGene00022196 | WBGene00011974 | WBGene00000784 | | WBGene00011348 | WBGene00009626 | WBGene00002038 | WBGene00001562 |
| WBGene00008952 | WBGene00016565 | WBGene00004184 | WBGene00001629 | | | WBGene00016524 | WBGene00022274 | WBGene00012168 | WBGene00000785 | | WBGene00011694 | WBGene00009674 | WBGene00002046 | WBGene00001564 |
| WBGene00008959 | WBGene00016662 | WBGene00004193 | WBGene00001637 | | | WBGene00016657 | WBGene00022665 | WBGene00012188 | WBGene00000786 | | WBGene00011738 | WBGene00009726 | WBGene00002059 | WBGene00001565 |
| WBGene00008972 | WBGene00016700 | WBGene00004223 | WBGene00001647 | | | WBGene00016860 | WBGene00022857 | WBGene00012216 | WBGene00000789 | | WBGene00011828 | WBGene00009888 | WBGene00002061 | WBGene00001582 |
| WBGene00009054 | WBGene00016742 | WBGene00004257 | WBGene00001650 | | | WBGene00016889 | WBGene00044392 | WBGene00012375 | WBGene00000793 | | WBGene00012007 | WBGene00010018 | WBGene00002066 | WBGene00001592 |
| WBGene00009060 | WBGene00016877 | WBGene00004278 | WBGene00001653 | | | WBGene00016914 | WBGene00044395 | WBGene00012487 | WBGene00000794 | | WBGene00012106 | WBGene00010079 | WBGene00002071 | WBGene00001604 |

| | | | | | | | | | | | |
|---|---|---|---|---|---|---|---|---|---|---|---|
| WBGene00009160 | WBGene00016951 | WBGene00004309 | WBGene00001660 | WBGene00016916 | WBGene00044413 | WBGene00012722 | WBGene00000796 | WBGene00012152 | WBGene00010115 | WBGene00002081 | WBGene00001631 |
| WBGene00009178 | WBGene00016960 | WBGene00004346 | WBGene00001709 | WBGene00017022 | WBGene00044445 | WBGene00013532 | WBGene00000812 | WBGene00012330 | WBGene00010117 | WBGene00002083 | WBGene00001649 |
| WBGene00009232 | WBGene00016962 | WBGene00004363 | WBGene00001731 | WBGene00017046 | WBGene00044530 | WBGene00013986 | WBGene00000813 | WBGene00012472 | WBGene00010224 | WBGene00002152 | WBGene00001684 |
| WBGene00009239 | WBGene00017350 | WBGene00004404 | WBGene00001749 | WBGene00017471 | WBGene00044895 | WBGene00014123 | WBGene00000814 | WBGene00013007 | WBGene00010251 | WBGene00002162 | WBGene00001685 |
| WBGene00009308 | WBGene00017388 | WBGene00004801 | WBGene00001752 | WBGene00017536 | WBGene00077728 | WBGene00014176 | WBGene00000822 | WBGene00013260 | WBGene00010356 | WBGene00002191 | WBGene00001713 |
| WBGene00009339 | WBGene00017442 | WBGene00004811 | WBGene00001757 | WBGene00017580 | WBGene00195068 | WBGene00014218 | WBGene00000832 | WBGene00013266 | WBGene00010409 | WBGene00002196 | WBGene00001725 |
| WBGene00009358 | WBGene00017767 | WBGene00004816 | WBGene00001758 | WBGene00017771 | WBGene00206515 | WBGene00014250 | WBGene00000834 | WBGene00013418 | WBGene00010525 | WBGene00002201 | WBGene00001744 |
| WBGene00009386 | WBGene00017802 | WBGene00004824 | WBGene00001772 | WBGene00017781 | WBGene00235268 | WBGene00015011 | WBGene00000836 | WBGene00013438 | WBGene00010730 | WBGene00002202 | WBGene00001748 |
| WBGene00009618 | WBGene00017902 | WBGene00004893 | WBGene00001808 | WBGene00017840 | WBGene00235269 | WBGene00015196 | WBGene00000838 | WBGene00013499 | WBGene00010731 | WBGene00002203 | WBGene00001843 |
| WBGene00009619 | WBGene00018018 | WBGene00004898 | WBGene00001812 | WBGene00017981 | | WBGene00015203 | WBGene00000840 | WBGene00013650 | WBGene00010766 | WBGene00002207 | WBGene00001851 |
| WBGene00009622 | WBGene00018153 | WBGene00004966 | WBGene00001813 | WBGene00018226 | | WBGene00015206 | WBGene00000846 | WBGene00013920 | WBGene00010770 | WBGene00002223 | WBGene00001955 |
| WBGene00009623 | WBGene00018278 | WBGene00004996 | WBGene00001820 | WBGene00018232 | | WBGene00015266 | WBGene00000865 | WBGene00013923 | WBGene00010778 | WBGene00002238 | WBGene00001993 |
| WBGene00009625 | WBGene00018292 | WBGene00005020 | WBGene00001830 | WBGene00018405 | | WBGene00015278 | WBGene00000867 | WBGene00014051 | WBGene00011003 | WBGene00002250 | WBGene00001994 |
| WBGene00009658 | WBGene00018296 | WBGene00005080 | WBGene00001833 | WBGene00018703 | | WBGene00015326 | WBGene00000868 | WBGene00014093 | WBGene00011015 | WBGene00002266 | WBGene00002000 |
| WBGene00009728 | WBGene00018338 | WBGene00005292 | WBGene00001835 | WBGene00019011 | | WBGene00015402 | WBGene00000869 | WBGene00015021 | WBGene00011071 | WBGene00002295 | WBGene00002005 |
| WBGene00009811 | WBGene00018473 | WBGene00005712 | WBGene00001836 | WBGene00019326 | | WBGene00015434 | WBGene00000870 | WBGene00015138 | WBGene00011292 | WBGene00002335 | WBGene00002007 |
| WBGene00009830 | WBGene00018591 | WBGene00006038 | WBGene00001837 | WBGene00019365 | | WBGene00015477 | WBGene00000874 | WBGene00015647 | WBGene00011415 | WBGene00002344 | WBGene00002008 |
| WBGene00009868 | WBGene00018706 | WBGene00006039 | WBGene00001844 | WBGene00019518 | | WBGene00015497 | WBGene00000877 | WBGene00016093 | WBGene00011451 | WBGene00002632 | WBGene00002010 |
| WBGene00009929 | WBGene00018774 | WBGene00006040 | WBGene00001865 | WBGene00019648 | | WBGene00015577 | WBGene00000886 | WBGene00016095 | WBGene00011527 | WBGene00002637 | WBGene00002024 |
| WBGene00009935 | WBGene00018840 | WBGene00006055 | WBGene00001875 | WBGene00019823 | | WBGene00015671 | WBGene00000896 | WBGene00016148 | WBGene00011540 | WBGene00002783 | WBGene00002025 |
| WBGene00009984 | WBGene00018954 | WBGene00006395 | WBGene00001876 | WBGene00019955 | | WBGene00015743 | WBGene00000899 | WBGene00016644 | WBGene00011735 | WBGene00002981 | WBGene00002037 |
| WBGene00010144 | WBGene00018999 | WBGene00006404 | WBGene00001877 | WBGene00020036 | | WBGene00016123 | WBGene00000910 | WBGene00016981 | WBGene00011756 | WBGene00003002 | WBGene00002045 |
| WBGene00010170 | WBGene00019006 | WBGene00006491 | WBGene00001878 | WBGene00020052 | | WBGene00016147 | WBGene00000911 | WBGene00017123 | WBGene00011814 | WBGene00003003 | WBGene00002053 |
| WBGene00010173 | WBGene00019084 | WBGene00006572 | WBGene00001879 | WBGene00020096 | | WBGene00016313 | WBGene00000929 | WBGene00017272 | WBGene00011832 | WBGene00003014 | WBGene00002065 |
| WBGene00010205 | WBGene00019197 | WBGene00006609 | WBGene00001880 | WBGene00020237 | | WBGene00016418 | WBGene00000931 | WBGene00017301 | WBGene00011885 | WBGene00003025 | WBGene00002070 |
| WBGene00010289 | WBGene00019255 | WBGene00006624 | WBGene00001881 | WBGene00020486 | | WBGene00016603 | WBGene00000935 | WBGene00018015 | WBGene00011953 | WBGene00003026 | WBGene00002074 |
| WBGene00010359 | WBGene00019376 | WBGene00006636 | WBGene00001891 | WBGene00020629 | | WBGene00016810 | WBGene00000936 | WBGene00018302 | WBGene00012004 | WBGene00003030 | WBGene00002077 |
| WBGene00010387 | WBGene00019536 | WBGene00006661 | WBGene00001892 | WBGene00020718 | | WBGene00016968 | WBGene00000937 | WBGene00018566 | WBGene00012005 | WBGene00003048 | WBGene00002078 |
| WBGene00010434 | WBGene00019640 | WBGene00006700 | WBGene00001905 | WBGene00020719 | | WBGene00017084 | WBGene00000942 | WBGene00018738 | WBGene00012337 | WBGene00003052 | WBGene00002080 |
| WBGene00010450 | WBGene00019688 | WBGene00006824 | WBGene00001906 | WBGene00020884 | | WBGene00017534 | WBGene00000958 | WBGene00018786 | WBGene00012347 | WBGene00003059 | WBGene00002127 |
| WBGene00010493 | WBGene00019962 | WBGene00006827 | WBGene00001911 | WBGene00021049 | | WBGene00017537 | WBGene00000959 | WBGene00018996 | WBGene00012348 | WBGene00003061 | WBGene00002134 |
| WBGene00010599 | WBGene00019964 | WBGene00006852 | WBGene00001912 | WBGene00021073 | | WBGene00017547 | WBGene00000961 | WBGene00019380 | WBGene00012351 | WBGene00003064 | WBGene00002135 |
| WBGene00010606 | WBGene00020019 | WBGene00006875 | WBGene00001914 | WBGene00021157 | | WBGene00017608 | WBGene00000969 | WBGene00019825 | WBGene00012358 | WBGene00003068 | WBGene00002163 |
| WBGene00010666 | WBGene00020318 | WBGene00006881 | WBGene00001922 | WBGene00021487 | | WBGene00017686 | WBGene00000970 | WBGene00019841 | WBGene00012440 | WBGene00003072 | WBGene00002189 |
| WBGene00010682 | WBGene00020500 | WBGene00006951 | WBGene00001933 | WBGene00021522 | | WBGene00017745 | WBGene00000985 | WBGene00019871 | WBGene00012484 | WBGene00003073 | WBGene00002258 |
| WBGene00010690 | WBGene00020571 | WBGene00006960 | WBGene00001934 | WBGene00021540 | | WBGene00017758 | WBGene00000988 | WBGene00019961 | WBGene00012978 | WBGene00003082 | WBGene00002262 |
| WBGene00010704 | WBGene00020593 | WBGene00006978 | WBGene00001935 | WBGene00021556 | | WBGene00017776 | WBGene00000991 | WBGene00019979 | WBGene00013008 | WBGene00003085 | WBGene00002264 |
| WBGene00010761 | WBGene00020594 | WBGene00006980 | WBGene00001936 | WBGene00021633 | | WBGene00017778 | WBGene00000993 | WBGene00020300 | WBGene00013146 | WBGene00003086 | WBGene00002265 |
| WBGene00010808 | WBGene00021175 | WBGene00007018 | WBGene00001937 | WBGene00021785 | | WBGene00017800 | WBGene00000996 | WBGene00020346 | WBGene00013217 | WBGene00003234 | WBGene00002363 |
| WBGene00010833 | WBGene00021269 | WBGene00007065 | WBGene00001938 | WBGene00021816 | | WBGene00017814 | WBGene00000998 | WBGene00020566 | WBGene00013543 | WBGene00003378 | WBGene00002717 |
| WBGene00010837 | WBGene00021309 | WBGene00007070 | WBGene00001941 | WBGene00021871 | | WBGene00017936 | WBGene00001000 | WBGene00020919 | WBGene00013577 | WBGene00003388 | WBGene00002845 |
| WBGene00010871 | WBGene00021333 | WBGene00007103 | WBGene00001942 | WBGene00021929 | | WBGene00017962 | WBGene00001017 | WBGene00020947 | WBGene00013639 | WBGene00003391 | WBGene00002879 |
| WBGene00010937 | WBGene00021451 | WBGene00007106 | WBGene00001945 | WBGene00022100 | | WBGene00018259 | WBGene00001019 | WBGene00021047 | WBGene00013688 | WBGene00003516 | WBGene00002889 |
| WBGene00011017 | WBGene00021596 | WBGene00007124 | WBGene00001946 | WBGene00022282 | | WBGene00018290 | WBGene00001025 | WBGene00021080 | WBGene00014001 | WBGene00003530 | WBGene00002891 |
| WBGene00011030 | WBGene00021611 | WBGene00007131 | WBGene00001973 | WBGene00022368 | | WBGene00018512 | WBGene00001026 | WBGene00022129 | WBGene00014075 | WBGene00003553 | WBGene00002915 |
| WBGene00011074 | WBGene00021679 | WBGene00007172 | WBGene00001995 | WBGene00022629 | | WBGene00018632 | WBGene00001028 | WBGene00022788 | WBGene00014092 | WBGene00003554 | WBGene00003009 |
| WBGene00011140 | WBGene00021715 | WBGene00007174 | WBGene00002004 | WBGene00022792 | | WBGene00018635 | WBGene00001035 | WBGene00044767 | WBGene00014171 | WBGene00003559 | WBGene00003012 |
| WBGene00011159 | WBGene00021976 | WBGene00007187 | WBGene00002021 | WBGene00023415 | | WBGene00018866 | WBGene00001037 | WBGene00044789 | WBGene00014262 | WBGene00003564 | WBGene00003034 |
| WBGene00011173 | WBGene00022112 | WBGene00007194 | WBGene00002028 | WBGene00044258 | | WBGene00018878 | WBGene00001041 | WBGene00045488 | WBGene00015008 | WBGene00003566 | WBGene00003040 |
| WBGene00011177 | WBGene00022117 | WBGene00007200 | WBGene00002032 | WBGene00044916 | | WBGene00019077 | WBGene00001049 | WBGene00013458 | WBGene00015141 | WBGene00003586 | WBGene00003044 |
| WBGene00011183 | WBGene00022256 | WBGene00007201 | WBGene00002034 | WBGene00045063 | | WBGene00019232 | WBGene00001051 | | WBGene00015208 | WBGene00003596 | WBGene00003055 |
| WBGene00011411 | WBGene00022457 | WBGene00007210 | WBGene00002035 | WBGene00045433 | | WBGene00019259 | WBGene00001082 | | WBGene00015235 | WBGene00003619 | WBGene00003063 |
| WBGene00011433 | WBGene00022627 | WBGene00007211 | WBGene00002041 | WBGene00077714 | | WBGene00019280 | WBGene00001087 | | WBGene00015291 | WBGene00003623 | WBGene00003071 |
| WBGene00011435 | WBGene00022808 | WBGene00007212 | WBGene00002043 | | | WBGene00019599 | WBGene00001101 | | WBGene00015430 | WBGene00003628 | WBGene00003112 |
| WBGene00011497 | WBGene00022837 | WBGene00007216 | WBGene00002052 | | | WBGene00019615 | WBGene00001112 | | WBGene00015484 | WBGene00003635 | WBGene00003124 |
| WBGene00011615 | WBGene00023425 | WBGene00007219 | WBGene00002056 | | | WBGene00019645 | WBGene00001113 | | WBGene00015508 | WBGene00003639 | WBGene00003132 |
| WBGene00011643 | WBGene00043120 | WBGene00007243 | WBGene00002067 | | | WBGene00019836 | WBGene00001132 | | WBGene00015684 | WBGene00003659 | WBGene00003144 |
| WBGene00011698 | WBGene00043279 | WBGene00007273 | WBGene00002069 | | | WBGene00019878 | WBGene00001152 | | WBGene00015759 | WBGene00003678 | WBGene00003151 |
| WBGene00011751 | WBGene00043308 | WBGene00007303 | WBGene00002072 | | | WBGene00019882 | WBGene00001153 | | WBGene00015769 | WBGene00003695 | WBGene00003161 |
| WBGene00011770 | WBGene00044227 | WBGene00007332 | WBGene00002123 | | | WBGene00019963 | WBGene00001155 | | WBGene00015866 | WBGene00003762 | WBGene00003162 |
| WBGene00011877 | WBGene00044257 | WBGene00007334 | WBGene00002136 | | | WBGene00019994 | WBGene00001161 | | WBGene00015887 | WBGene00003773 | WBGene00003163 |
| WBGene00011899 | WBGene00044287 | WBGene00007335 | WBGene00002144 | | | WBGene00020104 | WBGene00001162 | | WBGene00015896 | WBGene00003789 | WBGene00003178 |
| WBGene00011959 | WBGene00044293 | WBGene00007338 | WBGene00002146 | | | WBGene00020283 | WBGene00001166 | | WBGene00015919 | WBGene00003796 | WBGene00003214 |
| WBGene00012169 | WBGene00044539 | WBGene00007421 | WBGene00002153 | | | WBGene00020382 | WBGene00001172 | | WBGene00015926 | WBGene00003825 | WBGene00003218 |
| WBGene00012178 | WBGene00044568 | WBGene00007422 | WBGene00002176 | | | WBGene00020413 | WBGene00001187 | | WBGene00016020 | WBGene00003827 | WBGene00003367 |
| WBGene00012219 | WBGene00044637 | WBGene00007439 | WBGene00002179 | | | WBGene00020422 | WBGene00001189 | | WBGene00016119 | WBGene00003830 | WBGene00003400 |
| WBGene00012224 | WBGene00044901 | WBGene00007459 | WBGene00002216 | | | WBGene00020504 | WBGene00001230 | | WBGene00016269 | WBGene00003904 | WBGene00003476 |
| WBGene00012324 | WBGene00050939 | WBGene00007473 | WBGene00002217 | | | WBGene00020559 | WBGene00001231 | | WBGene00016292 | WBGene00003911 | WBGene00003482 |

| | | | | | | | | |
|---|---|---|---|---|---|---|---|---|
| WBGene00012336 | WBGene00077490 | WBGene00007487 | WBGene00002220 | WBGene00021202 | WBGene00001232 | WBGene00016325 | WBGene00003914 | WBGene00003485 |
| WBGene00012405 | WBGene00086546 | WBGene00007489 | WBGene00002225 | WBGene00021209 | WBGene00001235 | WBGene00016331 | WBGene00003920 | WBGene00003497 |
| WBGene00012408 | WBGene00175031 | WBGene00007507 | WBGene00002232 | WBGene00021473 | WBGene00001243 | WBGene00016408 | WBGene00003929 | WBGene00003511 |
| WBGene00012436 | WBGene00175035 | WBGene00007513 | WBGene00002239 | WBGene00021787 | WBGene00001244 | WBGene00016500 | WBGene00003947 | WBGene00003528 |
| WBGene00012710 | WBGene00185001 | WBGene00007531 | WBGene00002242 | WBGene00022148 | WBGene00001248 | WBGene00016507 | WBGene00003949 | WBGene00003588 |
| WBGene00012754 | WBGene00194737 | WBGene00007541 | WBGene00002245 | WBGene00022155 | WBGene00001258 | WBGene00016508 | WBGene00003953 | WBGene00003598 |
| WBGene00012791 | WBGene00194916 | WBGene00007543 | WBGene00002276 | WBGene00022313 | WBGene00001259 | WBGene00016535 | WBGene00003961 | WBGene00003600 |
| WBGene00012886 | WBGene00194986 | WBGene00007547 | WBGene00002278 | WBGene00022423 | WBGene00001263 | WBGene00016576 | WBGene00003982 | WBGene00003610 |
| WBGene00012942 | WBGene00195011 | WBGene00007585 | WBGene00002299 | WBGene00022427 | WBGene00001309 | WBGene00016611 | WBGene00004015 | WBGene00003613 |
| WBGene00012947 | WBGene00195239 | WBGene00007589 | WBGene00002368 | WBGene00022501 | WBGene00001320 | WBGene00016645 | WBGene00004025 | WBGene00003622 |
| WBGene00012966 | WBGene00206361 | WBGene00007613 | WBGene00002850 | WBGene00022739 | WBGene00001324 | WBGene00016655 | WBGene00004034 | WBGene00003625 |
| WBGene00013013 | WBGene00206374 | WBGene00007668 | WBGene00002982 | WBGene00022759 | WBGene00001328 | WBGene00016659 | WBGene00004040 | WBGene00003633 |
| WBGene00013187 | WBGene00206517 | WBGene00007682 | WBGene00002985 | WBGene00023450 | WBGene00001331 | WBGene00016661 | WBGene00004053 | WBGene00003636 |
| WBGene00013192 | WBGene00206525 | WBGene00007731 | WBGene00002989 | WBGene00023489 | WBGene00001333 | WBGene00016671 | WBGene00004075 | WBGene00003638 |
| WBGene00013244 | WBGene00219319 | WBGene00007734 | WBGene00002992 | WBGene00044582 | WBGene00001334 | WBGene00016728 | WBGene00004076 | WBGene00003656 |
| WBGene00013255 | WBGene00235340 | WBGene00007762 | WBGene00002996 | WBGene00050941 | WBGene00001335 | WBGene00016741 | WBGene00004077 | WBGene00003658 |
| WBGene00013287 | WBGene00235374 | WBGene00007774 | WBGene00003000 | WBGene00077500 | WBGene00001337 | WBGene00016894 | WBGene00004091 | WBGene00003681 |
| WBGene00013304 | WBGene00236789 | WBGene00007806 | WBGene00003006 | WBGene00077712 | WBGene00001345 | WBGene00016992 | WBGene00004124 | WBGene00003687 |
| WBGene00013436 | | WBGene00007825 | WBGene00003007 | | WBGene00001371 | WBGene00017023 | WBGene00004134 | WBGene00003704 |
| WBGene00013439 | | WBGene00007862 | WBGene00003017 | | WBGene00001385 | WBGene00017283 | WBGene00004136 | WBGene00003728 |
| WBGene00013507 | | WBGene00007863 | WBGene00003019 | | WBGene00001386 | WBGene00017293 | WBGene00004147 | WBGene00003736 |
| WBGene00013531 | | WBGene00007883 | WBGene00003020 | | WBGene00001389 | WBGene00017302 | WBGene00004152 | WBGene00003771 |
| WBGene00013568 | | WBGene00007900 | WBGene00003022 | | WBGene00001390 | WBGene00017390 | WBGene00004167 | WBGene00003776 |
| WBGene00013599 | | WBGene00007909 | WBGene00003043 | | WBGene00001391 | WBGene00017582 | WBGene00004187 | WBGene00003785 |
| WBGene00013871 | | WBGene00007916 | WBGene00003060 | | WBGene00001392 | WBGene00017735 | WBGene00004205 | WBGene00003793 |
| WBGene00013886 | | WBGene00007932 | WBGene00003093 | | WBGene00001394 | WBGene00017757 | WBGene00004208 | WBGene00003795 |
| WBGene00013902 | | WBGene00007938 | WBGene00003111 | | WBGene00001404 | WBGene00017770 | WBGene00004230 | WBGene00003818 |
| WBGene00014046 | | WBGene00007950 | WBGene00003133 | | WBGene00001406 | WBGene00017834 | WBGene00004235 | WBGene00003821 |
| WBGene00014089 | | WBGene00007963 | WBGene00003134 | | WBGene00001410 | WBGene00017885 | WBGene00004254 | WBGene00003822 |
| WBGene00014103 | | WBGene00008059 | WBGene00003136 | | WBGene00001425 | WBGene00017954 | WBGene00004259 | WBGene00003831 |
| WBGene00014128 | | WBGene00008076 | WBGene00003154 | | WBGene00001426 | WBGene00017982 | WBGene00004271 | WBGene00003847 |
| WBGene00014129 | | WBGene00008118 | WBGene00003156 | | WBGene00001427 | WBGene00018006 | WBGene00004274 | WBGene00003858 |
| WBGene00014235 | | WBGene00008155 | WBGene00003165 | | WBGene00001439 | WBGene00018131 | WBGene00004304 | WBGene00003887 |
| WBGene00014699 | | WBGene00008174 | WBGene00003169 | | WBGene00001444 | WBGene00018267 | WBGene00004306 | WBGene00003891 |
| WBGene00014965 | | WBGene00008176 | WBGene00003220 | | WBGene00001446 | WBGene00018349 | WBGene00004311 | WBGene00003902 |
| WBGene00015040 | | WBGene00008191 | WBGene00003224 | | WBGene00001449 | WBGene00018395 | WBGene00004339 | WBGene00003903 |
| WBGene00015044 | | WBGene00008198 | WBGene00003236 | | WBGene00001452 | WBGene00018400 | WBGene00004371 | WBGene00003915 |
| WBGene00015320 | | WBGene00008206 | WBGene00003241 | | WBGene00001457 | WBGene00018532 | WBGene00004373 | WBGene00003916 |
| WBGene00015373 | | WBGene00008216 | WBGene00003258 | | WBGene00001458 | WBGene00018657 | WBGene00004459 | WBGene00003918 |
| WBGene00015435 | | WBGene00008239 | WBGene00003373 | | WBGene00001459 | WBGene00018702 | WBGene00004463 | WBGene00003928 |
| WBGene00015458 | | WBGene00008273 | WBGene00003374 | | WBGene00001461 | WBGene00018710 | WBGene00004467 | WBGene00003936 |
| WBGene00015467 | | WBGene00008288 | WBGene00003380 | | WBGene00001462 | WBGene00018719 | WBGene00004501 | WBGene00003951 |
| WBGene00015474 | | WBGene00008304 | WBGene00003396 | | WBGene00001464 | WBGene00018739 | WBGene00004502 | WBGene00003962 |
| WBGene00015496 | | WBGene00008344 | WBGene00003405 | | WBGene00001470 | WBGene00018764 | WBGene00004503 | WBGene00003963 |
| WBGene00015558 | | WBGene00008364 | WBGene00003408 | | WBGene00001484 | WBGene00018804 | WBGene00004504 | WBGene00003964 |
| WBGene00015615 | | WBGene00008384 | WBGene00003411 | | WBGene00001485 | WBGene00018847 | WBGene00004682 | WBGene00003968 |
| WBGene00015740 | | WBGene00008402 | WBGene00003412 | | WBGene00001499 | WBGene00018974 | WBGene00004698 | WBGene00004014 |
| WBGene00015796 | | WBGene00008409 | WBGene00003413 | | WBGene00001509 | WBGene00019102 | WBGene00004701 | WBGene00004095 |
| WBGene00015858 | | WBGene00008413 | WBGene00003422 | | WBGene00001511 | WBGene00019111 | WBGene00004703 | WBGene00004109 |
| WBGene00015860 | | WBGene00008418 | WBGene00003424 | | WBGene00001513 | WBGene00019185 | WBGene00004738 | WBGene00004110 |
| WBGene00015994 | | WBGene00008421 | WBGene00003431 | | WBGene00001523 | WBGene00019211 | WBGene00004739 | WBGene00004120 |
| WBGene00016069 | | WBGene00008436 | WBGene00003432 | | WBGene00001572 | WBGene00019294 | WBGene00004756 | WBGene00004131 |
| WBGene00016105 | | WBGene00008439 | WBGene00003438 | | WBGene00001579 | WBGene00019298 | WBGene00004759 | WBGene00004132 |
| WBGene00016187 | | WBGene00008442 | WBGene00003442 | | WBGene00001581 | WBGene00019317 | WBGene00004767 | WBGene00004143 |
| WBGene00016196 | | WBGene00008447 | WBGene00003443 | | WBGene00001583 | WBGene00019457 | WBGene00004773 | WBGene00004164 |
| WBGene00016208 | | WBGene00008453 | WBGene00003456 | | WBGene00001585 | WBGene00019487 | WBGene00004788 | WBGene00004172 |
| WBGene00016320 | | WBGene00008483 | WBGene00003458 | | WBGene00001591 | WBGene00019509 | WBGene00004804 | WBGene00004175 |
| WBGene00016529 | | WBGene00008484 | WBGene00003470 | | WBGene00001595 | WBGene00019693 | WBGene00004807 | WBGene00004183 |
| WBGene00016533 | | WBGene00008500 | WBGene00003472 | | WBGene00001597 | WBGene00019697 | WBGene00004829 | WBGene00004189 |
| WBGene00016559 | | WBGene00008536 | WBGene00003474 | | WBGene00001598 | WBGene00019719 | WBGene00004855 | WBGene00004210 |
| WBGene00016613 | | WBGene00008557 | WBGene00003475 | | WBGene00001599 | WBGene00019726 | WBGene00004858 | WBGene00004217 |
| WBGene00016665 | | WBGene00008569 | WBGene00003477 | | WBGene00001602 | WBGene00019780 | WBGene00004884 | WBGene00004222 |
| WBGene00016767 | | WBGene00008576 | WBGene00003479 | | WBGene00001606 | WBGene00019827 | WBGene00004889 | WBGene00004227 |
| WBGene00016788 | | WBGene00008582 | WBGene00003504 | | WBGene00001607 | WBGene00019831 | WBGene00004897 | WBGene00004237 |
| WBGene00016927 | | WBGene00008589 | WBGene00003507 | | WBGene00001630 | WBGene00019946 | WBGene00004946 | WBGene00004245 |
| WBGene00016956 | | WBGene00008590 | WBGene00003519 | | WBGene00001632 | WBGene00020112 | WBGene00004951 | WBGene00004256 |
| WBGene00016976 | | WBGene00008640 | WBGene00003555 | | WBGene00001648 | WBGene00020181 | WBGene00005009 | WBGene00004258 |
| WBGene00016982 | | WBGene00008680 | WBGene00003561 | | WBGene00001662 | WBGene00020339 | WBGene00005015 | WBGene00004264 |

| | | | | | | |
|---|---|---|---|---|---|---|
| WBGene00016993 | WBGene00008681 | WBGene00003577 | WBGene00001678 | WBGene00020446 | WBGene00005019 | WBGene00004272 |
| WBGene00017082 | WBGene00008695 | WBGene00003578 | WBGene00001679 | WBGene00020497 | WBGene00005021 | WBGene00004300 |
| WBGene00017124 | WBGene00008726 | WBGene00003581 | WBGene00001681 | WBGene00020604 | WBGene00005022 | WBGene00004313 |
| WBGene00017160 | WBGene00008732 | WBGene00003584 | WBGene00001686 | WBGene00020636 | WBGene00005023 | WBGene00004336 |
| WBGene00017258 | WBGene00008768 | WBGene00003585 | WBGene00001692 | WBGene00020658 | WBGene00006307 | WBGene00004358 |
| WBGene00017326 | WBGene00008794 | WBGene00003590 | WBGene00001699 | WBGene00020693 | WBGene00006364 | WBGene00004397 |
| WBGene00017329 | WBGene00008824 | WBGene00003592 | WBGene00001708 | WBGene00020781 | WBGene00006367 | WBGene00004398 |
| WBGene00017401 | WBGene00008825 | WBGene00003594 | WBGene00001729 | WBGene00020917 | WBGene00006392 | WBGene00004408 |
| WBGene00017559 | WBGene00008861 | WBGene00003624 | WBGene00001743 | WBGene00020931 | WBGene00006410 | WBGene00004409 |
| WBGene00017568 | WBGene00008875 | WBGene00003640 | WBGene00001745 | WBGene00020948 | WBGene00006414 | WBGene00004410 |
| WBGene00017570 | WBGene00008910 | WBGene00003646 | WBGene00001746 | WBGene00020949 | WBGene00006423 | WBGene00004458 |
| WBGene00017629 | WBGene00008951 | WBGene00003647 | WBGene00001761 | WBGene00020951 | WBGene00006432 | WBGene00004464 |
| WBGene00017638 | WBGene00009008 | WBGene00003651 | WBGene00001776 | WBGene00020957 | WBGene00006434 | WBGene00004465 |
| WBGene00017655 | WBGene00009033 | WBGene00003652 | WBGene00001790 | WBGene00021043 | WBGene00006437 | WBGene00004469 |
| WBGene00017693 | WBGene00009090 | WBGene00003668 | WBGene00001793 | WBGene00021079 | WBGene00006438 | WBGene00004736 |
| WBGene00017694 | WBGene00009091 | WBGene00003669 | WBGene00001794 | WBGene00021149 | WBGene00006446 | WBGene00004751 |
| WBGene00017700 | WBGene00009106 | WBGene00003670 | WBGene00001815 | WBGene00021156 | WBGene00006465 | WBGene00004755 |
| WBGene00017707 | WBGene00009118 | WBGene00003676 | WBGene00001821 | WBGene00021170 | WBGene00006467 | WBGene00004760 |
| WBGene00017714 | WBGene00009128 | WBGene00003680 | WBGene00001827 | WBGene00021248 | WBGene00006474 | WBGene00004768 |
| WBGene00017722 | WBGene00009129 | WBGene00003697 | WBGene00001829 | WBGene00021292 | WBGene00006476 | WBGene00004783 |
| WBGene00017743 | WBGene00009131 | WBGene00003720 | WBGene00001831 | WBGene00021460 | WBGene00006494 | WBGene00004859 |
| WBGene00017795 | WBGene00009138 | WBGene00003723 | WBGene00001832 | WBGene00021562 | WBGene00006497 | WBGene00004862 |
| WBGene00017922 | WBGene00009176 | WBGene00003725 | WBGene00001834 | WBGene00021646 | WBGene00006530 | WBGene00004900 |
| WBGene00017971 | WBGene00009212 | WBGene00003733 | WBGene00001852 | WBGene00021845 | WBGene00006568 | WBGene00004902 |
| WBGene00018003 | WBGene00009213 | WBGene00003738 | WBGene00001853 | WBGene00021863 | WBGene00006574 | WBGene00004911 |
| WBGene00018009 | WBGene00009226 | WBGene00003747 | WBGene00001855 | WBGene00021890 | WBGene00006575 | WBGene00004930 |
| WBGene00018070 | WBGene00009257 | WBGene00003748 | WBGene00001856 | WBGene00021909 | WBGene00006579 | WBGene00004945 |
| WBGene00018111 | WBGene00009266 | WBGene00003754 | WBGene00001860 | WBGene00022059 | WBGene00006606 | WBGene00004947 |
| WBGene00018204 | WBGene00009269 | WBGene00003758 | WBGene00001862 | WBGene00022125 | WBGene00006617 | WBGene00004980 |
| WBGene00018206 | WBGene00009292 | WBGene00003764 | WBGene00001863 | WBGene00022127 | WBGene00006640 | WBGene00005016 |
| WBGene00018283 | WBGene00009304 | WBGene00003779 | WBGene00001874 | WBGene00022386 | WBGene00006708 | WBGene00005018 |
| WBGene00018373 | WBGene00009325 | WBGene00003783 | WBGene00001971 | WBGene00022532 | WBGene00006713 | WBGene00005663 |
| WBGene00018402 | WBGene00009333 | WBGene00003788 | WBGene00001974 | WBGene00022580 | WBGene00006720 | WBGene00006066 |
| WBGene00018425 | WBGene00009343 | WBGene00003790 | WBGene00001976 | WBGene00022583 | WBGene00006728 | WBGene00006220 |
| WBGene00018472 | WBGene00009356 | WBGene00003794 | WBGene00001977 | WBGene00022749 | WBGene00006733 | WBGene00006342 |
| WBGene00018514 | WBGene00009408 | WBGene00003797 | WBGene00001978 | WBGene00022750 | WBGene00006734 | WBGene00006352 |
| WBGene00018565 | WBGene00009442 | WBGene00003798 | WBGene00001980 | WBGene00022855 | WBGene00006760 | WBGene00006353 |
| WBGene00018571 | WBGene00009449 | WBGene00003802 | WBGene00001983 | WBGene00023418 | WBGene00006768 | WBGene00006366 |
| WBGene00018576 | WBGene00009466 | WBGene00003806 | WBGene00001996 | WBGene00044191 | WBGene00006780 | WBGene00006369 |
| WBGene00018607 | WBGene00009470 | WBGene00003812 | WBGene00002002 | WBGene00044326 | WBGene00006791 | WBGene00006373 |
| WBGene00018638 | WBGene00009475 | WBGene00003815 | WBGene00002003 | WBGene00044636 | WBGene00006796 | WBGene00006407 |
| WBGene00018740 | WBGene00009479 | WBGene00003846 | WBGene00002011 | WBGene00045272 | WBGene00006817 | WBGene00006439 |
| WBGene00018745 | WBGene00009496 | WBGene00003861 | WBGene00002016 | WBGene00045418 | WBGene00006870 | WBGene00006444 |
| WBGene00018748 | WBGene00009502 | WBGene00003870 | WBGene00002023 | WBGene00045475 | WBGene00006888 | WBGene00006462 |
| WBGene00018750 | WBGene00009503 | WBGene00003882 | WBGene00002027 | WBGene00077690 | WBGene00006912 | WBGene00006472 |
| WBGene00018789 | WBGene00009533 | WBGene00003898 | WBGene00002036 | | WBGene00006913 | WBGene00006483 |
| WBGene00018842 | WBGene00009543 | WBGene00003921 | WBGene00002044 | | WBGene00006918 | WBGene00006518 |
| WBGene00018844 | WBGene00009547 | WBGene00003932 | WBGene00002047 | | WBGene00006947 | WBGene00006523 |
| WBGene00018987 | WBGene00009588 | WBGene00003958 | WBGene00002050 | | WBGene00006975 | WBGene00006528 |
| WBGene00019047 | WBGene00009595 | WBGene00003992 | WBGene00002054 | | WBGene00006997 | WBGene00006529 |
| WBGene00019070 | WBGene00009606 | WBGene00003994 | WBGene00002055 | | WBGene00007000 | WBGene00006536 |
| WBGene00019109 | WBGene00009627 | WBGene00004016 | WBGene00002060 | | WBGene00007009 | WBGene00006537 |
| WBGene00019265 | WBGene00009639 | WBGene00004020 | WBGene00002064 | | WBGene00007024 | WBGene00006591 |
| WBGene00019297 | WBGene00009673 | WBGene00004024 | WBGene00002068 | | WBGene00007042 | WBGene00006592 |
| WBGene00019331 | WBGene00009680 | WBGene00004031 | WBGene00002073 | | WBGene00007048 | WBGene00006593 |
| WBGene00019408 | WBGene00009691 | WBGene00004036 | WBGene00002075 | | WBGene00007099 | WBGene00006595 |
| WBGene00019507 | WBGene00009730 | WBGene00004048 | WBGene00002076 | | WBGene00007110 | WBGene00006599 |
| WBGene00019578 | WBGene00009735 | WBGene00004068 | WBGene00002079 | | WBGene00007135 | WBGene00006603 |
| WBGene00019630 | WBGene00009750 | WBGene00004087 | WBGene00002113 | | WBGene00007195 | WBGene00006626 |
| WBGene00019669 | WBGene00009806 | WBGene00004088 | WBGene00002116 | | WBGene00007223 | WBGene00006686 |
| WBGene00019754 | WBGene00009840 | WBGene00004093 | WBGene00002130 | | WBGene00007257 | WBGene00006699 |
| WBGene00019832 | WBGene00009848 | WBGene00004094 | WBGene00002147 | | WBGene00007350 | WBGene00006702 |
| WBGene00019843 | WBGene00009851 | WBGene00004106 | WBGene00002148 | | WBGene00007355 | WBGene00006725 |
| WBGene00019846 | WBGene00009863 | WBGene00004111 | WBGene00002169 | | WBGene00007385 | WBGene00006727 |
| WBGene00019858 | WBGene00009865 | WBGene00004122 | WBGene00002175 | | WBGene00007402 | WBGene00006739 |
| WBGene00019925 | WBGene00009889 | WBGene00004123 | WBGene00002180 | | WBGene00007428 | WBGene00006769 |
| WBGene00019967 | WBGene00009938 | WBGene00004126 | WBGene00002183 | | WBGene00007505 | WBGene00006771 |
| WBGene00019980 | WBGene00009946 | WBGene00004144 | WBGene00002187 | | WBGene00007520 | WBGene00006773 |

| | | | | | |
|---|---|---|---|---|---|
| WBGene00020116 | WBGene00009953 | WBGene00004145 | WBGene00002190 | WBGene00007534 | WBGene00006779 |
| WBGene00020135 | WBGene00009963 | WBGene00004173 | WBGene00002198 | WBGene00007549 | WBGene00006786 |
| WBGene00020217 | WBGene00009964 | WBGene00004180 | WBGene00002210 | WBGene00007553 | WBGene00006787 |
| WBGene00020231 | WBGene00009969 | WBGene00004190 | WBGene00002219 | WBGene00007555 | WBGene00006803 |
| WBGene00020276 | WBGene00010009 | WBGene00004198 | WBGene00002222 | WBGene00007588 | WBGene00006815 |
| WBGene00020313 | WBGene00010011 | WBGene00004202 | WBGene00002226 | WBGene00007646 | WBGene00006833 |
| WBGene00020411 | WBGene00010016 | WBGene00004203 | WBGene00002228 | WBGene00007696 | WBGene00006840 |
| WBGene00020460 | WBGene00010062 | WBGene00004206 | WBGene00002229 | WBGene00007698 | WBGene00006861 |
| WBGene00020478 | WBGene00010083 | WBGene00004207 | WBGene00002231 | WBGene00007746 | WBGene00006876 |
| WBGene00020479 | WBGene00010084 | WBGene00004212 | WBGene00002241 | WBGene00007812 | WBGene00006882 |
| WBGene00020483 | WBGene00010086 | WBGene00004220 | WBGene00002244 | WBGene00007898 | WBGene00006890 |
| WBGene00020484 | WBGene00010093 | WBGene00004236 | WBGene00002247 | WBGene00007926 | WBGene00006910 |
| WBGene00020555 | WBGene00010114 | WBGene00004238 | WBGene00002249 | WBGene00007969 | WBGene00006911 |
| WBGene00020587 | WBGene00010136 | WBGene00004244 | WBGene00002251 | WBGene00008000 | WBGene00006914 |
| WBGene00020647 | WBGene00010149 | WBGene00004248 | WBGene00002253 | WBGene00008006 | WBGene00006917 |
| WBGene00020670 | WBGene00010150 | WBGene00004269 | WBGene00002254 | WBGene00008019 | WBGene00006919 |
| WBGene00020720 | WBGene00010175 | WBGene00004283 | WBGene00002255 | WBGene00008053 | WBGene00006920 |
| WBGene00020822 | WBGene00010191 | WBGene00004296 | WBGene00002257 | WBGene00008136 | WBGene00006921 |
| WBGene00020825 | WBGene00010230 | WBGene00004298 | WBGene00002259 | WBGene00008211 | WBGene00006923 |
| WBGene00020830 | WBGene00010231 | WBGene00004318 | WBGene00002261 | WBGene00008225 | WBGene00006924 |
| WBGene00020850 | WBGene00010242 | WBGene00004323 | WBGene00002263 | WBGene00008266 | WBGene00006934 |
| WBGene00020851 | WBGene00010305 | WBGene00004338 | WBGene00002267 | WBGene00008274 | WBGene00006941 |
| WBGene00020901 | WBGene00010325 | WBGene00004343 | WBGene00002268 | WBGene00008363 | WBGene00006950 |
| WBGene00020908 | WBGene00010350 | WBGene00004348 | WBGene00002269 | WBGene00008411 | WBGene00006956 |
| WBGene00020914 | WBGene00010374 | WBGene00004349 | WBGene00002271 | WBGene00008430 | WBGene00006959 |
| WBGene00020923 | WBGene00010466 | WBGene00004355 | WBGene00002272 | WBGene00008448 | WBGene00006996 |
| WBGene00020984 | WBGene00010475 | WBGene00004367 | WBGene00002273 | WBGene00008480 | WBGene00007012 |
| WBGene00021076 | WBGene00010507 | WBGene00004370 | WBGene00002275 | WBGene00008532 | WBGene00007016 |
| WBGene00021087 | WBGene00010509 | WBGene00004375 | WBGene00002280 | WBGene00008666 | WBGene00007030 |
| WBGene00021137 | WBGene00010510 | WBGene00004380 | WBGene00002297 | WBGene00008682 | WBGene00007053 |
| WBGene00021138 | WBGene00010516 | WBGene00004386 | WBGene00002348 | WBGene00008684 | WBGene00007058 |
| WBGene00021143 | WBGene00010528 | WBGene00004390 | WBGene00002694 | WBGene00008685 | WBGene00007071 |
| WBGene00021178 | WBGene00010538 | WBGene00004411 | WBGene00002696 | WBGene00008686 | WBGene00007137 |
| WBGene00021349 | WBGene00010542 | WBGene00004413 | WBGene00002855 | WBGene00008711 | WBGene00007142 |
| WBGene00021371 | WBGene00010564 | WBGene00004433 | WBGene00002957 | WBGene00008748 | WBGene00007143 |
| WBGene00021500 | WBGene00010569 | WBGene00004435 | WBGene00002978 | WBGene00008764 | WBGene00007150 |
| WBGene00021781 | WBGene00010573 | WBGene00004438 | WBGene00002979 | WBGene00008800 | WBGene00007178 |
| WBGene00021908 | WBGene00010592 | WBGene00004439 | WBGene00002980 | WBGene00008832 | WBGene00007227 |
| WBGene00022315 | WBGene00010632 | WBGene00004442 | WBGene00002986 | WBGene00008851 | WBGene00007264 |
| WBGene00022390 | WBGene00010633 | WBGene00004448 | WBGene00002994 | WBGene00008852 | WBGene00007270 |
| WBGene00022478 | WBGene00010672 | WBGene00004451 | WBGene00002998 | WBGene00008859 | WBGene00007275 |
| WBGene00022479 | WBGene00010676 | WBGene00004454 | WBGene00002999 | WBGene00008922 | WBGene00007352 |
| WBGene00022533 | WBGene00010698 | WBGene00004457 | WBGene00003010 | WBGene00008948 | WBGene00007365 |
| WBGene00022596 | WBGene00010717 | WBGene00004461 | WBGene00003021 | WBGene00008956 | WBGene00007384 |
| WBGene00022668 | WBGene00010734 | WBGene00004481 | WBGene00003036 | WBGene00008987 | WBGene00007409 |
| WBGene00022669 | WBGene00010737 | WBGene00004497 | WBGene00003047 | WBGene00008999 | WBGene00007449 |
| WBGene00022721 | WBGene00010811 | WBGene00004506 | WBGene00003062 | WBGene00009084 | WBGene00007463 |
| WBGene00022767 | WBGene00010812 | WBGene00004680 | WBGene00003065 | WBGene00009094 | WBGene00007474 |
| WBGene00022795 | WBGene00010835 | WBGene00004681 | WBGene00003066 | WBGene00009098 | WBGene00007492 |
| WBGene00022801 | WBGene00010850 | WBGene00004719 | WBGene00003076 | WBGene00009120 | WBGene00007514 |
| WBGene00022826 | WBGene00010904 | WBGene00004726 | WBGene00003079 | WBGene00009121 | WBGene00007545 |
| WBGene00022827 | WBGene00010934 | WBGene00004728 | WBGene00003080 | WBGene00009122 | WBGene00007586 |
| WBGene00022892 | WBGene00010984 | WBGene00004744 | WBGene00003087 | WBGene00009141 | WBGene00007651 |
| WBGene00023068 | WBGene00011019 | WBGene00004747 | WBGene00003090 | WBGene00009142 | WBGene00007679 |
| WBGene00023417 | WBGene00011021 | WBGene00004753 | WBGene00003091 | WBGene00009164 | WBGene00007736 |
| WBGene00044009 | WBGene00011028 | WBGene00004762 | WBGene00003096 | WBGene00009187 | WBGene00007779 |
| WBGene00044031 | WBGene00011076 | WBGene00004775 | WBGene00003097 | WBGene00009241 | WBGene00007836 |
| WBGene00044107 | WBGene00011077 | WBGene00004778 | WBGene00003102 | WBGene00009245 | WBGene00007841 |
| WBGene00044133 | WBGene00011139 | WBGene00004786 | WBGene00003119 | WBGene00009262 | WBGene00007848 |
| WBGene00044243 | WBGene00011147 | WBGene00004789 | WBGene00003123 | WBGene00009264 | WBGene00007877 |
| WBGene00044263 | WBGene00011154 | WBGene00004822 | WBGene00003129 | WBGene00009284 | WBGene00007903 |
| WBGene00044264 | WBGene00011172 | WBGene00004823 | WBGene00003142 | WBGene00009311 | WBGene00007934 |
| WBGene00044381 | WBGene00011217 | WBGene00004857 | WBGene00003149 | WBGene00009454 | WBGene00007947 |
| WBGene00044391 | WBGene00011261 | WBGene00004860 | WBGene00003150 | WBGene00009460 | WBGene00007999 |
| WBGene00044393 | WBGene00011271 | WBGene00004873 | WBGene00003153 | WBGene00009477 | WBGene00008032 |
| WBGene00044464 | WBGene00011274 | WBGene00004886 | WBGene00003155 | WBGene00009498 | WBGene00008089 |
| WBGene00044510 | WBGene00011291 | WBGene00004891 | WBGene00003157 | WBGene00009508 | WBGene00008117 |
| WBGene00044534 | WBGene00011308 | WBGene00004905 | WBGene00003158 | WBGene00009559 | WBGene00008205 |

| | | | | | |
|---|---|---|---|---|---|
| WBGene00044542 | WBGene00011416 | WBGene00004908 | WBGene00003159 | WBGene00009561 | WBGene00008215 |
| WBGene00044569 | WBGene00011418 | WBGene00004912 | WBGene00003160 | WBGene00009574 | WBGene00008224 |
| WBGene00044570 | WBGene00011419 | WBGene00004923 | WBGene00003164 | WBGene00009583 | WBGene00008233 |
| WBGene00044577 | WBGene00011429 | WBGene00004928 | WBGene00003175 | WBGene00009594 | WBGene00008270 |
| WBGene00044611 | WBGene00011432 | WBGene00004932 | WBGene00003182 | WBGene00009660 | WBGene00008362 |
| WBGene00044621 | WBGene00011439 | WBGene00004949 | WBGene00003183 | WBGene00009740 | WBGene00008404 |
| WBGene00044630 | WBGene00011442 | WBGene00004952 | WBGene00003184 | WBGene00009770 | WBGene00008419 |
| WBGene00044634 | WBGene00011469 | WBGene00004953 | WBGene00003185 | WBGene00009771 | WBGene00008505 |
| WBGene00044667 | WBGene00011484 | WBGene00004965 | WBGene00003196 | WBGene00009785 | WBGene00008506 |
| WBGene00044675 | WBGene00011487 | WBGene00004978 | WBGene00003209 | WBGene00009793 | WBGene00008521 |
| WBGene00044707 | WBGene00011494 | WBGene00004981 | WBGene00003210 | WBGene00009797 | WBGene00008549 |
| WBGene00044732 | WBGene00011495 | WBGene00004989 | WBGene00003221 | WBGene00009809 | WBGene00008622 |
| WBGene00044917 | WBGene00011499 | WBGene00005002 | WBGene00003222 | WBGene00009818 | WBGene00008652 |
| WBGene00045061 | WBGene00011500 | WBGene00005006 | WBGene00003225 | WBGene00009880 | WBGene00008664 |
| WBGene00045245 | WBGene00011508 | WBGene00005010 | WBGene00003228 | WBGene00009882 | WBGene00008683 |
| WBGene00045246 | WBGene00011533 | WBGene00005024 | WBGene00003229 | WBGene00009915 | WBGene00008688 |
| WBGene00045308 | WBGene00011606 | WBGene00005552 | WBGene00003230 | WBGene00009930 | WBGene00008707 |
| WBGene00045383 | WBGene00011610 | WBGene00005643 | WBGene00003231 | WBGene00009952 | WBGene00008743 |
| WBGene00050968 | WBGene00011624 | WBGene00005647 | WBGene00003235 | WBGene00009978 | WBGene00008803 |
| WBGene00077453 | WBGene00011642 | WBGene00005655 | WBGene00003239 | WBGene00009983 | WBGene00008857 |
| WBGene00077489 | WBGene00011663 | WBGene00005662 | WBGene00003242 | WBGene00009987 | WBGene00008920 |
| WBGene00077585 | WBGene00011669 | WBGene00006047 | WBGene00003243 | WBGene00009993 | WBGene00008926 |
| WBGene00077696 | WBGene00011692 | WBGene00006052 | WBGene00003247 | WBGene00009996 | WBGene00008940 |
| WBGene00077704 | WBGene00011707 | WBGene00006053 | WBGene00003254 | WBGene00010035 | WBGene00008964 |
| WBGene00077786 | WBGene00011727 | WBGene00006062 | WBGene00003368 | WBGene00010055 | WBGene00008975 |
| WBGene00189933 | WBGene00011731 | WBGene00006063 | WBGene00003369 | WBGene00010111 | WBGene00009050 |
| WBGene00194704 | WBGene00011733 | WBGene00006306 | WBGene00003370 | WBGene00010199 | WBGene00009081 |
| WBGene00194708 | WBGene00011739 | WBGene00006349 | WBGene00003371 | WBGene00010225 | WBGene00009082 |
| WBGene00194733 | WBGene00011744 | WBGene00006359 | WBGene00003375 | WBGene00010262 | WBGene00009112 |
| WBGene00194734 | WBGene00011745 | WBGene00006368 | WBGene00003389 | WBGene00010280 | WBGene00009119 |
| WBGene00194835 | WBGene00011759 | WBGene00006371 | WBGene00003392 | WBGene00010284 | WBGene00009126 |
| WBGene00194867 | WBGene00011793 | WBGene00006372 | WBGene00003393 | WBGene00010303 | WBGene00009158 |
| WBGene00195146 | WBGene00011795 | WBGene00006375 | WBGene00003395 | WBGene00010321 | WBGene00009177 |
| WBGene00195208 | WBGene00011819 | WBGene00006382 | WBGene00003401 | WBGene00010333 | WBGene00009218 |
| WBGene00195212 | WBGene00011838 | WBGene00006387 | WBGene00003402 | WBGene00010337 | WBGene00009306 |
| WBGene00206479 | WBGene00011854 | WBGene00006391 | WBGene00003404 | WBGene00010407 | WBGene00009334 |
| WBGene00206487 | WBGene00011860 | WBGene00006402 | WBGene00003406 | WBGene00010449 | WBGene00009340 |
| WBGene00206516 | WBGene00011906 | WBGene00006408 | WBGene00003410 | WBGene00010453 | WBGene00009342 |
| WBGene00235095 | WBGene00011914 | WBGene00006416 | WBGene00003415 | WBGene00010470 | WBGene00009349 |
| WBGene00235158 | WBGene00011927 | WBGene00006445 | WBGene00003417 | WBGene00010476 | WBGene00009367 |
| WBGene00235332 | WBGene00011936 | WBGene00006448 | WBGene00003423 | WBGene00010479 | WBGene00009405 |
| WBGene00269421 | WBGene00012014 | WBGene00006463 | WBGene00003426 | WBGene00010480 | WBGene00009450 |
| WBGene00271777 | WBGene00012016 | WBGene00006481 | WBGene00003435 | WBGene00010550 | WBGene00009453 |
| WBGene00271798 | WBGene00012040 | WBGene00006492 | WBGene00003444 | WBGene00010557 | WBGene00009482 |
| WBGene00271822 | WBGene00012059 | WBGene00006498 | WBGene00003452 | WBGene00010560 | WBGene00009532 |
| WBGene00302974 | WBGene00012095 | WBGene00006499 | WBGene00003457 | WBGene00010579 | WBGene00009542 |
| WBGene00302980 | WBGene00012157 | WBGene00006503 | WBGene00003463 | WBGene00010595 | WBGene00009552 |
| WBGene00302993 | WBGene00012164 | WBGene00006512 | WBGene00003465 | WBGene00010629 | WBGene00009584 |
| WBGene00303000 | WBGene00012282 | WBGene00006513 | WBGene00003467 | WBGene00010631 | WBGene00009587 |
| WBGene00303011 | WBGene00012293 | WBGene00006515 | WBGene00003469 | WBGene00010639 | WBGene00009621 |
| WBGene00303048 | WBGene00012307 | WBGene00006524 | WBGene00003473 | WBGene00010640 | WBGene00009628 |
| WBGene00303057 | WBGene00012312 | WBGene00006541 | WBGene00003495 | WBGene00010641 | WBGene00009661 |
| WBGene00303079 | WBGene00012357 | WBGene00006562 | WBGene00003508 | WBGene00010681 | WBGene00009678 |
| WBGene00303088 | WBGene00012363 | WBGene00006567 | WBGene00003509 | WBGene00010699 | WBGene00009712 |
| WBGene00303093 | WBGene00012371 | WBGene00006582 | WBGene00003510 | WBGene00010736 | WBGene00009717 |
| WBGene00303105 | WBGene00012373 | WBGene00006586 | WBGene00003514 | WBGene00010740 | WBGene00009723 |
| WBGene00303423 | WBGene00012396 | WBGene00006590 | WBGene00003515 | WBGene00010780 | WBGene00009724 |
| WBGene00304794 | WBGene00012407 | WBGene00006635 | WBGene00003565 | WBGene00010785 | WBGene00009772 |
| WBGene00304808 | WBGene00012412 | WBGene00006648 | WBGene00003567 | WBGene00010790 | WBGene00009783 |
| WBGene00304824 | WBGene00012420 | WBGene00006652 | WBGene00003576 | WBGene00010796 | WBGene00009787 |
| WBGene00304992 | WBGene00012424 | WBGene00006662 | WBGene00003582 | WBGene00010828 | WBGene00009812 |
| WBGene00305157 | WBGene00012457 | WBGene00006671 | WBGene00003587 | WBGene00010840 | WBGene00009813 |
| WBGene00305173 | WBGene00012459 | WBGene00006672 | WBGene00003591 | WBGene00010878 | WBGene00009829 |
| WBGene00016067 | WBGene00012462 | WBGene00006675 | WBGene00003599 | WBGene00010882 | WBGene00009918 |
| WBGene00001588 | WBGene00012489 | WBGene00006705 | WBGene00003604 | WBGene00010943 | WBGene00009926 |
| WBGene00011305 | WBGene00012605 | WBGene00006707 | WBGene00003607 | WBGene00010945 | WBGene00009947 |
| | WBGene00012628 | WBGene00006709 | WBGene00003620 | WBGene00010993 | WBGene00009980 |
| | WBGene00012634 | WBGene00006710 | WBGene00003648 | WBGene00011033 | WBGene00009982 |

| | | | | |
|---|---|---|---|---|
| WBGene00012635 | WBGene00006714 | WBGene00003693 | WBGene00011036 | WBGene00010037 |
| WBGene00012640 | WBGene00006724 | WBGene00003721 | WBGene00011050 | WBGene00010052 |
| WBGene00012703 | WBGene00006729 | WBGene00003739 | WBGene00011105 | WBGene00010135 |
| WBGene00012728 | WBGene00006735 | WBGene00003741 | WBGene00011111 | WBGene00010221 |
| WBGene00012755 | WBGene00006741 | WBGene00003744 | WBGene00011122 | WBGene00010260 |
| WBGene00012776 | WBGene00006745 | WBGene00003745 | WBGene00011227 | WBGene00010279 |
| WBGene00012781 | WBGene00006747 | WBGene00003749 | WBGene00011240 | WBGene00010291 |
| WBGene00012795 | WBGene00006749 | WBGene00003750 | WBGene00011259 | WBGene00010317 |
| WBGene00012889 | WBGene00006753 | WBGene00003751 | WBGene00011298 | WBGene00010326 |
| WBGene00012901 | WBGene00006767 | WBGene00003752 | WBGene00011311 | WBGene00010419 |
| WBGene00012908 | WBGene00006795 | WBGene00003753 | WBGene00011312 | WBGene00010425 |
| WBGene00012960 | WBGene00006807 | WBGene00003756 | WBGene00011333 | WBGene00010454 |
| WBGene00013017 | WBGene00006810 | WBGene00003759 | WBGene00011408 | WBGene00010456 |
| WBGene00013030 | WBGene00006816 | WBGene00003766 | WBGene00011409 | WBGene00010478 |
| WBGene00013035 | WBGene00006825 | WBGene00003769 | WBGene00011488 | WBGene00010502 |
| WBGene00013039 | WBGene00006826 | WBGene00003777 | WBGene00011501 | WBGene00010503 |
| WBGene00013078 | WBGene00006836 | WBGene00003784 | WBGene00011526 | WBGene00010556 |
| WBGene00013079 | WBGene00006843 | WBGene00003786 | WBGene00011528 | WBGene00010621 |
| WBGene00013109 | WBGene00006856 | WBGene00003787 | WBGene00011538 | WBGene00010627 |
| WBGene00013124 | WBGene00006896 | WBGene00003791 | WBGene00011564 | WBGene00010661 |
| WBGene00013130 | WBGene00006922 | WBGene00003792 | WBGene00011580 | WBGene00010664 |
| WBGene00013135 | WBGene00006940 | WBGene00003799 | WBGene00011629 | WBGene00010665 |
| WBGene00013141 | WBGene00006943 | WBGene00003800 | WBGene00011639 | WBGene00010667 |
| WBGene00013142 | WBGene00006944 | WBGene00003801 | WBGene00011696 | WBGene00010677 |
| WBGene00013171 | WBGene00006945 | WBGene00003803 | WBGene00011713 | WBGene00010697 |
| WBGene00013212 | WBGene00006963 | WBGene00003804 | WBGene00011802 | WBGene00010700 |
| WBGene00013218 | WBGene00006968 | WBGene00003805 | WBGene00011815 | WBGene00010755 |
| WBGene00013220 | WBGene00006970 | WBGene00003813 | WBGene00011833 | WBGene00010759 |
| WBGene00013227 | WBGene00006974 | WBGene00003826 | WBGene00011835 | WBGene00010809 |
| WBGene00013272 | WBGene00007001 | WBGene00003828 | WBGene00011887 | WBGene00010834 |
| WBGene00013290 | WBGene00007019 | WBGene00003829 | WBGene00011890 | WBGene00010867 |
| WBGene00013299 | WBGene00007020 | WBGene00003835 | WBGene00011932 | WBGene00010868 |
| WBGene00013329 | WBGene00007044 | WBGene00003836 | WBGene00011962 | WBGene00010889 |
| WBGene00013333 | WBGene00007063 | WBGene00003860 | WBGene00011965 | WBGene00010941 |
| WBGene00013365 | WBGene00007068 | WBGene00003864 | WBGene00011970 | WBGene00010988 |
| WBGene00013479 | WBGene00007087 | WBGene00003879 | WBGene00011971 | WBGene00011042 |
| WBGene00013553 | WBGene00007093 | WBGene00003893 | WBGene00011978 | WBGene00011059 |
| WBGene00013588 | WBGene00007101 | WBGene00003901 | WBGene00011985 | WBGene00011110 |
| WBGene00013637 | WBGene00007113 | WBGene00003905 | WBGene00011994 | WBGene00011116 |
| WBGene00013673 | WBGene00007118 | WBGene00003907 | WBGene00012031 | WBGene00011128 |
| WBGene00013678 | WBGene00007130 | WBGene00003912 | WBGene00012094 | WBGene00011146 |
| WBGene00013693 | WBGene00007153 | WBGene00003917 | WBGene00012097 | WBGene00011230 |
| WBGene00013746 | WBGene00007168 | WBGene00003919 | WBGene00012121 | WBGene00011232 |
| WBGene00013754 | WBGene00007171 | WBGene00003923 | WBGene00012123 | WBGene00011235 |
| WBGene00013819 | WBGene00007185 | WBGene00003924 | WBGene00012136 | WBGene00011263 |
| WBGene00013845 | WBGene00007188 | WBGene00003925 | WBGene00012144 | WBGene00011273 |
| WBGene00013852 | WBGene00007190 | WBGene00003926 | WBGene00012148 | WBGene00011278 |
| WBGene00013856 | WBGene00007221 | WBGene00003927 | WBGene00012167 | WBGene00011300 |
| WBGene00013877 | WBGene00007225 | WBGene00003930 | WBGene00012179 | WBGene00011304 |
| WBGene00013893 | WBGene00007234 | WBGene00003931 | WBGene00012184 | WBGene00011393 |
| WBGene00013958 | WBGene00007260 | WBGene00003933 | WBGene00012193 | WBGene00011398 |
| WBGene00013960 | WBGene00007282 | WBGene00003934 | WBGene00012194 | WBGene00011404 |
| WBGene00013968 | WBGene00007283 | WBGene00003941 | WBGene00012203 | WBGene00011436 |
| WBGene00013975 | WBGene00007302 | WBGene00003948 | WBGene00012230 | WBGene00011480 |
| WBGene00013984 | WBGene00007308 | WBGene00003952 | WBGene00012243 | WBGene00011481 |
| WBGene00013989 | WBGene00007312 | WBGene00003955 | WBGene00012245 | WBGene00011522 |
| WBGene00014002 | WBGene00007315 | WBGene00003956 | WBGene00012339 | WBGene00011529 |
| WBGene00014034 | WBGene00007331 | WBGene00003966 | WBGene00012344 | WBGene00011662 |
| WBGene00014055 | WBGene00007339 | WBGene00003967 | WBGene00012354 | WBGene00011679 |
| WBGene00014066 | WBGene00007347 | WBGene00003970 | WBGene00012359 | WBGene00011729 |
| WBGene00014074 | WBGene00007349 | WBGene00003978 | WBGene00012360 | WBGene00011737 |
| WBGene00014182 | WBGene00007386 | WBGene00003980 | WBGene00012444 | WBGene00011760 |
| WBGene00014183 | WBGene00007387 | WBGene00003986 | WBGene00012497 | WBGene00011763 |
| WBGene00014184 | WBGene00007414 | WBGene00003989 | WBGene00012538 | WBGene00011768 |
| WBGene00014188 | WBGene00007450 | WBGene00003990 | WBGene00012591 | WBGene00011773 |
| WBGene00014203 | WBGene00007455 | WBGene00003996 | WBGene00012608 | WBGene00011775 |
| WBGene00014240 | WBGene00007480 | WBGene00004013 | WBGene00012704 | WBGene00011820 |
| WBGene00014697 | WBGene00007486 | WBGene00004027 | WBGene00012803 | WBGene00011831 |

| | | | | |
|---|---|---|---|---|
| WBGene00014938 | WBGene00007493 | WBGene00004028 | WBGene00012829 | WBGene00011849 |
| WBGene00015001 | WBGene00007496 | WBGene00004032 | WBGene00012869 | WBGene00011850 |
| WBGene00015014 | WBGene00007499 | WBGene00004042 | WBGene00012896 | WBGene00011867 |
| WBGene00015024 | WBGene00007508 | WBGene00004044 | WBGene00012903 | WBGene00011883 |
| WBGene00015050 | WBGene00007517 | WBGene00004046 | WBGene00012929 | WBGene00011884 |
| WBGene00015067 | WBGene00007521 | WBGene00004047 | WBGene00012943 | WBGene00011891 |
| WBGene00015105 | WBGene00007574 | WBGene00004049 | WBGene00013103 | WBGene00011977 |
| WBGene00015145 | WBGene00007579 | WBGene00004050 | WBGene00013127 | WBGene00012033 |
| WBGene00015157 | WBGene00007580 | WBGene00004051 | WBGene00013143 | WBGene00012085 |
| WBGene00015158 | WBGene00007584 | WBGene00004055 | WBGene00013203 | WBGene00012107 |
| WBGene00015204 | WBGene00007590 | WBGene00004056 | WBGene00013219 | WBGene00012118 |
| WBGene00015218 | WBGene00007606 | WBGene00004060 | WBGene00013236 | WBGene00012149 |
| WBGene00015229 | WBGene00007616 | WBGene00004062 | WBGene00013263 | WBGene00012186 |
| WBGene00015262 | WBGene00007622 | WBGene00004063 | WBGene00013379 | WBGene00012256 |
| WBGene00015271 | WBGene00007637 | WBGene00004064 | WBGene00013497 | WBGene00012261 |
| WBGene00015322 | WBGene00007640 | WBGene00004078 | WBGene00013597 | WBGene00012317 |
| WBGene00015327 | WBGene00007649 | WBGene00004083 | WBGene00013734 | WBGene00012343 |
| WBGene00015339 | WBGene00007652 | WBGene00004085 | WBGene00013892 | WBGene00012376 |
| WBGene00015345 | WBGene00007664 | WBGene00004086 | WBGene00013895 | WBGene00012443 |
| WBGene00015352 | WBGene00007681 | WBGene00004089 | WBGene00014000 | WBGene00012463 |
| WBGene00015369 | WBGene00007690 | WBGene00004096 | WBGene00014013 | WBGene00012471 |
| WBGene00015392 | WBGene00007697 | WBGene00004101 | WBGene00014016 | WBGene00012546 |
| WBGene00015393 | WBGene00007702 | WBGene00004105 | WBGene00014096 | WBGene00012553 |
| WBGene00015433 | WBGene00007703 | WBGene00004112 | WBGene00014109 | WBGene00012717 |
| WBGene00015457 | WBGene00007705 | WBGene00004113 | WBGene00014111 | WBGene00012762 |
| WBGene00015471 | WBGene00007751 | WBGene00004116 | WBGene00014151 | WBGene00012768 |
| WBGene00015487 | WBGene00007752 | WBGene00004117 | WBGene00014164 | WBGene00012769 |
| WBGene00015517 | WBGene00007757 | WBGene00004121 | WBGene00014165 | WBGene00012999 |
| WBGene00015527 | WBGene00007771 | WBGene00004125 | WBGene00014219 | WBGene00013025 |
| WBGene00015539 | WBGene00007780 | WBGene00004128 | WBGene00014244 | WBGene00013034 |
| WBGene00015568 | WBGene00007785 | WBGene00004130 | WBGene00015074 | WBGene00013106 |
| WBGene00015574 | WBGene00007797 | WBGene00004135 | WBGene00015125 | WBGene00013139 |
| WBGene00015593 | WBGene00007807 | WBGene00004138 | WBGene00015155 | WBGene00013311 |
| WBGene00015605 | WBGene00007811 | WBGene00004148 | WBGene00015164 | WBGene00013352 |
| WBGene00015619 | WBGene00007837 | WBGene00004154 | WBGene00015168 | WBGene00013461 |
| WBGene00015632 | WBGene00007857 | WBGene00004156 | WBGene00015172 | WBGene00013462 |
| WBGene00015634 | WBGene00007859 | WBGene00004161 | WBGene00015197 | WBGene00013463 |
| WBGene00015642 | WBGene00007860 | WBGene00004166 | WBGene00015298 | WBGene00013489 |
| WBGene00015648 | WBGene00007892 | WBGene00004178 | WBGene00015329 | WBGene00013560 |
| WBGene00015679 | WBGene00007912 | WBGene00004181 | WBGene00015330 | WBGene00013575 |
| WBGene00015688 | WBGene00007914 | WBGene00004186 | WBGene00015344 | WBGene00013578 |
| WBGene00015729 | WBGene00007918 | WBGene00004188 | WBGene00015349 | WBGene00013591 |
| WBGene00015765 | WBGene00007922 | WBGene00004194 | WBGene00015391 | WBGene00013672 |
| WBGene00015790 | WBGene00007924 | WBGene00004199 | WBGene00015418 | WBGene00013709 |
| WBGene00015793 | WBGene00007956 | WBGene00004201 | WBGene00015460 | WBGene00013736 |
| WBGene00015843 | WBGene00007965 | WBGene00004213 | WBGene00015463 | WBGene00013766 |
| WBGene00015885 | WBGene00007971 | WBGene00004214 | WBGene00015481 | WBGene00013859 |
| WBGene00015897 | WBGene00007972 | WBGene00004215 | WBGene00015485 | WBGene00013866 |
| WBGene00015899 | WBGene00007976 | WBGene00004239 | WBGene00015660 | WBGene00013878 |
| WBGene00015910 | WBGene00007987 | WBGene00004241 | WBGene00015703 | WBGene00013924 |
| WBGene00015911 | WBGene00008002 | WBGene00004242 | WBGene00015734 | WBGene00013926 |
| WBGene00015929 | WBGene00008005 | WBGene00004243 | WBGene00015778 | WBGene00013969 |
| WBGene00015947 | WBGene00008018 | WBGene00004266 | WBGene00015791 | WBGene00013985 |
| WBGene00015972 | WBGene00008027 | WBGene00004267 | WBGene00015794 | WBGene00014012 |
| WBGene00016003 | WBGene00008031 | WBGene00004268 | WBGene00015810 | WBGene00014025 |
| WBGene00016009 | WBGene00008040 | WBGene00004270 | WBGene00015811 | WBGene00014058 |
| WBGene00016028 | WBGene00008080 | WBGene00004273 | WBGene00015813 | WBGene00014095 |
| WBGene00016038 | WBGene00008081 | WBGene00004276 | WBGene00015814 | WBGene00014098 |
| WBGene00016070 | WBGene00008119 | WBGene00004277 | WBGene00015920 | WBGene00014115 |
| WBGene00016108 | WBGene00008127 | WBGene00004282 | WBGene00015941 | WBGene00014148 |
| WBGene00016145 | WBGene00008139 | WBGene00004284 | WBGene00016002 | WBGene00014172 |
| WBGene00016153 | WBGene00008143 | WBGene00004286 | WBGene00016015 | WBGene00014173 |
| WBGene00016160 | WBGene00008145 | WBGene00004302 | WBGene00016019 | WBGene00014202 |
| WBGene00016177 | WBGene00008167 | WBGene00004303 | WBGene00016045 | WBGene00014300 |
| WBGene00016209 | WBGene00008169 | WBGene00004305 | WBGene00016059 | WBGene00015002 |
| WBGene00016218 | WBGene00008172 | WBGene00004307 | WBGene00016140 | WBGene00015007 |
| WBGene00016221 | WBGene00008173 | WBGene00004308 | WBGene00016146 | WBGene00015046 |
| WBGene00016231 | WBGene00008213 | WBGene00004314 | WBGene00016167 | WBGene00015056 |

| | | | | |
|---|---|---|---|---|
| WBGene00016245 | WBGene00008214 | WBGene00004315 | WBGene00016170 | WBGene00015064 |
| WBGene00016248 | WBGene00008228 | WBGene00004316 | WBGene00016173 | WBGene00015143 |
| WBGene00016295 | WBGene00008238 | WBGene00004317 | WBGene00016200 | WBGene00015156 |
| WBGene00016379 | WBGene00008254 | WBGene00004320 | WBGene00016235 | WBGene00015163 |
| WBGene00016383 | WBGene00008281 | WBGene00004322 | WBGene00016250 | WBGene00015207 |
| WBGene00016404 | WBGene00008298 | WBGene00004333 | WBGene00016258 | WBGene00015285 |
| WBGene00016423 | WBGene00008317 | WBGene00004337 | WBGene00016261 | WBGene00015294 |
| WBGene00016427 | WBGene00008318 | WBGene00004340 | WBGene00016310 | WBGene00015332 |
| WBGene00016429 | WBGene00008334 | WBGene00004341 | WBGene00016318 | WBGene00015410 |
| WBGene00016449 | WBGene00008338 | WBGene00004345 | WBGene00016326 | WBGene00015464 |
| WBGene00016456 | WBGene00008341 | WBGene00004350 | WBGene00016354 | WBGene00015512 |
| WBGene00016463 | WBGene00008346 | WBGene00004356 | WBGene00016405 | WBGene00015514 |
| WBGene00016490 | WBGene00008348 | WBGene00004357 | WBGene00016439 | WBGene00015545 |
| WBGene00016504 | WBGene00008354 | WBGene00004361 | WBGene00016452 | WBGene00015547 |
| WBGene00016545 | WBGene00008358 | WBGene00004364 | WBGene00016489 | WBGene00015676 |
| WBGene00016561 | WBGene00008361 | WBGene00004374 | WBGene00016494 | WBGene00015802 |
| WBGene00016566 | WBGene00008369 | WBGene00004377 | WBGene00016716 | WBGene00015809 |
| WBGene00016567 | WBGene00008385 | WBGene00004381 | WBGene00016735 | WBGene00015943 |
| WBGene00016590 | WBGene00008387 | WBGene00004382 | WBGene00016803 | WBGene00016006 |
| WBGene00016598 | WBGene00008393 | WBGene00004383 | WBGene00016955 | WBGene00016011 |
| WBGene00016616 | WBGene00008398 | WBGene00004384 | WBGene00016989 | WBGene00016018 |
| WBGene00016642 | WBGene00008399 | WBGene00004385 | WBGene00016995 | WBGene00016103 |
| WBGene00016664 | WBGene00008400 | WBGene00004388 | WBGene00017004 | WBGene00016133 |
| WBGene00016701 | WBGene00008420 | WBGene00004389 | WBGene00017075 | WBGene00016195 |
| WBGene00016752 | WBGene00008422 | WBGene00004391 | WBGene00017119 | WBGene00016201 |
| WBGene00016760 | WBGene00008428 | WBGene00004392 | WBGene00017163 | WBGene00016264 |
| WBGene00016778 | WBGene00008438 | WBGene00004394 | WBGene00017244 | WBGene00016355 |
| WBGene00016782 | WBGene00008444 | WBGene00004403 | WBGene00017274 | WBGene00016415 |
| WBGene00016817 | WBGene00008456 | WBGene00004405 | WBGene00017311 | WBGene00016417 |
| WBGene00016865 | WBGene00008457 | WBGene00004460 | WBGene00017372 | WBGene00016419 |
| WBGene00016885 | WBGene00008476 | WBGene00004466 | WBGene00017419 | WBGene00016422 |
| WBGene00016892 | WBGene00008477 | WBGene00004468 | WBGene00017431 | WBGene00016493 |
| WBGene00016900 | WBGene00008490 | WBGene00004505 | WBGene00017480 | WBGene00016495 |
| WBGene00016903 | WBGene00008502 | WBGene00004507 | WBGene00017563 | WBGene00016496 |
| WBGene00016911 | WBGene00008503 | WBGene00004508 | WBGene00017571 | WBGene00016499 |
| WBGene00016940 | WBGene00008504 | WBGene00004510 | WBGene00017606 | WBGene00016505 |
| WBGene00016947 | WBGene00008515 | WBGene00004679 | WBGene00017648 | WBGene00016509 |
| WBGene00016953 | WBGene00008522 | WBGene00004684 | WBGene00017717 | WBGene00016596 |
| WBGene00016964 | WBGene00008531 | WBGene00004700 | WBGene00017759 | WBGene00016630 |
| WBGene00016965 | WBGene00008533 | WBGene00004702 | WBGene00017760 | WBGene00016653 |
| WBGene00016978 | WBGene00008541 | WBGene00004705 | WBGene00017773 | WBGene00016739 |
| WBGene00017002 | WBGene00008547 | WBGene00004706 | WBGene00017785 | WBGene00016749 |
| WBGene00017017 | WBGene00008554 | WBGene00004723 | WBGene00017786 | WBGene00016750 |
| WBGene00017042 | WBGene00008561 | WBGene00004727 | WBGene00017816 | WBGene00016790 |
| WBGene00017068 | WBGene00008578 | WBGene00004729 | WBGene00017852 | WBGene00016794 |
| WBGene00017071 | WBGene00008580 | WBGene00004732 | WBGene00017855 | WBGene00016907 |
| WBGene00017073 | WBGene00008584 | WBGene00004735 | WBGene00017891 | WBGene00016987 |
| WBGene00017077 | WBGene00008585 | WBGene00004737 | WBGene00017920 | WBGene00016997 |
| WBGene00017078 | WBGene00008586 | WBGene00004743 | WBGene00017928 | WBGene00017012 |
| WBGene00017112 | WBGene00008588 | WBGene00004749 | WBGene00017964 | WBGene00017088 |
| WBGene00017134 | WBGene00008624 | WBGene00004752 | WBGene00017970 | WBGene00017121 |
| WBGene00017143 | WBGene00008648 | WBGene00004754 | WBGene00018008 | WBGene00017125 |
| WBGene00017169 | WBGene00008665 | WBGene00004758 | WBGene00018051 | WBGene00017126 |
| WBGene00017193 | WBGene00008690 | WBGene00004765 | WBGene00018132 | WBGene00017166 |
| WBGene00017218 | WBGene00008691 | WBGene00004766 | WBGene00018151 | WBGene00017217 |
| WBGene00017265 | WBGene00008710 | WBGene00004774 | WBGene00018152 | WBGene00017220 |
| WBGene00017279 | WBGene00008713 | WBGene00004782 | WBGene00018208 | WBGene00017262 |
| WBGene00017289 | WBGene00008733 | WBGene00004787 | WBGene00018218 | WBGene00017385 |
| WBGene00017297 | WBGene00008749 | WBGene00004795 | WBGene00018250 | WBGene00017430 |
| WBGene00017310 | WBGene00008775 | WBGene00004798 | WBGene00018285 | WBGene00017438 |
| WBGene00017312 | WBGene00008805 | WBGene00004800 | WBGene00018316 | WBGene00017463 |
| WBGene00017327 | WBGene00008860 | WBGene00004803 | WBGene00018341 | WBGene00017483 |
| WBGene00017348 | WBGene00008868 | WBGene00004806 | WBGene00018421 | WBGene00017506 |
| WBGene00017381 | WBGene00008885 | WBGene00004808 | WBGene00018426 | WBGene00017565 |
| WBGene00017384 | WBGene00008890 | WBGene00004809 | WBGene00018475 | WBGene00017569 |
| WBGene00017413 | WBGene00008918 | WBGene00004810 | WBGene00018491 | WBGene00017698 |
| WBGene00017420 | WBGene00008924 | WBGene00004825 | WBGene00018509 | WBGene00017765 |
| WBGene00017426 | WBGene00008961 | WBGene00004874 | WBGene00018510 | WBGene00017772 |

| | | | | |
|---|---|---|---|---|
| WBGene00017427 | WBGene00008974 | WBGene00004875 | WBGene00018522 | WBGene00017789 |
| WBGene00017448 | WBGene00008979 | WBGene00004879 | WBGene00018533 | WBGene00017838 |
| WBGene00017490 | WBGene00008985 | WBGene00004880 | WBGene00018562 | WBGene00017841 |
| WBGene00017498 | WBGene00008990 | WBGene00004881 | WBGene00018572 | WBGene00017864 |
| WBGene00017499 | WBGene00009000 | WBGene00004887 | WBGene00018682 | WBGene00017904 |
| WBGene00017500 | WBGene00009007 | WBGene00004888 | WBGene00018697 | WBGene00017923 |
| WBGene00017501 | WBGene00009056 | WBGene00004890 | WBGene00018701 | WBGene00017968 |
| WBGene00017503 | WBGene00009058 | WBGene00004892 | WBGene00018734 | WBGene00017979 |
| WBGene00017507 | WBGene00009069 | WBGene00004896 | WBGene00018756 | WBGene00017991 |
| WBGene00017514 | WBGene00009092 | WBGene00004914 | WBGene00018849 | WBGene00017998 |
| WBGene00017526 | WBGene00009100 | WBGene00004915 | WBGene00018967 | WBGene00018014 |
| WBGene00017535 | WBGene00009110 | WBGene00004916 | WBGene00018986 | WBGene00018031 |
| WBGene00017541 | WBGene00009116 | WBGene00004918 | WBGene00019001 | WBGene00018073 |
| WBGene00017567 | WBGene00009124 | WBGene00004919 | WBGene00019061 | WBGene00018144 |
| WBGene00017647 | WBGene00009139 | WBGene00004920 | WBGene00019118 | WBGene00018145 |
| WBGene00017688 | WBGene00009173 | WBGene00004921 | WBGene00019162 | WBGene00018268 |
| WBGene00017696 | WBGene00009174 | WBGene00004924 | WBGene00019212 | WBGene00018282 |
| WBGene00017741 | WBGene00009211 | WBGene00004927 | WBGene00019250 | WBGene00018293 |
| WBGene00017792 | WBGene00009224 | WBGene00004931 | WBGene00019272 | WBGene00018294 |
| WBGene00017811 | WBGene00009237 | WBGene00004933 | WBGene00019296 | WBGene00018297 |
| WBGene00017829 | WBGene00009254 | WBGene00004955 | WBGene00019301 | WBGene00018321 |
| WBGene00017832 | WBGene00009256 | WBGene00004974 | WBGene00019305 | WBGene00018330 |
| WBGene00017865 | WBGene00009270 | WBGene00004975 | WBGene00019334 | WBGene00018339 |
| WBGene00017888 | WBGene00009274 | WBGene00004984 | WBGene00019504 | WBGene00018403 |
| WBGene00017924 | WBGene00009276 | WBGene00004986 | WBGene00019521 | WBGene00018418 |
| WBGene00017955 | WBGene00009286 | WBGene00004987 | WBGene00019620 | WBGene00018482 |
| WBGene00017969 | WBGene00009289 | WBGene00004988 | WBGene00019628 | WBGene00018487 |
| WBGene00017975 | WBGene00009321 | WBGene00004990 | WBGene00019678 | WBGene00018488 |
| WBGene00017990 | WBGene00009323 | WBGene00004993 | WBGene00019680 | WBGene00018518 |
| WBGene00018016 | WBGene00009346 | WBGene00004995 | WBGene00019686 | WBGene00018586 |
| WBGene00018040 | WBGene00009347 | WBGene00004999 | WBGene00019748 | WBGene00018700 |
| WBGene00018043 | WBGene00009352 | WBGene00005000 | WBGene00019816 | WBGene00018744 |
| WBGene00018046 | WBGene00009368 | WBGene00005003 | WBGene00019877 | WBGene00018802 |
| WBGene00018061 | WBGene00009372 | WBGene00005007 | WBGene00019890 | WBGene00018811 |
| WBGene00018085 | WBGene00009377 | WBGene00005014 | WBGene00019893 | WBGene00018846 |
| WBGene00018100 | WBGene00009383 | WBGene00005017 | WBGene00019940 | WBGene00018869 |
| WBGene00018119 | WBGene00009388 | WBGene00005025 | WBGene00019953 | WBGene00018953 |
| WBGene00018122 | WBGene00009403 | WBGene00005026 | WBGene00019960 | WBGene00018969 |
| WBGene00018134 | WBGene00009404 | WBGene00005077 | WBGene00019983 | WBGene00018990 |
| WBGene00018140 | WBGene00009406 | WBGene00006050 | WBGene00020022 | WBGene00019007 |
| WBGene00018146 | WBGene00009430 | WBGene00006051 | WBGene00020033 | WBGene00019074 |
| WBGene00018188 | WBGene00009441 | WBGene00006311 | WBGene00020038 | WBGene00019122 |
| WBGene00018210 | WBGene00009448 | WBGene00006321 | WBGene00020047 | WBGene00019148 |
| WBGene00018239 | WBGene00009456 | WBGene00006351 | WBGene00020102 | WBGene00019204 |
| WBGene00018273 | WBGene00009478 | WBGene00006381 | WBGene00020107 | WBGene00019249 |
| WBGene00018305 | WBGene00009507 | WBGene00006384 | WBGene00020148 | WBGene00019268 |
| WBGene00018307 | WBGene00009512 | WBGene00006385 | WBGene00020184 | WBGene00019300 |
| WBGene00018328 | WBGene00009521 | WBGene00006390 | WBGene00020185 | WBGene00019304 |
| WBGene00018337 | WBGene00009529 | WBGene00006394 | WBGene00020216 | WBGene00019322 |
| WBGene00018380 | WBGene00009553 | WBGene00006396 | WBGene00020245 | WBGene00019327 |
| WBGene00018386 | WBGene00009554 | WBGene00006405 | WBGene00020297 | WBGene00019464 |
| WBGene00018390 | WBGene00009556 | WBGene00006406 | WBGene00020301 | WBGene00019465 |
| WBGene00018392 | WBGene00009563 | WBGene00006415 | WBGene00020398 | WBGene00019466 |
| WBGene00018410 | WBGene00009577 | WBGene00006433 | WBGene00020423 | WBGene00019489 |
| WBGene00018422 | WBGene00009578 | WBGene00006436 | WBGene00020441 | WBGene00019490 |
| WBGene00018459 | WBGene00009582 | WBGene00006442 | WBGene00020464 | WBGene00019492 |
| WBGene00018460 | WBGene00009617 | WBGene00006452 | WBGene00020481 | WBGene00019510 |
| WBGene00018461 | WBGene00009637 | WBGene00006460 | WBGene00020496 | WBGene00019537 |
| WBGene00018470 | WBGene00009640 | WBGene00006478 | WBGene00020551 | WBGene00019540 |
| WBGene00018489 | WBGene00009642 | WBGene00006479 | WBGene00020553 | WBGene00019572 |
| WBGene00018536 | WBGene00009654 | WBGene00006489 | WBGene00020554 | WBGene00019656 |
| WBGene00018547 | WBGene00009663 | WBGene00006490 | WBGene00020592 | WBGene00019760 |
| WBGene00018548 | WBGene00009668 | WBGene00006495 | WBGene00020628 | WBGene00019819 |
| WBGene00018578 | WBGene00009672 | WBGene00006496 | WBGene00020649 | WBGene00019855 |
| WBGene00018580 | WBGene00009729 | WBGene00006508 | WBGene00020679 | WBGene00019875 |
| WBGene00018730 | WBGene00009731 | WBGene00006510 | WBGene00020683 | WBGene00019895 |
| WBGene00018747 | WBGene00009736 | WBGene00006516 | WBGene00020721 | WBGene00019900 |
| WBGene00018763 | WBGene00009743 | WBGene00006519 | WBGene00020762 | WBGene00019947 |

| | | | | |
|---|---|---|---|---|
| WBGene00018783 | WBGene00009745 | WBGene00006527 | WBGene00020770 | WBGene00020007 |
| WBGene00018787 | WBGene00009776 | WBGene00006535 | WBGene00020801 | WBGene00020031 |
| WBGene00018795 | WBGene00009784 | WBGene00006539 | WBGene00020806 | WBGene00020092 |
| WBGene00018835 | WBGene00009801 | WBGene00006542 | WBGene00020821 | WBGene00020097 |
| WBGene00018836 | WBGene00009804 | WBGene00006563 | WBGene00020823 | WBGene00020131 |
| WBGene00018839 | WBGene00009820 | WBGene00006573 | WBGene00020868 | WBGene00020139 |
| WBGene00018845 | WBGene00009823 | WBGene00006577 | WBGene00020911 | WBGene00020160 |
| WBGene00018863 | WBGene00009824 | WBGene00006578 | WBGene00020936 | WBGene00020169 |
| WBGene00018899 | WBGene00009825 | WBGene00006583 | WBGene00021296 | WBGene00020219 |
| WBGene00018909 | WBGene00009877 | WBGene00006584 | WBGene00021326 | WBGene00020269 |
| WBGene00018946 | WBGene00009884 | WBGene00006585 | WBGene00021391 | WBGene00020274 |
| WBGene00018968 | WBGene00009885 | WBGene00006587 | WBGene00021466 | WBGene00020275 |
| WBGene00018976 | WBGene00009886 | WBGene00006589 | WBGene00021468 | WBGene00020284 |
| WBGene00019005 | WBGene00009896 | WBGene00006602 | WBGene00021470 | WBGene00020315 |
| WBGene00019030 | WBGene00009897 | WBGene00006604 | WBGene00021532 | WBGene00020335 |
| WBGene00019079 | WBGene00009898 | WBGene00006607 | WBGene00021628 | WBGene00020347 |
| WBGene00019143 | WBGene00009902 | WBGene00006608 | WBGene00021685 | WBGene00020348 |
| WBGene00019149 | WBGene00009917 | WBGene00006613 | WBGene00021692 | WBGene00020366 |
| WBGene00019166 | WBGene00009937 | WBGene00006647 | WBGene00021732 | WBGene00020391 |
| WBGene00019262 | WBGene00009958 | WBGene00006697 | WBGene00021834 | WBGene00020425 |
| WBGene00019264 | WBGene00009959 | WBGene00006698 | WBGene00021888 | WBGene00020507 |
| WBGene00019276 | WBGene00009968 | WBGene00006701 | WBGene00021899 | WBGene00020530 |
| WBGene00019283 | WBGene00009973 | WBGene00006703 | WBGene00021912 | WBGene00020556 |
| WBGene00019285 | WBGene00009981 | WBGene00006711 | WBGene00021956 | WBGene00020696 |
| WBGene00019318 | WBGene00009994 | WBGene00006715 | WBGene00021986 | WBGene00020824 |
| WBGene00019341 | WBGene00010000 | WBGene00006730 | WBGene00022037 | WBGene00020827 |
| WBGene00019388 | WBGene00010002 | WBGene00006737 | WBGene00022069 | WBGene00020831 |
| WBGene00019411 | WBGene00010006 | WBGene00006742 | WBGene00022131 | WBGene00020846 |
| WBGene00019448 | WBGene00010007 | WBGene00006750 | WBGene00022169 | WBGene00020865 |
| WBGene00019456 | WBGene00010010 | WBGene00006751 | WBGene00022170 | WBGene00020891 |
| WBGene00019482 | WBGene00010036 | WBGene00006752 | WBGene00022199 | WBGene00020915 |
| WBGene00019530 | WBGene00010044 | WBGene00006754 | WBGene00022235 | WBGene00020930 |
| WBGene00019535 | WBGene00010047 | WBGene00006755 | WBGene00022245 | WBGene00020937 |
| WBGene00019561 | WBGene00010051 | WBGene00006757 | WBGene00022279 | WBGene00020950 |
| WBGene00019564 | WBGene00010054 | WBGene00006759 | WBGene00022456 | WBGene00020952 |
| WBGene00019570 | WBGene00010056 | WBGene00006762 | WBGene00022500 | WBGene00021068 |
| WBGene00019624 | WBGene00010077 | WBGene00006763 | WBGene00022631 | WBGene00021088 |
| WBGene00019631 | WBGene00010089 | WBGene00006764 | WBGene00022643 | WBGene00021095 |
| WBGene00019636 | WBGene00010094 | WBGene00006770 | WBGene00022680 | WBGene00021128 |
| WBGene00019649 | WBGene00010109 | WBGene00006776 | WBGene00022703 | WBGene00021281 |
| WBGene00019730 | WBGene00010131 | WBGene00006784 | WBGene00022712 | WBGene00021348 |
| WBGene00019751 | WBGene00010132 | WBGene00006785 | WBGene00022718 | WBGene00021350 |
| WBGene00019759 | WBGene00010140 | WBGene00006788 | WBGene00022748 | WBGene00021352 |
| WBGene00019812 | WBGene00010188 | WBGene00006789 | WBGene00022783 | WBGene00021369 |
| WBGene00019833 | WBGene00010196 | WBGene00006793 | WBGene00022787 | WBGene00021420 |
| WBGene00019892 | WBGene00010198 | WBGene00006798 | WBGene00022793 | WBGene00021427 |
| WBGene00019906 | WBGene00010223 | WBGene00006801 | WBGene00022794 | WBGene00021430 |
| WBGene00019931 | WBGene00010227 | WBGene00006802 | WBGene00022814 | WBGene00021469 |
| WBGene00019939 | WBGene00010238 | WBGene00006804 | WBGene00022852 | WBGene00021486 |
| WBGene00019948 | WBGene00010281 | WBGene00006805 | WBGene00022880 | WBGene00021533 |
| WBGene00019989 | WBGene00010288 | WBGene00006808 | WBGene00022882 | WBGene00021535 |
| WBGene00019992 | WBGene00010304 | WBGene00006811 | WBGene00044026 | WBGene00021561 |
| WBGene00020003 | WBGene00010307 | WBGene00006812 | WBGene00044069 | WBGene00021597 |
| WBGene00020029 | WBGene00010308 | WBGene00006814 | WBGene00044071 | WBGene00021645 |
| WBGene00020040 | WBGene00010330 | WBGene00006819 | WBGene00044077 | WBGene00021835 |
| WBGene00020071 | WBGene00010332 | WBGene00006820 | WBGene00044324 | WBGene00021883 |
| WBGene00020083 | WBGene00010334 | WBGene00006823 | WBGene00044396 | WBGene00021960 |
| WBGene00020089 | WBGene00010336 | WBGene00006829 | WBGene00044483 | WBGene00021975 |
| WBGene00020093 | WBGene00010338 | WBGene00006830 | WBGene00044553 | WBGene00022010 |
| WBGene00020095 | WBGene00010348 | WBGene00006863 | WBGene00044779 | WBGene00022033 |
| WBGene00020103 | WBGene00010353 | WBGene00006868 | WBGene00044989 | WBGene00022042 |
| WBGene00020130 | WBGene00010362 | WBGene00006874 | WBGene00045419 | WBGene00022089 |
| WBGene00020189 | WBGene00010367 | WBGene00006889 | WBGene00077526 | WBGene00022103 |
| WBGene00020193 | WBGene00010368 | WBGene00006915 | WBGene00077697 | WBGene00022114 |
| WBGene00020194 | WBGene00010383 | WBGene00006925 | WBGene00077761 | WBGene00022119 |
| WBGene00020197 | WBGene00010397 | WBGene00006926 | WBGene00175030 | WBGene00022122 |
| WBGene00020248 | WBGene00010420 | WBGene00006927 | WBGene00195248 | WBGene00022139 |
| WBGene00020263 | WBGene00010422 | WBGene00006928 | | WBGene00022158 |

| | | | |
|---|---|---|---|
| WBGene00020282 | WBGene00010439 | WBGene00006929 | WBGene00022160 |
| WBGene00020285 | WBGene00010444 | WBGene00006930 | WBGene00022231 |
| WBGene00020299 | WBGene00010458 | WBGene00006931 | WBGene00022233 |
| WBGene00020350 | WBGene00010481 | WBGene00006932 | WBGene00022307 |
| WBGene00020353 | WBGene00010496 | WBGene00006933 | WBGene00022336 |
| WBGene00020386 | WBGene00010512 | WBGene00006935 | WBGene00022358 |
| WBGene00020444 | WBGene00010513 | WBGene00006936 | WBGene00022517 |
| WBGene00020450 | WBGene00010537 | WBGene00006937 | WBGene00022538 |
| WBGene00020451 | WBGene00010546 | WBGene00006942 | WBGene00022599 |
| WBGene00020467 | WBGene00010552 | WBGene00006949 | WBGene00022678 |
| WBGene00020502 | WBGene00010563 | WBGene00006957 | WBGene00022679 |
| WBGene00020678 | WBGene00010572 | WBGene00006961 | WBGene00022683 |
| WBGene00020703 | WBGene00010582 | WBGene00006977 | WBGene00022722 |
| WBGene00020713 | WBGene00010593 | WBGene00006986 | WBGene00022743 |
| WBGene00020722 | WBGene00010608 | WBGene00006993 | WBGene00022766 |
| WBGene00020783 | WBGene00010622 | WBGene00006994 | WBGene00022781 |
| WBGene00020817 | WBGene00010624 | WBGene00006999 | WBGene00022797 |
| WBGene00020837 | WBGene00010646 | WBGene00007007 | WBGene00022800 |
| WBGene00020840 | WBGene00010660 | WBGene00007010 | WBGene00022817 |
| WBGene00020841 | WBGene00010663 | WBGene00007022 | WBGene00022850 |
| WBGene00020844 | WBGene00010678 | WBGene00007023 | WBGene00022856 |
| WBGene00020852 | WBGene00010679 | WBGene00007025 | WBGene00022861 |
| WBGene00020862 | WBGene00010687 | WBGene00007026 | WBGene00023500 |
| WBGene00020883 | WBGene00010702 | WBGene00007027 | WBGene00044019 |
| WBGene00020895 | WBGene00010703 | WBGene00007028 | WBGene00044061 |
| WBGene00020918 | WBGene00010713 | WBGene00007029 | WBGene00044294 |
| WBGene00020920 | WBGene00010724 | WBGene00007050 | WBGene00045399 |
| WBGene00020941 | WBGene00010726 | WBGene00007054 | WBGene00077693 |
| WBGene00020983 | WBGene00010732 | WBGene00007055 | WBGene00185014 |
| WBGene00021089 | WBGene00010744 | WBGene00007059 | WBGene00219376 |
| WBGene00021114 | WBGene00010756 | WBGene00007088 | WBGene00235102 |
| WBGene00021195 | WBGene00010767 | WBGene00007089 | WBGene00017984 |
| WBGene00021212 | WBGene00010772 | WBGene00007090 | WBGene00015470 |
| WBGene00021219 | WBGene00010810 | WBGene00007091 | WBGene00010896 |
| WBGene00021236 | WBGene00010814 | WBGene00007107 | |
| WBGene00021253 | WBGene00010844 | WBGene00007111 | |
| WBGene00021270 | WBGene00010848 | WBGene00007122 | |
| WBGene00021279 | WBGene00010856 | WBGene00007129 | |
| WBGene00021295 | WBGene00010872 | WBGene00007136 | |
| WBGene00021319 | WBGene00010875 | WBGene00007144 | |
| WBGene00021321 | WBGene00010888 | WBGene00007145 | |
| WBGene00021345 | WBGene00010892 | WBGene00007155 | |
| WBGene00021347 | WBGene00010912 | WBGene00007158 | |
| WBGene00021366 | WBGene00010913 | WBGene00007167 | |
| WBGene00021375 | WBGene00010922 | WBGene00007175 | |
| WBGene00021394 | WBGene00010931 | WBGene00007184 | |
| WBGene00021398 | WBGene00010936 | WBGene00007186 | |
| WBGene00021403 | WBGene00010975 | WBGene00007192 | |
| WBGene00021406 | WBGene00010980 | WBGene00007193 | |
| WBGene00021419 | WBGene00011000 | WBGene00007196 | |
| WBGene00021472 | WBGene00011020 | WBGene00007197 | |
| WBGene00021475 | WBGene00011029 | WBGene00007217 | |
| WBGene00021497 | WBGene00011032 | WBGene00007226 | |
| WBGene00021529 | WBGene00011035 | WBGene00007235 | |
| WBGene00021557 | WBGene00011037 | WBGene00007252 | |
| WBGene00021594 | WBGene00011040 | WBGene00007261 | |
| WBGene00021630 | WBGene00011043 | WBGene00007277 | |
| WBGene00021635 | WBGene00011045 | WBGene00007278 | |
| WBGene00021638 | WBGene00011047 | WBGene00007284 | |
| WBGene00021641 | WBGene00011064 | WBGene00007296 | |
| WBGene00021736 | WBGene00011067 | WBGene00007297 | |
| WBGene00021740 | WBGene00011069 | WBGene00007301 | |
| WBGene00021741 | WBGene00011070 | WBGene00007304 | |
| WBGene00021776 | WBGene00011087 | WBGene00007307 | |
| WBGene00021812 | WBGene00011094 | WBGene00007324 | |
| WBGene00021846 | WBGene00011106 | WBGene00007329 | |
| WBGene00021855 | WBGene00011109 | WBGene00007330 | |
| WBGene00021898 | WBGene00011123 | WBGene00007333 | |

| | | |
|---|---|---|
| WBGene00021926 | WBGene00011124 | WBGene00007348 |
| WBGene00021933 | WBGene00011131 | WBGene00007353 |
| WBGene00021945 | WBGene00011134 | WBGene00007390 |
| WBGene00021967 | WBGene00011142 | WBGene00007400 |
| WBGene00021983 | WBGene00011143 | WBGene00007413 |
| WBGene00021997 | WBGene00011182 | WBGene00007429 |
| WBGene00022002 | WBGene00011193 | WBGene00007430 |
| WBGene00022005 | WBGene00011194 | WBGene00007433 |
| WBGene00022024 | WBGene00011215 | WBGene00007443 |
| WBGene00022031 | WBGene00011221 | WBGene00007445 |
| WBGene00022038 | WBGene00011224 | WBGene00007448 |
| WBGene00022039 | WBGene00011231 | WBGene00007458 |
| WBGene00022043 | WBGene00011242 | WBGene00007500 |
| WBGene00022057 | WBGene00011250 | WBGene00007501 |
| WBGene00022076 | WBGene00011252 | WBGene00007516 |
| WBGene00022126 | WBGene00011253 | WBGene00007525 |
| WBGene00022128 | WBGene00011268 | WBGene00007554 |
| WBGene00022154 | WBGene00011286 | WBGene00007561 |
| WBGene00022162 | WBGene00011293 | WBGene00007564 |
| WBGene00022252 | WBGene00011309 | WBGene00007565 |
| WBGene00022298 | WBGene00011310 | WBGene00007577 |
| WBGene00022302 | WBGene00011314 | WBGene00007591 |
| WBGene00022309 | WBGene00011320 | WBGene00007593 |
| WBGene00022343 | WBGene00011330 | WBGene00007599 |
| WBGene00022371 | WBGene00011331 | WBGene00007602 |
| WBGene00022377 | WBGene00011352 | WBGene00007609 |
| WBGene00022378 | WBGene00011399 | WBGene00007617 |
| WBGene00022385 | WBGene00011410 | WBGene00007621 |
| WBGene00022392 | WBGene00011424 | WBGene00007624 |
| WBGene00022415 | WBGene00011428 | WBGene00007625 |
| WBGene00022418 | WBGene00011441 | WBGene00007626 |
| WBGene00022428 | WBGene00011444 | WBGene00007627 |
| WBGene00022441 | WBGene00011489 | WBGene00007628 |
| WBGene00022455 | WBGene00011509 | WBGene00007643 |
| WBGene00022474 | WBGene00011531 | WBGene00007645 |
| WBGene00022546 | WBGene00011560 | WBGene00007647 |
| WBGene00022562 | WBGene00011583 | WBGene00007653 |
| WBGene00022576 | WBGene00011600 | WBGene00007666 |
| WBGene00022591 | WBGene00011625 | WBGene00007683 |
| WBGene00022597 | WBGene00011637 | WBGene00007684 |
| WBGene00022609 | WBGene00011645 | WBGene00007686 |
| WBGene00022611 | WBGene00011648 | WBGene00007689 |
| WBGene00022612 | WBGene00011683 | WBGene00007707 |
| WBGene00022670 | WBGene00011708 | WBGene00007709 |
| WBGene00022708 | WBGene00011720 | WBGene00007710 |
| WBGene00022728 | WBGene00011723 | WBGene00007712 |
| WBGene00022730 | WBGene00011740 | WBGene00007720 |
| WBGene00022746 | WBGene00011741 | WBGene00007730 |
| WBGene00022786 | WBGene00011755 | WBGene00007733 |
| WBGene00022845 | WBGene00011761 | WBGene00007758 |
| WBGene00022848 | WBGene00011788 | WBGene00007761 |
| WBGene00022851 | WBGene00011794 | WBGene00007765 |
| WBGene00023422 | WBGene00011800 | WBGene00007766 |
| WBGene00023424 | WBGene00011806 | WBGene00007772 |
| WBGene00023491 | WBGene00011813 | WBGene00007784 |
| WBGene00023493 | WBGene00011824 | WBGene00007786 |
| WBGene00023504 | WBGene00011830 | WBGene00007799 |
| WBGene00043057 | WBGene00011844 | WBGene00007824 |
| WBGene00043302 | WBGene00011864 | WBGene00007846 |
| WBGene00043948 | WBGene00011868 | WBGene00007855 |
| WBGene00043994 | WBGene00011870 | WBGene00007880 |
| WBGene00044008 | WBGene00011872 | WBGene00007881 |
| WBGene00044024 | WBGene00011875 | WBGene00007907 |
| WBGene00044045 | WBGene00011886 | WBGene00007921 |
| WBGene00044063 | WBGene00011892 | WBGene00007923 |
| WBGene00044078 | WBGene00011912 | WBGene00007925 |
| WBGene00044094 | WBGene00011915 | WBGene00007931 |
| WBGene00044120 | WBGene00011916 | WBGene00007941 |

| | | |
|---|---|---|
| WBGene00044163 | WBGene00011920 | WBGene00007942 |
| WBGene00044201 | WBGene00011923 | WBGene00007943 |
| WBGene00044236 | WBGene00011940 | WBGene00007952 |
| WBGene00044245 | WBGene00011979 | WBGene00007955 |
| WBGene00044250 | WBGene00011984 | WBGene00007957 |
| WBGene00044316 | WBGene00011988 | WBGene00007958 |
| WBGene00044322 | WBGene00012000 | WBGene00007959 |
| WBGene00044323 | WBGene00012002 | WBGene00007966 |
| WBGene00044330 | WBGene00012073 | WBGene00007978 |
| WBGene00044333 | WBGene00012087 | WBGene00007979 |
| WBGene00044361 | WBGene00012100 | WBGene00007980 |
| WBGene00044374 | WBGene00012125 | WBGene00007984 |
| WBGene00044385 | WBGene00012126 | WBGene00007993 |
| WBGene00044394 | WBGene00012143 | WBGene00008020 |
| WBGene00044401 | WBGene00012150 | WBGene00008034 |
| WBGene00044421 | WBGene00012182 | WBGene00008035 |
| WBGene00044447 | WBGene00012192 | WBGene00008041 |
| WBGene00044463 | WBGene00012197 | WBGene00008082 |
| WBGene00044472 | WBGene00012205 | WBGene00008107 |
| WBGene00044484 | WBGene00012213 | WBGene00008110 |
| WBGene00044487 | WBGene00012217 | WBGene00008121 |
| WBGene00044492 | WBGene00012231 | WBGene00008131 |
| WBGene00044578 | WBGene00012241 | WBGene00008134 |
| WBGene00044589 | WBGene00012251 | WBGene00008140 |
| WBGene00044615 | WBGene00012252 | WBGene00008147 |
| WBGene00044623 | WBGene00012276 | WBGene00008149 |
| WBGene00044643 | WBGene00012277 | WBGene00008164 |
| WBGene00044654 | WBGene00012279 | WBGene00008170 |
| WBGene00044672 | WBGene00012291 | WBGene00008194 |
| WBGene00044697 | WBGene00012295 | WBGene00008219 |
| WBGene00044720 | WBGene00012306 | WBGene00008229 |
| WBGene00044762 | WBGene00012315 | WBGene00008237 |
| WBGene00044776 | WBGene00012316 | WBGene00008258 |
| WBGene00044796 | WBGene00012341 | WBGene00008262 |
| WBGene00045036 | WBGene00012353 | WBGene00008284 |
| WBGene00045146 | WBGene00012369 | WBGene00008311 |
| WBGene00045177 | WBGene00012385 | WBGene00008330 |
| WBGene00045269 | WBGene00012389 | WBGene00008339 |
| WBGene00045275 | WBGene00012395 | WBGene00008343 |
| WBGene00045355 | WBGene00012423 | WBGene00008345 |
| WBGene00045356 | WBGene00012445 | WBGene00008356 |
| WBGene00045392 | WBGene00012446 | WBGene00008359 |
| WBGene00045397 | WBGene00012458 | WBGene00008360 |
| WBGene00045411 | WBGene00012465 | WBGene00008366 |
| WBGene00050875 | WBGene00012469 | WBGene00008371 |
| WBGene00050885 | WBGene00012474 | WBGene00008377 |
| WBGene00050957 | WBGene00012479 | WBGene00008378 |
| WBGene00077450 | WBGene00012481 | WBGene00008380 |
| WBGene00077521 | WBGene00012496 | WBGene00008386 |
| WBGene00077685 | WBGene00012512 | WBGene00008390 |
| WBGene00077691 | WBGene00012514 | WBGene00008408 |
| WBGene00077764 | WBGene00012559 | WBGene00008410 |
| WBGene00138711 | WBGene00012560 | WBGene00008415 |
| WBGene00138717 | WBGene00012564 | WBGene00008417 |
| WBGene00138724 | WBGene00012595 | WBGene00008435 |
| WBGene00175027 | WBGene00012644 | WBGene00008440 |
| WBGene00185005 | WBGene00012659 | WBGene00008446 |
| WBGene00185048 | WBGene00012663 | WBGene00008452 |
| WBGene00185079 | WBGene00012665 | WBGene00008454 |
| WBGene00185118 | WBGene00012674 | WBGene00008458 |
| WBGene00194677 | WBGene00012676 | WBGene00008459 |
| WBGene00194699 | WBGene00012678 | WBGene00008530 |
| WBGene00194727 | WBGene00012693 | WBGene00008538 |
| WBGene00194738 | WBGene00012696 | WBGene00008546 |
| WBGene00194795 | WBGene00012701 | WBGene00008548 |
| WBGene00194849 | WBGene00012715 | WBGene00008564 |
| WBGene00194894 | WBGene00012718 | WBGene00008571 |
| WBGene00194905 | WBGene00012730 | WBGene00008572 |

| | | |
|---|---|---|
| WBGene00194913 | WBGene00012732 | WBGene00008603 |
| WBGene00194952 | WBGene00012734 | WBGene00008607 |
| WBGene00195004 | WBGene00012735 | WBGene00008610 |
| WBGene00195051 | WBGene00012740 | WBGene00008629 |
| WBGene00195142 | WBGene00012750 | WBGene00008639 |
| WBGene00195147 | WBGene00012756 | WBGene00008645 |
| WBGene00195185 | WBGene00012757 | WBGene00008670 |
| WBGene00195209 | WBGene00012802 | WBGene00008677 |
| WBGene00206354 | WBGene00012804 | WBGene00008694 |
| WBGene00206360 | WBGene00012807 | WBGene00008729 |
| WBGene00206389 | WBGene00012810 | WBGene00008739 |
| WBGene00206418 | WBGene00012830 | WBGene00008741 |
| WBGene00206482 | WBGene00012834 | WBGene00008745 |
| WBGene00206500 | WBGene00012835 | WBGene00008750 |
| WBGene00206502 | WBGene00012854 | WBGene00008760 |
| WBGene00219308 | WBGene00012859 | WBGene00008767 |
| WBGene00219329 | WBGene00012861 | WBGene00008777 |
| WBGene00219378 | WBGene00012879 | WBGene00008820 |
| WBGene00219434 | WBGene00012880 | WBGene00008831 |
| WBGene00235164 | WBGene00012885 | WBGene00008841 |
| WBGene00235267 | WBGene00012890 | WBGene00008865 |
| WBGene00235271 | WBGene00012904 | WBGene00008914 |
| WBGene00235352 | WBGene00012909 | WBGene00008915 |
| | WBGene00012910 | WBGene00008919 |
| | WBGene00012911 | WBGene00008921 |
| | WBGene00012928 | WBGene00008925 |
| | WBGene00012935 | WBGene00008930 |
| | WBGene00012936 | WBGene00008932 |
| | WBGene00012948 | WBGene00008934 |
| | WBGene00012950 | WBGene00008944 |
| | WBGene00012951 | WBGene00008973 |
| | WBGene00012968 | WBGene00008980 |
| | WBGene00012977 | WBGene00008986 |
| | WBGene00012982 | WBGene00008991 |
| | WBGene00012987 | WBGene00009004 |
| | WBGene00012989 | WBGene00009006 |
| | WBGene00012992 | WBGene00009010 |
| | WBGene00012994 | WBGene00009011 |
| | WBGene00012995 | WBGene00009012 |
| | WBGene00012997 | WBGene00009031 |
| | WBGene00013000 | WBGene00009035 |
| | WBGene00013006 | WBGene00009045 |
| | WBGene00013015 | WBGene00009049 |
| | WBGene00013020 | WBGene00009051 |
| | WBGene00013021 | WBGene00009052 |
| | WBGene00013024 | WBGene00009059 |
| | WBGene00013040 | WBGene00009065 |
| | WBGene00013042 | WBGene00009078 |
| | WBGene00013074 | WBGene00009096 |
| | WBGene00013075 | WBGene00009097 |
| | WBGene00013083 | WBGene00009103 |
| | WBGene00013120 | WBGene00009113 |
| | WBGene00013129 | WBGene00009127 |
| | WBGene00013137 | WBGene00009140 |
| | WBGene00013156 | WBGene00009157 |
| | WBGene00013164 | WBGene00009161 |
| | WBGene00013167 | WBGene00009180 |
| | WBGene00013169 | WBGene00009186 |
| | WBGene00013176 | WBGene00009189 |
| | WBGene00013177 | WBGene00009191 |
| | WBGene00013193 | WBGene00009200 |
| | WBGene00013196 | WBGene00009201 |
| | WBGene00013208 | WBGene00009207 |
| | WBGene00013214 | WBGene00009215 |
| | WBGene00013224 | WBGene00009220 |
| | WBGene00013231 | WBGene00009221 |
| | WBGene00013243 | WBGene00009223 |
| | WBGene00013256 | WBGene00009234 |

WBGene00013265
WBGene00013268
WBGene00013269
WBGene00013292
WBGene00013327
WBGene00013344
WBGene00013350
WBGene00013359
WBGene00013362
WBGene00013380
WBGene00013391
WBGene00013392
WBGene00013393
WBGene00013394
WBGene00013405
WBGene00013441
WBGene00013460
WBGene00013473
WBGene00013477
WBGene00013482
WBGene00013506
WBGene00013515
WBGene00013530
WBGene00013538
WBGene00013540
WBGene00013549
WBGene00013552
WBGene00013557
WBGene00013561
WBGene00013587
WBGene00013593
WBGene00013594
WBGene00013596
WBGene00013598
WBGene00013601
WBGene00013604
WBGene00013632
WBGene00013654
WBGene00013668
WBGene00013670
WBGene00013674
WBGene00013679
WBGene00013687
WBGene00013691
WBGene00013697
WBGene00013700
WBGene00013710
WBGene00013713
WBGene00013720
WBGene00013725
WBGene00013728
WBGene00013729
WBGene00013730
WBGene00013737
WBGene00013738
WBGene00013768
WBGene00013804
WBGene00013835
WBGene00013854
WBGene00013857
WBGene00013860
WBGene00013862
WBGene00013869
WBGene00013870
WBGene00013896
WBGene00013925
WBGene00013946
WBGene00013947
WBGene00009246
WBGene00009259
WBGene00009267
WBGene00009272
WBGene00009277
WBGene00009288
WBGene00009290
WBGene00009294
WBGene00009307
WBGene00009319
WBGene00009322
WBGene00009336
WBGene00009337
WBGene00009353
WBGene00009365
WBGene00009366
WBGene00009369
WBGene00009374
WBGene00009385
WBGene00009393
WBGene00009394
WBGene00009396
WBGene00009435
WBGene00009436
WBGene00009440
WBGene00009445
WBGene00009455
WBGene00009457
WBGene00009505
WBGene00009514
WBGene00009531
WBGene00009534
WBGene00009551
WBGene00009562
WBGene00009565
WBGene00009575
WBGene00009580
WBGene00009620
WBGene00009624
WBGene00009632
WBGene00009634
WBGene00009636
WBGene00009638
WBGene00009650
WBGene00009657
WBGene00009664
WBGene00009665
WBGene00009666
WBGene00009667
WBGene00009671
WBGene00009677
WBGene00009679
WBGene00009681
WBGene00009682
WBGene00009688
WBGene00009701
WBGene00009711
WBGene00009732
WBGene00009734
WBGene00009739
WBGene00009744
WBGene00009778
WBGene00009796
WBGene00009803
WBGene00009807
WBGene00009881
WBGene00009895
WBGene00009919

| | |
|---|---|
| WBGene00013962 | WBGene00009920 |
| WBGene00013966 | WBGene00009921 |
| WBGene00013967 | WBGene00009922 |
| WBGene00013987 | WBGene00009924 |
| WBGene00013994 | WBGene00009940 |
| WBGene00013997 | WBGene00009945 |
| WBGene00014003 | WBGene00009957 |
| WBGene00014023 | WBGene00009966 |
| WBGene00014035 | WBGene00009974 |
| WBGene00014078 | WBGene00009975 |
| WBGene00014081 | WBGene00009992 |
| WBGene00014083 | WBGene00009995 |
| WBGene00014090 | WBGene00010005 |
| WBGene00014099 | WBGene00010013 |
| WBGene00014114 | WBGene00010019 |
| WBGene00014120 | WBGene00010028 |
| WBGene00014124 | WBGene00010040 |
| WBGene00014150 | WBGene00010041 |
| WBGene00014152 | WBGene00010049 |
| WBGene00014177 | WBGene00010050 |
| WBGene00014213 | WBGene00010053 |
| WBGene00014215 | WBGene00010061 |
| WBGene00014224 | WBGene00010070 |
| WBGene00014228 | WBGene00010075 |
| WBGene00014230 | WBGene00010081 |
| WBGene00014233 | WBGene00010096 |
| WBGene00014247 | WBGene00010097 |
| WBGene00014261 | WBGene00010113 |
| WBGene00014666 | WBGene00010124 |
| WBGene00014848 | WBGene00010130 |
| WBGene00014999 | WBGene00010133 |
| WBGene00015023 | WBGene00010137 |
| WBGene00015043 | WBGene00010139 |
| WBGene00015047 | WBGene00010159 |
| WBGene00015057 | WBGene00010174 |
| WBGene00015061 | WBGene00010177 |
| WBGene00015073 | WBGene00010178 |
| WBGene00015091 | WBGene00010192 |
| WBGene00015092 | WBGene00010197 |
| WBGene00015095 | WBGene00010204 |
| WBGene00015096 | WBGene00010236 |
| WBGene00015104 | WBGene00010244 |
| WBGene00015115 | WBGene00010259 |
| WBGene00015122 | WBGene00010263 |
| WBGene00015132 | WBGene00010266 |
| WBGene00015134 | WBGene00010272 |
| WBGene00015160 | WBGene00010286 |
| WBGene00015189 | WBGene00010306 |
| WBGene00015194 | WBGene00010320 |
| WBGene00015205 | WBGene00010339 |
| WBGene00015216 | WBGene00010351 |
| WBGene00015217 | WBGene00010354 |
| WBGene00015219 | WBGene00010357 |
| WBGene00015238 | WBGene00010369 |
| WBGene00015245 | WBGene00010405 |
| WBGene00015267 | WBGene00010406 |
| WBGene00015282 | WBGene00010408 |
| WBGene00015295 | WBGene00010416 |
| WBGene00015303 | WBGene00010427 |
| WBGene00015308 | WBGene00010428 |
| WBGene00015313 | WBGene00010435 |
| WBGene00015347 | WBGene00010437 |
| WBGene00015389 | WBGene00010440 |
| WBGene00015407 | WBGene00010465 |
| WBGene00015408 | WBGene00010471 |
| WBGene00015425 | WBGene00010484 |
| WBGene00015429 | WBGene00010488 |
| WBGene00015449 | WBGene00010491 |

WBGene00015451
WBGene00015454
WBGene00015462
WBGene00015469
WBGene00015483
WBGene00015492
WBGene00015499
WBGene00015504
WBGene00015515
WBGene00015554
WBGene00015557
WBGene00015559
WBGene00015560
WBGene00015567
WBGene00015579
WBGene00015580
WBGene00015621
WBGene00015658
WBGene00015678
WBGene00015683
WBGene00015686
WBGene00015692
WBGene00015697
WBGene00015702
WBGene00015705
WBGene00015709
WBGene00015713
WBGene00015731
WBGene00015739
WBGene00015744
WBGene00015747
WBGene00015754
WBGene00015779
WBGene00015783
WBGene00015805
WBGene00015815
WBGene00015817
WBGene00015818
WBGene00015847
WBGene00015913
WBGene00015916
WBGene00015937
WBGene00015938
WBGene00015955
WBGene00015956
WBGene00015975
WBGene00016046
WBGene00016058
WBGene00016073
WBGene00016074
WBGene00016084
WBGene00016097
WBGene00016102
WBGene00016114
WBGene00016124
WBGene00016134
WBGene00016138
WBGene00016142
WBGene00016143
WBGene00016151
WBGene00016157
WBGene00016182
WBGene00016192
WBGene00016197
WBGene00016210
WBGene00016237
WBGene00016249
WBGene00016260
WBGene00010492
WBGene00010498
WBGene00010500
WBGene00010539
WBGene00010551
WBGene00010565
WBGene00010605
WBGene00010609
WBGene00010615
WBGene00010616
WBGene00010620
WBGene00010626
WBGene00010645
WBGene00010670
WBGene00010671
WBGene00010685
WBGene00010701
WBGene00010709
WBGene00010725
WBGene00010727
WBGene00010738
WBGene00010742
WBGene00010769
WBGene00010783
WBGene00010793
WBGene00010794
WBGene00010806
WBGene00010807
WBGene00010813
WBGene00010815
WBGene00010842
WBGene00010845
WBGene00010870
WBGene00010879
WBGene00010890
WBGene00010891
WBGene00010897
WBGene00010902
WBGene00010905
WBGene00010908
WBGene00010909
WBGene00010910
WBGene00010918
WBGene00010923
WBGene00010935
WBGene00010942
WBGene00010944
WBGene00010972
WBGene00010985
WBGene00010990
WBGene00011016
WBGene00011034
WBGene00011038
WBGene00011039
WBGene00011054
WBGene00011060
WBGene00011061
WBGene00011063
WBGene00011072
WBGene00011089
WBGene00011102
WBGene00011112
WBGene00011115
WBGene00011119
WBGene00011144
WBGene00011145
WBGene00011155
WBGene00011156

| | |
|---|---|
| WBGene00016289 | WBGene00011176 |
| WBGene00016294 | WBGene00011181 |
| WBGene00016317 | WBGene00011195 |
| WBGene00016362 | WBGene00011198 |
| WBGene00016373 | WBGene00011206 |
| WBGene00016387 | WBGene00011226 |
| WBGene00016390 | WBGene00011237 |
| WBGene00016399 | WBGene00011248 |
| WBGene00016406 | WBGene00011270 |
| WBGene00016409 | WBGene00011272 |
| WBGene00016412 | WBGene00011276 |
| WBGene00016434 | WBGene00011279 |
| WBGene00016443 | WBGene00011281 |
| WBGene00016444 | WBGene00011282 |
| WBGene00016454 | WBGene00011283 |
| WBGene00016458 | WBGene00011289 |
| WBGene00016474 | WBGene00011299 |
| WBGene00016522 | WBGene00011303 |
| WBGene00016530 | WBGene00011306 |
| WBGene00016534 | WBGene00011318 |
| WBGene00016539 | WBGene00011319 |
| WBGene00016541 | WBGene00011323 |
| WBGene00016562 | WBGene00011329 |
| WBGene00016568 | WBGene00011334 |
| WBGene00016570 | WBGene00011335 |
| WBGene00016575 | WBGene00011338 |
| WBGene00016587 | WBGene00011349 |
| WBGene00016609 | WBGene00011353 |
| WBGene00016620 | WBGene00011374 |
| WBGene00016624 | WBGene00011375 |
| WBGene00016632 | WBGene00011376 |
| WBGene00016643 | WBGene00011383 |
| WBGene00016652 | WBGene00011391 |
| WBGene00016658 | WBGene00011392 |
| WBGene00016663 | WBGene00011412 |
| WBGene00016669 | WBGene00011423 |
| WBGene00016676 | WBGene00011446 |
| WBGene00016678 | WBGene00011449 |
| WBGene00016684 | WBGene00011460 |
| WBGene00016702 | WBGene00011462 |
| WBGene00016704 | WBGene00011464 |
| WBGene00016725 | WBGene00011474 |
| WBGene00016754 | WBGene00011490 |
| WBGene00016792 | WBGene00011498 |
| WBGene00016793 | WBGene00011502 |
| WBGene00016795 | WBGene00011503 |
| WBGene00016799 | WBGene00011507 |
| WBGene00016812 | WBGene00011510 |
| WBGene00016816 | WBGene00011511 |
| WBGene00016836 | WBGene00011532 |
| WBGene00016838 | WBGene00011556 |
| WBGene00016845 | WBGene00011558 |
| WBGene00016848 | WBGene00011559 |
| WBGene00016862 | WBGene00011561 |
| WBGene00016871 | WBGene00011571 |
| WBGene00016879 | WBGene00011573 |
| WBGene00016893 | WBGene00011586 |
| WBGene00016906 | WBGene00011587 |
| WBGene00016918 | WBGene00011597 |
| WBGene00016920 | WBGene00011604 |
| WBGene00016924 | WBGene00011605 |
| WBGene00016944 | WBGene00011634 |
| WBGene00016957 | WBGene00011636 |
| WBGene00016966 | WBGene00011638 |
| WBGene00016971 | WBGene00011647 |
| WBGene00016996 | WBGene00011661 |
| WBGene00017028 | WBGene00011680 |
| WBGene00017037 | WBGene00011681 |

| | |
|---|---|
| WBGene00017051 | WBGene00011687 |
| WBGene00017054 | WBGene00011688 |
| WBGene00017058 | WBGene00011699 |
| WBGene00017064 | WBGene00011718 |
| WBGene00017065 | WBGene00011722 |
| WBGene00017074 | WBGene00011730 |
| WBGene00017079 | WBGene00011732 |
| WBGene00017085 | WBGene00011734 |
| WBGene00017101 | WBGene00011742 |
| WBGene00017132 | WBGene00011743 |
| WBGene00017137 | WBGene00011746 |
| WBGene00017138 | WBGene00011753 |
| WBGene00017147 | WBGene00011758 |
| WBGene00017158 | WBGene00011767 |
| WBGene00017164 | WBGene00011771 |
| WBGene00017177 | WBGene00011805 |
| WBGene00017210 | WBGene00011807 |
| WBGene00017237 | WBGene00011827 |
| WBGene00017239 | WBGene00011834 |
| WBGene00017240 | WBGene00011839 |
| WBGene00017241 | WBGene00011856 |
| WBGene00017245 | WBGene00011857 |
| WBGene00017261 | WBGene00011858 |
| WBGene00017268 | WBGene00011859 |
| WBGene00017269 | WBGene00011869 |
| WBGene00017273 | WBGene00011879 |
| WBGene00017280 | WBGene00011880 |
| WBGene00017282 | WBGene00011897 |
| WBGene00017298 | WBGene00011898 |
| WBGene00017300 | WBGene00011904 |
| WBGene00017321 | WBGene00011911 |
| WBGene00017336 | WBGene00011917 |
| WBGene00017344 | WBGene00011939 |
| WBGene00017362 | WBGene00011941 |
| WBGene00017373 | WBGene00011944 |
| WBGene00017376 | WBGene00011945 |
| WBGene00017380 | WBGene00011955 |
| WBGene00017436 | WBGene00011964 |
| WBGene00017513 | WBGene00011966 |
| WBGene00017530 | WBGene00011967 |
| WBGene00017532 | WBGene00011969 |
| WBGene00017542 | WBGene00011975 |
| WBGene00017574 | WBGene00011980 |
| WBGene00017578 | WBGene00011986 |
| WBGene00017583 | WBGene00011995 |
| WBGene00017597 | WBGene00011997 |
| WBGene00017600 | WBGene00012015 |
| WBGene00017607 | WBGene00012018 |
| WBGene00017611 | WBGene00012019 |
| WBGene00017613 | WBGene00012020 |
| WBGene00017621 | WBGene00012029 |
| WBGene00017646 | WBGene00012030 |
| WBGene00017653 | WBGene00012037 |
| WBGene00017657 | WBGene00012104 |
| WBGene00017683 | WBGene00012110 |
| WBGene00017692 | WBGene00012113 |
| WBGene00017724 | WBGene00012117 |
| WBGene00017726 | WBGene00012122 |
| WBGene00017730 | WBGene00012124 |
| WBGene00017748 | WBGene00012128 |
| WBGene00017749 | WBGene00012139 |
| WBGene00017763 | WBGene00012140 |
| WBGene00017766 | WBGene00012156 |
| WBGene00017768 | WBGene00012158 |
| WBGene00017797 | WBGene00012166 |
| WBGene00017813 | WBGene00012185 |
| WBGene00017818 | WBGene00012187 |
| WBGene00017825 | WBGene00012204 |

| | |
|---|---|
| WBGene00017826 | WBGene00012206 |
| WBGene00017827 | WBGene00012208 |
| WBGene00017831 | WBGene00012221 |
| WBGene00017837 | WBGene00012226 |
| WBGene00017886 | WBGene00012239 |
| WBGene00017908 | WBGene00012244 |
| WBGene00017919 | WBGene00012253 |
| WBGene00017938 | WBGene00012258 |
| WBGene00017939 | WBGene00012274 |
| WBGene00017945 | WBGene00012289 |
| WBGene00017951 | WBGene00012319 |
| WBGene00017974 | WBGene00012326 |
| WBGene00017978 | WBGene00012342 |
| WBGene00017989 | WBGene00012352 |
| WBGene00017997 | WBGene00012362 |
| WBGene00017999 | WBGene00012364 |
| WBGene00018013 | WBGene00012366 |
| WBGene00018037 | WBGene00012382 |
| WBGene00018044 | WBGene00012383 |
| WBGene00018056 | WBGene00012386 |
| WBGene00018060 | WBGene00012393 |
| WBGene00018075 | WBGene00012428 |
| WBGene00018076 | WBGene00012431 |
| WBGene00018088 | WBGene00012434 |
| WBGene00018137 | WBGene00012435 |
| WBGene00018138 | WBGene00012439 |
| WBGene00018150 | WBGene00012452 |
| WBGene00018156 | WBGene00012466 |
| WBGene00018184 | WBGene00012468 |
| WBGene00018193 | WBGene00012478 |
| WBGene00018214 | WBGene00012522 |
| WBGene00018249 | WBGene00012528 |
| WBGene00018258 | WBGene00012530 |
| WBGene00018262 | WBGene00012543 |
| WBGene00018303 | WBGene00012544 |
| WBGene00018340 | WBGene00012548 |
| WBGene00018366 | WBGene00012551 |
| WBGene00018367 | WBGene00012555 |
| WBGene00018369 | WBGene00012556 |
| WBGene00018370 | WBGene00012557 |
| WBGene00018384 | WBGene00012558 |
| WBGene00018407 | WBGene00012562 |
| WBGene00018408 | WBGene00012584 |
| WBGene00018409 | WBGene00012585 |
| WBGene00018417 | WBGene00012600 |
| WBGene00018427 | WBGene00012602 |
| WBGene00018483 | WBGene00012610 |
| WBGene00018497 | WBGene00012615 |
| WBGene00018511 | WBGene00012641 |
| WBGene00018520 | WBGene00012643 |
| WBGene00018563 | WBGene00012645 |
| WBGene00018569 | WBGene00012646 |
| WBGene00018599 | WBGene00012650 |
| WBGene00018605 | WBGene00012658 |
| WBGene00018613 | WBGene00012660 |
| WBGene00018637 | WBGene00012664 |
| WBGene00018643 | WBGene00012666 |
| WBGene00018672 | WBGene00012668 |
| WBGene00018678 | WBGene00012692 |
| WBGene00018679 | WBGene00012694 |
| WBGene00018704 | WBGene00012697 |
| WBGene00018714 | WBGene00012700 |
| WBGene00018721 | WBGene00012712 |
| WBGene00018769 | WBGene00012713 |
| WBGene00018777 | WBGene00012714 |
| WBGene00018781 | WBGene00012716 |
| WBGene00018785 | WBGene00012738 |
| WBGene00018788 | WBGene00012739 |

WBGene00018793
WBGene00018794
WBGene00018797
WBGene00018806
WBGene00018814
WBGene00018830
WBGene00018831
WBGene00018833
WBGene00018853
WBGene00018854
WBGene00018857
WBGene00018886
WBGene00018902
WBGene00018923
WBGene00018955
WBGene00018966
WBGene00018995
WBGene00018998
WBGene00019004
WBGene00019009
WBGene00019022
WBGene00019026
WBGene00019064
WBGene00019068
WBGene00019075
WBGene00019076
WBGene00019087
WBGene00019090
WBGene00019092
WBGene00019116
WBGene00019120
WBGene00019121
WBGene00019125
WBGene00019130
WBGene00019156
WBGene00019164
WBGene00019170
WBGene00019180
WBGene00019182
WBGene00019187
WBGene00019199
WBGene00019218
WBGene00019245
WBGene00019247
WBGene00019274
WBGene00019275
WBGene00019289
WBGene00019324
WBGene00019347
WBGene00019362
WBGene00019368
WBGene00019400
WBGene00019410
WBGene00019427
WBGene00019432
WBGene00019460
WBGene00019481
WBGene00019513
WBGene00019544
WBGene00019546
WBGene00019554
WBGene00019563
WBGene00019592
WBGene00019600
WBGene00019607
WBGene00019608
WBGene00019627
WBGene00019629

WBGene00012758
WBGene00012759
WBGene00012765
WBGene00012766
WBGene00012782
WBGene00012783
WBGene00012794
WBGene00012812
WBGene00012813
WBGene00012819
WBGene00012825
WBGene00012840
WBGene00012853
WBGene00012860
WBGene00012868
WBGene00012875
WBGene00012888
WBGene00012891
WBGene00012894
WBGene00012895
WBGene00012897
WBGene00012907
WBGene00012926
WBGene00012930
WBGene00012938
WBGene00012964
WBGene00012973
WBGene00012980
WBGene00012983
WBGene00012988
WBGene00012990
WBGene00012996
WBGene00013001
WBGene00013002
WBGene00013011
WBGene00013014
WBGene00013018
WBGene00013023
WBGene00013029
WBGene00013038
WBGene00013077
WBGene00013081
WBGene00013093
WBGene00013094
WBGene00013095
WBGene00013096
WBGene00013123
WBGene00013132
WBGene00013136
WBGene00013144
WBGene00013150
WBGene00013151
WBGene00013160
WBGene00013161
WBGene00013168
WBGene00013178
WBGene00013181
WBGene00013188
WBGene00013189
WBGene00013200
WBGene00013202
WBGene00013209
WBGene00013210
WBGene00013213
WBGene00013223
WBGene00013225
WBGene00013228
WBGene00013235

WBGene00019643
WBGene00019650
WBGene00019655
WBGene00019657
WBGene00019664
WBGene00019684
WBGene00019692
WBGene00019711
WBGene00019715
WBGene00019717
WBGene00019720
WBGene00019728
WBGene00019733
WBGene00019744
WBGene00019749
WBGene00019770
WBGene00019774
WBGene00019777
WBGene00019781
WBGene00019782
WBGene00019808
WBGene00019818
WBGene00019820
WBGene00019822
WBGene00019830
WBGene00019842
WBGene00019847
WBGene00019862
WBGene00019864
WBGene00019867
WBGene00019881
WBGene00019902
WBGene00019907
WBGene00019914
WBGene00019915
WBGene00019918
WBGene00019938
WBGene00019959
WBGene00019965
WBGene00019993
WBGene00019998
WBGene00020012
WBGene00020026
WBGene00020051
WBGene00020058
WBGene00020068
WBGene00020090
WBGene00020094
WBGene00020115
WBGene00020152
WBGene00020168
WBGene00020172
WBGene00020199
WBGene00020207
WBGene00020208
WBGene00020230
WBGene00020264
WBGene00020273
WBGene00020298
WBGene00020311
WBGene00020320
WBGene00020334
WBGene00020341
WBGene00020343
WBGene00020376
WBGene00020390
WBGene00020397
WBGene00020403

WBGene00013237
WBGene00013238
WBGene00013239
WBGene00013240
WBGene00013241
WBGene00013253
WBGene00013259
WBGene00013270
WBGene00013289
WBGene00013307
WBGene00013308
WBGene00013321
WBGene00013323
WBGene00013325
WBGene00013339
WBGene00013340
WBGene00013343
WBGene00013347
WBGene00013354
WBGene00013355
WBGene00013358
WBGene00013360
WBGene00013361
WBGene00013366
WBGene00013373
WBGene00013378
WBGene00013406
WBGene00013407
WBGene00013422
WBGene00013434
WBGene00013435
WBGene00013442
WBGene00013446
WBGene00013465
WBGene00013481
WBGene00013520
WBGene00013529
WBGene00013544
WBGene00013545
WBGene00013550
WBGene00013558
WBGene00013567
WBGene00013576
WBGene00013580
WBGene00013585
WBGene00013595
WBGene00013603
WBGene00013605
WBGene00013606
WBGene00013633
WBGene00013647
WBGene00013666
WBGene00013667
WBGene00013671
WBGene00013677
WBGene00013680
WBGene00013682
WBGene00013683
WBGene00013689
WBGene00013690
WBGene00013694
WBGene00013695
WBGene00013702
WBGene00013703
WBGene00013704
WBGene00013708
WBGene00013718
WBGene00013719

WBGene00020407
WBGene00020408
WBGene00020436
WBGene00020442
WBGene00020443
WBGene00020461
WBGene00020463
WBGene00020466
WBGene00020475
WBGene00020480
WBGene00020491
WBGene00020498
WBGene00020508
WBGene00020509
WBGene00020549
WBGene00020557
WBGene00020558
WBGene00020563
WBGene00020585
WBGene00020588
WBGene00020589
WBGene00020600
WBGene00020601
WBGene00020630
WBGene00020633
WBGene00020661
WBGene00020674
WBGene00020705
WBGene00020715
WBGene00020725
WBGene00020735
WBGene00020742
WBGene00020756
WBGene00020763
WBGene00020796
WBGene00020798
WBGene00020811
WBGene00020819
WBGene00020842
WBGene00020843
WBGene00020847
WBGene00020860
WBGene00020866
WBGene00020867
WBGene00020886
WBGene00020905
WBGene00020910
WBGene00020932
WBGene00020939
WBGene00020955
WBGene00020964
WBGene00021000
WBGene00021005
WBGene00021020
WBGene00021022
WBGene00021038
WBGene00021051
WBGene00021052
WBGene00021056
WBGene00021057
WBGene00021058
WBGene00021059
WBGene00021061
WBGene00021082
WBGene00021103
WBGene00021113
WBGene00021123
WBGene00021155
WBGene00013723
WBGene00013724
WBGene00013732
WBGene00013735
WBGene00013740
WBGene00013742
WBGene00013765
WBGene00013790
WBGene00013803
WBGene00013805
WBGene00013807
WBGene00013808
WBGene00013847
WBGene00013867
WBGene00013880
WBGene00013881
WBGene00013882
WBGene00013884
WBGene00013887
WBGene00013894
WBGene00013898
WBGene00013899
WBGene00013919
WBGene00013921
WBGene00013934
WBGene00013964
WBGene00013983
WBGene00014006
WBGene00014011
WBGene00014015
WBGene00014017
WBGene00014019
WBGene00014022
WBGene00014036
WBGene00014053
WBGene00014054
WBGene00014057
WBGene00014086
WBGene00014088
WBGene00014108
WBGene00014112
WBGene00014117
WBGene00014119
WBGene00014125
WBGene00014140
WBGene00014204
WBGene00014207
WBGene00014208
WBGene00014209
WBGene00014214
WBGene00014222
WBGene00014232
WBGene00014234
WBGene00014243
WBGene00014249
WBGene00014251
WBGene00014252
WBGene00014253
WBGene00014256
WBGene00014793
WBGene00014826
WBGene00015009
WBGene00015025
WBGene00015075
WBGene00015083
WBGene00015088
WBGene00015099
WBGene00015101

| | |
|---|---|
| WBGene00021161 | WBGene00015102 |
| WBGene00021200 | WBGene00015116 |
| WBGene00021205 | WBGene00015126 |
| WBGene00021237 | WBGene00015131 |
| WBGene00021238 | WBGene00015133 |
| WBGene00021246 | WBGene00015146 |
| WBGene00021249 | WBGene00015150 |
| WBGene00021260 | WBGene00015159 |
| WBGene00021263 | WBGene00015161 |
| WBGene00021276 | WBGene00015162 |
| WBGene00021305 | WBGene00015169 |
| WBGene00021311 | WBGene00015173 |
| WBGene00021312 | WBGene00015181 |
| WBGene00021313 | WBGene00015182 |
| WBGene00021320 | WBGene00015184 |
| WBGene00021322 | WBGene00015185 |
| WBGene00021323 | WBGene00015186 |
| WBGene00021324 | WBGene00015232 |
| WBGene00021327 | WBGene00015237 |
| WBGene00021346 | WBGene00015248 |
| WBGene00021365 | WBGene00015249 |
| WBGene00021370 | WBGene00015250 |
| WBGene00021372 | WBGene00015251 |
| WBGene00021412 | WBGene00015280 |
| WBGene00021423 | WBGene00015283 |
| WBGene00021442 | WBGene00015296 |
| WBGene00021443 | WBGene00015297 |
| WBGene00021444 | WBGene00015301 |
| WBGene00021482 | WBGene00015312 |
| WBGene00021508 | WBGene00015328 |
| WBGene00021509 | WBGene00015340 |
| WBGene00021510 | WBGene00015346 |
| WBGene00021541 | WBGene00015350 |
| WBGene00021546 | WBGene00015355 |
| WBGene00021549 | WBGene00015356 |
| WBGene00021552 | WBGene00015404 |
| WBGene00021555 | WBGene00015405 |
| WBGene00021563 | WBGene00015413 |
| WBGene00021605 | WBGene00015461 |
| WBGene00021613 | WBGene00015468 |
| WBGene00021626 | WBGene00015476 |
| WBGene00021629 | WBGene00015480 |
| WBGene00021634 | WBGene00015486 |
| WBGene00021637 | WBGene00015501 |
| WBGene00021647 | WBGene00015507 |
| WBGene00021648 | WBGene00015509 |
| WBGene00021657 | WBGene00015510 |
| WBGene00021686 | WBGene00015513 |
| WBGene00021687 | WBGene00015524 |
| WBGene00021693 | WBGene00015525 |
| WBGene00021743 | WBGene00015538 |
| WBGene00021751 | WBGene00015540 |
| WBGene00021752 | WBGene00015549 |
| WBGene00021754 | WBGene00015551 |
| WBGene00021757 | WBGene00015552 |
| WBGene00021758 | WBGene00015553 |
| WBGene00021764 | WBGene00015555 |
| WBGene00021765 | WBGene00015565 |
| WBGene00021768 | WBGene00015581 |
| WBGene00021779 | WBGene00015591 |
| WBGene00021783 | WBGene00015633 |
| WBGene00021789 | WBGene00015635 |
| WBGene00021808 | WBGene00015644 |
| WBGene00021813 | WBGene00015650 |
| WBGene00021815 | WBGene00015677 |
| WBGene00021817 | WBGene00015680 |
| WBGene00021827 | WBGene00015687 |
| WBGene00021828 | WBGene00015691 |

WBGene00021831
WBGene00021832
WBGene00021856
WBGene00021869
WBGene00021872
WBGene00021879
WBGene00021880
WBGene00021882
WBGene00021894
WBGene00021895
WBGene00021901
WBGene00021905
WBGene00021906
WBGene00021923
WBGene00021943
WBGene00021948
WBGene00021977
WBGene00021984
WBGene00021985
WBGene00022014
WBGene00022021
WBGene00022029
WBGene00022034
WBGene00022036
WBGene00022051
WBGene00022052
WBGene00022055
WBGene00022073
WBGene00022090
WBGene00022092
WBGene00022093
WBGene00022099
WBGene00022104
WBGene00022113
WBGene00022118
WBGene00022121
WBGene00022124
WBGene00022134
WBGene00022138
WBGene00022141
WBGene00022149
WBGene00022150
WBGene00022153
WBGene00022167
WBGene00022168
WBGene00022173
WBGene00022176
WBGene00022182
WBGene00022191
WBGene00022192
WBGene00022193
WBGene00022200
WBGene00022230
WBGene00022232
WBGene00022244
WBGene00022246
WBGene00022269
WBGene00022277
WBGene00022281
WBGene00022296
WBGene00022300
WBGene00022351
WBGene00022354
WBGene00022363
WBGene00022367
WBGene00022369
WBGene00022372
WBGene00022388
WBGene00015698
WBGene00015745
WBGene00015746
WBGene00015752
WBGene00015755
WBGene00015756
WBGene00015776
WBGene00015789
WBGene00015800
WBGene00015801
WBGene00015803
WBGene00015823
WBGene00015868
WBGene00015907
WBGene00015922
WBGene00015927
WBGene00015928
WBGene00015942
WBGene00015954
WBGene00015971
WBGene00015976
WBGene00015978
WBGene00015981
WBGene00016014
WBGene00016017
WBGene00016027
WBGene00016033
WBGene00016048
WBGene00016052
WBGene00016055
WBGene00016057
WBGene00016060
WBGene00016061
WBGene00016062
WBGene00016066
WBGene00016068
WBGene00016101
WBGene00016112
WBGene00016113
WBGene00016115
WBGene00016116
WBGene00016118
WBGene00016120
WBGene00016121
WBGene00016128
WBGene00016131
WBGene00016139
WBGene00016144
WBGene00016150
WBGene00016154
WBGene00016158
WBGene00016161
WBGene00016162
WBGene00016166
WBGene00016171
WBGene00016178
WBGene00016194
WBGene00016203
WBGene00016234
WBGene00016238
WBGene00016244
WBGene00016263
WBGene00016278
WBGene00016291
WBGene00016311
WBGene00016316
WBGene00016321
WBGene00016323

WBGene00022391
WBGene00022399
WBGene00022413
WBGene00022447
WBGene00022450
WBGene00022458
WBGene00022460
WBGene00022466
WBGene00022470
WBGene00022471
WBGene00022493
WBGene00022505
WBGene00022515
WBGene00022531
WBGene00022569
WBGene00022610
WBGene00022615
WBGene00022620
WBGene00022633
WBGene00022634
WBGene00022644
WBGene00022660
WBGene00022664
WBGene00022667
WBGene00022673
WBGene00022694
WBGene00022697
WBGene00022742
WBGene00022762
WBGene00022774
WBGene00022775
WBGene00022785
WBGene00022802
WBGene00022803
WBGene00022805
WBGene00022812
WBGene00022821
WBGene00022836
WBGene00022847
WBGene00022849
WBGene00022868
WBGene00022887
WBGene00023404
WBGene00023407
WBGene00023414
WBGene00023419
WBGene00023421
WBGene00023487
WBGene00023490
WBGene00023497
WBGene00023498
WBGene00023520
WBGene00043051
WBGene00043056
WBGene00043147
WBGene00043156
WBGene00044001
WBGene00044029
WBGene00044035
WBGene00044049
WBGene00044067
WBGene00044068
WBGene00044080
WBGene00044134
WBGene00044143
WBGene00044176
WBGene00044200
WBGene00044205
WBGene00016330
WBGene00016333
WBGene00016351
WBGene00016357
WBGene00016360
WBGene00016361
WBGene00016374
WBGene00016376
WBGene00016378
WBGene00016381
WBGene00016384
WBGene00016391
WBGene00016392
WBGene00016393
WBGene00016395
WBGene00016396
WBGene00016400
WBGene00016407
WBGene00016411
WBGene00016416
WBGene00016432
WBGene00016435
WBGene00016438
WBGene00016440
WBGene00016442
WBGene00016447
WBGene00016469
WBGene00016501
WBGene00016564
WBGene00016573
WBGene00016582
WBGene00016583
WBGene00016588
WBGene00016589
WBGene00016592
WBGene00016594
WBGene00016601
WBGene00016602
WBGene00016606
WBGene00016608
WBGene00016610
WBGene00016614
WBGene00016622
WBGene00016627
WBGene00016629
WBGene00016635
WBGene00016636
WBGene00016638
WBGene00016641
WBGene00016650
WBGene00016722
WBGene00016746
WBGene00016781
WBGene00016797
WBGene00016808
WBGene00016811
WBGene00016837
WBGene00016844
WBGene00016866
WBGene00016867
WBGene00016868
WBGene00016888
WBGene00016898
WBGene00016902
WBGene00016905
WBGene00016913
WBGene00016915
WBGene00016937

WBGene00044210
WBGene00044260
WBGene00044280
WBGene00044310
WBGene00044319
WBGene00044325
WBGene00044329
WBGene00044345
WBGene00044376
WBGene00044420
WBGene00044431
WBGene00044438
WBGene00044440
WBGene00044500
WBGene00044509
WBGene00044514
WBGene00044527
WBGene00044535
WBGene00044554
WBGene00044556
WBGene00044603
WBGene00044605
WBGene00044608
WBGene00044648
WBGene00044698
WBGene00044745
WBGene00044772
WBGene00044773
WBGene00044788
WBGene00044922
WBGene00044924
WBGene00045038
WBGene00045078
WBGene00045208
WBGene00045242
WBGene00045271
WBGene00045305
WBGene00045357
WBGene00045474
WBGene00045495
WBGene00045517
WBGene00077439
WBGene00077757
WBGene00086566
WBGene00164985
WBGene00185078
WBGene00185086
WBGene00189949
WBGene00194649
WBGene00194703
WBGene00194706
WBGene00195093
WBGene00195189
WBGene00206362
WBGene00206382
WBGene00219425
WBGene00235310

WBGene00016943
WBGene00016961
WBGene00016977
WBGene00016983
WBGene00016990
WBGene00017011
WBGene00017016
WBGene00017025
WBGene00017031
WBGene00017044
WBGene00017045
WBGene00017053
WBGene00017062
WBGene00017066
WBGene00017070
WBGene00017080
WBGene00017098
WBGene00017127
WBGene00017128
WBGene00017159
WBGene00017161
WBGene00017162
WBGene00017165
WBGene00017195
WBGene00017201
WBGene00017238
WBGene00017275
WBGene00017284
WBGene00017285
WBGene00017286
WBGene00017292
WBGene00017303
WBGene00017304
WBGene00017313
WBGene00017316
WBGene00017317
WBGene00017319
WBGene00017328
WBGene00017342
WBGene00017347
WBGene00017349
WBGene00017355
WBGene00017356
WBGene00017358
WBGene00017369
WBGene00017374
WBGene00017423
WBGene00017429
WBGene00017435
WBGene00017440
WBGene00017445
WBGene00017446
WBGene00017464
WBGene00017472
WBGene00017482
WBGene00017484
WBGene00017485
WBGene00017546
WBGene00017548
WBGene00017549
WBGene00017591
WBGene00017605
WBGene00017620
WBGene00017640
WBGene00017641
WBGene00017642
WBGene00017643
WBGene00017644

WBGene00017658
WBGene00017659
WBGene00017663
WBGene00017671
WBGene00017675
WBGene00017676
WBGene00017687
WBGene00017691
WBGene00017699
WBGene00017719
WBGene00017727
WBGene00017733
WBGene00017734
WBGene00017742
WBGene00017746
WBGene00017756
WBGene00017774
WBGene00017775
WBGene00017777
WBGene00017799
WBGene00017801
WBGene00017828
WBGene00017850
WBGene00017851
WBGene00017874
WBGene00017881
WBGene00017887
WBGene00017889
WBGene00017892
WBGene00017895
WBGene00017898
WBGene00017900
WBGene00017925
WBGene00017926
WBGene00017929
WBGene00017931
WBGene00017934
WBGene00017961
WBGene00017967
WBGene00017985
WBGene00017986
WBGene00017988
WBGene00017993
WBGene00017996
WBGene00018007
WBGene00018011
WBGene00018017
WBGene00018024
WBGene00018045
WBGene00018052
WBGene00018064
WBGene00018071
WBGene00018072
WBGene00018090
WBGene00018112
WBGene00018117
WBGene00018149
WBGene00018154
WBGene00018159
WBGene00018160
WBGene00018161
WBGene00018164
WBGene00018181
WBGene00018187
WBGene00018195
WBGene00018216
WBGene00018217
WBGene00018221

WBGene00018230
WBGene00018237
WBGene00018238
WBGene00018257
WBGene00018269
WBGene00018270
WBGene00018271
WBGene00018275
WBGene00018286
WBGene00018314
WBGene00018317
WBGene00018319
WBGene00018342
WBGene00018350
WBGene00018361
WBGene00018362
WBGene00018364
WBGene00018371
WBGene00018376
WBGene00018391
WBGene00018393
WBGene00018399
WBGene00018420
WBGene00018424
WBGene00018431
WBGene00018462
WBGene00018465
WBGene00018467
WBGene00018468
WBGene00018474
WBGene00018481
WBGene00018508
WBGene00018516
WBGene00018517
WBGene00018519
WBGene00018564
WBGene00018585
WBGene00018602
WBGene00018604
WBGene00018609
WBGene00018610
WBGene00018611
WBGene00018612
WBGene00018625
WBGene00018636
WBGene00018656
WBGene00018674
WBGene00018681
WBGene00018731
WBGene00018735
WBGene00018743
WBGene00018757
WBGene00018758
WBGene00018762
WBGene00018765
WBGene00018776
WBGene00018778
WBGene00018782
WBGene00018784
WBGene00018803
WBGene00018812
WBGene00018819
WBGene00018837
WBGene00018864
WBGene00018872
WBGene00018890
WBGene00018892
WBGene00018893

WBGene00018894
WBGene00018898
WBGene00018900
WBGene00018925
WBGene00018932
WBGene00018934
WBGene00018935
WBGene00018948
WBGene00018950
WBGene00018957
WBGene00018959
WBGene00018960
WBGene00018961
WBGene00018963
WBGene00018975
WBGene00019003
WBGene00019008
WBGene00019010
WBGene00019017
WBGene00019018
WBGene00019097
WBGene00019108
WBGene00019124
WBGene00019126
WBGene00019127
WBGene00019157
WBGene00019158
WBGene00019160
WBGene00019163
WBGene00019167
WBGene00019168
WBGene00019188
WBGene00019196
WBGene00019203
WBGene00019205
WBGene00019207
WBGene00019209
WBGene00019217
WBGene00019220
WBGene00019234
WBGene00019237
WBGene00019240
WBGene00019241
WBGene00019244
WBGene00019246
WBGene00019261
WBGene00019263
WBGene00019282
WBGene00019284
WBGene00019295
WBGene00019323
WBGene00019329
WBGene00019332
WBGene00019333
WBGene00019351
WBGene00019353
WBGene00019354
WBGene00019357
WBGene00019361
WBGene00019381
WBGene00019396
WBGene00019399
WBGene00019401
WBGene00019402
WBGene00019403
WBGene00019429
WBGene00019433
WBGene00019478

| |
|---|
| WBGene00019479 |
| WBGene00019480 |
| WBGene00019488 |
| WBGene00019511 |
| WBGene00019522 |
| WBGene00019543 |
| WBGene00019593 |
| WBGene00019597 |
| WBGene00019598 |
| WBGene00019606 |
| WBGene00019609 |
| WBGene00019612 |
| WBGene00019617 |
| WBGene00019619 |
| WBGene00019647 |
| WBGene00019663 |
| WBGene00019666 |
| WBGene00019673 |
| WBGene00019679 |
| WBGene00019682 |
| WBGene00019698 |
| WBGene00019713 |
| WBGene00019716 |
| WBGene00019727 |
| WBGene00019747 |
| WBGene00019762 |
| WBGene00019766 |
| WBGene00019767 |
| WBGene00019771 |
| WBGene00019778 |
| WBGene00019779 |
| WBGene00019786 |
| WBGene00019800 |
| WBGene00019801 |
| WBGene00019803 |
| WBGene00019805 |
| WBGene00019807 |
| WBGene00019811 |
| WBGene00019824 |
| WBGene00019837 |
| WBGene00019838 |
| WBGene00019844 |
| WBGene00019863 |
| WBGene00019872 |
| WBGene00019883 |
| WBGene00019888 |
| WBGene00019937 |
| WBGene00019941 |
| WBGene00019943 |
| WBGene00019944 |
| WBGene00019957 |
| WBGene00019971 |
| WBGene00019978 |
| WBGene00020025 |
| WBGene00020030 |
| WBGene00020035 |
| WBGene00020049 |
| WBGene00020088 |
| WBGene00020091 |
| WBGene00020101 |
| WBGene00020110 |
| WBGene00020111 |
| WBGene00020113 |
| WBGene00020114 |
| WBGene00020126 |
| WBGene00020142 |
| WBGene00020143 |
| WBGene00020146 |

WBGene00020149
WBGene00020155
WBGene00020156
WBGene00020158
WBGene00020166
WBGene00020171
WBGene00020190
WBGene00020192
WBGene00020209
WBGene00020214
WBGene00020215
WBGene00020218
WBGene00020229
WBGene00020234
WBGene00020246
WBGene00020251
WBGene00020259
WBGene00020268
WBGene00020286
WBGene00020287
WBGene00020296
WBGene00020314
WBGene00020316
WBGene00020317
WBGene00020322
WBGene00020340
WBGene00020369
WBGene00020374
WBGene00020375
WBGene00020379
WBGene00020383
WBGene00020389
WBGene00020392
WBGene00020393
WBGene00020399
WBGene00020402
WBGene00020417
WBGene00020426
WBGene00020437
WBGene00020445
WBGene00020447
WBGene00020462
WBGene00020499
WBGene00020501
WBGene00020510
WBGene00020511
WBGene00020516
WBGene00020517
WBGene00020540
WBGene00020561
WBGene00020562
WBGene00020577
WBGene00020581
WBGene00020596
WBGene00020651
WBGene00020652
WBGene00020662
WBGene00020682
WBGene00020684
WBGene00020687
WBGene00020695
WBGene00020700
WBGene00020706
WBGene00020709
WBGene00020710
WBGene00020732
WBGene00020736
WBGene00020738

WBGene00020740
WBGene00020757
WBGene00020765
WBGene00020779
WBGene00020782
WBGene00020784
WBGene00020808
WBGene00020812
WBGene00020838
WBGene00020839
WBGene00020854
WBGene00020894
WBGene00020896
WBGene00020904
WBGene00020909
WBGene00020921
WBGene00020933
WBGene00020962
WBGene00020970
WBGene00020993
WBGene00020994
WBGene00020999
WBGene00021004
WBGene00021009
WBGene00021012
WBGene00021021
WBGene00021024
WBGene00021026
WBGene00021035
WBGene00021036
WBGene00021044
WBGene00021048
WBGene00021063
WBGene00021078
WBGene00021093
WBGene00021097
WBGene00021191
WBGene00021208
WBGene00021210
WBGene00021239
WBGene00021277
WBGene00021286
WBGene00021294
WBGene00021316
WBGene00021325
WBGene00021331
WBGene00021332
WBGene00021334
WBGene00021335
WBGene00021351
WBGene00021355
WBGene00021359
WBGene00021363
WBGene00021392
WBGene00021395
WBGene00021400
WBGene00021401
WBGene00021421
WBGene00021440
WBGene00021441
WBGene00021446
WBGene00021456
WBGene00021467
WBGene00021474
WBGene00021506
WBGene00021507
WBGene00021512
WBGene00021514

WBGene00021534
WBGene00021536
WBGene00021539
WBGene00021543
WBGene00021545
WBGene00021548
WBGene00021551
WBGene00021554
WBGene00021558
WBGene00021559
WBGene00021564
WBGene00021595
WBGene00021614
WBGene00021625
WBGene00021636
WBGene00021644
WBGene00021658
WBGene00021660
WBGene00021675
WBGene00021681
WBGene00021683
WBGene00021689
WBGene00021691
WBGene00021697
WBGene00021698
WBGene00021702
WBGene00021733
WBGene00021735
WBGene00021738
WBGene00021748
WBGene00021749
WBGene00021753
WBGene00021756
WBGene00021763
WBGene00021766
WBGene00021771
WBGene00021784
WBGene00021800
WBGene00021810
WBGene00021811
WBGene00021820
WBGene00021826
WBGene00021829
WBGene00021830
WBGene00021839
WBGene00021841
WBGene00021842
WBGene00021843
WBGene00021844
WBGene00021847
WBGene00021849
WBGene00021852
WBGene00021854
WBGene00021857
WBGene00021868
WBGene00021870
WBGene00021876
WBGene00021900
WBGene00021902
WBGene00021904
WBGene00021910
WBGene00021915
WBGene00021921
WBGene00021922
WBGene00021930
WBGene00021934
WBGene00021936
WBGene00021950

WBGene00021952
WBGene00021959
WBGene00021973
WBGene00021993
WBGene00021996
WBGene00022015
WBGene00022019
WBGene00022025
WBGene00022027
WBGene00022044
WBGene00022045
WBGene00022046
WBGene00022048
WBGene00022053
WBGene00022060
WBGene00022063
WBGene00022072
WBGene00022074
WBGene00022075
WBGene00022077
WBGene00022078
WBGene00022087
WBGene00022101
WBGene00022115
WBGene00022130
WBGene00022144
WBGene00022146
WBGene00022152
WBGene00022178
WBGene00022185
WBGene00022188
WBGene00022194
WBGene00022201
WBGene00022202
WBGene00022203
WBGene00022248
WBGene00022251
WBGene00022254
WBGene00022257
WBGene00022268
WBGene00022271
WBGene00022273
WBGene00022276
WBGene00022278
WBGene00022280
WBGene00022286
WBGene00022297
WBGene00022301
WBGene00022306
WBGene00022310
WBGene00022329
WBGene00022347
WBGene00022348
WBGene00022350
WBGene00022360
WBGene00022361
WBGene00022366
WBGene00022373
WBGene00022380
WBGene00022381
WBGene00022382
WBGene00022389
WBGene00022393
WBGene00022400
WBGene00022401
WBGene00022402
WBGene00022410
WBGene00022420

WBGene00022425
WBGene00022426
WBGene00022434
WBGene00022448
WBGene00022452
WBGene00022453
WBGene00022459
WBGene00022462
WBGene00022463
WBGene00022473
WBGene00022489
WBGene00022492
WBGene00022497
WBGene00022499
WBGene00022516
WBGene00022518
WBGene00022534
WBGene00022579
WBGene00022603
WBGene00022618
WBGene00022642
WBGene00022646
WBGene00022647
WBGene00022649
WBGene00022666
WBGene00022702
WBGene00022704
WBGene00022705
WBGene00022717
WBGene00022719
WBGene00022734
WBGene00022738
WBGene00022745
WBGene00022751
WBGene00022765
WBGene00022769
WBGene00022776
WBGene00022791
WBGene00022798
WBGene00022820
WBGene00022822
WBGene00022830
WBGene00022831
WBGene00022832
WBGene00022835
WBGene00022858
WBGene00022864
WBGene00022865
WBGene00022883
WBGene00023405
WBGene00023408
WBGene00023431
WBGene00023451
WBGene00023486
WBGene00023492
WBGene00043064
WBGene00043097
WBGene00043743
WBGene00044025
WBGene00044065
WBGene00044070
WBGene00044072
WBGene00044079
WBGene00044081
WBGene00044132
WBGene00044177
WBGene00044204
WBGene00044206

WBGene00044212
WBGene00044321
WBGene00044341
WBGene00044350
WBGene00044389
WBGene00044467
WBGene00044638
WBGene00044640
WBGene00044650
WBGene00044651
WBGene00044686
WBGene00044689
WBGene00044724
WBGene00044736
WBGene00044737
WBGene00044743
WBGene00044784
WBGene00044891
WBGene00044921
WBGene00045192
WBGene00045249
WBGene00045255
WBGene00045261
WBGene00045276
WBGene00045387
WBGene00045390
WBGene00045394
WBGene00045434
WBGene00045483
WBGene00050914
WBGene00050936
WBGene00050944
WBGene00077643
WBGene00077701
WBGene00077732
WBGene00077771
WBGene00077773
WBGene00185067
WBGene00194641
WBGene00194674
WBGene00194700
WBGene00194908
WBGene00195063
WBGene00219208
WBGene00235291

**Supplementary Table S7 Single-cell RNA-seq values for subset of genes expressed in hypodermis determined by RAPID**

| gene | gene.id | cell.bin | raw.tpm.estimate | adjusted.tpm.estimate | bootstrap.median.tpm | ci.95p.lb | ci.80p.lb | ci.80p.ub | ci.95p.ub |
|---|---|---|---|---|---|---|---|---|---|
| aat-5 | WBGene00000006 | hyp1_hyp2:330_390 | 13.3 | 0 | 13.3 | 0 | 0 | 28.1 | 42.9 |
| acr-5 | WBGene00000044 | hyp1_hyp2:330_390 | 0 | 0 | 0 | 0 | 0 | 0 | 0 |
| adm-4 | WBGene00000075 | hyp7_C_lineage:270_330 | 5.6 | 3.7 | 5.3 | 1.8 | 2.7 | 8.9 | 10.9 |
| aex-1 | WBGene00000084 | hyp4_to_7_and_P_cells:580_650 | 8.9 | 5.7 | 8.6 | 0.2 | 2.6 | 15.9 | 20.3 |
| cat-1 | WBGene00000295 | Seam_cell:330_390 | 583 | 593.4 | 581.5 | 522.5 | 544.8 | 620.4 | 640.3 |
| cdh-9 | WBGene00000401 | hyp3:450_510 | 1446.2 | 1783.4 | 1414.6 | 1093.3 | 1195.6 | 1642 | 1741.8 |
| cdr-1 | WBGene00000412 | hyp4_hyp5_hyp6:390_450 | 39.2 | 35 | 38.1 | 18.7 | 25 | 52.9 | 62.1 |
| ced-8 | WBGene00000422 | hyp1_hyp2:390_450 | 132.7 | 112 | 122 | 0 | 29.2 | 233 | 292.6 |
| ckb-2 | WBGene00000512 | hyp3:330_390 | 81.9 | 72.5 | 79.4 | 14.5 | 34.8 | 130.6 | 155.1 |
| cnc-5 | WBGene00000559 | hyp1_hyp2:330_390 | 0 | 0 | 0 | 0 | 0 | 0 | 0 |
| cng-3 | WBGene00000563 | hyp7_AB_lineage:390_450 | 1.2 | 0.3 | 1.2 | 0 | 0 | 2.6 | 3.7 |
| col-72 | WBGene00000648 | Seam_cell:gt_650 | 17.1 | 0 | 16.7 | 0 | 0 | 34.3 | 50.7 |
| col-78 | WBGene00000654 | hyp7_C_lineage:270_330 | 0.5 | 0.1 | 0.4 | 0 | 0 | 1.2 | 1.6 |
| col-99 | WBGene00000674 | hyp7_C_lineage:270_330 | 35.7 | 34.6 | 35.3 | 26.4 | 29.1 | 41.9 | 44.9 |
| cyn-8 | WBGene00000884 | hyp3:330_390 | 563.4 | 616.1 | 555.9 | 367.2 | 424.1 | 690.5 | 772.1 |
| deg-1 | WBGene00000950 | hyp7_C_lineage:270_330 | 11 | 9.1 | 10.7 | 5.6 | 7.3 | 14.7 | 17.5 |
| deg-3 | WBGene00000951 | hyp3:450_510 | 95.7 | 46.9 | 93.2 | 0 | 0 | 188 | 250.8 |
| dpy-19 | WBGene00001078 | hyp1_hyp2:390_450 | 169.4 | 162.2 | 159.7 | 34.9 | 69.8 | 273.4 | 327.5 |
| egl-46 | WBGene00001210 | hyp1_hyp2:330_390 | 97.9 | 74.2 | 95.2 | 21.9 | 44.5 | 156.6 | 197.8 |
| frh-1 | WBGene00001486 | hyp4_hyp5_hyp6:270_330 | 20.5 | 16.9 | 20.2 | 8.1 | 11.4 | 30.2 | 35.1 |
| gcy-4 | WBGene00001531 | hyp1_hyp2:330_390 | 0 | 0 | 0 | 0 | 0 | 0 | 0 |
| gcy-5 | WBGene00001532 | hyp1_hyp2:330_390 | 0 | 0 | 0 | 0 | 0 | 0 | 0 |
| gcy-11 | WBGene00001537 | hyp7_C_lineage:270_330 | 12.9 | 11 | 12.3 | 7.1 | 9 | 16.9 | 19.6 |
| ggr-3 | WBGene00001588 | hyp1_hyp2:330_390 | 0 | 0 | 0 | 0 | 0 | 0 | 0 |
| gly-19 | WBGene00001644 | hyp1_hyp2:330_390 | 0 | 0 | 0 | 0 | 0 | 0 | 0 |
| grl-3 | WBGene00001712 | hyp1_hyp2:330_390 | 0 | 0 | 0 | 0 | 0 | 0 | 0 |
| grl-11 | WBGene00001720 | hyp4_hyp5_hyp6:390_450 | 21.9 | 10.6 | 19.1 | 0 | 0 | 46.9 | 60.5 |
| grl-24 | WBGene00001733 | hyp4_hyp5_hyp6:330_390 | 6.5 | 4.5 | 6 | 1.5 | 2.7 | 10.5 | 12.6 |
| gro-1 | WBGene00001740 | hyp4_hyp5_hyp6:270_330 | 39.8 | 38.4 | 39.4 | 23.3 | 28.2 | 51.3 | 58.8 |
| gst-8 | WBGene00001756 | hyp7_AB_lineage:270_330 | 1.3 | 0.6 | 1.3 | 0 | 0 | 2.6 | 3.3 |
| gst-30 | WBGene00001778 | hyp1_hyp2:330_390 | 0 | 0 | 0 | 0 | 0 | 0 | 0 |
| gst-33 | WBGene00001781 | Seam_cell:580_650 | 4.3 | 1.2 | 4.1 | 0 | 0 | 9.3 | 12.8 |
| hnd-1 | WBGene00001981 | hyp1_hyp2:330_390 | 40.8 | 22.4 | 38.9 | 0 | 1.5 | 82.4 | 106.9 |
| ifa-3 | WBGene00002051 | hyp7_AB_lineage:390_450 | 1953.9 | 2019 | 1951.9 | 1812.9 | 1856.1 | 2036.7 | 2082.4 |
| ins-9 | WBGene00002092 | hyp1_hyp2:330_390 | 0 | 0 | 0 | 0 | 0 | 0 | 0 |

| | | | | | | | | | |
|---|---|---|---|---|---|---|---|---|---|
| klp-6 | WBGene00002218 | hyp4_hyp5_hyp6:270_330 | 2.4 | 1.1 | 2.4 | 0 | 0 | 5 | 6.7 |
| lev-9 | WBGene00002976 | Seam_cell:510_580 | 2.5 | 0.6 | 2.1 | 0 | 0 | 6.1 | 8.4 |
| lin-32 | WBGene00003018 | hyp1_hyp2:450_510 | 384.8 | 119.2 | 346.2 | 0 | 0 | 793.5 | 1103 |
| mom-1 | WBGene00003394 | hyp1_hyp2:510_580 | 18.9 | 8.2 | 18.6 | 0 | 0 | 40.7 | 54.2 |
| nhr-44 | WBGene00003634 | hyp1_hyp2:330_390 | 223.5 | 231.1 | 219.8 | 143.4 | 165.9 | 280 | 312.3 |
| nhr-53 | WBGene00003643 | hyp1_hyp2:330_390 | 13.2 | 9.4 | 12.9 | 2.8 | 5.7 | 22.5 | 29.3 |
| nhr-104 | WBGene00003694 | hyp4_hyp5_hyp6:330_390 | 170.8 | 173.3 | 169.8 | 139.3 | 149.1 | 191.8 | 201.7 |
| nhr-109 | WBGene00003699 | hyp4_hyp5_hyp6:390_450 | 135 | 133.6 | 132.4 | 87.4 | 102.1 | 171.1 | 189.2 |
| nhr-111 | WBGene00003701 | hyp7_AB_lineage:270_330 | 1 | 0.2 | 1 | 0 | 0 | 2.4 | 3.1 |
| nhr-115 | WBGene00003705 | hyp1_hyp2:330_390 | 0 | 0 | 0 | 0 | 0 | 0 | 0 |
| nhx-1 | WBGene00003729 | hyp7_C_lineage:390_450 | 619.8 | 630.7 | 618.9 | 570.6 | 586.2 | 649.6 | 667.1 |
| odr-1 | WBGene00003848 | hyp1_hyp2:330_390 | 0 | 0 | 0 | 0 | 0 | 0 | 0 |
| osm-1 | WBGene00003883 | hyp1_hyp2:510_580 | 17.4 | 3.6 | 17.1 | 0 | 0 | 39.1 | 53.8 |
| osm-9 | WBGene00003889 | hyp1_hyp2:330_390 | 0 | 0 | 0 | 0 | 0 | 0 | 0 |
| pgl-2 | WBGene00003993 | hyp7_AB_lineage:390_450 | 86.7 | 81.2 | 83.9 | 51 | 60.4 | 112.5 | 126.1 |

The 471 genes identified uniquely by RAPID were ranked according to gene.id; the first 50 were examined in the database for single-cell RNA-seq from Packer et al (2019) at https://cello.shinyapps.io/celegans/

We consider reliably detected genes for which the lower 95% confidence interval (ci.95p.lb) is above zero.

Packer, J. S., Q. Zhu, C. Huynh, P. Sivaramakrishnan, E. Preston, H. Dueck, D. Stefanik, K. Tan, C. Trapnell, J. Kim, R. H. Waterston and J. I. Murray (2019). A lineage-resolved molecular atlas of C. elegans embryogenesis at single-cell resolution. Science: eaax1971.
